# Bone Marrow Mesenchymal Stem Cells Therapy for Premature Ovarian Insufficiency: A Systematic Review and Meta-analysis of Preclinical Studies

**DOI:** 10.64898/2026.07.02.736116

**Authors:** Jaime Plane, Francisco Torres, Pilar Vera, David Vantman, Barbara A. Andrews, Juan A. Asenjo, Pablo Caviedes, Anamaría Daza

## Abstract

**Background:** Premature ovarian insufficiency (POI) affects approximately 1% of women under 40 and is characterized by elevated levels of gonadotropins, reduced estradiol, impaired folliculogenesis, and infertility. Bone marrow–derived mesenchymal stem cell (BM-MSC)-based therapy has emerged as a promising regenerative strategy in preclinical POI models. This systematic review and meta-analysis evaluated BM-MSC-based interventions, including cell transplantation and secretome/extracellular vesicle administration, in animal models of POI.

**Methods:** A systematic review and meta-analysis was conducted following PRISMA guidelines. PubMed, Web of Science, Scopus, ScienceDirect, and the Cochrane Library were searched from inception to February 19, 2025. Preclinical studies assessing BM-MSC-based interventions in animal models of POI were included.

**Results:** Thirty-four studies comprising 1,357 animals were included. Compared with controls, BM-MSC-based therapy increased serum estradiol (standardized mean difference [SMD] 3.11; 95% confidence interval [CI] 2.38–3.84) and anti-Müllerian hormone (SMD 1.86; 95% CI 1.03–2.69), while reducing follicle-stimulating hormone (SMD −3.54; 95% CI −4.37 to −2.71) and luteinizing hormone (SMD −3.44; 95% CI −5.17 to −1.70). Follicular counts increased across developmental stages, with fewer atretic follicles. Reproductive outcomes improved, including normal estrous cycles (risk ratio [RR] 7.80; 95% CI 3.15–19.34), pregnancy occurrence (RR 3.72; 95% CI 2.14–6.44), and offspring number (SMD 1.57; 95% CI 1.04–2.09).

**Conclusion:** BM-MSC-based therapy consistently improved hormonal, follicular, and reproductive outcomes in preclinical POI models. More well-designed, standardized, and adequately controlled studies to confirm these findings are warranted.

Systematic review registration: CRD42023449053

## 1. Introduction

Premature ovarian insufficiency (POI) is a clinical syndrome characterized by the loss of ovarian function before the age of 40. The condition is characterized by hypogonadism, elevated gonadotropin levels—primarily luteinizing hormone (LH) and follicle-stimulating hormone (FSH)—and decreased estradiol levels [1]. This condition affects approximately 1% of women of reproductive age and is associated with hormonal imbalances that can lead to amenorrhea and infertility [2]. Additionally, POI increases the risk of long-term complications such as cardiovascular diseases, reduced bone mineral density, and psychological disorders, significantly impacting patients’ quality of life [3].

In this context, mesenchymal stem cell (MSC) therapy, particularly those derived from bone marrow (BM-MSC), has emerged as a promising therapeutic strategy. These cells can differentiate into multiple cell types and secrete factors that promote angiogenesis, cell survival, and tissue regeneration [4]. Recent studies suggest that MSC administration can activate ovarian follicles and improve hormonal profiles in POI patients, representing an innovative intervention with significant therapeutic potential [5].

The etiology of POI is complex and multifactorial, and in many cases, it remains poorly understood. Known factors include genetic abnormalities, autoimmune disorders, and iatrogenic damage associated with cancer treatments such as chemotherapy and radiotherapy [3,6]. Autoimmune diseases, including systemic lupus erythematosus and Hashimoto’s thyroiditis, can trigger immune responses that affect ovarian tissue, contributing to the development of POI [7,8]. This diversity of causal factors underscores the need for a multidisciplinary approach to diagnosis and treatment. On the other hand, conventional therapies such as hormone replacement therapy (HRT) have shown limitations, especially for women with reproductive aspirations. HRT can mitigate menopausal symptoms and reduce the risk of osteoporosis and cardiovascular disease, although it does not restore ovarian function [5,9]. Alternatively, strategies such as oocyte or embryo cryopreservation represent viable options for some women, although their availability and effectiveness remain limited [10,11].

Despite advances in stem cell research for treating POI, current evidence remains fragmented and, at times, contradictory. It is crucial to conduct rigorous clinical studies and meta-analyses to comprehensively evaluate the safety and efficacy of MSCs and their derived products as therapy [12,13]. This review examines preclinical studies that have explored the role of BM-MSCs in POI models, detailing the underlying mechanisms of their action and analyzing their clinical implications [14]. Since POI is a complex condition requiring a comprehensive approach, stem cell research offers new hope for affected women, with the potential to restore ovarian function and improve their quality of life [15,16]. Thus, conducting a systematic review and meta-analysis of the available evidence is crucial to consolidate findings and clarify these fundamental aspects. This approach will facilitate data collection and synthesis to detect patterns and discrepancies, establishing a solid knowledge base to guide future clinical trials and contribute to defining treatment standards. Ultimately, a thorough analysis is essential for advancing toward safe and effective clinical applications, providing new hope to women affected by this condition[17–19].

Therefore, this systematic review and meta-analysis aims to determine the effectiveness of Bone Marrow Mesenchymal Stem Cells Therapy for Premature Ovarian Insufficiency in preclinical studies.

## 2. Methods

The present study was carried out following the Cochrane Collaboration Manual for Systematic Reviews of Intervention Studies [20], and Preferred Reporting Items for Systematic Reviews and Meta-Analysis (PRISMA) [21] statement guidelines (completed PRISMA 2020 checklist is provided as Supplementary Material). The protocol was registered into PROSPERO (ID CRD42023449053) prior to the literature search and data analysis.

### 2.1. Eligibility Criteria

Inclusion in this meta-analysis was restricted to studies that met all the following eligibility criteria: (1) preclinical studies involving *in vivo* models; (2) successful POI model establishment; (3) comparing BM-MSC therapy, either transplantation of mesenchymal stem cells or their secretomes; (4) studies available in English or Spanish; (5) no restriction in follow-up time was considered. In addition, studies were included only if at least one of the outcomes of interest were included. Studies classified as observational, case reports, or non-original research (e.g., reviews, editorials, non-investigative letters, and protocols), as well as *in vitro* studies and clinical trials, were excluded.

### 2.2. Search strategy and data extraction

The search was performed from inception to 19 February 2025 in the following databases: PubMed, Web of Science, Scopus, ScienceDirect and Cochrane Library. The search strategy included the following terms and synonyms: “Premature Ovarian Insufficiency” and “Mesenchymal Stem Cells”. The detailed search strategy for each consulted database is available in Table S1, supplementary material. To identify additional eligible studies, backward and forward snowballing were performed by screening the reference lists of included studies and tracking citing articles.

Two authors (JP and FT) performed the study screening and data extraction independently using Covidence software [22], any conflicts were solved by consensus or with guidance from a third senior author (AD), and the data extraction was validated by AD. The Webplotdigitizer software [23] was utilized to extract data directly from graphs. When any of the above information was unclear, we contacted the authors to provide clarification.

### 2.3. Endpoints and subgroup analyses

The outcomes of interest were: (1) Serum hormone production; (2) Folliculogenesis; (3) Estrous cycle; and (4) Fertility endpoints, such as pregnancy and number of offspring. Subgroup analyses were performed according to the type of BM-MSC therapy: (1) BM-MSCs; (2) BM-MSCs secretome; and (3) BM-MSCs miR-21 exosomes. In studies reporting outcomes at multiple time points, the assessment closest to the most frequently reported follow-up period was included, corresponding to 4 weeks.

### 2.4. Quality assessment

The methodological quality of each included study was assessed using the SYRCLE risk of bias tool [24]. Two reviewers independently assessed the quality of each included study, with disagreements resolved by a third reviewer. Publication bias was evaluated in all outcomes, including more than 10 studies, using Funnel plots, and a qualitative analysis of their symmetry.

### 2.5. Assessment of heterogeneity

Heterogeneity was assessed using I² statistics and the Cochran Q test; p-values<0.10 and I²> 50% were considered significant heterogeneity. The DerSimonian and Laird random-effects model was applied for significant heterogeneity; otherwise, the fixed-effects model was used. Sensitivity analysis was performed by removing each individual study from the outcome assessment. Furthermore, subgroup analyses were performed to investigate the possible factors of heterogeneity such as: (1) animal model; (2) hormone measurement techniques; (3) model establishment method; and (4) route of administration.

### 2.6. Statistical Analysis

All statistical analyses were conducted using Review Manager Web (version 8.2.0). For studies with three or more arms, the BM-MSC therapy arms were combined by calculating a pooled mean and standard deviation using Cochrane’s formula [20] and were entered as a single intervention group in the overall analysis. Standardized Mean Difference (SMD) was adopted for continuous data, and Risk Ratio (RR) was used for categorical outcomes, both with a 95% confidence interval (95% CI). P-values less than 0.05 were considered statistically significant.

## 3. Results

### 3.1. Study selection and baseline characteristics

As illustrated in the PRISMA flow diagram (Figure 1), the initial search yielded 1,423 studies. After duplicate removal and title/abstract screening, 62 studies were retained for full-text assessment, of which 34 met the inclusion criteria and were included in this systematic review and meta-analysis, comprising a total of 1,357 animals.

**Fig. 1.**
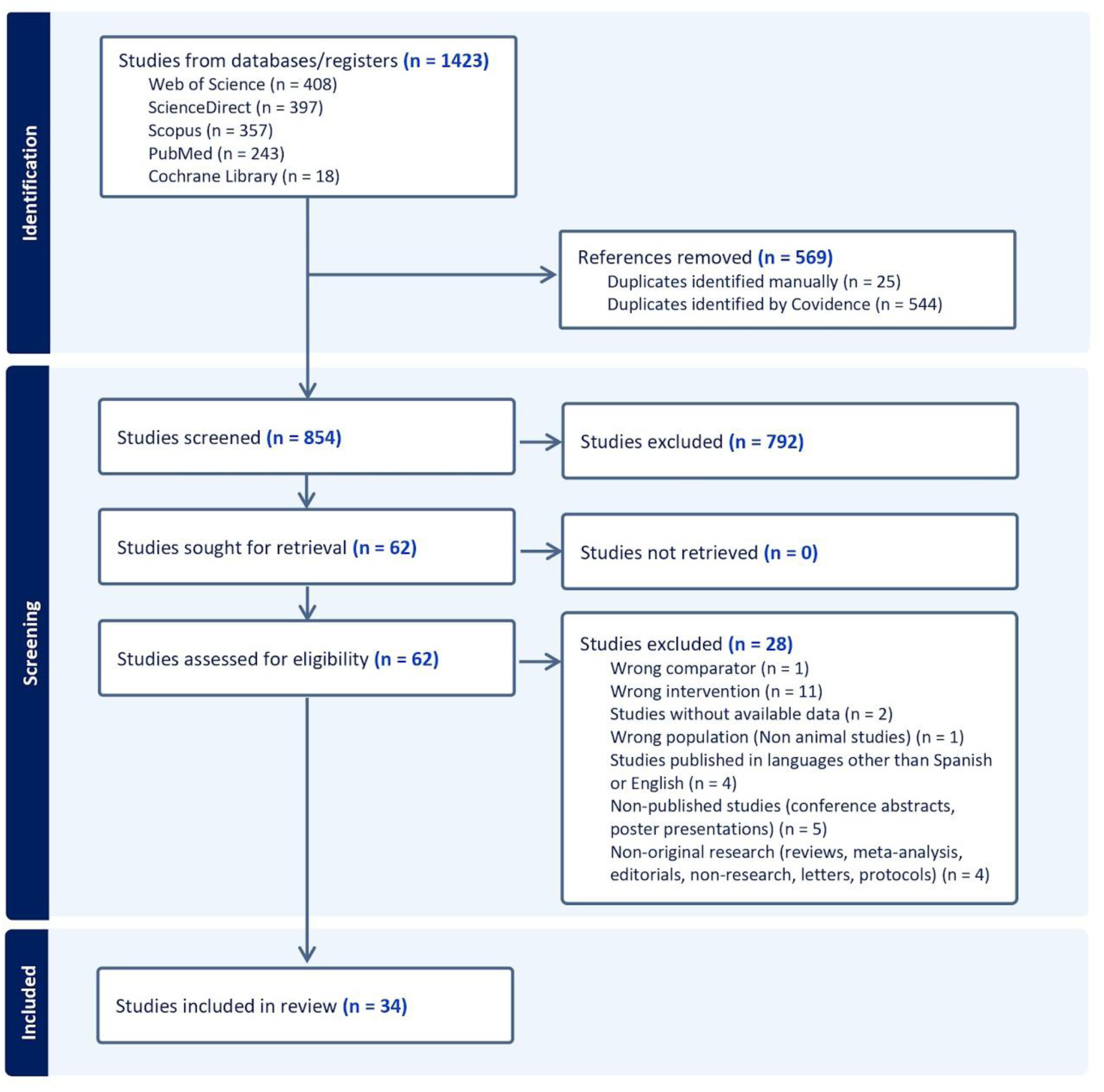
Flow diagram on searching procedure of Web of Science, ScienceDirect, Scopus, PubMed, and Cochrane Library, and the exclusion criteria

The geographical distribution of the included publications was concentrated in China (n=15) and Egypt (n=8), followed by Iran (n=4) and the United States (n=4). Most experiments used rats (n=21), predominantly Sprague-Dawley and Wistar strains, followed by mice (n=12) and a single rabbit study (n=1), providing a diverse representation of the effects of BM-MSC therapy on ovarian function.

Analysis of the POI induction methods showed that the most frequent model was chemotherapy-induced toxicity with cyclophosphamide (n=26). Less common methods included cisplatin administration (n=3), autoimmune induction (n=3), radiotherapy (n=1), and a combination of cyclophosphamide and cisplatin (n=1).

In terms of therapeutic interventions, BM-MSC cell transplantation (n=26) was the predominant approach, followed by BM-MSC secretome (n=12) including exosomes (n=9), extracellular vesicles (n=1) and conditioned medium (n=2). Two studies included a miR-21 exosomes intervention group. Regarding the route of administration, systemic intravenous (IV) delivery was the most frequently used method, accounting for 47.1% (n=16) of the included studies, followed by localised intraovarian (IO) injection (38.2%; n=13) and intraperitoneal (IP) administration (14.7%; n=5).

Our analysis evidenced that dosing varied by intervention type. For cell-based therapies, administered doses ranged from 5×10⁵ to 2×10⁶ cells, with 1×10⁶ being the most frequently used concentration (n=9). For exosome-based therapies, doses were generally standardized between 100 and 150 µg of total protein, with 150 µg being the most represented dose (n=5). Outcome measures across the included studies comprised serum hormone levels (E2, FSH, LH, AMH), estrous-cycle regularity, follicular counts, and fertility endpoints such as pregnancy and litter size (Detailed information is provided in Table 1).

**Table 1.**
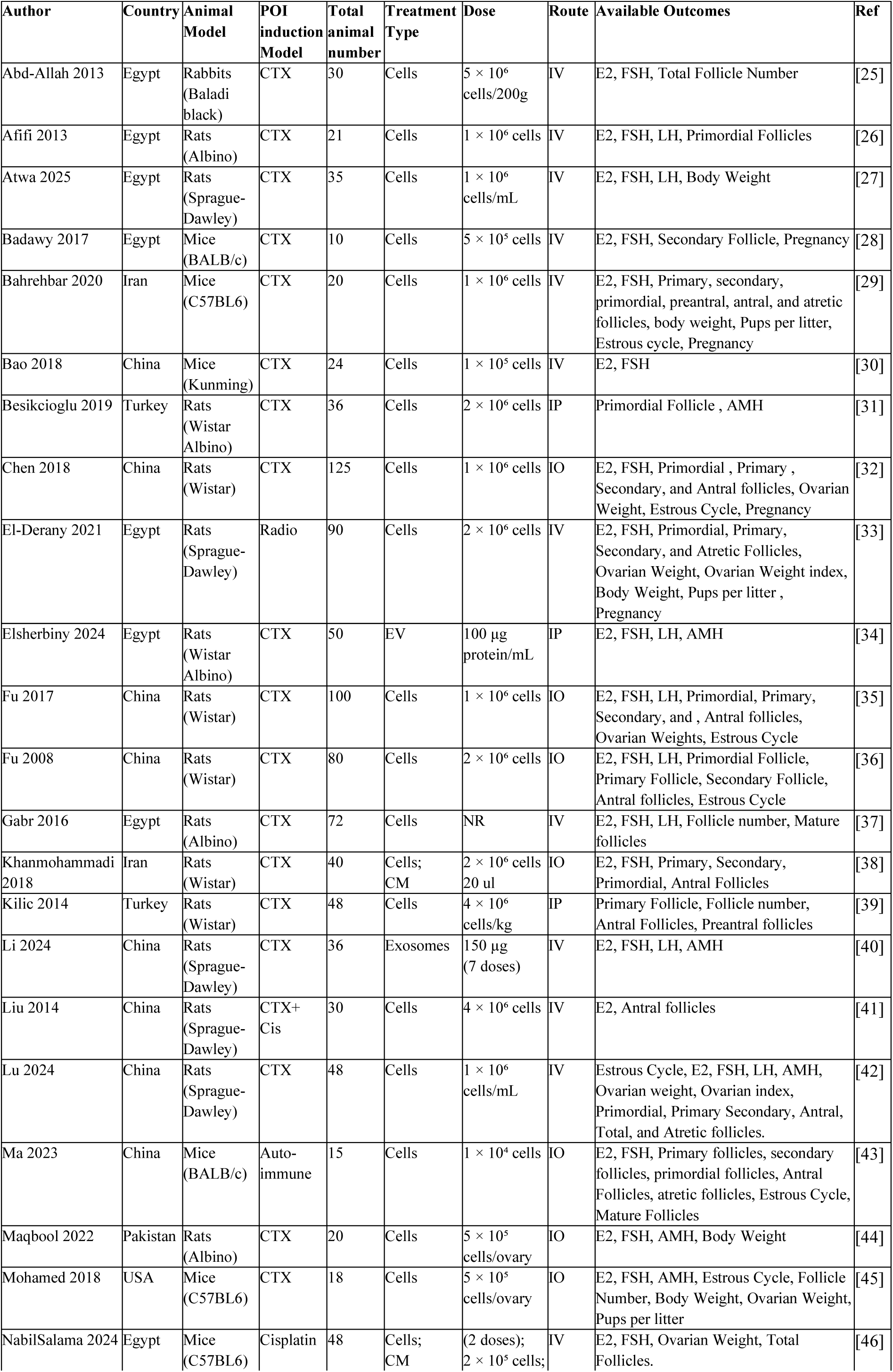
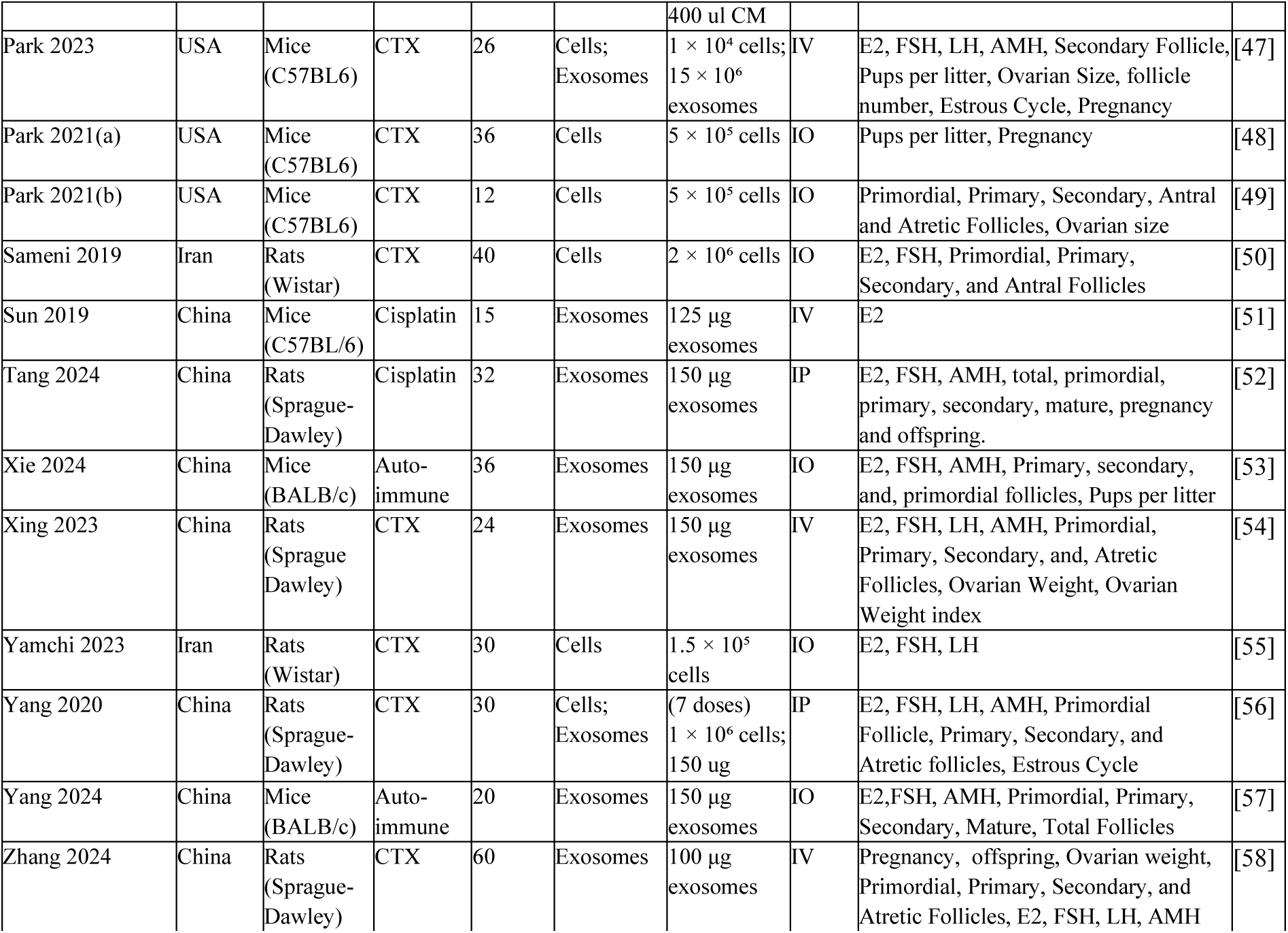
Baseline characteristics of the included studies. Thirty-four studies encompassing 1,357 animals were included. Studies are listed in alphabetical order by the first author. POI: premature ovarian insufficiency; CTX: cyclophosphamide; Cis: cisplatin; Radio: ionizing radiation; CM: conditioned medium; EV: extracellular vesicles; IV: intravenous; IO: intraovarian; IP: intraperitoneal; E2: estradiol; FSH: follicle-stimulating hormone; LH: luteinizing hormone; AMH: anti-Müllerian hormone; NR: not reported. Dose reflects the amount administered per animal unless otherwise specified by body weight or ovary.

| Author | Country | Animal Model | POI induction Model | Total animal number | Treatment Type | Dose | Route | Available Outcomes | Ref |
| --- | --- | --- | --- | --- | --- | --- | --- | --- | --- |
| Abd-Allah 2013 | Egypt | Rabbits (Baladi black) | CTX | 30 | Cells | $5 \times 10^6$ cells/200g | IV | E2, FSH, Total Follicle Number | [25] |
| Afifi 2013 | Egypt | Rats (Albino) | CTX | 21 | Cells | $1 \times 10^6$ cells | IV | E2, FSH, LH, Primordial Follicles | [26] |
| Atwa 2025 | Egypt | Rats (Sprague-Dawley) | CTX | 35 | Cells | $1 \times 10^6$ cells/mL | IV | E2, FSH, LH, Body Weight | [27] |
| Badawy 2017 | Egypt | Mice (BALB/c) | CTX | 10 | Cells | $5 \times 10^5$ cells | IV | E2, FSH, Secondary Follicle, Pregnancy | [28] |
| Bahrebar 2020 | Iran | Mice (C57BL6) | CTX | 20 | Cells | $1 \times 10^6$ cells | IV | E2, FSH, Primary, secondary, primordial, preantral, antral, and atretic follicles, body weight, Pups per litter, Estrous cycle, Pregnancy | [29] |
| Bao 2018 | China | Mice (Kunming) | CTX | 24 | Cells | $1 \times 10^5$ cells | IV | E2, FSH | [30] |
| Besikcioglu 2019 | Turkey | Rats (Wistar Albino) | CTX | 36 | Cells | $2 \times 10^6$ cells | IP | Primordial Follicle , AMH | [31] |
| Chen 2018 | China | Rats (Wistar) | CTX | 125 | Cells | $1 \times 10^6$ cells | IO | E2, FSH, Primordial , Primary , Secondary, and Antral follicles, Ovarian Weight, Estrous Cycle, Pregnancy | [32] |
| El-Derany 2021 | Egypt | Rats (Sprague-Dawley) | Radio | 90 | Cells | $2 \times 10^6$ cells | IV | E2, FSH, Primordial, Primary, Secondary, and Atretic Follicles, Ovarian Weight, Ovarian Weight index, Body Weight, Pups per litter , Pregnancy | [33] |
| Elsherbiny 2024 | Egypt | Rats (Wistar Albino) | CTX | 50 | EV | 100 μg protein/mL | IP | E2, FSH, LH, AMH | [34] |
| Fu 2017 | China | Rats (Wistar) | CTX | 100 | Cells | $1 \times 10^6$ cells | IO | E2, FSH, LH, Primordial, Primary, Secondary, and , Antral follicles, Ovarian Weights, Estrous Cycle | [35] |
| Fu 2008 | China | Rats (Wistar) | CTX | 80 | Cells | $2 \times 10^6$ cells | IO | E2, FSH, LH, Primordial Follicle, Primary Follicle, Secondary Follicle, Antral follicles, Estrous Cycle | [36] |
| Gabr 2016 | Egypt | Rats (Albino) | CTX | 72 | Cells | NR | IV | E2, FSH, LH, Follicle number, Mature follicles | [37] |
| Khanmohammadi 2018 | Iran | Rats (Wistar) | CTX | 40 | Cells; CM | $2 \times 10^6$ cells<br>20 ul | IO | E2, FSH, Primary, Secondary, Primordial, Antral Follicles | [38] |
| Kilic 2014 | Turkey | Rats (Wistar) | CTX | 48 | Cells | $4 \times 10^6$ cells/kg | IP | Primary Follicle, Follicle number, Antral Follicles, Preantral follicles | [39] |
| Li 2024 | China | Rats (Sprague-Dawley) | CTX | 36 | Exosomes | 150 μg (7 doses) | IV | E2, FSH, LH, AMH | [40] |
| Liu 2014 | China | Rats (Sprague-Dawley) | CTX+ Cis | 30 | Cells | $4 \times 10^6$ cells | IV | E2, Antral follicles | [41] |
| Lu 2024 | China | Rats (Sprague-Dawley) | CTX | 48 | Cells | $1 \times 10^6$ cells/mL | IV | Estrous Cycle, E2, FSH, LH, AMH, Ovarian weight, Ovarian index, Primordial, Primary Secondary, Antral, Total, and Atretic follicles. | [42] |
| Ma 2023 | China | Mice (BALB/c) | Auto-immune | 15 | Cells | $1 \times 10^4$ cells | IO | E2, FSH, Primary follicles, secondary follicles, primordial follicles, Antral Follicles, atretic follicles, Estrous Cycle, Mature Follicles | [43] |
| Maqbool 2022 | Pakistan | Rats (Albino) | CTX | 20 | Cells | $5 \times 10^5$ cells/ovary | IO | E2, FSH, AMH, Body Weight | [44] |
| Mohamed 2018 | USA | Mice (C57BL6) | CTX | 18 | Cells | $5 \times 10^5$ cells/ovary | IO | E2, FSH, AMH, Estrous Cycle, Follicle Number, Body Weight, Ovarian Weight, Pups per litter | [45] |
| NabilSalama 2024 | Egypt | Mice (C57BL6) | Cisplatin | 48 | Cells; CM | (2 doses);<br>$2 \times 10^5$ cells; | IV | E2, FSH, Ovarian Weight, Total Follicles. | [46] |
|  |  |  |  |  |  | 400 ul CM |  |  |  |
| Park 2023 | USA | Mice (C57BL/6) | CTX | 26 | Cells; Exosomes | $1 \times 10^4$ cells; $15 \times 10^6$ exosomes | IV | E2, FSH, LH, AMH, Secondary Follicle, Pups per litter, Ovarian Size, follicle number, Estrous Cycle, Pregnancy | [47] |
| Park 2021(a) | USA | Mice (C57BL/6) | CTX | 36 | Cells | $5 \times 10^5$ cells | IO | Pups per litter, Pregnancy | [48] |
| Park 2021(b) | USA | Mice (C57BL/6) | CTX | 12 | Cells | $5 \times 10^5$ cells | IO | Primordial, Primary, Secondary, Antral and Atretic Follicles, Ovarian size | [49] |
| Sameni 2019 | Iran | Rats (Wistar) | CTX | 40 | Cells | $2 \times 10^6$ cells | IO | E2, FSH, Primordial, Primary, Secondary, and Antral Follicles | [50] |
| Sun 2019 | China | Mice (C57BL/6) | Cisplatin | 15 | Exosomes | 125 μg exosomes | IV | E2 | [51] |
| Tang 2024 | China | Rats (Sprague-Dawley) | Cisplatin | 32 | Exosomes | 150 μg exosomes | IP | E2, FSH, AMH, total, primordial, primary, secondary, mature, pregnancy and offspring. | [52] |
| Xie 2024 | China | Mice (BALB/c) | Auto-immune | 36 | Exosomes | 150 μg exosomes | IO | E2, FSH, AMH, Primary, secondary, and, primordial follicles, Pups per litter | [53] |
| Xing 2023 | China | Rats (Sprague Dawley) | CTX | 24 | Exosomes | 150 μg exosomes | IV | E2, FSH, LH, AMH, Primordial, Primary, Secondary, and, Atretic Follicles, Ovarian Weight, Ovarian Weight index | [54] |
| Yamchi 2023 | Iran | Rats (Wistar) | CTX | 30 | Cells | $1.5 \times 10^5$ cells | IO | E2, FSH, LH | [55] |
| Yang 2020 | China | Rats (Sprague-Dawley) | CTX | 30 | Cells; Exosomes | (7 doses) $1 \times 10^6$ cells; 150 μg | IP | E2, FSH, LH, AMH, Primordial Follicle, Primary, Secondary, and Atretic follicles, Estrous Cycle | [56] |
| Yang 2024 | China | Mice (BALB/c) | Auto-immune | 20 | Exosomes | 150 μg exosomes | IO | E2, FSH, AMH, Primordial, Primary, Secondary, Mature, Total Follicles | [57] |
| Zhang 2024 | China | Rats (Sprague-Dawley) | CTX | 60 | Exosomes | 100 μg exosomes | IV | Pregnancy, offspring, Ovarian weight, Primordial, Primary, Secondary, and Atretic Follicles, E2, FSH, LH, AMH | [58] |

### 3.2. Outcomes

#### 3.2.1. Pooled Analysis of included studies

##### 3.2.1.1. Serum hormonal function

Animals treated with BM-MSC therapy had a significant improvement in hormonal function, increasing E2 (SMD 3.11; 95% confidence interval (CI:) [2.38, 3.84]; p<0.00001; I² = 83%; Figure 2) and AMH secretion (SMD 1.86; 95% CI: [1.03, 2.69]; p<0.0001; I² = 78%; Figure 3), decreasing FSH (SMD -3.54; 95%; CI: [-4.37, -2.71]; p<0.00001; I² = 86%; Figure 4) and LH (SMD -3.44; 95% CI: [-5.17, -1.70]; p=0.0001; I² = 89%, Supplemental Figure 1).

**Fig. 2.**
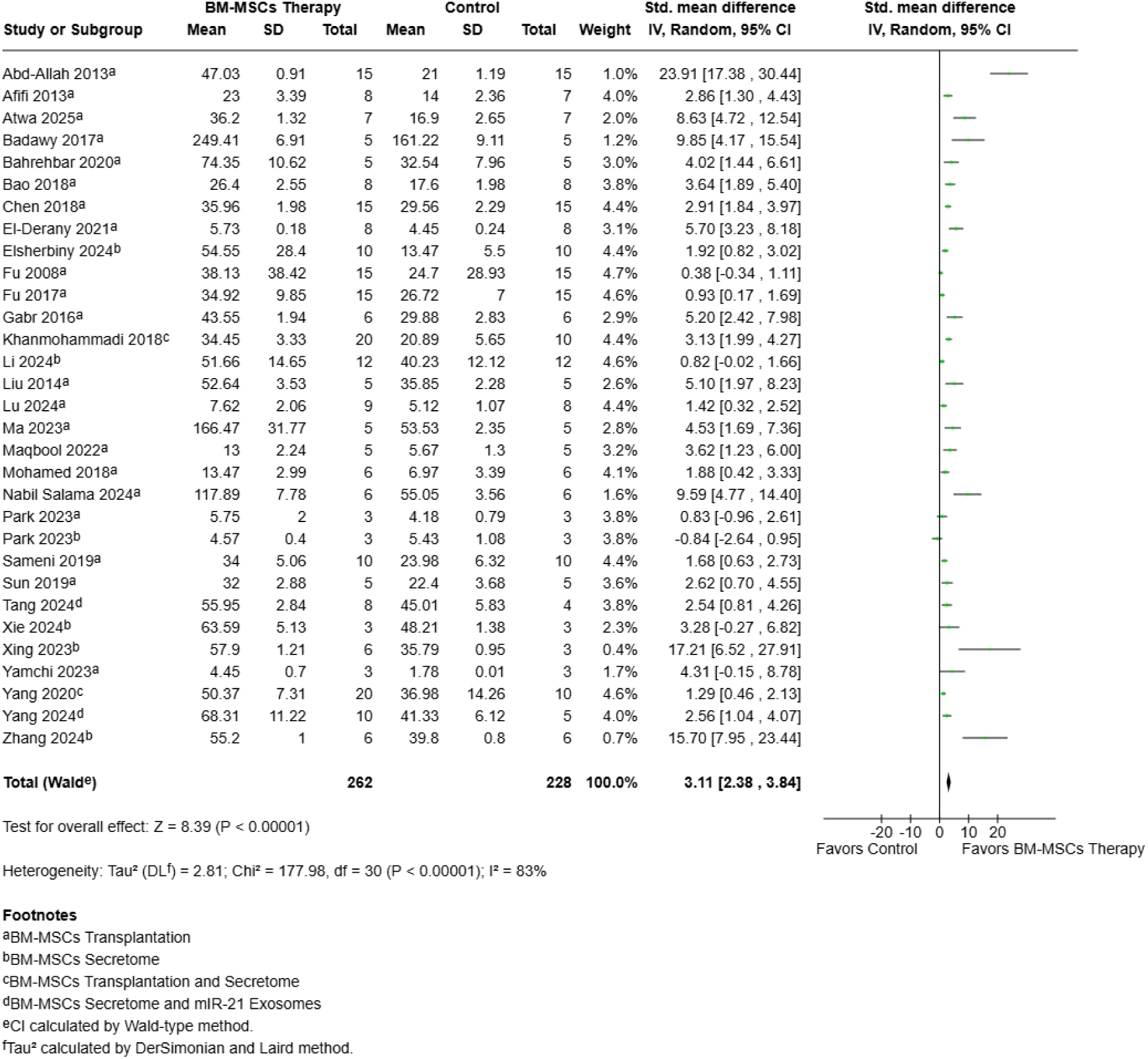
Forest plot of the pooled effect of BM-MSC therapy on serum estradiol levels. Forest plot showing the standardized mean difference (SMD) in serum estradiol (E2) levels between animals treated with bone marrow–derived mesenchymal stem cell (BM-MSC)-based therapy and control groups. Effect estimates were pooled using a random-effects inverse-variance model and are presented with 95% confidence intervals (CIs). Individual study estimates are shown as squares, with the size of each square reflecting the study weight, and the pooled effect estimate is shown as a diamond. BM-MSC-based therapy significantly increased serum E2 levels compared with controls (SMD = 3.11, 95% CI 2.38 to 3.84; P < 0.00001). Substantial heterogeneity was observed across studies (I² = 83%). Values to the right of the line of no effect favor BM-MSC therapy. BM-MSC therapy included BM-MSC transplantation, BM-MSC secretome, and BM-MSC miR-21 exosomes.

**Fig. 3.**
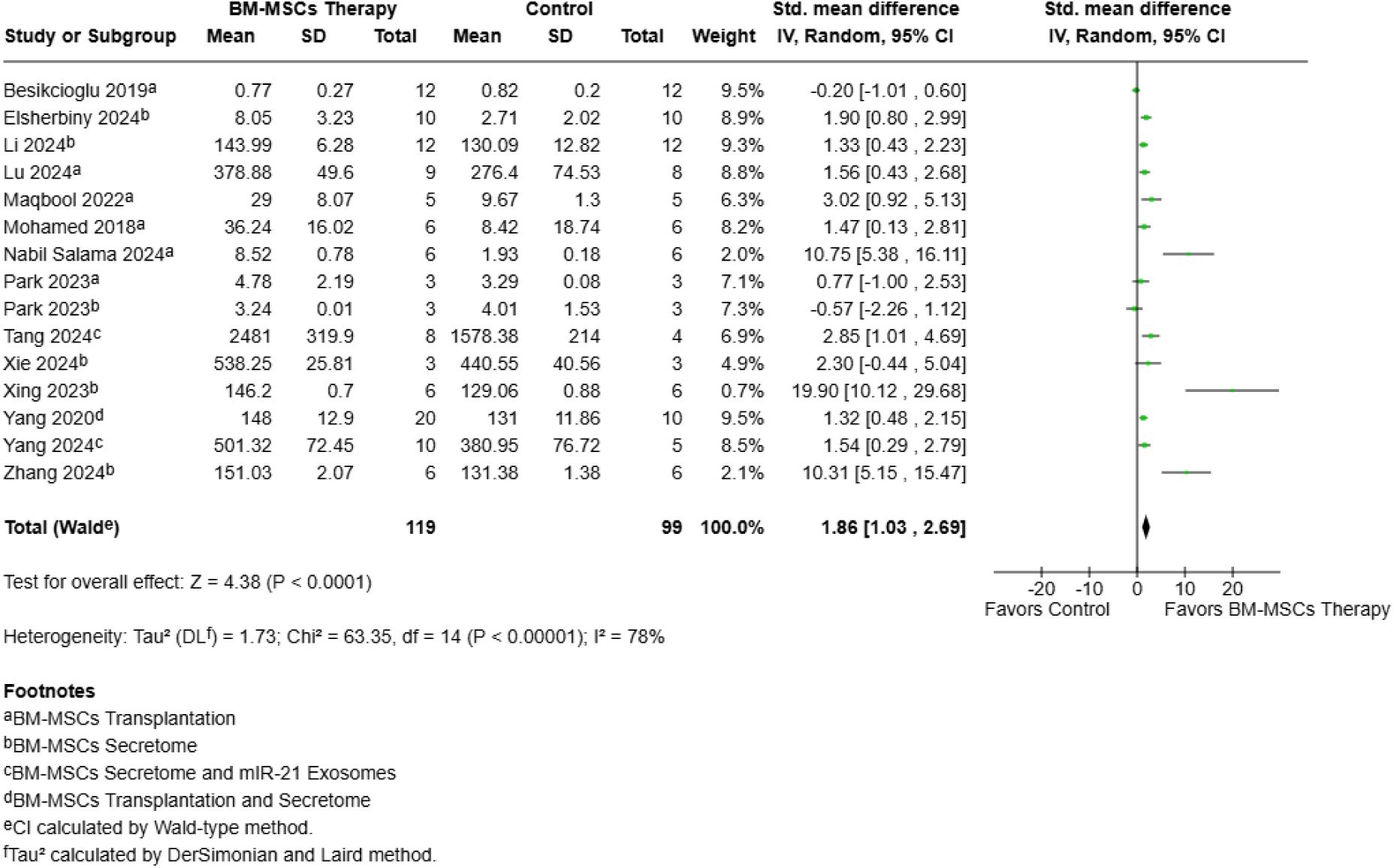
Forest plot of the pooled effect of BM-MSC therapy on serum anti-Müllerian hormone levels. Forest plot showing the standardized mean difference (SMD) in serum anti-Müllerian hormone (AMH) levels between animals treated with bone marrow–derived mesenchymal stem cell (BM-MSC)-based therapy and control groups. Effect estimates were pooled using a random-effects inverse-variance model and are presented with 95% confidence intervals (CIs). Individual study estimates are shown as squares, with the size of each square reflecting the study weight, and the pooled effect estimate is shown as a diamond. BM-MSC-based therapy significantly increased serum AMH levels compared with controls (SMD = 1.86, 95% CI 1.03 to 2.69; P < 0.0001). Substantial heterogeneity was observed across studies (I² = 78%). BM-MSC therapy included BM-MSC transplantation, BM-MSC secretome, and BM-MSC miR-21 exosomes. Values to the right of the line of no effect favor BM-MSC therapy. Park 2023 corresponds to a single article reporting two independent experiments: one evaluating BM-MSC transplantation and the other evaluating BM-MSC-derived secretome.

**Fig. 4.**
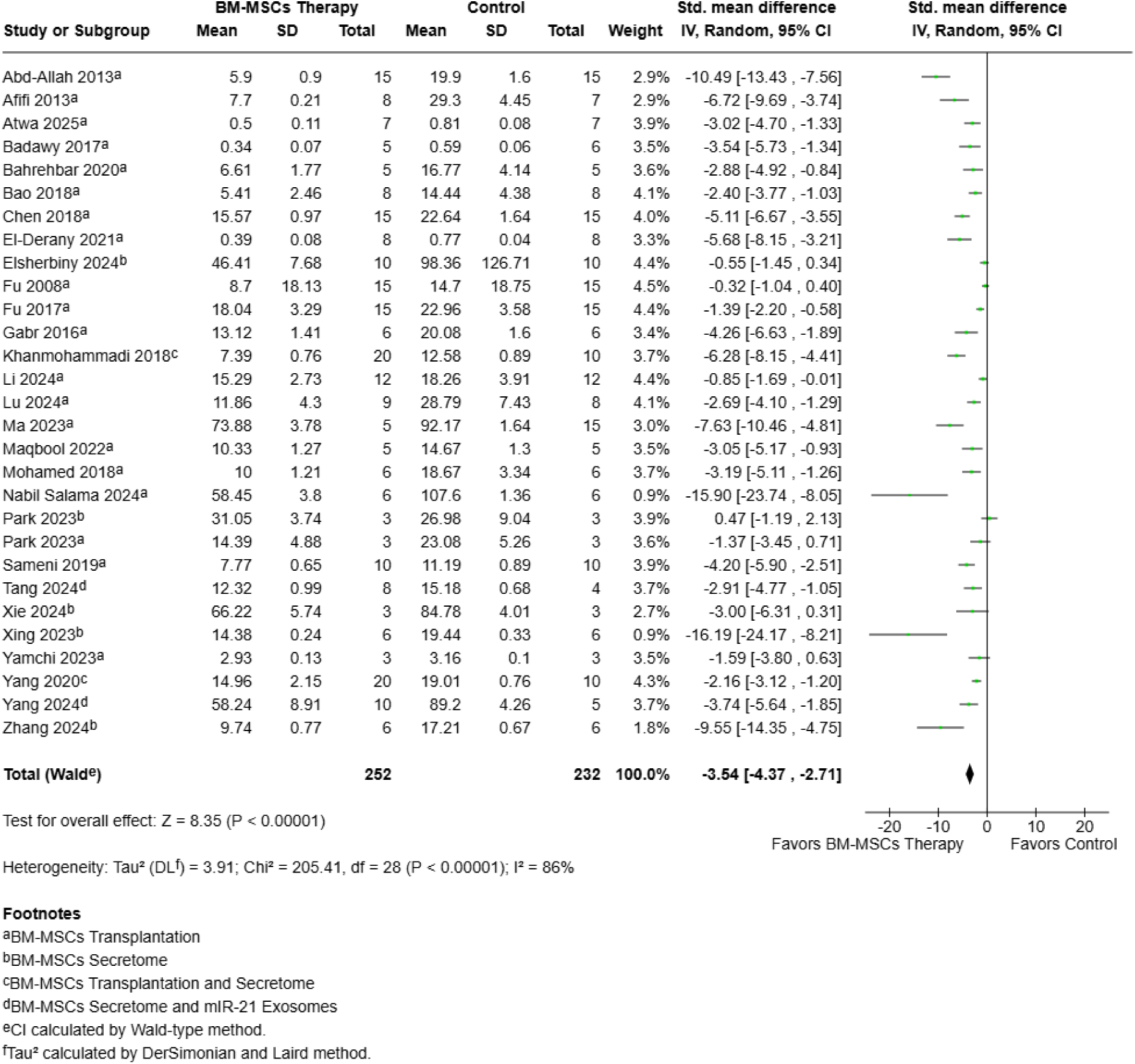
Forest plot of the pooled effect of BM-MSC therapy on serum follicle-stimulating hormone levels. Forest plot showing the standardized mean difference (SMD) in serum follicle-stimulating hormone (FSH) levels between animals treated with bone marrow–derived mesenchymal stem cell (BM-MSC)-based therapy and control groups. Effect estimates were pooled using a random-effects inverse-variance model and are presented with 95% confidence intervals (CIs). Individual study estimates are shown as squares, with the size of each square reflecting the study weight, and the pooled effect estimate is shown as a diamond. BM-MSC-based therapy significantly reduced serum FSH levels compared with controls (SMD = −3.54, 95% CI −4.37 to −2.71; P < 0.00001). Substantial heterogeneity was observed across studies (I² = 86%). Values to the left of the line of no effect favor BM-MSC therapy. BM-MSC therapy included BM-MSC transplantation, BM-MSC secretome, and BM-MSC miR-21 exosomes. Park 2023 corresponds to a single article reporting two independent experiments: one evaluating BM-MSC transplantation and the other evaluating BM-MSC-derived secretome.

##### 3.2.1.2. Folliculogenesis

Follicular production was significantly increased in the BM-MSC therapy group, including primordial (SMD 2.11; 95% CI: [1.56, 2.66]; p<0.00001; I² = 59%), primary (SMD 1.24; 95% CI: [0.92, 1.56]; p<0.00001; I² = 46%), secondary (SMD 1.62; 95% CI: [0.95, 2.28]; p<0.00001; I² = 67%), preantral (SMD 1.30; 95% CI: [0.42, 2.18]; p=0.004; I² = 38%), antral (SMD 1.45; 95% CI: [0.71, 2.19]; p=0.0001; I² = 65%), mature (SMD 2.07; 95% CI: [1.16, 2.97]; p<0.00001; I^2^ = 0%) and total follicles (SMD 4.47; 95% CI: [2.72, 6.22]; p<0.00001; I² = 86%). In contrast, there was a significant decrease in the production of atretic follicles (SMD -2.39; 95% CI: [-3.22, -1.56]; p<0.00001; I² = 58%). Forest plots are shown in Supplemental Figures 2-9.

##### 3.2.1.3. Fertility outcomes

Concerning the fertility outcomes, BM-MSC therapy was associated with a higher proportion of animals conserving normal estrous cycle (38.83% vs 1.19%; RR 7.80; 95% CI: [3.15, 19.34]; p<0.00001; I^2^ = 0%; Figure 5). There was an improvement in pregnancy (73.68% vs 16.98%; RR 3.72; 95% CI: [2.14, 6.44]; p<0.00001; I^2^ = 0%; Figure 6) and the number of offspring (SMD 1.57; 95% CI: [1.04, 2.09]; p<0.00001; I^2^ = 0%; Figure 7) was greater compared to the control group.

**Fig. 5.**
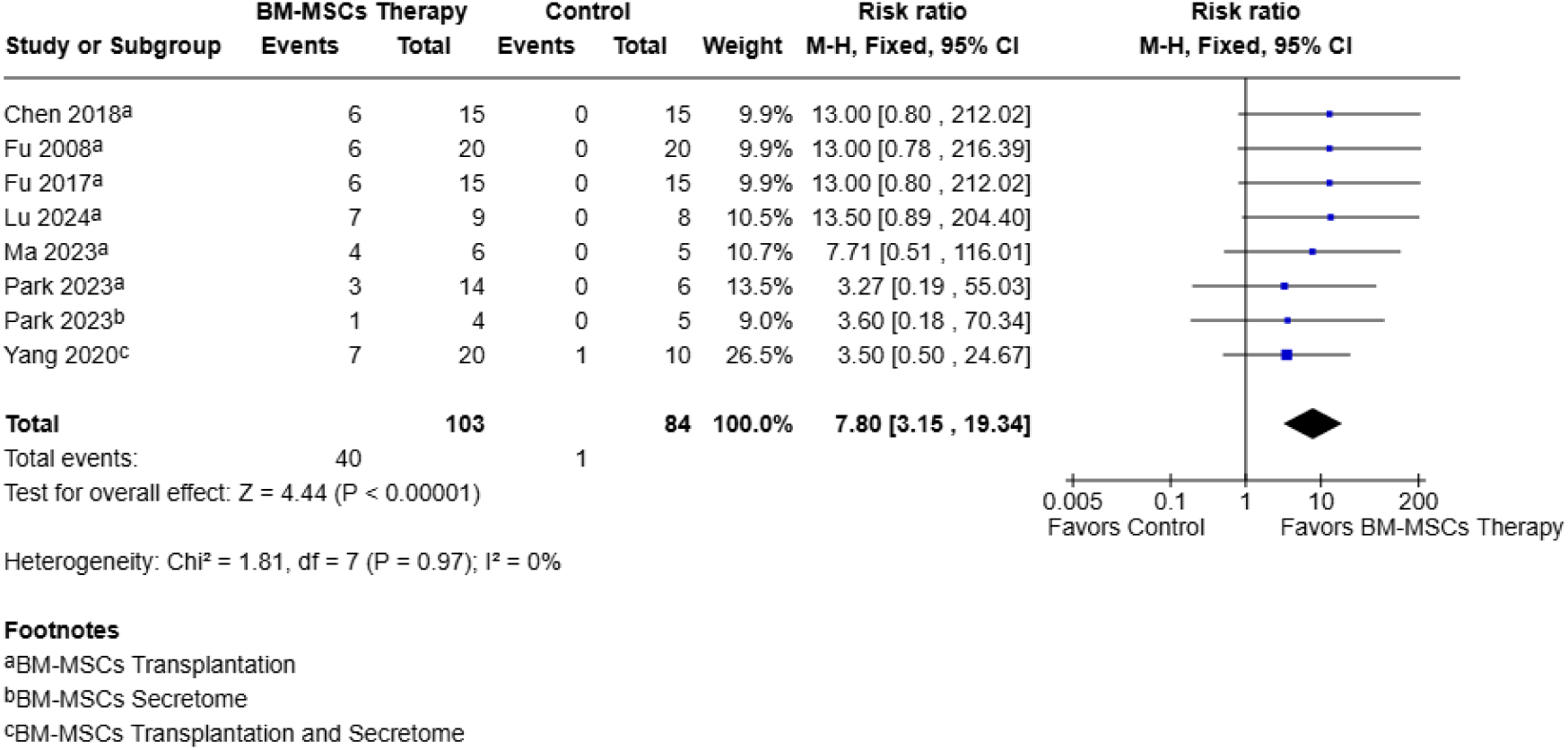
Forest plot of the pooled effect of BM-MSC therapy on restoration of a normal estrous cycle. Forest plot showing the risk ratio (RR) for restoration of a normal estrous cycle in animals treated with bone marrow–derived mesenchymal stem cell (BM-MSC)-based therapy compared with control groups. Effect estimates were pooled using a fixed-effect Mantel–Haenszel model and are presented with 95% confidence intervals (CIs). Individual study estimates are shown as squares, with the size of each square reflecting the study weight, and the pooled effect estimate is shown as a diamond. BM-MSC-based therapy was associated with a significantly higher likelihood of restoring a normal estrous cycle compared with controls (38.83% vs 1.19%; RR = 7.80, 95% CI 3.15 to 19.34; P < 0.00001). No statistical heterogeneity was observed across studies (I² = 0%). Values to the right of the line of no effect favor BM-MSC therapy. BM-MSC therapy included BM-MSC transplantation, BM-MSC secretome, and BM-MSC miR-21 exosomes.

**Fig. 6.**
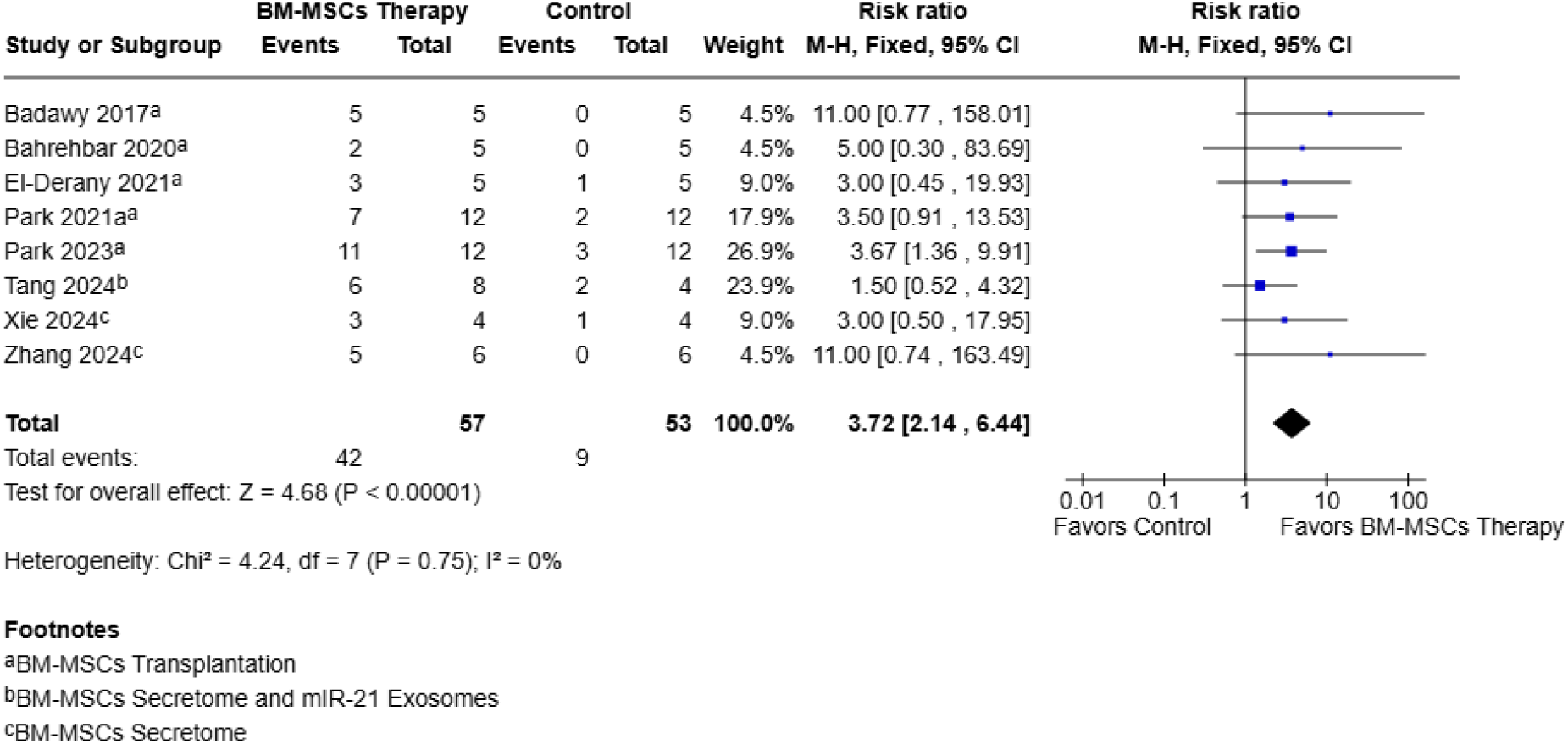
Forest plot of the pooled effect of BM-MSC therapy on pregnancy occurrence. Forest plot showing the risk ratio (RR) for pregnancy occurrence in animals treated with bone marrow–derived mesenchymal stem cell (BM-MSC)-based therapy compared with control groups. Effect estimates were pooled using a fixed-effect Mantel–Haenszel model and are presented with 95% confidence intervals (CIs). Individual study estimates are shown as squares, with the size of each square reflecting the study weight, and the pooled effect estimate is shown as a diamond. BM-MSC-based therapy was associated with a significantly higher likelihood of pregnancy occurrence compared with controls (73.68% vs 16.98%; RR = 3.72, 95% CI 2.14 to 6.44; P < 0.00001). No statistical heterogeneity was observed across studies (I² = 0%). Values to the right of the line of no effect favor BM-MSC therapy. BM-MSC therapy included BM-MSC transplantation, BM-MSC secretome, and BM-MSC miR-21 exosomes.

**Fig. 7.**
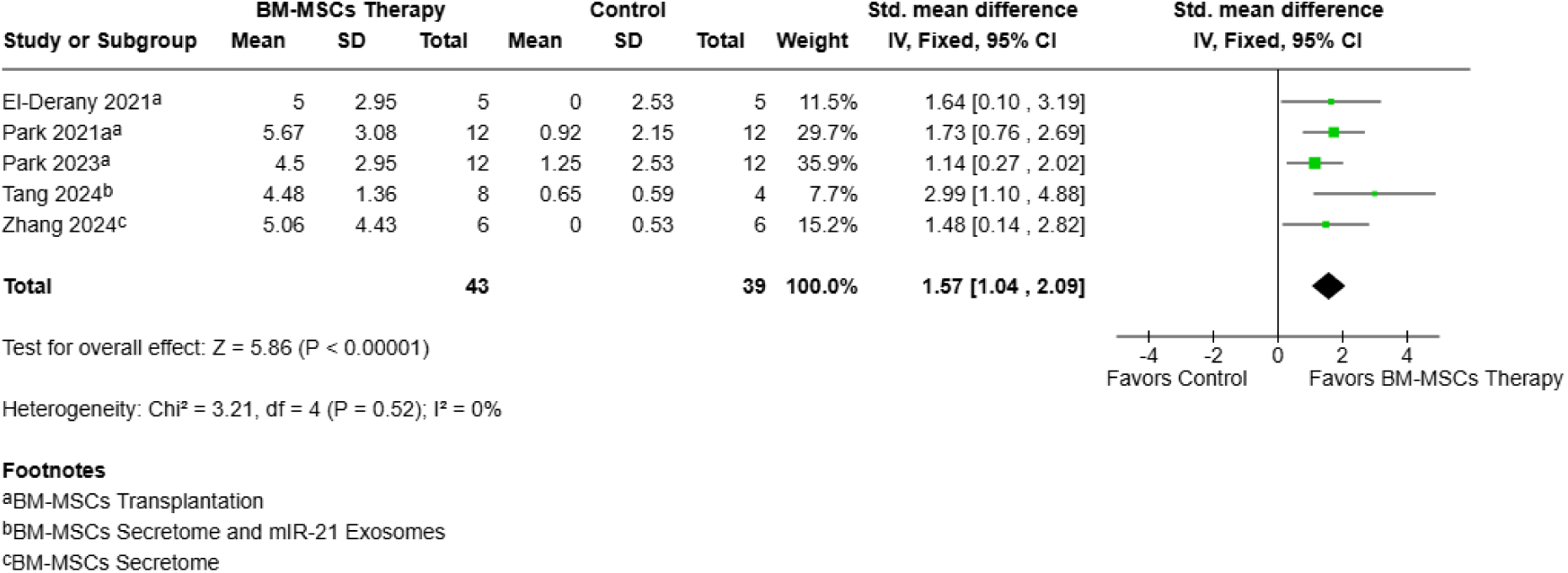
Forest plot of the pooled effect of BM-MSC therapy on the number of offspring. Forest plot showing the standardized mean difference (SMD) in the number of offspring between animals treated with bone marrow–derived mesenchymal stem cell (BM-MSC)-based therapy and control groups. Effect estimates were pooled using a fixed-effect inverse-variance model and are presented with 95% confidence intervals (CIs). Individual study estimates are shown as squares, with the size of each square reflecting the study weight, and the pooled effect estimate is shown as a diamond. BM-MSC-based therapy was associated with a significantly higher number of offspring compared with controls (SMD = 1.57, 95% CI 1.04 to 2.09; P < 0.00001). No statistical heterogeneity was observed across studies (I² = 0%). Values to the right of the line of no effect favor BM-MSC therapy. BM-MSC therapy included BM-MSC transplantation, BM-MSC secretome, and BM-MSC miR-21 exosomes.

##### 3.2.1.4. Body and ovarian weight

Ovarian size increased significantly in the BM-MSC therapy group (SMD 2.66; 95% CI: [0.79, 4.53]; p=0.005; I^2^ = 0%), following a similar trend regarding ovarian weight (SMD 3.60; 95% CI: [1.78, 5.42]; p=0.0001; I^2^ = 79%), in turn deriving in a significantly increased ovarian weight index (SMD 2.01; 95% CI: [0.83, 3.20]; p=0.0009; I² = 53%). No significant difference was found between groups in body weight (SMD 0.62; 95% CI: [-0.87, 2.10]; p=0.42; I2 = 85%). Forest plots are shown in Supplemental Figures 10-13.

#### 3.2.2. Subgroup analysis

In a subanalysis of BM-MSCs transplantation limited to cellular therapy, comparable outcomes were observed. Specifically, E2 and AMH levels were significantly increased, whereas LH and FSH levels were significantly reduced. Folliculogenesis was markedly improved, as evidenced by significant increases in primordial, primary, secondary, preantral, antral, and total follicle counts, accompanied by a decrease in atretic follicles. Fertility outcomes also improved, with a significantly higher proportion of animals exhibiting a normal estrous cycle, along with increased pregnancy percentage and offspring numbers. Additionally, ovarian size, ovarian weight, and ovarian weight index were significantly greater in the cellular therapy group. No statistical difference in body weight was found in the cellular therapy group. Due to the limited number of studies reporting mature follicles (n = 1), a subgroup analysis could not be performed.

Among acellular therapies, BM-MSC secretome showed significant effects on hormonal secretion, characterized by increased E2 and AMH levels and decreased FSH and LH levels. Regarding follicular outcomes, BM-MSC secretome were associated with significant increases in primordial, primary, secondary, and total follicle counts, together with a significant reduction in atretic follicles. In addition, ovarian weight was significantly increased in the secretome group. However, in contrast to the overall analysis, no statistically significant increase was observed in the proportion of normal estrous cycles, pregnancy, number of offspring, or mature follicle count. Preantral follicles, antral follicles, body weight, ovarian size, and ovarian weight index subgroup analysis was not possible for this subgroup.

BM-MSC-derived miR-21 exosomes showed a similar hormonal profile, with increased E2 and AMH levels and reduced FSH levels. In follicular outcomes, miR-21 exosomes significantly increased primary, mature, and total follicle counts, whereas the increase in primordial and secondary follicles was not statistically significant. LH, preantral follicles, antral follicles, atretic follicles, estrous cyclicity, pregnancy, number of offspring, body weight, and ovarian morphology subgroup analysis was not possible for this subgroup. More detailed information is provided in Table 2.

**Table 2.** Subgroup analyses. Results are expressed as standardized mean differences (SMD) with 95% confidence intervals (CI) for continuous outcomes, and as risk ratios (RR) with 95% CI for dichotomous outcomes (normal estrous cycle and pregnancy). Arrows indicate the direction of effect: ↑ favorable increase; ↓ favorable decrease. Heterogeneity was quantified using I² statistics. NA: not applicable (outcome not reported or insufficient studies for quantitative analysis).

| Outcomes | Subgroup analysis |  |  |  |
| --- | --- | --- | --- | --- |
|  | Cells (BM-MSCs Transplantation) | Secretome (BM-MSCs Secretome) | miR-21 Exosomes (BM-MSCs miR-21 Exosomes) | BM-MSC therapy (Overall) |
| **E2** | ↑ [SMD] 3.46; 95% CI: [2.55, 4.37]; $p<0.00001$; $I^2 = 85\%$; 22 studies | ↑ [SMD] 2.14; 95% CI: [1.08, 3.21]; $p<0.0001$; $I^2 = 77\%$; 11 studies | ↑ [SMD] 3.11; 95% CI: [1.42, 4.79]; $p=0.0003$; $I^2 = 25\%$; 2 studies | ↑ [SMD] 3.11; 95% CI: [2.38, 3.84]; $p<0.00001$; $I^2 = 83\%$; 30 studies |
| **FSH** | ↓ [SMD] -3.54; 95% CI: [-4.45, -2.62]; $p<0.00001$; $I^2 = 84\%$; 21 studies | ↓ [SMD] -2.78; 95% CI: [-4.42, -1.14]; $p=0.0009$; $I^2 = 89\%$; 10 studies | ↓ [SMD] -5.09; 95% CI: [-7.49, -2.68]; $p<0.0001$; $I^2 = 0\%$; 2 studies | ↓ [SMD] -3.54; 95% CI: [-4.37, -2.71]; $p<0.00001$; $I^2 = 86\%$; 28 studies |
| **AMH** | ↑ [SMD] 1.49; 95% CI: [0.40, 2.59]; $p=0.008$; $I^2 = 77\%$; 7 studies | ↑ [SMD] 1.97; 95% CI: [0.77, 3.16]; $p=0.001$; $I^2 = 78\%$; 9 studies | ↑ [SMD] 2.69; 95% CI: [1.10, 4.27]; $p=0.0009$; $I^2 = 48\%$; 2 studies | ↑ [SMD] 1.86; 95% CI: [1.03, 2.69]; $p<0.0001$; $I^2 = 78\%$; 14 studies |
| **LH** | ↓ [SMD] -3.17; 95% CI: [-5.02, -1.33]; $p=0.0007$; $I^2 = 80\%$; 5 studies | ↓ [SMD] -2.92; 95% CI: [-3.46, -2.38]; $p<0.00001$; $I^2 = 93\%$; 5 studies | NA | ↓ [SMD] -3.44; 95% CI: [-5.17, -1.70]; $p=0.0001$; $I^2 = 89\%$; 9 studies |
| **Primordial Follicles** | ↑ [SMD] 1.83; 95% CI: [1.24, 2.42]; $p<0.00001$; $I^2 = 54\%$; 13 studies | ↑ [SMD] 2.44; 95% CI: [1.82, 3.06]; $p<0.00001$; $I^2 = 49\%$; 7 studies | [SMD] 5.03; 95% CI: [-2.80, 12.85]; $p=0.21$; $I^2 = 80\%$; 2 studies | ↑ [SMD] 2.11; 95% CI: [1.56, 2.66]; $p<0.00001$; $I^2 = 59\%$; 18 studies |
| **Primary Follicles** | ↑ [SMD] 0.88; 95% CI: [0.51, 1.24]; $p<0.00001$; $I^2 = 0\%$; 11 studies | ↑ [SMD] 2.19; 95% CI: [1.31, 3.08]; $p<0.00001$; $I^2 = 52\%$; 7 studies | ↑ [SMD] 5.13; 95% CI: [2.63, 7.62]; $p<0.0001$; $I^2 = 13\%$; 2 studies | ↑ [SMD] 1.24; 95% CI: [0.92, 1.56]; $p<0.00001$; $I^2 = 46\%$; 16 studies |
| **Secondary Follicles** | ↑ [SMD] 1.05; 95% CI: [0.40, 1.71]; | ↑ [SMD] 2.02; 95% CI: [1.04, 3.00]; | [SMD] 4.72; 95% CI: [-2.02, 11.46]; $p=0.17$; $I^2 =$ | ↑ [SMD] 1.62; 95% CI: [0.95, 2.28]; |
|  | p=0.002; I<sup>2</sup> = 56%;<br>9 studies | p<0.0001; I<sup>2</sup> = 61%;<br>7 studies | 78%; 2 studies | p<0.00001; I<sup>2</sup> = 67%;<br>14 studies |
| **Preantral Follicles** | ↑ [SMD] 1.30; 95% CI: [0.42, 2.18];<br>p=0.004; I<sup>2</sup> = 38%;<br>2 studies | NA | NA | ↑ [SMD] 1.30; 95% CI: [0.42, 2.18];<br>p=0.004; I<sup>2</sup> = 38%;<br>2 studies |
| **Antral Follicles** | ↑ [SMD] 1.46; 95% CI: [0.70, 2.22];<br>p=0.0002; I<sup>2</sup> = 65%;<br>10 studies | NA | NA | ↑ [SMD] 1.45; 95% CI: [0.71, 2.19];<br>p=0.0001; I<sup>2</sup> = 65%;<br>10 studies |
| **Total Follicles** | ↑ [SMD] 5.69; 95% CI: [2.27, 8.60];<br>p=0.0001; I<sup>2</sup> = 90%;<br>7 studies | ↑[SMD] 4.25; 95% CI: [1.94, 6.57];<br>p=0.003; I<sup>2</sup> = 70%;<br>5 studies | ↑ [SMD] 6.42; 95% CI: [0.84, 12.00]; p=0.02;<br>I<sup>2</sup>= 56%; 2 studies | ↑ [SMD] 4.47; 95% CI: [2.72, 6.22];<br>p<0.00001; I<sup>2</sup> = 86%;<br>10 studies |
| **Mature Follicles** | NA | [SMD] 2.69; 95% CI: [-1.47, 6.85]; p=0.21;<br>I<sup>2</sup> = 77% ; 2 studies | ↑ [SMD] 3.18; 95% CI: [1.52, 4.85]; p=0.0002;<br>I<sup>2</sup> = 0%; 2 studies | ↑ [SMD] 2.07; 95% CI: [1.16, 2.97];<br>p<0.00001; I<sup>2</sup> = 0%;<br>3 studies |
| **Atretic Follicles** | ↓ [SMD] -1.62; 95% CI: [-2.29, -0.94];<br>p<0.00001; I<sup>2</sup> = 26%;<br>4 studies | ↓ [SMD] -3.14; 95% CI: [-3.86, -2.41];<br>p<0.00001; I<sup>2</sup> = 0%;<br>4 studies | NA | ↓ [SMD] -2.39; 95% CI: [-3.22, -1.56];<br>p<0.00001; I<sup>2</sup> = 58%;<br>7 studies |
| **Normal estrous cycle** | ↑ [RR] 7.35; 95% CI: [2.72, 19.88];<br>p<0.0001; I<sup>2</sup> = 0%;<br>7 studies | [RR] 3.87; 95% CI: [0.73, 20.44]; p=0.11;<br>I<sup>2</sup> = 0%; 2 studies | NA | ↑ [RR] 7.80; 95% CI: [3.15, 19.34];<br>p<0.00001; I<sup>2</sup> = 0%;<br>7 studies |
| **Pregnancy** | ↑ [RR] 3.87; 95% CI: [1.94, 7.71];<br>p=0.0001; I<sup>2</sup> = 0%;<br>5 studies | [RR] 2.23; 95% CI: [0.91, 5.50]; p=0.08;<br>I<sup>2</sup> = 0%; 3 studies | NA | ↑ [RR] 3.72; 95% CI: [2.14, 6.44];<br>p<0.00001; I<sup>2</sup> = 0%;<br>8 studies |
| **Number of offspring** | ↑ [SMD] 1.33; 95% CI: [0.74, 1.91];<br>p<0.00001; I<sup>2</sup> = 0%;<br>3 studies | [SMD] 2.31; 95% CI: [0.04, 4.59]; p=0.05;<br>I<sup>2</sup> = 52%; 2 studies | NA | ↑ [SMD] 1.57; 95% CI: [1.04, 2.09];<br>p<0.00001; I<sup>2</sup> = 0%;<br>5 studies |
| **Ovarian Size** | ↑ [SMD] 4.23; 95% CI: [1.12, 7.34];<br>p=0.008; I<sup>2</sup> = 0%;<br>2 studies | NA | NA | ↑ [SMD] 2.66; 95% CI: [0.79, 4.53];<br>p=0.005; I<sup>2</sup> = 0%;<br>2 studies |
| **Ovarian Weight** | ↑ [SMD] 2.02; 95% CI: [1.40, 2.65];<br>p<0.00001; I<sup>2</sup> = 49%;<br>6 studies | ↑ [SMD] 10.07; 95% CI: [1.25, 18.89];<br>p=0.03; I<sup>2</sup> = 86%;<br>3 studies | NA | ↑ [SMD] 3.60; 95% CI: [1.78, 5.42];<br>p=0.0001; I<sup>2</sup> = 79%;<br>7 studies |
| **Body Weight** | [SMD] 0.73; 95% CI: [-1.42, 2.87]; p=0.51;<br>I<sup>2</sup> = 88%;<br>5 studies | NA | NA | [SMD] 0.62; 95% CI: [-0.87, 2.10]; p=0.42;<br>I<sup>2</sup> = 85%;<br>6 studies |
| **Ovarian Weight Index** | ↑ [SMD] 1.53; 95% CI: [0.72, 2.33];<br>p=0.0002; I<sup>2</sup> = 0%; | NA | NA | ↑ [SMD] 2.01; 95% CI: [0.83, 3.20];<br>p=0.0009; I<sup>2</sup> = 53%; |
|  | 2 studies |  |  | 3 studies |

#### 3.2.3 Sensitivity analysis (Leave-one-out) and subgroup analysis of heterogeneity and funnel plots

Sensitivity analyses by leave-one-out showed no significant variation in E2, FSH, AMH, pregnancy, pups per litter, and total follicles after sequential exclusion of individual studies. Leave-one-out analysis identified Ma et al. (2023) [43] as the main driver of heterogeneity in follicular outcomes: its removal increased heterogeneity for mature follicles from 0% to 38%, while reducing it for atretic follicles from 58% to 9%. Detailed analyses can be found in Supplementary material.

Subgroup analysis of possible heterogeneity factors was performed in E2, FSH, AMH, primordial follicles, primary follicles, secondary follicles, estrous cycle, pregnancy percentage, and number of offspring including by animal model, route of administration, POI induction model and measurement technique for hormonal production outcomes. None of the subgroup analyses achieved a significant reduction in heterogeneity. Detailed analyses can be found in Supplementary material X.

Publication bias was assessed using funnel plot analysis for outcomes including 10 or more studies. Evidence of potential publication bias, reflected by an asymmetric distribution of studies in the funnel plot, was observed for E2, FSH, AMH, primordial follicles, primary follicles, and total follicles. In contrast, funnel plots for secondary follicles and antral follicles did not show relevant asymmetry. Funnel plots can be found in Supplementary materials.

### 3.3. Quality assessment

The methodological quality of the studies included in this systematic review and meta-analysis was rigorously assessed using the SYRCLE risk of bias tool, specifically designed to assess the quality of animal studies. Two reviewers performed this assessment independently, and any discrepancies were resolved by consensus or with the guidance of a third senior reviewer. The evaluation focused on several key areas: sequence generation, allocation concealment, randomization, blinding, and randomized outcome assessment. The quality assessment results revealed that only three studies [45,47,49] were classified as a high risk of bias, mainly due to deficiencies in the above mentioned domains. While quality assessment highlighted several methodological limitations in the included studies, it also stressed the importance of rigorous design and reporting standards in preclinical research. These findings emphasize the need for better methodological practices in future studies to improve the reliability and applicability of research findings in the field of POI and stem cell therapy Table 3.

**Table 3.**
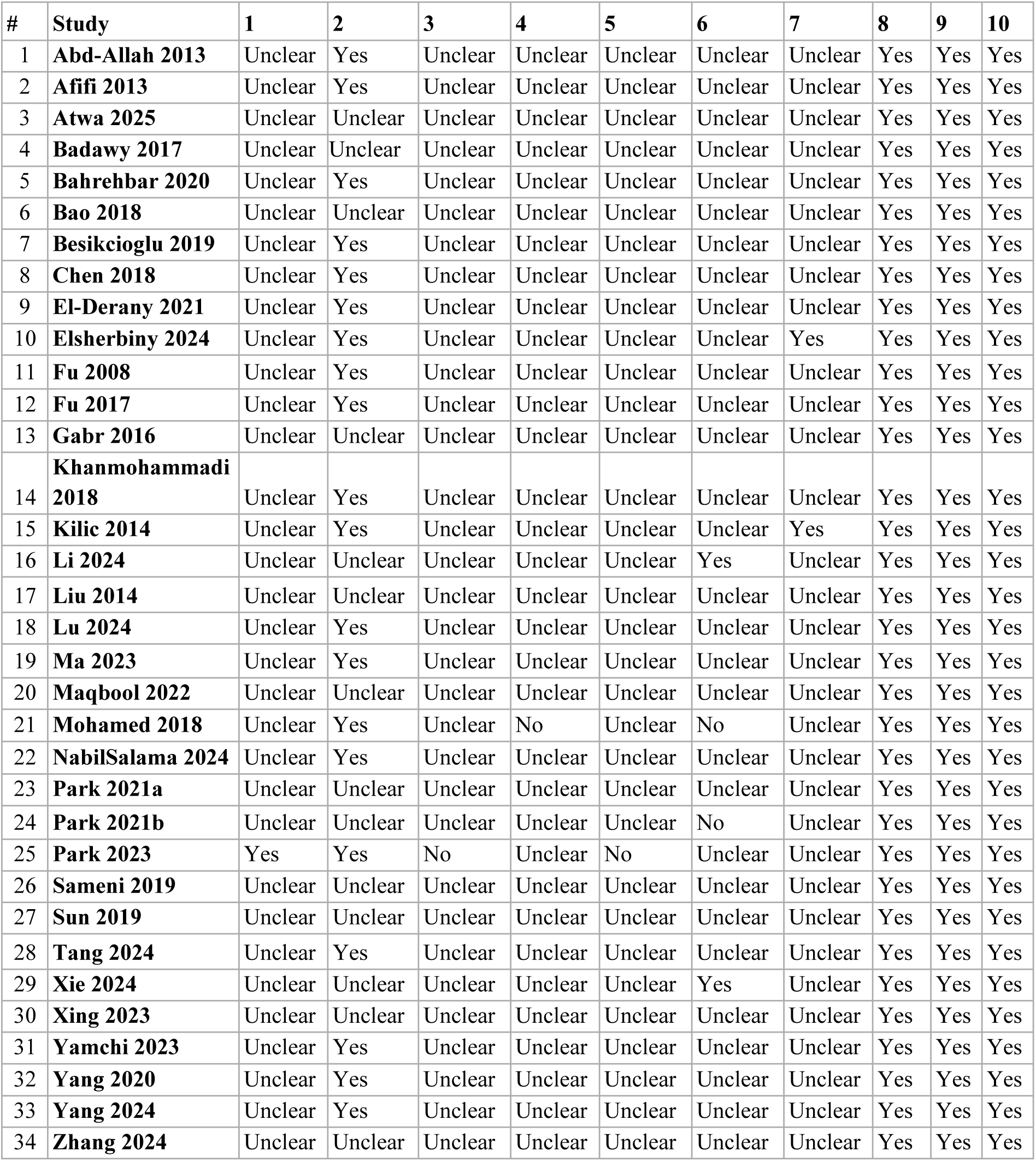
Risk of bias assessment of included studies using the SYRCLE Tool. Each domain was rated as Yes (low risk of bias), No (high risk of bias), or Unclear (insufficient information to make a judgment). Domains assessed: (1) The distribution sequence production is sufficient. (2) The baselines of groups are the same. (3) Distribution concealing is sufficient. (4) Experiment animals are randomly fed. (5) Blinding to the researcher is ensured. (6) Animals are randomly selected to assess the result. (7) There is blinding to the result evaluator. (8) No incomplete data. (9) No selective result reporting. (10) No other biases are present. SYRCLE: Systematic Review Centre for Laboratory Animal Experimentation.

## 4. Discussion

### 4.1. Summary of findings

This systematic review and meta-analysis included 34 preclinical studies comprising 1,357 animals and evaluated the effectiveness of BM-MSC therapy, encompassing both cellular transplantation and secretome administration, for the treatment of POI in animal models. The principal findings were: (1) a significant improvement in ovarian endocrine function, with increased serum E2 and AMH levels and decreased FSH and LH levels; (2) enhanced folliculogenesis across all developmental stages—primordial, primary, secondary, preantral, antral, mature and total follicle counts—accompanied by a significant reduction in atretic follicles; (3) preservation of fertility, with a higher proportion of animals maintaining a normal estrous cycle, improved pregnancy percentage and larger litter sizes compared with controls; and (4) recovery of ovarian morphology, reflected by greater ovarian size, ovarian weight and ovarian weight index, with no significant difference in overall body weight. Subgroup analyses indicated that both cell- and secretome-based therapies yielded consistent endocrine and follicular benefits. However, the effect of secretome-only therapy on estrous-cycle normalisation, pregnancy, and ovarian morphology did not reach statistical significance, likely reflecting the smaller number of studies in that subgroup. These benefits were observed despite substantial between-study heterogeneity in most endocrine outcomes and funnel-plot asymmetry suggestive of publication bias for E2, FSH, AMH, primordial, primary, and total follicle counts.

### 4.2. Comparison with other studies involving BM-MSC therapy and other sources of MSCs

To the best of our knowledge, no previous meta-analysis has focused exclusively on BM-MSC-based therapy for POI in animal models. Guo et al. in 2023 performed a systematic review and meta-analysis assessing MSCs for the prevention of POI, including 11 studies [59], using BM-MSCs, four of these met our language eligibility criterion and were incorporated into the present review [28,33,36,37], while the remaining seven were excluded as they were published in Chinese. In their BM-MSC subgroup analysis, they reported findings consistent with the present study regarding hormonal recovery, with increased E2 levels (10 studies; SMD 3.59; 95% CI 2.33 to 4.84), decreased FSH levels (11 studies; SMD −2.63; 95% CI −3.48 to −1.78), and increased total follicle counts (2 studies; SMD 1.06; 95% CI 0.08 to 2.05). However, no significant differences were observed for LH levels (2 studies; SMD −4.21; 95% CI −8.90 to 0.47) or primordial follicle counts (2 studies; SMD 6.09; 95% CI −4.54 to 16.72), which may be explained by the limited statistical power of these subgroup analyses. Moreover, several outcomes, including primary, secondary, antral, atretic, and mature follicle counts, as well as the proportion of animals with normal estrous cycles, were represented by only one study, precluding quantitative synthesis. AMH was not assessed in any BM-MSC study included in that review.

Other meta-analyses have evaluated MSC-based therapy from different cellular sources. Wang et al. (2023) [60] assessed human umbilical cord-derived MSCs (hUCMSCs) and reported a similar pattern of ovarian recovery, including improved hormonal profiles, normalization of the estrous cycle, and enhanced folliculogenesis. However, they found no significant difference in atretic follicle counts, and fertility-related outcomes such as pregnancy and number of offspring were not assessed. Similarly, Zhao et al. (2026) recently evaluated UCMSC transplantation in animal models of premature ovarian failure, including 10 randomized animal studies comprising 365 animals. Their meta-analysis showed significant increases in primordial, primary, secondary, and antral follicle counts, as well as increased E2 and AMH levels and decreased FSH levels. These findings are directionally consistent with our BM-MSC-focused results, supporting the broader regenerative potential of MSC-based therapies in POI models. Nevertheless, Zhao et al. (2026) also reported substantial heterogeneity across all pooled outcomes and highlighted methodological limitations, particularly related to allocation concealment, blinding, and incomplete outcome reporting [61].

Compared with these previous studies, the present meta-analysis expands the available evidence by focusing specifically on BM-MSC-based interventions and by including both cellular therapy and acellular therapy through BM-MSC-derived secretome. In addition, our review incorporates a larger number of studies and animals, and evaluates a broader range of outcomes, including hormonal markers, follicular development, atretic follicles, estrous cycle recovery, pregnancy, and litter size. These fertility-related endpoints provide a more functionally relevant assessment of ovarian recovery than hormonal and histological outcomes alone, and have not been evaluated in previous meta-analyses of BM-MSCs cellular therapy.

Regarding acellular therapy, two meta-analyses published in 2024 evaluated MSC-derived or stem cell-derived extracellular vesicles in POI models; however, neither focused exclusively on BM-MSC-derived extracellular vesicles [62,63]. Firouzabadi et al. (2024) [62] included four studies involving BM-MSC-derived secretome [47,51,54,56]. Subgroup analyses stratified by human and murine sources were performed for hormonal and follicular outcomes, but pooled analysis was only possible for E2 in human BM-MSC-derived exosome studies, showing a significant increase in E2 levels (2 studies; SMD 2.93; 95% CI 1.51 to 4.33). Other subgroup analyses included only one study each, limiting quantitative interpretation. Luo et al. (2024) [63], evaluated stem cell-derived extracellular vesicles more broadly, not restricting inclusion to MSC-derived vesicles. Their subgroup analysis included four studies assessing BM-MSC-derived secretome [47,51,54,56] and reported significant improvements in primordial follicles (2 studies; SMD 5.29; 95% CI 3.64 to 6.93), primary follicles (2 studies; SMD 3.97; 95% CI 2.48 to 5.47), and secondary follicles (2 studies; SMD 3.76; 95% CI 2.49 to 5.03). However, no other outcomes were reported for this specific subgroup. A recent network meta-analysis from 2026 by Yu et al. [64] compared exosomes derived from different mesenchymal stem cell sources, including human adipose tissue, amniotic fluid, umbilical cord, and bone marrow. This review included six studies evaluating BM-MSC-derived exosomes or secretome-based interventions. Of these, five studies overlapped with the present BM-MSC secretome subgroup, while Park et al. (2024) [65] was excluded from our analysis because the intervention was administered preventively, before induction of the POI model. In their subgroup analysis, Yu et al. (2026) reported significant improvements in the primary outcomes, including increased AMH levels (SMD = 3.55; 95% CI, 1.74 to 5.37) and antral follicle counts (SMD = 4.33; 95% CI, 2.06 to 6.60), along with reduced FSH levels (SMD = −3.18; 95% CI, −4.78 to −1.58). Secondary outcomes also favored exosome-based therapy, with increased E2 levels (SMD = 3.85; 95% CI, 2.16 to 5.53), primordial follicle counts (SMD = 3.18; 95% CI, 1.35 to 5.02), and pregnancy rate (SMD = 2.17; 95% CI, 1.09 to 4.35) [64].

Overall, these previous meta-analyses support the biological plausibility and therapeutic potential of MSC-based and extracellular vesicle-based interventions for POI. However, the present study provides a more comprehensive and source-specific synthesis of BM-MSC therapy, including both cellular and acellular strategies.

### 4.3. Pathophysiology of BM-MSC Therapy

BM-MSC therapy may improve POI predominantly through paracrine-mediated repair of the ovarian microenvironment rather than direct follicular replacement [14,41]. In a chemotherapy-induced POI model, Fu et al. (2008) showed that BM-MSC transplantation improved ovarian structure and function, reduced granulosa cell apoptosis, upregulated Bcl-2, and that BM-MSCs secreted VEGF, HGF, and IGF-1, supporting anti-apoptotic and trophic mechanisms [36]. More recent experimental evidence further supports this paracrine model: human BM-MSC-conditioned medium enhanced granulosa cell proliferation, increased steroidogenic markers such as aromatase (CYP19A1) and StAR gene expression restoring granulosa cell function [49]. In addition, BM-MSC secretome has been shown to enhance angiogenic signaling through increased markers including endogline, Tie-2, VEGF-R2, VEGF, and VE-Cadherin in human ovarian microvascular endothelial cells, supporting vascular regeneration as a potential contributor to restore folliculogenesis and ovulation [66]. In radiotherapy-induced POI model, BM-MSCs also improved ovarian function through regulation of TGF-β, canonical Wnt/β-catenin, and Hippo pathway signaling, suggesting that BM-MSC therapy may modulate expression of genes involved in folliculogenesis/follicle development [33].

MSC-derived secretome/exosome represent an acellular therapeutic approach that may reproduce these paracrine effects by delivering regulatory miRNAs and other bioactive molecules to ovarian granulosa and stromal cells [67]. Sun et al. (2019) showed that BM-MSC-derived exosomal miR-644-5p inhibited ovarian granulosa cell apoptosis by regulating p53, providing direct evidence for exosome-mediated anti-apoptotic signaling [51]. Yang et al. (2020) demonstrated that BM-MSC-derived exosomal miR-144-5p inhibits the apoptosis of cyclophosphamide-damaged granulosa cells by targeting PTEN, supporting involvement of the PI3K-Akt pathway [56]. Similarly, miR-21 overexpression in BM-MSCs improved ovarian structure and function in chemotherapy-induced ovarian damage by targeting PDCD4 and PTEN to inhibit granulosa cell apoptosis [35]. More recently, miR-21-5p-loaded BM-MSC-derived exosomes were shown to repair ovarian function in autoimmune POI by targeting MSX1/Notch signaling pathway, with associated effects on granulosa cell proliferation, apoptosis and hormone synthesis [57]. Together, these studies suggest that BM-MSC-derived secretome act as concentrated paracrine mediators capable of regulating apoptosis, follicular survival, and steroidogenic recovery in preclinical POI models.

### 4.4. Strengths and limitations

This systematic review and meta-analysis is among the first to specifically focus on BM-MSC-based therapy for POI, providing a more homogeneous synthesis of the available evidence. Compared with previous reviews, this study adds a substantial number of newly published articles, including 22 studies evaluating BM-MSC transplantation, 3 studies assessing BM-MSC secretome, and a subgroup of 2 studies with miR-21-enriched exosomes intervention groups. To our knowledge, this is also the first meta-analysis of BM-MSCs cellular therapy to assess fertility-related outcomes beyond hormonal levels and estrous cyclicity, including pregnancy and pups per litter. Also, no previous meta-analysis has included miR-21 exosomes subgroup analysis. Methodologically, heterogeneity was addressed using a random-effects model with the DerSimonian and Laird method, and subgroup analyses were performed to explore potential causing variables across the primary outcomes. In addition, the robustness of the findings was evaluated through leave-one-out sensitivity analyses, which generally showed consistent results.

This study has several limitations. First, many included studies lacked detailed methodological reporting, which led to unclear risk-of-bias judgments, particularly in domains such as sequence generation, allocation concealment, random housing, blinding, and random outcome assessment. Second, significant publication bias was identified for several outcomes, including E2, FSH, AMH, primordial follicles, primary follicles, and total follicles, which may affect the certainty of the evidence. Third, safety outcomes were poorly reported across the included studies. This is particularly relevant given the experimental nature of BM-MSC-based therapies, and most studies had relatively short follow-up periods, ranging from 7 days to 17 weeks, limiting the assessment of long-term safety and efficacy. Finally, substantial inter-study heterogeneity was observed across several methodological and biological factors, including animal models, POI induction methods, route of administration, timing of transplantation, follow-up duration, and outcome measurement techniques and units. Although detailed subgroup analyses were conducted to explore potential sources of heterogeneity, no single factor consistently explained the observed variability.

### 4.5. Implications for Clinical Practice and Future Research

Although BM-MSC therapy remains at a preclinical stage and cannot yet be recommended for routine clinical use in POI, the consistent endocrine, follicular, and fertility benefits observed across animal studies support further translational research. This is also consistent with the recent clinical meta-analysis by Umer et al. (2024), which suggested that stem cell transplantation may improve ovarian function in women with POI, including changes in E2, AMH, follicle count, pregnancy, and live birth outcomes; however, this evidence was limited by the small number of human studies, small sample sizes, heterogeneous stem cell sources, variable protocols, and inconsistent follow-up [68]. Therefore, these findings should be interpreted cautiously, particularly because most preclinical studies rely on cyclophosphamide-induced POI models, which do not fully capture the heterogeneous etiologies of human POI [6,69]. Future studies should improve the standardization of BM-MSC and extracellular vesicle characterization, explore dose–response effects and administration routes, and prospectively incorporate methodological safeguards aligned with SYRCLE and CAMARADES risk-of-bias tools [24,70], including randomization, allocation concealment, blinding, sample size calculation, and transparent reporting of exclusions and outcomes. Long-term safety endpoints should also be embedded before clinical translation.

## 5. Conclusion

This systematic review and meta-analysis of 34 preclinical studies and 1,357 animals suggests that BM-MSC-based therapy, delivered either as cellular transplantation or secretome therapy, improves ovarian function in animal models of POI. These benefits included endocrine recovery, enhanced folliculogenesis, reduced follicular atresia, improved estrous cyclicity, higher pregnancy percentage, and larger litter sizes. However, the evidence remains limited by substantial heterogeneity, incomplete methodological reporting, short follow-up, and lack of formal safety outcomes. Future preclinical studies should follow SYRCLE and CAMARADES recommendations, standardize experimental methods, and include long-term safety assessment before clinical translation.

## 6. Declarations

### 6.1 Acknowledgements

This work was supported by the Agencia Nacional de Investigación y Desarrollo (ANID, Chile) through the following grants: FONDECYT Iniciación 11230208 (to A.D.) and the Beca de Magíster Nacional 22240802 (to P.V.). Furthermore, J.P., F.M.T., P.V., B.A.A., P.C., J.A.A., and A.D. acknowledge the support of the Institute for Cell Dynamics and Biotechnology (ICDB) and the Centre for Biotechnology and Bioengineering (CeBiB) via the ANID PIA/Basal projects FB0001 and AFB240001.

### 6.2. Author contribution statement

Conceptualization: J.P., F.M.T., and A.D.; Methodology: J.P., F.M.T., and A.D.; Validation: J.P., F.M.T., and A.D.; Formal analysis: J.P., F.M.T., and A.D.; Investigation: J.P., F.M.T., P.V., and A.D.; Writing – original draft: J.P., F.M.T., and A.D.; Writing – review & editing: J.P., F.M.T., P.V., D.V., B.A.A., P.C., J.A.A., and A.D.; Visualization: J.P., F.M.T., P.V., and A.D.; Supervision: A.D.; Project administration: A.D.; Funding acquisition: B.A.A., P.C., J.A.A., and A.D. All authors have read and approved the final version of the manuscript.

### 6.4. Ethics approval

Not applicable

### 6.5. Consent to participate

Not applicable

### 6.6. Consent for publication

Not applicable

### 6.7. Availability of data and material

All data supporting the findings of this study are available within the paper and its Supplementary Information.

### 6.8. Code availability

Not applicable

### Declaration of generative AI and AI-assisted technologies in the writing process

During the preparation of this work, the author(s) used ChatGPT and Claude AI assisted to improve organization, grammar and readability. After using this tool/service, the author(s) reviewed and edited the content as needed and take(s) full responsibility for the content of the publication.

