## Supplementary Material for "Bone Marrow Mesenchymal Stem Cells Therapy for Premature Ovarian Insufficiency: A Systematic Review and Meta-analysis of Preclinical Studies"

**Table S1. Search strategy used across bibliographic databases**. Detailed search strategies used for each consulted database, including PubMed, Web of Science, Cochrane Library, Scopus, and ScienceDirect. The search strategy was developed using controlled vocabulary and free-text terms related to “premature ovarian insufficiency” and “mesenchymal stem cells,” together with their respective synonyms and database-specific adaptations.

| **Searches** | **Search strategy** |
|---|---|
| PubMed | #1:(“Primary Ovarian Insufficiency” [MeSH Terms])<br>#2: ((Premature ovarian insufficiency [Title/Abstract]) OR (Premature ovarian failure [Title/Abstract]) OR (chemotherapy-induced ovarian failure [Title/Abstract]) OR (chemotherapy-induced ovarian insufficiency [Title/Abstract]) OR (cisplatin-induced ovarian failure [Title/Abstract]) OR (cisplatin-induced ovarian insufficiency [Title/Abstract]) OR (idiopathic premature ovarian failure [Title/Abstract]) OR (idiopathic premature ovarian insufficiency[Title/Abstract]))<br>#3: #1 OR #2<br>#4: (“Mesenchymal Stem Cells” [MeSH Terms]) 50,685<br>#5 ((Mesenchymal stem cell*[Title/Abstract]) OR (Bone Marrow Stem Cell*[Title/Abstract]) OR (Bone Marrow Stromal Cell*[Title/Abstract]) OR (Mesenchymal Progenitor Cell*[Title/Abstract]) OR (Mesenchymal Stromal Cell*[Title/Abstract]) OR (Human Umbilical Cord Mesenchymal Stem Cell* [Title/Abstract]) OR (Wharton’s Jelly Mesenchymal Stem Cell* [Title/Abstract]) OR (Adipose-Derived Mesenchymal Stem Cell* [Title/Abstract]) OR (Adipose Derived Mesenchymal Stem Cell* [Title/Abstract]) OR (Human Adipose Derived Mesenchymal Stem Cell* [Title/Abstract]) OR (Adipose Tissue-Derived Mesenchymal Stem Cell* [Title/Abstract]) OR (Adipose Tissue Derived Mesenchymal Stem Cell* [Title/Abstract]) OR (Human Endometrial Stem Cell* [Title/Abstract]) OR (Human Menstrual Blood Stem Cell* [Title/Abstract]))<br>#6: #4 OR #5<br>#7: #3 AND #6 |

| | |
|---|---|
| Web Of Science | #1: TS=('Mesenchymal stem cell*' OR 'Bone Marrow Stromal Cell*' OR 'Bone Marrow Stem Cell*' OR 'Mesenchymal Progenitor Cell*' OR 'Mesenchymal Stromal Cell*' OR 'Human Umbilical Cord Mesenchymal Stem Cell*' OR 'Wharton’s Jelly Mesenchymal Stem Cell*' OR 'Adipose-Derived Mesenchymal Stem Cell*' OR 'Adipose Derived Mesenchymal Stem Cell*' OR 'Human Adipose Derived Mesenchymal Stem Cell*' OR 'Adipose Tissue-Derived Mesenchymal Stem Cell*' OR 'Adipose Tissue Derived Mesenchymal Stem Cell*' OR 'Human Endometrial Stem Cell*' OR 'Human Menstrual Blood Stem Cell*' )<br><br>#2: TS=('Primary Ovarian Insufficiency' OR 'Premature ovarian insufficiency' OR 'Premature ovarian failure' OR 'chemotherapy-induced ovarian failure' OR 'chemotherapy-induced ovarian insufficiency' OR 'cisplatin-induced ovarian failure' OR 'cisplatin-induced ovarian insufficiency' OR 'idiopathic premature ovarian failure' OR 'idiopathic premature ovarian insufficiency')<br><br>#3: #1 AND #2 |
| Cochrane Library | #1 MeSH descriptor: [Primary Ovarian Insufficiency] explode all trees<br>#2 “Premature ovarian failure”<br>#3 "Premature ovarian Insufficiency"<br>#4 "chemotherapy-induced ovarian failure”<br>#5 "chemotherapy-induced ovarian insufficiency"<br>#6 "cisplatin-induced ovarian failure"<br>#7 "cisplatin-induced ovarian insufficiency"<br>#8 "idiopathic premature ovarian failure"<br>#9 "idiopathic premature ovarian insufficiency"<br>#10 #1 OR #2 OR #3 OR #4 OR #5 OR #6 OR #7 OR #8 OR #9<br><br>#11 MeSH descriptor: [Mesenchymal Stem Cells] explode all trees<br><br>#12 “Mesenchymal Stem Cells”<br>#13 “Bone Marrow Stem Cells”<br>#14 “Human Umbilical Cord Mesenchymal Stem Cells”<br>#15 "Human endometrial stem cells"<br>#16 “Mesenchymal Stromal Cells”<br>#17 “Bone Marrow Stromal Cells”<br>#18 “Mesenchymal Progenitor Cells”<br>#19 "Cord Mesenchymal Stem Cell"<br>#20 "Wharton’s Jelly Mesenchymal Stem Cell"<br>#21 "Adipose-Derived Mesenchymal Stem Cell"<br>#22 "Adipose Derived Mesenchymal Stem Cell"<br>#23 "Human Adipose Derived Mesenchymal Stem Cell"<br>#24 #11 OR #12 OR #13 OR #14 OR #15 OR #16 OR #17 OR #18 OR #19 OR #20 OR #21 OR #22 OR #23<br><br>#25 #10 AND #24 |

| | |
|---|---|
| SCOPUS | #1: (TITLE-ABS-KEY("Mesenchymal stem cell*") OR TITLE-ABS-KEY("Bone Marrow Stromal Cell*") OR TITLE-ABS-KEY("Bone Marrow Stem Cell*") OR TITLE-ABS-KEY("Mesenchymal Progenitor Cell*") OR TITLE-ABS-KEY("Mesenchymal Stromal Cell*") OR TITLE-ABS-KEY("Human Umbilical Cord Mesenchymal Stem Cell*") OR TITLE-ABS-KEY("Wharton's Jelly Mesenchymal Stem Cell*") OR TITLE-ABS-KEY("Adipose-Derived Mesenchymal Stem Cell*") OR TITLE-ABS-KEY("Adipose Derived Mesenchymal Stem Cell*") OR TITLE-ABS-KEY("Human Adipose Derived Mesenchymal Stem Cell*") OR TITLE-ABS-KEY("Adipose Tissue-Derived Mesenchymal Stem Cell*") OR TITLE-ABS-KEY("Adipose Tissue Derived Mesenchymal Stem Cell*") OR TITLE-ABS-KEY("Human Endometrial Stem Cell*") OR TITLE-ABS-KEY("Human Menstrual Blood Stem Cell*"))<br><br>#2: (TITLE-ABS-KEY("Primary Ovarian Insufficiency") OR TITLE-ABS-KEY("Premature ovarian insufficiency") OR TITLE-ABS-KEY("Premature ovarian failure") OR TITLE-ABS-KEY("chemotherapy-induced ovarian failure") OR TITLE-ABS-KEY("chemotherapy-induced ovarian insufficiency") OR TITLE-ABS-KEY("cisplatin-induced ovarian failure") OR TITLE-ABS-KEY("cisplatin-induced ovarian insufficiency") OR TITLE-ABS-KEY("idiopathic premature ovarian failure") OR TITLE-ABS-KEY("idiopathic premature ovarian insufficiency"))<br><br>#3: #1 AND #2 |
| ScienceDirect | (("Primary Ovarian Insufficiency") OR ("Premature Ovarian Insufficiency") OR ("Premature ovarian Failure") OR ("cisplatin-induced ovarian insufficiency") OR ("idiopathic premature ovarian failure")) AND (("Mesenchymal Stem Cells") OR ("Mesenchymal Stromal Cells") OR ("Mesenchymal Progenitor Cells")) |

### Excluded studies

#### Supplementary Table S2. Studies excluded after full-text assessment, with reasons for exclusion.

Twenty-eight studies assessed at full text were excluded. Studies are listed grouped by reason for exclusion and, within each reason, alphabetically by first author.

| Reason for exclusion | n |
|---|---|
| Wrong intervention | 11 |
| Non-published studies (conference abstracts, poster presentations) | 5 |
| Studies published in languages other than Spanish or English | 4 |
| Non-original research (reviews, meta-analyses, editorials, letters, protocols) | 4 |
| Studies without available data | 2 |
| Wrong population (non-animal studies) | 1 |
| Wrong comparator | 1 |
| **Total** | **28** |

| # | Study | Journal | Year | Ref | DOI | Reason for exclusion |
|---|---|---|---|---|---|---|
| 1 | Buigues 2021 | *Am J Obstet Gynecol* | 2021 | [1] | 10.1016/j.ajog.2021.01.023 | Wrong intervention |
| 2 | Chen 2025 | *Redox Rep* | 2025 | [2] | 10.1080/13510002.2025.2455914 | Wrong intervention |
| 3 | Feng 2010 | *Transplantation* | 2010 | [3] | 10.1097/TP.0b013e3181ca86bb | Wrong intervention |
| 4 | Ghadami 2012 | *PLoS One* | 2012 | [4] | 10.1371/journal.pone.0032462 | Wrong intervention |
| 5 | Herraiz 2018 | *Fertil Steril* | 2018 | [5] | 10.1016/j.fertnstert.2018.01.004 | Wrong intervention |
| 6 | Kozub 2017 | *Exp Oncol* | 2017 | [6] | — | Wrong intervention |
| 7 | Lee 2007 | *J Clin Oncol* | 2007 | [7] | 10.1200/JCO.2006.10.3028 | Wrong intervention |
| 8 | Li 2024 | *Stem Cell Res Ther* | 2024 | [8] | 10.1186/s13287-024-03718-z | Wrong intervention |
| 9 | Mousaei Ghasroldasht 2024 | *J Pers Med* | 2024 | [9] | 10.3390/jpm14050482 | Wrong intervention |
| 10 | Zhang 2025 | *Mater Today Bio* | 2025 | [10] | 10.1016/j.mtbio.2025.101469 | Wrong intervention |
| 11 | Zohni 2021 | *Cancer Lett* | 2021 | [11] | 10.1016/j.canlet.2020.12.035 | Wrong intervention |
| 12 | Aboalsoud 2020 | *Reprod Sci* | 2020 | [12] | — | Non-published (conference abstract/poster) |

| | | | | | | |
|---|---|---|---|---|---|---|
| 13 | Gabr 2015 | *Cytotherapy* | 2015 | [13] | — | Non-published (conference abstract/poster) |
| 14 | Mohamed 2015 | *Fertil Steril* | 2015 | [14] | 10.1016/j.fertnstert.2015.07.010 | Non-published (conference abstract/poster) |
| 15 | Park 2021 | *Reprod Sci* | 2021 | [15] | — | Non-published (conference abstract/poster) |
| 16 | Sun 2019 | *Fertil Steril* | 2019 | [16] | 10.1016/j.fertnstert.2019.07.1287 | Non-published (conference abstract/poster) |
| 17 | Munan 2023 | *Chin J Tissue Eng Res* | 2023 | [17] | 10.12307/2022.722 | Language other than Spanish/English |
| 18 | Peng 2018 | *Zhong Nan Da Xue Xue Bao Yi Xue Ban* | 2018 | [18] | 10.11817/j.issn.1672-7347.2018.01.002 | Language other than Spanish/English |
| 19 | Salehi 2023 | *Iran J Obstet Gynecol Infertil* | 2023 | [19] | 10.22038/IJOGI.2023.74197.5815 | Language other than Spanish/English |
| 20 | Ye 2015 | *Chin J Tissue Eng Res* | 2015 | [20] | 10.3969/j.issn.2095-4344.2015.10.022 | Language other than Spanish/English |
| 21 | Alsaab 2025 | *J Reprod Immunol* | 2025 | [21] | 10.1016/j.jri.2024.104403 | Non-original research |
| 22 | Martirosyan 2023 | *Life (Basel)* | 2023 | [22] | 10.3390/life13122247 | Non-original research |
| 23 | Park 2024 | *Am J Obstet Gynecol* | 2024 | [23] | 10.1016/j.ajog.2024.02.023 | Wrong intervention |
| 24 | Sadeghi 2024 | *J Reprod Infertil* | 2024 | [24] | 10.18502/jri.v25i1.15192 | Non-original research |
| 25 | Chen 2018 | *Med J Chin People's Liberation Army* | 2018 | [25] | 10.11855/j.issn.0577-7402.2018.05.08 | Studies without available data |
| 26 | Wang 2019 | *Med J Chin People's Liberation Army* | 2019 | [26] | 10.11855/j.issn.0577-7402.2019.03.03 | Studies without available data |
| 27 | Huang 2024 | *Phytomedicine* | 2024 | [27] | 10.1016/j.phymed.2024.155630 | Wrong comparator |
| 28 | Ashour 2019 | *Reprod Sci* | 2019 | [28] | — | Wrong population (non-animal) |

### Supplemental Figures

#### Forest Plots

| Study or Subgroup | BM-MSCs Therapy Mean | SD | Total | Control Mean | SD | Total | Weight | Std. mean difference IV, Random, 95% CI |
|---|---|---|---|---|---|---|---|---|
| Afifi 2013[a] | 21.3 | 2.43 | 8 | 39.1 | 4.55 | 7 | 12.9% | -4.69 [-6.90 , -2.49] |
| Atwa 2025[a] | 0.62 | 0.03 | 7 | 0.9 | 0.08 | 7 | 13.0% | -4.34 [-6.50 , -2.17] |
| Elsherbiny 2024[b] | 8.12 | 6.86 | 10 | 28.18 | 26.56 | 10 | 15.6% | -0.99 [-1.93 , -0.05] |
| Li 2024[b] | 6.7 | 2.32 | 12 | 8.67 | 3.75 | 12 | 15.8% | -0.61 [-1.43 , 0.21] |
| Lu 2024[a] | 69.6 | 22.34 | 9 | 83.15 | 11.35 | 8 | 15.5% | -0.71 [-1.70 , 0.28] |
| Xing 2023[b] | 6.8 | 0.06 | 6 | 9.15 | 0.06 | 6 | 0.9% | -36.15 [-53.84 , -18.47] |
| Yamchi 2023[a] | 2.78 | 0.07 | 3 | 3.23 | 0.13 | 3 | 9.4% | -3.45 [-7.14 , 0.25] |
| Yang 2020[c] | 6.34 | 0.4 | 20 | 10.92 | 1.42 | 10 | 14.4% | -5.12 [-6.71 , -3.54] |
| Zhang 2024[b] | 5.96 | 0.13 | 6 | 8.81 | 0.13 | 6 | 2.6% | -20.24 [-30.18 , -10.29] |
| **Total (Wald[d])** | | | **81** | | | **69** | **100.0%** | **-3.44 [-5.17 , -1.70]** |

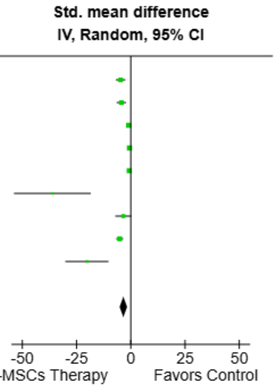


Test for overall effect: Z = 3.89 (P = 0.0001)

Heterogeneity: Tau² (DL[e]) = 4.79; Chi² = 72.10, df = 8 (P < 0.00001); I² = 89%

**Footnotes**
[a]BM-MSCs Transplantation
[b]BM-MSCs Secretome
[c]BM-MSCs Transplantation and Secretome
[d]CI calculated by Wald-type method.
[e]Tau² calculated by DerSimonian and Laird method.

**Supplemental Fig. 1. Forest plot of the pooled effect of BM-MSC therapy on serum luteinizing hormone levels.** Forest plot showing the standardized mean difference (SMD) in serum luteinizing hormone (LH) levels between animals treated with bone marrow-derived mesenchymal stem cell (BM-MSC)-based therapy and control groups. Effect estimates were pooled using a random-effects inverse-variance model and are presented with 95% confidence intervals (CIs). Individual study estimates are shown as squares, with the size of each square reflecting the study weight, and the pooled effect estimate is shown as a diamond. BM-MSC-based therapy was associated with significantly lower serum LH levels compared with controls (SMD = −3.44, 95% CI −5.17 to −1.70; P = 0.0001). Substantial statistical heterogeneity was observed across studies (I² = 89%). Values to the left of the line of no effect favor BM-MSC therapy. BM-MSC therapy included BM-MSC transplantation, and BM-MSC secretome.

| Study or Subgroup | BM-MSCs Therapy Mean | SD | Total | Control Mean | SD | Total | Weight | Std. mean difference IV, Random, 95% CI |
|---|---|---|---|---|---|---|---|---|
| Afifi 2013[a] | 30.2 | 10.4 | 8 | 10.3 | 2.6 | 7 | 6.1% | 2.39 [0.97 , 3.81] |
| Bahrehbar 2020[a] | 77.95 | 15.25 | 5 | 59.01 | 10.3 | 5 | 6.1% | 1.31 [-0.13 , 2.76] |
| Besikcioglu 2019[a] | 26.18 | 5.38 | 12 | 19.91 | 5.24 | 12 | 8.2% | 1.14 [0.27 , 2.01] |
| Chen 2018[a] | 383.54 | 5.7 | 5 | 360.76 | 9.49 | 5 | 4.6% | 2.63 [0.70 , 4.56] |
| El-Derany 2021[a] | 10.24 | 1.99 | 5 | 4.84 | 1.74 | 5 | 4.6% | 2.61 [0.69 , 4.53] |
| Fu 2008[a] | 383.57 | 32.27 | 5 | 364.33 | 34.95 | 5 | 6.7% | 0.52 [-0.76 , 1.79] |
| Fu 2017[a] | 357.32 | 19.51 | 5 | 357.32 | 12 | 5 | 6.8% | 0.00 [-1.24 , 1.24] |
| Khanmohammadi 2018[b] | 36.42 | 8.38 | 20 | 21.5 | 4.33 | 10 | 8.0% | 1.98 [1.05 , 2.91] |
| Lu 2024[a] | 65.02 | 11.61 | 6 | 26.01 | 13.93 | 6 | 5.0% | 2.81 [1.03 , 4.59] |
| Ma 2023[a] | 10.04 | 1.81 | 5 | 1.52 | 0.26 | 5 | 1.9% | 5.95 [2.38 , 9.52] |
| Park 2021b[a] | 4.34 | 2.13 | 3 | 1.54 | 1.21 | 3 | 4.3% | 1.29 [-0.74 , 3.32] |
| Sameni 2019[a] | 32.94 | 4.92 | 10 | 23.83 | 3.07 | 10 | 7.1% | 2.13 [0.98 , 3.27] |
| Tang 2024[c] | 210.82 | 38.57 | 8 | 104.3 | 12.4 | 4 | 4.7% | 2.98 [1.10 , 4.87] |
| Xie 2024[d] | 4.74 | 0.56 | 5 | 3.28 | 0.3 | 5 | 4.2% | 2.94 [0.87 , 5.00] |
| Xing 2023[d] | 14.23 | 1.29 | 6 | 5.98 | 1.55 | 6 | 2.8% | 5.34 [2.50 , 8.18] |
| Yang 2020[b] | 15.17 | 1.9 | 20 | 6.03 | 3.16 | 10 | 6.7% | 3.74 [2.47 , 5.01] |
| Yang 2024[c] | 5.26 | 1.43 | 10 | 3.05 | 1.03 | 5 | 6.7% | 1.58 [0.32 , 2.83] |
| Zhang 2024[d] | 5.97 | 0.89 | 6 | 2.98 | 1.48 | 6 | 5.6% | 2.26 [0.68 , 3.84] |
| **Total (Wald[e])** | | | **144** | | | **114** | **100.0%** | **2.11 [1.56 , 2.66]** |

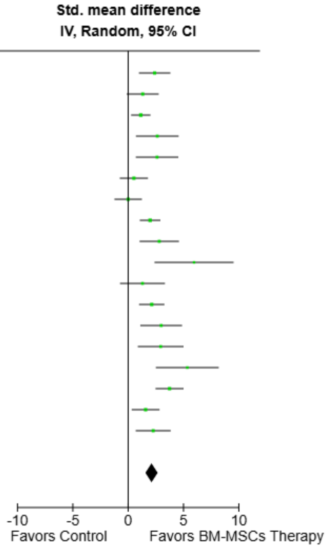


Test for overall effect: Z = 7.47 (P < 0.00001)

Heterogeneity: Tau² (DL[f]) = 0.77; Chi² = 41.48, df = 17 (P = 0.0008); I² = 59%

**Footnotes**
[a]BM-MSCs Transplantation
[b]BM-MSCs Transplantation and Secretome
[c]BM-MSCs Secretome and miR-21 Exosomes
[d]BM-MSCs Secretome
[e]CI calculated by Wald-type method.
[f]Tau² calculated by DerSimonian and Laird method.

**Supplemental Fig. 2. Forest plot of the pooled effect of BM-MSC therapy on primordial follicle count.** Forest plot showing the standardized mean difference (SMD) in primordial follicle count between animals treated with bone marrow–derived mesenchymal stem cell (BM-MSC)-based therapy and control groups. Effect estimates were pooled using a random-effects inverse-variance model and are presented with 95% confidence intervals (CIs). Individual study estimates are shown as squares, with the size of each square reflecting the study weight, and the pooled effect estimate is shown as a diamond. BM-MSC-based therapy was associated with a significantly higher primordial follicle count compared with controls (SMD = 2.11, 95% CI 1.56 to 2.66; P < 0.00001). Moderate statistical heterogeneity was observed across studies ($I^2$ = 59%). Values to the right of the line of no effect favor BM-MSC therapy. BM-MSC therapy included BM-MSC transplantation, BM-MSC secretome and BM-MSCs miR-21 exosomes.

| Study or Subgroup | BM-MSCs Therapy Mean | SD | Total | Control Mean | SD | Total | Weight | Std. mean difference IV, Fixed, 95% CI |
|---|---|---|---|---|---|---|---|---|
| Bahrehbar 2020[a] | 51.41 | 7.62 | 5 | 32.57 | 12.21 | 5 | 4.3% | 1.67 [0.12 , 3.23] |
| Chen 2018[a] | 122.5 | 4.16 | 5 | 115 | 3.33 | 5 | 4.0% | 1.80 [0.20 , 3.40] |
| El-Derany 2021[a] | 1.81 | 0.8 | 5 | 0.78 | 0.34 | 5 | 4.6% | 1.51 [0.01 , 3.02] |
| Fu 2008[a] | 132.41 | 29.92 | 5 | 119.02 | 29.94 | 5 | 6.5% | 0.40 [-0.86 , 1.66] |
| Fu 2017[a] | 117.09 | 14.33 | 5 | 116.6 | 7.41 | 5 | 6.7% | 0.04 [-1.20 , 1.28] |
| Khanmohammadi 2018[b] | 29.83 | 6.74 | 20 | 22.5 | 9.67 | 10 | 16.2% | 0.91 [0.12 , 1.71] |
| Lu 2024[a] | 20.77 | 10.46 | 6 | 7.85 | 7.08 | 6 | 6.1% | 1.34 [0.03 , 2.64] |
| Ma 2023[a] | 3.48 | 1 | 5 | 2.06 | 0.4 | 5 | 4.2% | 1.68 [0.12 , 3.24] |
| Park 2021b[a] | 3.93 | 1.65 | 3 | 2.89 | 1.65 | 3 | 3.7% | 0.50 [-1.17 , 2.18] |
| Sameni 2019[a] | 22.77 | 3.17 | 10 | 20.85 | 6.24 | 10 | 13.2% | 0.37 [-0.51 , 1.26] |
| Tang 2024[c] | 139.9 | 31.84 | 8 | 65.2 | 9.45 | 4 | 3.5% | 2.54 [0.81 , 4.27] |
| Xie 2024[d] | 4.02 | 0.8 | 5 | 0.98 | 0.48 | 5 | 1.5% | 4.16 [1.51 , 6.81] |
| Xing 2023[d] | 11.04 | 1.26 | 6 | 5.29 | 1.73 | 6 | 2.5% | 3.51 [1.45 , 5.56] |
| Yang 2020[b] | 10.02 | 3.39 | 20 | 5.35 | 3.73 | 10 | 14.7% | 1.30 [0.46 , 2.13] |
| Yang 2024[c] | 4.95 | 1.08 | 10 | 2.68 | 0.65 | 5 | 5.2% | 2.21 [0.79 , 3.62] |
| Zhang 2024[d] | 5.11 | 0.8 | 6 | 2.58 | 0.89 | 6 | 3.3% | 2.76 [1.00 , 4.52] |
| **Total** | | | **124** | | | **95** | **100.0%** | **1.24 [0.92 , 1.56]** |

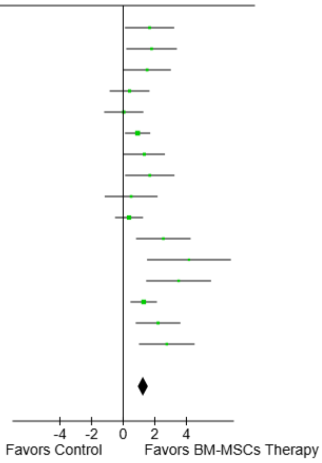


Test for overall effect: Z = 7.57 (P < 0.00001)

Heterogeneity: Chi² = 27.81, df = 15 (P = 0.02); I² = 46%

**Footnotes**
[a]BM-MSCs Transplantation
[b]BM-MSCs Transplantation and Secretome
[c]BM-MSCs Secretome and miR-21 Exosomes
[d]BM-MSCs Secretome

**Supplemental Fig. 3. Forest plot of the pooled effect of BM-MSC therapy on primary follicle count.** Forest plot showing the standardized mean difference (SMD) in primary follicle count between animals treated with bone marrow–derived mesenchymal stem cell (BM-MSC)-based therapy and control groups. Effect estimates were pooled using a fixed-effect inverse-variance model and are presented with 95% confidence intervals (CIs). Individual study estimates are shown as squares, with the size of each square reflecting the study weight, and the pooled effect estimate is shown as a diamond. BM-MSC-based therapy was associated with a significantly higher primary follicle count compared with controls (SMD = 1.24, 95% CI 0.92 to 1.56; P < 0.00001). Moderate statistical heterogeneity was observed across studies (I² = 46%). Values to the right of the line of no effect favor BM-MSC therapy. BM-MSC therapy included BM-MSC transplantation, BM-MSC secretome, and BM-MSC miR-21 exosomes.

| Study or Subgroup | BM-MSCs Therapy Mean | SD | Total | Control Mean | SD | Total | Weight | Std. mean difference IV, Random, 95% CI |
|---|---|---|---|---|---|---|---|---|
| Chen 2018[a] | 51.18 | 2.03 | 5 | 44.11 | 2.02 | 5 | 5.3% | 3.15 [0.99 , 5.32] |
| El-Derany 2021[a] | 2.82 | 0.76 | 5 | 0.78 | 0.87 | 5 | 6.4% | 2.26 [0.48 , 4.03] |
| Fu 2008[a] | 51.98 | 8.47 | 5 | 45.94 | 14.78 | 5 | 8.3% | 0.45 [-0.81 , 1.72] |
| Fu 2017[a] | 47.33 | 10.29 | 5 | 42.18 | 3.17 | 5 | 8.2% | 0.61 [-0.68 , 1.90] |
| Khanmohammadi 2018[b] | 28.34 | 5.53 | 20 | 21.67 | 6.17 | 10 | 10.1% | 1.13 [0.31 , 1.95] |
| Ma 2023[a] | 1.3 | 0.81 | 5 | 1.47 | 0.5 | 5 | 8.4% | -0.23 [-1.47 , 1.02] |
| Park 2021b[a] | 4.12 | 1.4 | 3 | 3.2 | 1.69 | 3 | 6.8% | 0.47 [-1.19 , 2.14] |
| Sameni 2019[a] | 30.79 | 3.17 | 10 | 17.88 | 6.19 | 10 | 8.4% | 2.51 [1.28 , 3.75] |
| Tang 2024[c] | 123.63 | 9.08 | 8 | 43.56 | 6.82 | 4 | 1.9% | 8.73 [4.30 , 13.16] |
| Xie 2024[d] | 2.23 | 0.29 | 5 | 0.7 | 0.45 | 5 | 4.7% | 3.65 [1.25 , 6.05] |
| Xing 2023[d] | 8.31 | 1.77 | 6 | 2.97 | 1.33 | 6 | 6.0% | 3.15 [1.24 , 5.06] |
| Yang 2020[b] | 7.84 | 5.4 | 20 | 3.98 | 3.35 | 10 | 10.2% | 0.78 [-0.01 , 1.56] |
| Yang 2024[c] | 4.02 | 1.2 | 10 | 2.68 | 0.82 | 5 | 8.7% | 1.15 [-0.03 , 2.33] |
| Zhang 2024[d] | 4.09 | 0.8 | 6 | 1.72 | 0.83 | 6 | 6.6% | 2.68 [0.95 , 4.42] |
| **Total (Wald[e])** | | | **113** | | | **84** | **100.0%** | **1.62 [0.95 , 2.28]** |

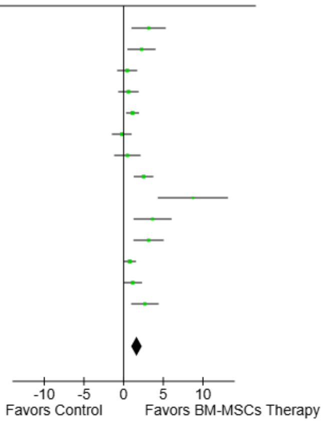


Test for overall effect: Z = 4.77 (P < 0.00001)

Heterogeneity: Tau² (DL[f]) = 0.97; Chi² = 39.66, df = 13 (P = 0.0002); I² = 67%

**Footnotes**
[a]BM-MSCs Transplantation
[b]BM-MSCs Transplantation and Secretome
[c]BM-MSCs Secretome and miR-21 Exosomes
[d]BM-MSCs Secretome
[e]CI calculated by Wald-type method.
[f]Tau² calculated by DerSimonian and Laird method.

**Supplemental Fig. 4. Forest plot of the pooled effect of BM-MSC therapy on secondary follicle count.** Forest plot showing the standardized mean difference (SMD) in secondary follicle count between animals treated with bone marrow–derived mesenchymal stem cell (BM-MSC)-based therapy and control groups. Effect estimates were pooled using a random-effects inverse-variance model and are presented with 95% confidence intervals (CI). Individual study estimates are shown as squares, with the size of each square reflecting the study weight, and the pooled effect estimate is shown as a diamond. BM-MSC-based therapy was associated with a significantly higher secondary follicle count compared with controls (SMD = 1.62, 95% CI 0.95 to 2.28; P < 0.00001). Significant statistical heterogeneity was observed across studies ($I^2$ = 67%). Values to the right of the line of no effect favor BM-MSC therapy. BM-MSC therapy included BM-MSC transplantation , BM-MSC secretome, and BM-MSC miR-21 exosomes.

| | BM-MSCs Therapy | | | Control | | | | Std. mean difference |
|---|---|---|---|---|---|---|---|---|
| **Study or Subgroup** | **Mean** | **SD** | **Total** | **Mean** | **SD** | **Total** | **Weight** | **IV, Fixed, 95% CI** |
| Bahrehbar 2020[a] | 42.64 | 10.98 | 5 | 20 | 5.9 | 5 | 24.1% | 2.32 [0.52 , 4.12] |
| Kilic 2014[a] | 3.9 | 0.83 | 11 | 3.1 | 0.69 | 7 | 75.9% | 0.98 [-0.04 , 1.99] |
| **Total** | | | **16** | | | **12** | **100.0%** | **1.30 [0.42 , 2.18]** |

Std. mean difference IV, Fixed, 95% CI

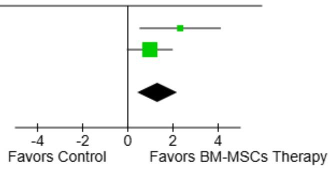


Test for overall effect: Z = 2.88 (P = 0.004)

Heterogeneity: $Chi^2$ = 1.62, df = 1 (P = 0.20); $I^2$ = 38%

**Footnotes**
[a]BM-MSCs Transplantation

**Supplemental Fig. 5. Forest plot of the pooled effect of BM-MSC therapy on preantral follicle count.** Forest plot showing the standardized mean difference (SMD) in preantral follicle count between animals treated with bone marrow–derived mesenchymal stem cell (BM-MSC)-based therapy and control groups. Effect estimates were pooled using a fixed-effect inverse-variance model and are presented with 95% confidence intervals (CI). Individual study estimates are shown as squares, with the size of each square reflecting the study weight, and the pooled effect estimate is shown as a diamond. BM-MSC-based therapy was associated with a significantly higher preantral follicle count compared with controls (SMD = 1.30, 95% CI 0.42 to 2.18; P = 0.004). No significant statistical heterogeneity was observed across studies ($I^2$ = 38%). Values to the right of the line of no effect favor BM-MSC therapy. BM-MSC therapy included exclusively BM-MSC transplantation.

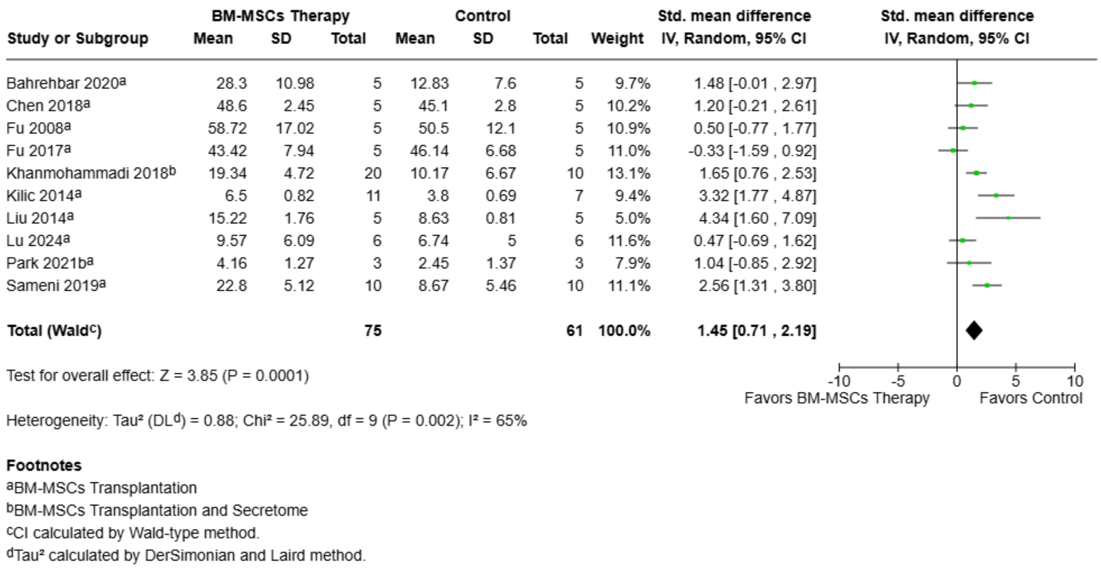


**Supplemental Fig. 6. Forest plot of the pooled effect of BM-MSC therapy on antral follicle count.** Forest plot showing the standardized mean difference (SMD) in antral follicle count between animals treated with bone marrow–derived mesenchymal stem cell (BM-MSC)-based therapy and control groups. Effect estimates were pooled using a random-effects inverse-variance model and are presented with 95% confidence intervals (CI). Individual study estimates are shown as squares, with the size of each square reflecting the study weight, and the pooled effect estimate is shown as a diamond. BM-MSC-based therapy was associated with a significantly higher antral follicle count compared with controls (SMD = 1.45, 95% CI 0.71 to 2.19; P = 0.0001). Significant statistical heterogeneity was observed across studies ($I^2$ = 65%). Values to the right of the line of no effect favor BM-MSC therapy. BM-MSC therapy included BM-MSC transplantation and BM-MSC secretome.

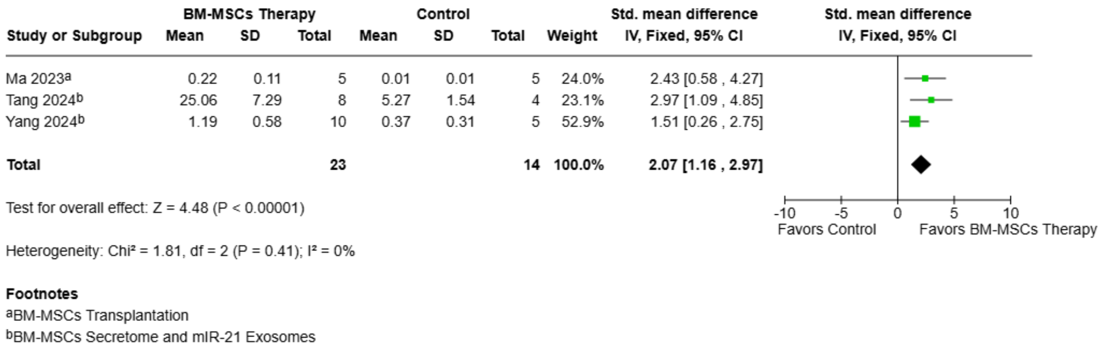


**Supplemental Fig. 7. Forest plot of the pooled effect of BM-MSC therapy on mature follicle count.** Forest plot showing the standardized mean difference (SMD) in mature follicle count between animals treated with bone marrow–derived mesenchymal stem cell (BM-MSC)-based therapy and control groups. Effect estimates were pooled using a fixed-effect inverse-variance model and are presented with 95% confidence intervals (CI). Individual study estimates are shown as squares, with the size of each square reflecting the study weight, and the pooled effect estimate is shown as a diamond. BM-MSC-based therapy was associated with a significantly higher mature follicle count compared with controls (SMD = 2.07, 95% CI 1.16 to 2.97; P < 0.00001). No significant statistical heterogeneity was observed across studies ($I^2$ =

0%). Values to the right of the line of no effect favor BM-MSC therapy. BM-MSC therapy included BM-MSC transplantation, BM-MSC secretome, and BM-MSC miR-21 exosomes.

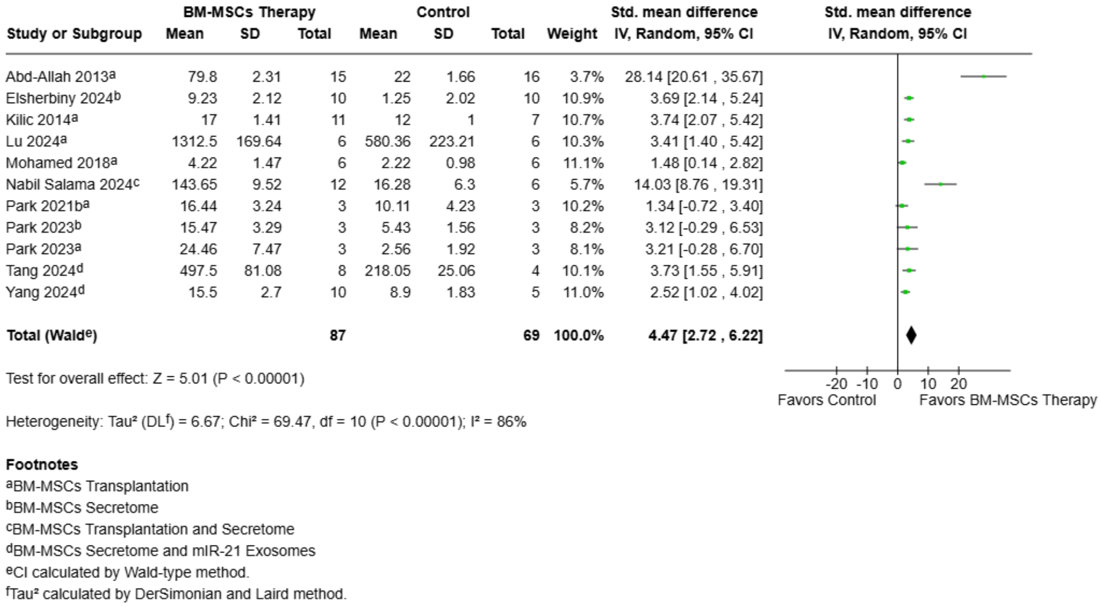


**Supplemental Fig. 8. Forest plot of the pooled effect of BM-MSC therapy on total follicle count.** Forest plot showing the standardized mean difference (SMD) in total follicle count between animals treated with bone marrow–derived mesenchymal stem cell (BM-MSC)-based therapy and control groups. Effect estimates were pooled using a random-effects inverse-variance model and are presented with 95% confidence intervals (CI). Individual study estimates are shown as squares, with the size of each square reflecting the study weight, and the pooled effect estimate is shown as a diamond. BM-MSC-based therapy was associated with a significantly higher total follicle count compared with controls (SMD = 4.47, 95% CI 2.72 to 6.22; P < 0.00001). Significant statistical heterogeneity was observed across studies ($I^2$ = 86%). Values to the right of the line of no effect favor BM-MSC therapy. BM-MSC therapy included BM-MSC transplantation, BM-MSC secretome, and BM-MSC miR-21 exosomes.

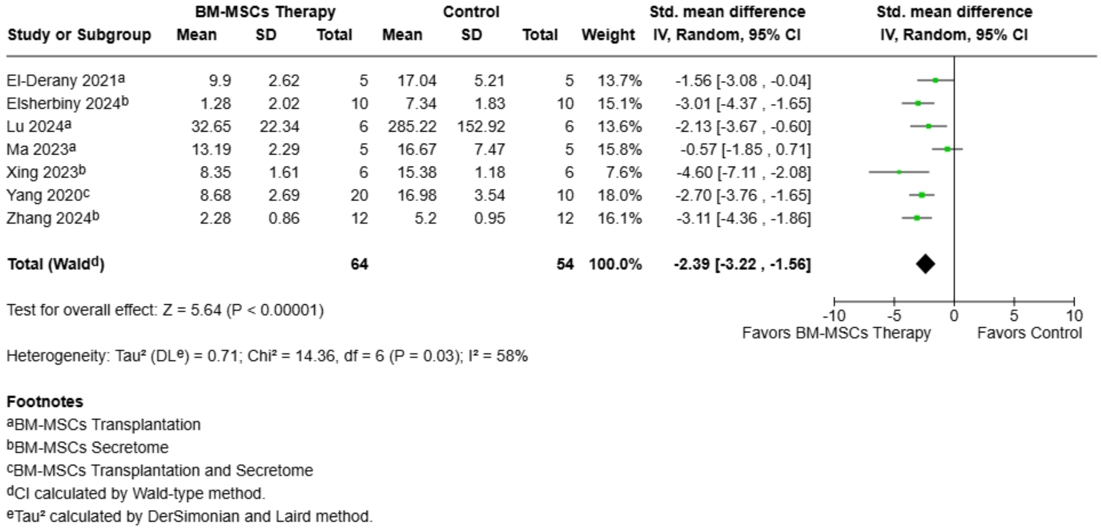

**Supplemental Fig. 9. Forest plot of the pooled effect of BM-MSC therapy on atretic follicle count.** Forest plot showing the standardized mean difference (SMD) in atretic follicle count between animals treated with bone marrow–derived mesenchymal stem cell (BM–MSC)–based therapy and control groups. Effect estimates were pooled using a random–effects inverse–variance model and are presented with 95% confidence intervals (CI). Individual study estimates are shown as squares, with the size of each square reflecting the study weight, and the pooled effect estimate is shown as a diamond. BM–MSC–based therapy was associated with a significantly lower atretic follicle count compared with controls (SMD = −2.39, 95% CI −3.22 to −1.56; P < 0.00001). Significant statistical heterogeneity was observed across studies ($I^2$ = 58%). Values to the left of the line of no effect favor BM–MSC therapy. BM–MSC therapy included BM–MSC transplantation, and BM–MSC secretome.

| | BM-MSCs Therapy | | | Control | | | | Std. mean difference |
|---|---|---|---|---|---|---|---|---|
| Study or Subgroup | Mean | SD | Total | Mean | SD | Total | Weight | IV, Fixed, 95% CI |
| Park 2021b[a] | 1.67 | 0.07 | 3 | 0.79 | 0.24 | 3 | 20.1% | 3.98 [-0.18 , 8.15] |
| Park 2023[b] | 1.09 | 0.38 | 3 | 0.48 | 0.09 | 3 | 63.8% | 1.77 [-0.57 , 4.11] |
| Park 2023[a] | 1.9 | 0.27 | 3 | 0.79 | 0.06 | 3 | 16.0% | 4.54 [-0.13 , 9.21] |
| **Total** | | | **9** | | | **9** | **100.0%** | **2.66 [0.79 , 4.53]** |

Test for overall effect: Z = 2.79 (P = 0.005)

Heterogeneity: $Chi^2$ = 1.57, df = 2 (P = 0.46); $I^2$ = 0%

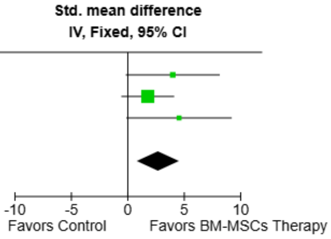


**Footnotes**
[a]BM-MSCs Transplantation
[b]BM-MSCs Secretome

**Supplemental Fig. 10. Forest plot of the pooled effect of BM-MSC therapy on ovarian size.** Forest plot showing the standardized mean difference (SMD) in ovarian size between animals treated with bone marrow–derived mesenchymal stem cell (BM-MSC)-based therapy and control groups. Effect estimates were pooled using a fixed-effect inverse-variance model and are presented with 95% confidence intervals (CI). Individual study estimates are shown as squares, with the size of each square reflecting the study weight, and the pooled effect estimate is shown as a diamond. BM-MSC-based therapy was associated with a significantly larger ovarian size compared with controls (SMD = 2.66, 95% CI 0.79 to 4.53; P = 0.005). No significant statistical heterogeneity was observed across studies ($I^2$ = 0%). Values to the right of the line of no effect favor BM-MSC therapy. BM-MSC therapy included BM-MSC transplantation and BM-MSC secretome.

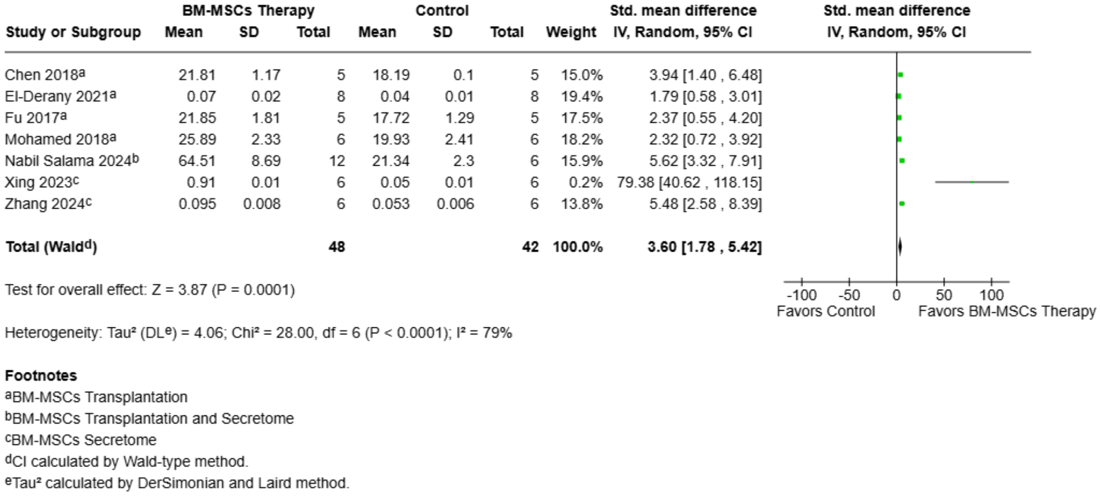


**Supplemental Fig. 11. Forest plot of the pooled effect of BM-MSC therapy on ovarian weight.** Forest plot showing the standardized mean difference (SMD) in ovarian weight between animals treated with bone marrow–derived mesenchymal stem cell (BM-MSC)-based therapy and control groups. Effect estimates were pooled using a random-effects inverse-variance model and are presented with 95% confidence intervals (CIs). Individual study estimates are shown as squares, with the size of each square reflecting the study weight, and the pooled effect estimate is shown as a diamond. BM-MSC-based therapy was associated with a significantly higher ovarian weight compared with controls (SMD = 3.60, 95% CI 1.78 to 5.42; P = 0.0001). Substantial heterogeneity was observed across studies ($I^2$ = 79%). Values to the right of the line of no effect favor BM-MSC therapy. BM-MSC therapy included BM-MSC transplantation, and BM-MSC secretome.

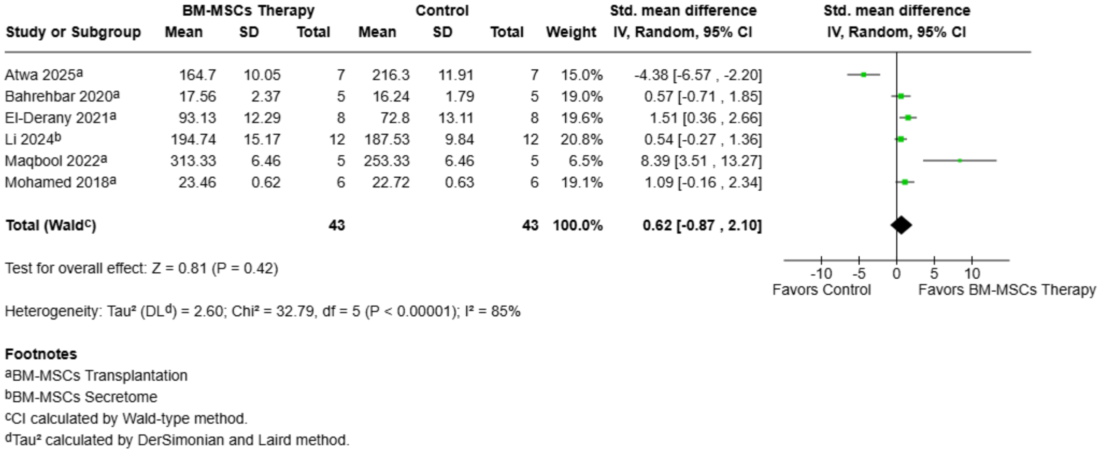


**Supplemental Fig. 12. Forest plot of the pooled effect of BM-MSC therapy on body weight.** Forest plot showing the standardized mean difference (SMD) in body weight between animals treated with bone marrow–derived mesenchymal stem cell (BM-MSC)-based therapy and control groups. Effect estimates were pooled using a random-effects inverse-variance model and are presented with 95% confidence intervals (CIs). Individual study estimates are shown as squares, with the size of each square reflecting the study weight, and the pooled effect estimate is shown as a diamond. BM-MSC-based therapy was not associated with a significant difference in body weight compared with controls (SMD = 0.62, 95% CI −0.87 to 2.10; P = 0.42). Significant heterogeneity was observed across studies ($I^2$ = 85%). Values to the right of the line of no effect favor BM-MSC therapy. BM-MSC therapy included BM-MSC transplantation and BM-MSC secretome.

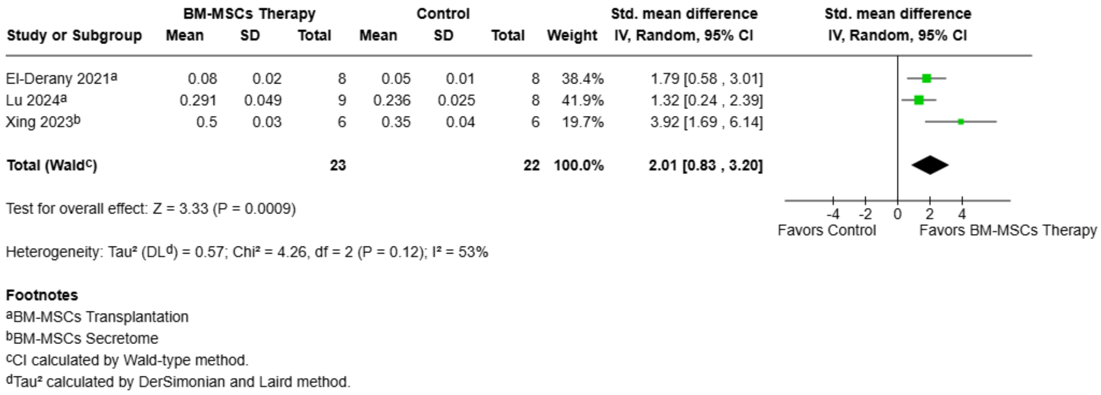


**Supplemental Fig. 13. Forest plot of the pooled effect of BM-MSC therapy on ovarian weight index.** Forest plot showing the standardized mean difference (SMD) in ovarian weight index between animals treated with bone marrow–derived mesenchymal stem cell (BM-MSC)-based therapy and control groups. Effect estimates were pooled using a random-effects inverse-variance model and are presented with 95% confidence intervals (CIs). Individual study estimates are shown as squares, with the size of each square reflecting the study weight, and the pooled effect estimate is shown as a diamond. BM-MSC-based therapy was associated with a significantly higher ovarian weight index compared with controls (SMD = 2.01, 95% CI 0.83 to 3.20; P = 0.0009). Moderate heterogeneity was observed across studies ($I^2$ = 53%). Values to the right of the line of no effect favor BM-MSC therapy. BM-MSC therapy included BM-MSC transplantation and BM-MSC secretome.

#### Subgroup Analysis According to BM-MSC Therapy Type

##### BM-MSCs Transplantation (Cell therapy)

| Study or Subgroup | BM-MSCs Transplantation Mean | SD | Total | Control Mean | SD | Total | Weight | Std. mean difference IV, Random, 95% CI |
|---|---|---|---|---|---|---|---|---|
| Abd-Allah 2013 | 47.03 | 0.91 | 15 | 21 | 1.19 | 15 | 1.5% | 23.91 [17.38 , 30.44] |
| Afifi 2013 | 23 | 3.39 | 8 | 14 | 2.36 | 7 | 5.5% | 2.86 [1.30 , 4.43] |
| Atwa 2025 | 36.2 | 1.32 | 7 | 16.9 | 2.65 | 7 | 3.0% | 8.63 [4.72 , 12.54] |
| Badawy 2017 | 249.41 | 6.91 | 5 | 161.22 | 9.11 | 5 | 1.8% | 9.85 [4.17 , 15.54] |
| Bahrehbar 2020 | 74.35 | 10.62 | 5 | 32.54 | 7.96 | 5 | 4.3% | 4.02 [1.44 , 6.61] |
| Bao 2018 | 26.4 | 2.55 | 8 | 17.6 | 1.98 | 8 | 5.3% | 3.64 [1.89 , 5.40] |
| Chen 2018 | 35.96 | 1.98 | 15 | 29.56 | 2.29 | 15 | 6.0% | 2.91 [1.84 , 3.97] |
| El-Derany 2021 | 5.73 | 0.18 | 8 | 4.45 | 0.24 | 8 | 4.4% | 5.70 [3.23 , 8.18] |
| Fu 2008 | 38.13 | 38.42 | 15 | 24.7 | 28.93 | 15 | 6.3% | 0.38 [-0.34 , 1.11] |
| Fu 2017 | 34.92 | 9.85 | 15 | 26.72 | 7 | 15 | 6.3% | 0.93 [0.17 , 1.69] |
| Gabr 2016 | 43.55 | 1.94 | 6 | 29.88 | 2.83 | 6 | 4.1% | 5.20 [2.42 , 7.98] |
| Khanmohammadi 2018 | 35.86 | 3.81 | 10 | 20.89 | 5.65 | 10 | 5.7% | 2.98 [1.62 , 4.33] |
| Liu 2014 | 52.64 | 3.53 | 5 | 35.85 | 2.28 | 5 | 3.7% | 5.10 [1.97 , 8.23] |
| Lu 2024 | 7.62 | 2.06 | 9 | 5.12 | 1.07 | 8 | 6.0% | 1.42 [0.32 , 2.52] |
| Ma 2023 | 166.47 | 31.77 | 5 | 53.53 | 2.35 | 5 | 4.0% | 4.53 [1.69 , 7.36] |
| Maqbool 2022 | 13 | 2.24 | 5 | 5.67 | 1.3 | 5 | 4.5% | 3.62 [1.23 , 6.00] |
| Mohamed 2018 | 13.47 | 2.99 | 6 | 6.97 | 3.39 | 6 | 5.6% | 1.88 [0.42 , 3.33] |
| Nabil Salama 2024 | 117.89 | 7.78 | 6 | 55.05 | 3.56 | 6 | 2.3% | 9.59 [4.77 , 14.40] |
| Park 2023 | 5.75 | 2 | 3 | 4.18 | 0.79 | 3 | 5.2% | 0.83 [-0.96 , 2.61] |
| Sameni 2019 | 34 | 5.06 | 10 | 23.98 | 6.32 | 10 | 6.0% | 1.68 [0.63 , 2.73] |
| Yamchi 2023 | 4.45 | 0.7 | 3 | 1.78 | 0.01 | 3 | 2.5% | 4.31 [-0.15 , 8.78] |
| Yang 2020 | 49.05 | 7.49 | 10 | 36.98 | 14.26 | 10 | 6.1% | 1.01 [0.07 , 1.96] |
| **Total (Wald[a])** | | | **179** | | | **177** | **100.0%** | **3.46 [2.55 , 4.37]** |

Test for overall effect: Z = 7.47 (P < 0.00001)

Heterogeneity: Tau² (DL[b]) = 3.27; Chi² = 139.71, df = 21 (P < 0.00001); I² = 85%

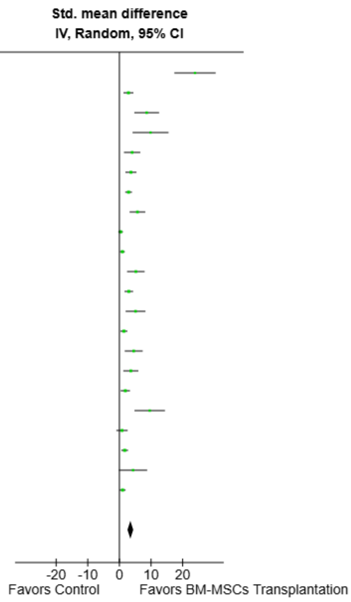


**Footnotes**
[a]CI calculated by Wald-type method.
[b]Tau² calculated by DerSimonian and Laird method.

**Supplemental Fig. 14. Forest plot of the pooled effect of BM-MSC transplantation on serum estradiol levels.** Forest plot showing the standardized mean difference (SMD) in serum estradiol (E2) levels between animals treated with bone marrow–derived mesenchymal stem cell (BM-MSC) transplantation and control groups. Effect estimates were pooled using a random-effects inverse-variance model and are presented with 95% confidence intervals (CIs). Individual study estimates are shown as squares, with the size of each square reflecting the study weight, and the pooled effect estimate is shown as a diamond. BM-MSC transplantation was associated with significantly higher serum E2 levels compared with controls (SMD = 3.46, 95% CI 2.55 to 4.37; P < 0.00001). Considerable heterogeneity was observed across studies (I² = 85%). Values to the right of the line of no effect favor BM-MSC transplantation.

| Study or Subgroup | BM-MSCs Transplantation Mean | SD | Total | Control Mean | SD | Total | Weight | Std. mean difference IV, Random, 95% CI |
|---|---|---|---|---|---|---|---|---|
| Abd-Allah 2013 | 5.9 | 0.9 | 15 | 19.9 | 1.6 | 15 | 3.8% | -10.49 [-13.43 , -7.56] |
| Afifi 2013 | 7.7 | 0.21 | 8 | 29.3 | 4.45 | 7 | 3.8% | -6.72 [-9.69 , -3.74] |
| Atwa 2025 | 0.5 | 0.11 | 7 | 0.81 | 0.08 | 7 | 5.2% | -3.02 [-4.70 , -1.33] |
| Badawy 2017 | 0.34 | 0.07 | 5 | 0.59 | 0.06 | 5 | 4.5% | -3.46 [-5.77 , -1.15] |
| Bahrehbar 2020 | 6.61 | 1.77 | 5 | 16.77 | 4.14 | 5 | 4.8% | -2.88 [-4.92 , -0.84] |
| Bao 2018 | 5.41 | 2.46 | 8 | 14.44 | 4.38 | 8 | 5.5% | -2.40 [-3.77 , -1.03] |
| Chen 2018 | 15.57 | 0.97 | 15 | 22.64 | 1.64 | 15 | 5.3% | -5.11 [-6.67 , -3.55] |
| El-Derany 2021 | 0.39 | 0.08 | 8 | 0.77 | 0.04 | 8 | 4.3% | -5.68 [-8.15 , -3.21] |
| Fu 2008 | 8.7 | 18.13 | 15 | 14.7 | 18.75 | 15 | 6.1% | -0.32 [-1.04 , 0.40] |
| Fu 2017 | 18.04 | 3.29 | 15 | 22.96 | 3.58 | 15 | 6.0% | -1.39 [-2.20 , -0.58] |
| Gabr 2016 | 13.12 | 1.41 | 6 | 20.08 | 3.58 | 6 | 5.3% | -2.36 [-3.98 , -0.75] |
| Khanmohammadi 2018 | 7.13 | 0.93 | 10 | 12.58 | 0.89 | 10 | 4.7% | -5.73 [-7.90 , -3.57] |
| Lu 2024 | 11.86 | 4.3 | 9 | 28.79 | 7.43 | 8 | 5.5% | -2.69 [-4.10 , -1.29] |
| Ma 2023 | 73.88 | 3.78 | 5 | 92.17 | 3.77 | 5 | 4.0% | -4.38 [-7.13 , -1.62] |
| Maqbool 2022 | 10.33 | 1.27 | 5 | 14.67 | 1.3 | 5 | 4.7% | -3.05 [-5.17 , -0.93] |
| Mohamed 2018 | 10 | 1.21 | 6 | 18.67 | 3.34 | 6 | 4.9% | -3.19 [-5.11 , -1.26] |
| Nabil Salama 2024 | 58.45 | 3.8 | 6 | 107.6 | 1.36 | 6 | 1.1% | -15.90 [-23.74 , -8.05] |
| Park 2023 | 14.39 | 4.88 | 3 | 23.08 | 5.26 | 3 | 4.8% | -1.37 [-3.45 , 0.71] |
| Sameni 2019 | 7.77 | 0.65 | 10 | 11.19 | 0.89 | 10 | 5.2% | -4.20 [-5.90 , -2.51] |
| Yamchi 2023 | 2.93 | 0.13 | 3 | 3.16 | 0.1 | 3 | 4.6% | -1.59 [-3.80 , 0.63] |
| Yang 2020 | 14.99 | 2.4 | 10 | 19.01 | 0.76 | 10 | 5.7% | -2.16 [-3.32 , -1.01] |
| **Total (Wald[a])** | | | **174** | | | **172** | **100.0%** | **-3.54 [-4.45 , -2.62]** |

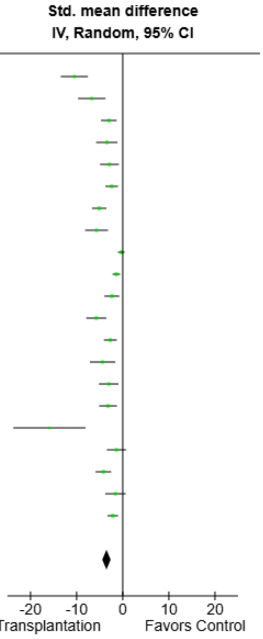


Test for overall effect: Z = 7.59 (P < 0.00001)

Heterogeneity: Tau² (DL[b]) = 3.44; Chi² = 124.71, df = 20 (P < 0.00001); I² = 84%

**Footnotes**

[a]CI calculated by Wald-type method.

[b]Tau² calculated by DerSimonian and Laird method.

**Supplemental Fig. 15. Forest plot of the pooled effect of BM-MSC transplantation on serum follicle-stimulating hormone levels.** Forest plot showing the standardized mean difference (SMD) in serum follicle-stimulating hormone (FSH) levels between animals treated with bone marrow-derived mesenchymal stem cell (BM-MSC) transplantation and control groups. Effect estimates were pooled using a random-effects inverse-variance model and are presented with 95% confidence intervals (CIs). Individual study estimates are shown as squares, with the size of each square reflecting the study weight, and the pooled effect estimate is shown as a diamond. BM-MSC transplantation was associated with significantly lower serum FSH levels compared with controls (SMD = −3.54, 95% CI −4.45 to −2.62; P < 0.00001). Considerable heterogeneity was observed across studies ($I^2$ = 84%). Values to the left of the line of no effect favor BM-MSC transplantation.

| Study or Subgroup | BM-MSCs Transplantation Mean | SD | Total | Control Mean | SD | Total | Weight | Std. mean difference IV, Random, 95% CI |
|---|---|---|---|---|---|---|---|---|
| Besikcioglu 2019 | 0.77 | 0.27 | 12 | 0.82 | 0.2 | 12 | 19.1% | -0.20 [-1.01 , 0.60] |
| Lu 2024 | 378.88 | 49.6 | 9 | 276.4 | 74.53 | 8 | 17.4% | 1.56 [0.43 , 2.68] |
| Maqbool 2022 | 29 | 8.07 | 5 | 9.67 | 1.3 | 5 | 11.9% | 3.02 [0.92 , 5.13] |
| Mohamed 2018 | 36.24 | 16.02 | 6 | 8.42 | 18.74 | 6 | 16.1% | 1.47 [0.13 , 2.81] |
| Nabil Salama 2024 | 8.52 | 0.78 | 6 | 1.93 | 0.18 | 6 | 3.5% | 10.75 [5.38 , 16.11] |
| Park 2023 | 4.78 | 2.19 | 3 | 3.29 | 0.08 | 3 | 13.7% | 0.77 [-1.00 , 2.53] |
| Yang 2020 | 145.75 | 15.81 | 10 | 131 | 11.86 | 10 | 18.4% | 1.01 [0.07 , 1.95] |
| **Total (Wald[a])** | | | **51** | | | **50** | **100.0%** | **1.49 [0.40 , 2.59]** |

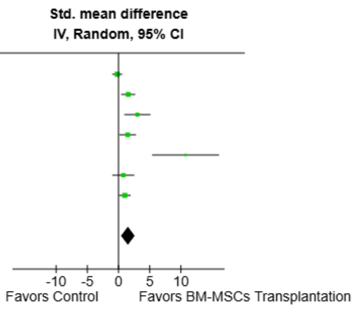


Test for overall effect: Z = 2.67 (P = 0.008)

Heterogeneity: Tau² (DL[b]) = 1.47; Chi² = 26.18, df = 6 (P = 0.0002); I² = 77%

**Footnotes**

[a]CI calculated by Wald-type method.

[b]Tau² calculated by DerSimonian and Laird method.

**Supplemental Fig. 16. Forest plot of the pooled effect of BM-MSC transplantation on serum anti-Müllerian hormone levels.** Forest plot showing the standardized mean difference (SMD) in serum anti-Müllerian hormone (AMH) levels between animals treated with bone marrow–derived mesenchymal stem cell (BM-MSC) transplantation and control groups. Effect estimates were pooled using a random-effects inverse-variance model and are presented with 95% confidence intervals (CIs). Individual study estimates are shown as squares, with the size of each square reflecting the study weight, and the pooled effect estimate is shown as a diamond. BM-MSC transplantation was associated with significantly higher serum AMH levels compared with controls (SMD = 1.49, 95% CI 0.40 to 2.59; P = 0.008). Substantial heterogeneity was observed across studies (I² = 77%). Values to the right of the line of no effect favor BM-MSC transplantation.

| Study or Subgroup | BM-MSCs Transplantation Mean | SD | Total | Control Mean | SD | Total | Weight | Std. mean difference IV, Random, 95% CI |
|---|---|---|---|---|---|---|---|---|
| Afifi 2013 | 21.3 | 2.43 | 8 | 39.1 | 4.55 | 7 | 19.4% | -4.69 [-6.90 , -2.49] |
| Atwa 2025 | 0.62 | 0.03 | 7 | 0.9 | 0.08 | 7 | 19.6% | -4.34 [-6.50 , -2.17] |
| Lu 2024 | 69.6 | 22.34 | 9 | 83.15 | 11.35 | 8 | 25.0% | -0.71 [-1.70 , 0.28] |
| Yamchi 2023 | 2.78 | 0.07 | 3 | 3.23 | 0.13 | 3 | 12.9% | -3.45 [-7.14 , 0.25] |
| Yang 2020 | 6.58 | 0.98 | 10 | 10.92 | 1.42 | 10 | 23.0% | -3.41 [-4.88 , -1.94] |
| **Total (Wald[a])** | | | **37** | | | **35** | **100.0%** | **-3.17 [-5.02 , -1.33]** |

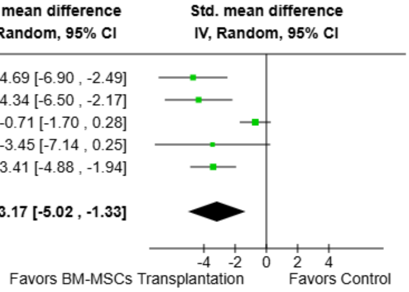


Test for overall effect: Z = 3.37 (P = 0.0007)

Heterogeneity: Tau² (DL[b]) = 3.29; Chi² = 20.35, df = 4 (P = 0.0004); I² = 80%

**Footnotes**

[a]CI calculated by Wald-type method.

[b]Tau² calculated by DerSimonian and Laird method.

**Supplemental Fig. 17. Forest plot of the pooled effect of BM-MSC transplantation on serum luteinizing hormone levels.** Forest plot showing the standardized mean difference (SMD) in serum luteinizing hormone (LH) levels between animals treated with bone marrow–derived mesenchymal stem cell (BM-MSC) transplantation and control groups. Effect estimates were pooled using a random-effects inverse-variance model and are presented with 95% confidence intervals (CIs). Individual study estimates are shown as squares, with the size of each square reflecting the study weight, and the pooled effect estimate is shown as a diamond. BM-MSC transplantation was associated with significantly lower serum LH levels compared with controls (SMD =

−3.17, 95% CI −5.02 to −1.33; P = 0.0007). Considerable heterogeneity was observed across studies (I² = 80%). Values to the left of the line of no effect favor BM-MSC transplantation.

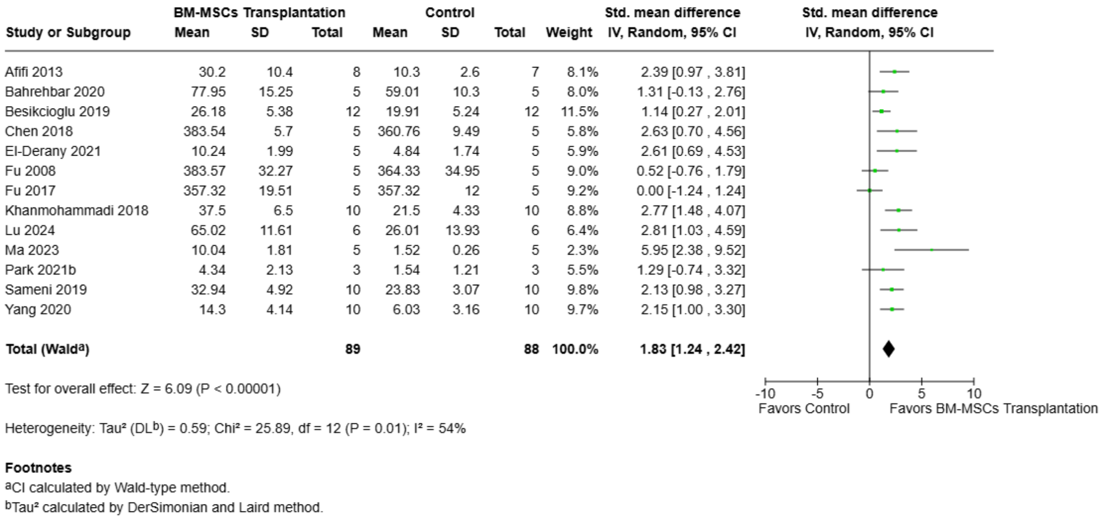


**Supplemental Fig. 18. Forest plot of the pooled effect of BM-MSC transplantation on primordial follicle count.** Forest plot showing the standardized mean difference (SMD) in primordial follicle count between animals treated with bone marrow–derived mesenchymal stem cell (BM-MSC) transplantation and control groups. Effect estimates were pooled using a random-effects inverse-variance model and are presented with 95% confidence intervals (CIs). Individual study estimates are shown as squares, with the size of each square reflecting the study weight, and the pooled effect estimate is shown as a diamond. BM-MSC transplantation was associated with a significantly higher primordial follicle count compared with controls (SMD = 1.83, 95% CI 1.24 to 2.42; P < 0.00001). Moderate heterogeneity was observed across studies (I² = 54%). Values to the right of the line of no effect favor BM-MSC transplantation.

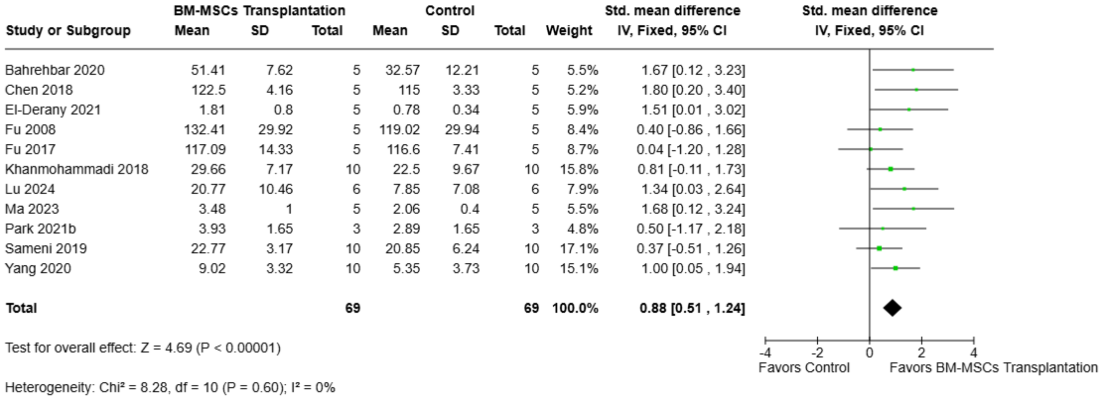


**Supplemental Fig. 19. Forest plot of the pooled effect of BM-MSC transplantation on primary follicle count.** Forest plot showing the standardized mean difference (SMD) in primary follicle count between animals treated with bone marrow–derived mesenchymal stem cell (BM-MSC) transplantation and control groups. Effect estimates were pooled using a fixed-effect inverse-variance model and are presented with 95% confidence intervals (CIs). Individual study estimates are shown as squares, with the size of each square reflecting the study weight, and the pooled effect estimate is shown as a diamond. BM-MSC transplantation was associated with a significantly higher primary follicle count compared with controls (SMD = 0.88, 95% CI 0.51 to 1.24; P < 0.00001). No statistical heterogeneity was observed across studies (I² = 0%). Values to the right of the line of no effect favor BM-MSC transplantation.

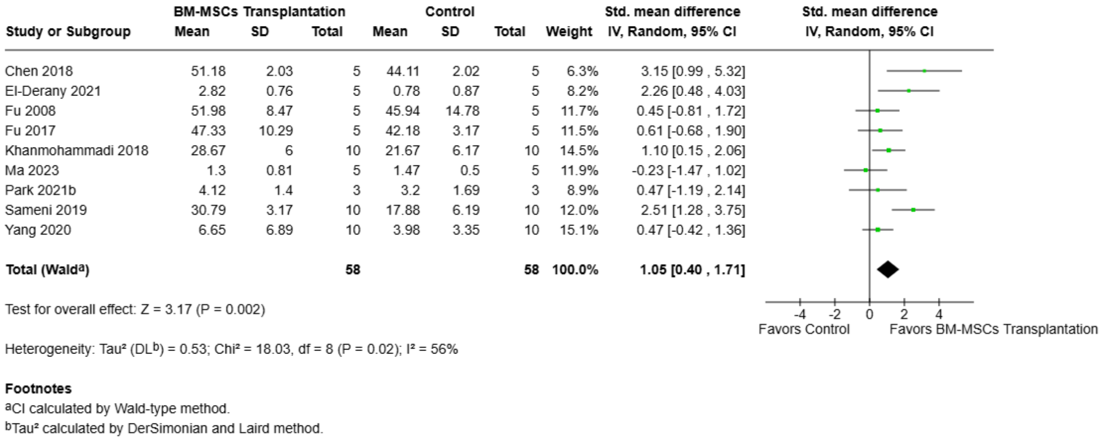


**Supplemental Fig. 20. Forest plot of the pooled effect of BM-MSC transplantation on secondary follicle count.** Forest plot showing the standardized mean difference (SMD) in secondary follicle count between animals treated with bone marrow–derived mesenchymal stem cell (BM-MSC) transplantation and control groups. Effect estimates were pooled using a random-effects inverse-variance model and are presented with 95% confidence intervals (CIs). Individual study estimates are shown as squares, with the size of each square reflecting the study weight, and the pooled effect estimate is shown as a diamond. BM-MSC transplantation was associated with a significantly higher secondary follicle count compared with controls (SMD = 1.05, 95% CI 0.40 to 1.71; P = 0.002). Moderate heterogeneity was observed across studies (I² = 56%). Values to the right of the line of no effect favor BM-MSC transplantation.

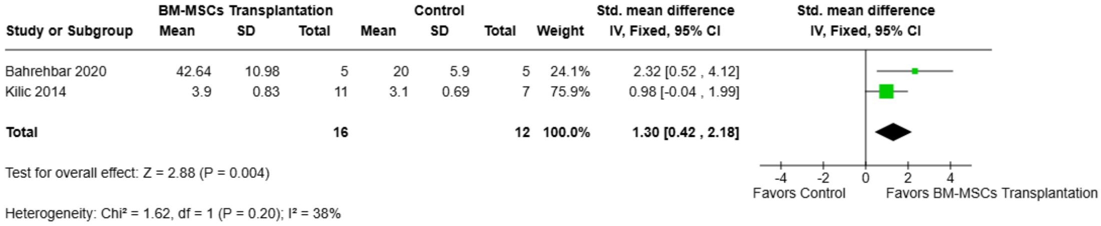


**Supplemental Fig. 21. Forest plot of the pooled effect of BM-MSC transplantation on preantral follicle count.** Forest plot showing the standardized mean difference (SMD) in preantral follicle count between animals treated with bone marrow–derived mesenchymal stem cell (BM-MSC) transplantation and control groups. Effect estimates were pooled using a fixed-effect inverse-variance model and are presented with 95% confidence intervals (CIs). Individual study estimates are shown as squares, with the size of each square reflecting the study weight, and the pooled effect estimate is shown as a diamond. BM-MSC transplantation was associated with a significantly higher preantral follicle count compared with controls (SMD = 1.30, 95% CI 0.42 to 2.18; P = 0.004). Moderate heterogeneity was observed across studies (I² = 38%). Values to the right of the line of no effect favor BM-MSC transplantation.

| Study or Subgroup | BM-MSCs Transplantation Mean | SD | Total | Control Mean | SD | Total | Weight | Std. mean difference IV, Random, 95% CI |
|---|---|---|---|---|---|---|---|---|
| Bahrehbar 2020 | 28.3 | 10.98 | 5 | 12.83 | 7.6 | 5 | 9.9% | 1.48 [-0.01 , 2.97] |
| Chen 2018 | 48.6 | 2.45 | 5 | 45.1 | 2.8 | 5 | 10.3% | 1.20 [-0.21 , 2.61] |
| Fu 2008 | 58.72 | 17.02 | 5 | 50.5 | 12.1 | 5 | 11.0% | 0.50 [-0.77 , 1.77] |
| Fu 2017 | 43.42 | 7.94 | 5 | 46.14 | 6.68 | 5 | 11.1% | -0.33 [-1.59 , 0.92] |
| Khanmohammadi 2018 | 20.17 | 4.33 | 10 | 10.17 | 6.67 | 10 | 12.2% | 1.70 [0.65 , 2.76] |
| Kilic 2014 | 6.5 | 0.82 | 11 | 3.8 | 0.69 | 7 | 9.6% | 3.32 [1.77 , 4.87] |
| Liu 2014 | 15.22 | 1.76 | 5 | 8.63 | 0.81 | 5 | 5.2% | 4.34 [1.60 , 7.09] |
| Lu 2024 | 9.57 | 6.09 | 6 | 6.74 | 5 | 6 | 11.7% | 0.47 [-0.69 , 1.62] |
| Park 2021b | 4.16 | 1.27 | 3 | 2.45 | 1.37 | 3 | 8.0% | 1.04 [-0.85 , 2.92] |
| Sameni 2019 | 22.8 | 5.12 | 10 | 8.67 | 5.46 | 10 | 11.2% | 2.56 [1.31 , 3.80] |
| **Total (Wald[a])** | | | **65** | | | **61** | **100.0%** | **1.46 [0.70 , 2.22]** |

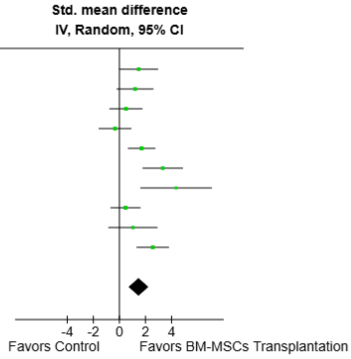


Test for overall effect: Z = 3.78 (P = 0.0002)

Heterogeneity: Tau² (DL[b]) = 0.93; Chi² = 25.89, df = 9 (P = 0.002); I² = 65%

**Footnotes**
[a]CI calculated by Wald-type method.
[b]Tau² calculated by DerSimonian and Laird method.

**Supplemental Fig. 22. Forest plot of the pooled effect of BM-MSC transplantation on antral follicle count.** Forest plot showing the standardized mean difference (SMD) in antral follicle count between animals treated with bone marrow–derived mesenchymal stem cell (BM-MSC) transplantation and control groups. Effect estimates were pooled using a random-effects inverse-variance model and are presented with 95% confidence intervals (CIs). Individual study estimates are shown as squares, with the size of each square reflecting the study weight, and the pooled effect estimate is shown as a diamond. BM-MSC transplantation was associated with a significantly higher antral follicle count compared with controls (SMD = 1.46, 95% CI 0.70 to 2.22; P = 0.0002). Substantial heterogeneity was observed across studies (I² = 65%). Values to the right of the line of no effect favor BM-MSC transplantation.

| Study or Subgroup | BM-MSCs Transplantation Mean | SD | Total | Control Mean | SD | Total | Weight | Std. mean difference IV, Random, 95% CI |
|---|---|---|---|---|---|---|---|---|
| Abd-Allah 2013 | 79.8 | 2.31 | 15 | 22 | 1.66 | 16 | 8.2% | 28.14 [20.61 , 35.67] |
| Kilic 2014 | 17 | 1.41 | 11 | 12 | 1 | 7 | 17.0% | 3.74 [2.07 , 5.42] |
| Lu 2024 | 1312.5 | 169.64 | 6 | 580.36 | 223.21 | 6 | 16.6% | 3.41 [1.40 , 5.42] |
| Mohamed 2018 | 4.22 | 1.47 | 6 | 2.22 | 0.98 | 6 | 17.4% | 1.48 [0.14 , 2.82] |
| Nabil Salama 2024 | 142.33 | 11.55 | 6 | 16.28 | 6.3 | 6 | 9.9% | 12.51 [6.30 , 18.72] |
| Park 2021b | 16.44 | 3.24 | 3 | 10.11 | 4.23 | 3 | 16.6% | 1.34 [-0.72 , 3.40] |
| Park 2023 | 24.46 | 7.47 | 3 | 2.56 | 1.92 | 3 | 14.3% | 3.21 [-0.28 , 6.70] |
| **Total (Wald[a])** | | | **50** | | | **47** | **100.0%** | **5.69 [2.77 , 8.60]** |

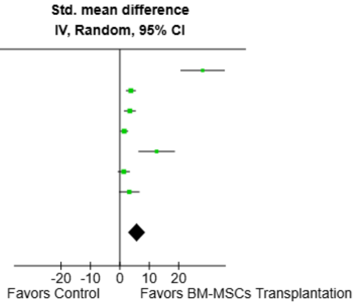


Test for overall effect: Z = 3.83 (P = 0.0001)

Heterogeneity: Tau² (DL[b]) = 12.24; Chi² = 60.11, df = 6 (P < 0.00001); I² = 90%

**Footnotes**
[a]CI calculated by Wald-type method.
[b]Tau² calculated by DerSimonian and Laird method.

**Supplemental Fig. 23. Forest plot of the pooled effect of BM-MSC transplantation on total follicle count.** Forest plot showing the standardized mean difference (SMD) in total follicle count between animals treated with bone marrow–derived mesenchymal stem cell (BM-MSC) transplantation and control groups. Effect estimates were pooled using a random-effects inverse-variance model and are presented with 95% confidence intervals (CIs). Individual study estimates are shown as squares, with the size of each square reflecting the study weight, and the pooled effect estimate is shown as a diamond. BM-MSC transplantation was associated with a significantly higher total follicle count compared with controls (SMD = 5.69, 95% CI 2.77 to 8.60; P = 0.0001). Considerable heterogeneity was observed across studies (I² = 90%). Values to the right of the line of no effect favor BM-MSC transplantation.

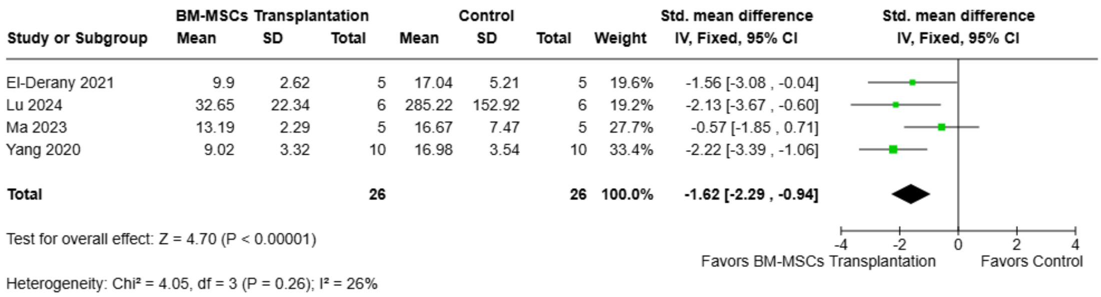


**Supplemental Fig. 24. Forest plot of the pooled effect of BM-MSC transplantation on atretic follicle count.** Forest plot showing the standardized mean difference (SMD) in atretic follicle count between animals treated with bone marrow-derived mesenchymal stem cell (BM-MSC) transplantation and control groups. Effect estimates were pooled using a fixed-effect inverse-variance model and are presented with 95% confidence intervals (CIs). Individual study estimates are shown as squares, with the size of each square reflecting the study weight, and the pooled effect estimate is shown as a diamond. BM-MSC transplantation was associated with a significantly lower atretic follicle count compared with controls (SMD = −1.62, 95% CI −2.29 to −0.94; $P < 0.00001$). Low heterogeneity was observed across studies ($I^2$ = 26%). Values to the left of the line of no effect favor BM-MSC transplantation.

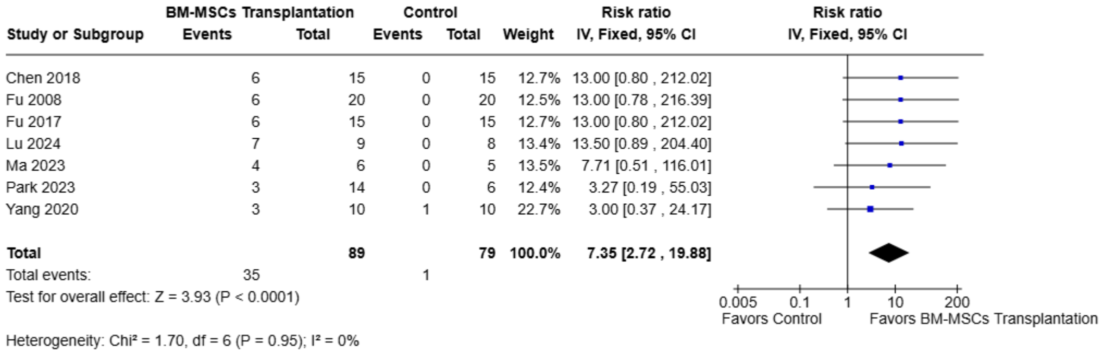


**Supplemental Fig. 25. Forest plot of the pooled effect of BM-MSC transplantation on normal estrous cycle.** Forest plot showing the risk ratio (RR) for the presence of a normal estrous cycle between animals treated with bone marrow–derived mesenchymal stem cell (BM-MSC) transplantation and control groups. Effect estimates were pooled using a fixed-effect inverse-variance model and are presented with 95% confidence intervals (CIs). Individual study estimates are shown as squares, with the size of each square reflecting the study weight, and the pooled effect estimate is shown as a diamond. BM-MSC transplantation was associated with a significantly higher likelihood of normal estrous cycle recovery compared with controls (RR = 7.35, 95% CI 2.72 to 19.88; $P < 0.0001$). No statistical heterogeneity was observed across studies ($I^2$ = 0%). Values to the right of the line of no effect favor BM-MSC transplantation.

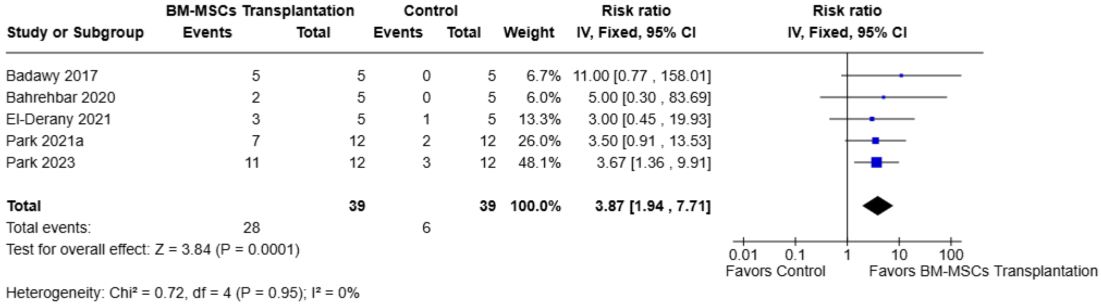


**Supplemental Fig. 26. Forest plot of the pooled effect of BM-MSC transplantation on pregnancy occurrence.** Forest plot showing the risk ratio (RR) for pregnancy occurrence between animals treated with bone marrow–derived mesenchymal stem cell (BM-MSC) transplantation and control groups. Effect estimates were pooled using a fixed-effect inverse-variance model and are presented with 95% confidence intervals (CIs). Individual study estimates are shown as squares, with the size of each square reflecting the study weight, and the pooled effect estimate is shown as a diamond. BM-MSC transplantation was associated with a significantly higher likelihood of pregnancy occurrence compared with controls (RR = 3.87, 95% CI 1.94 to 7.71; P = 0.0001). No statistical heterogeneity was observed across studies ($I^2$ = 0%). Values to the right of the line of no effect favor BM-MSC transplantation.

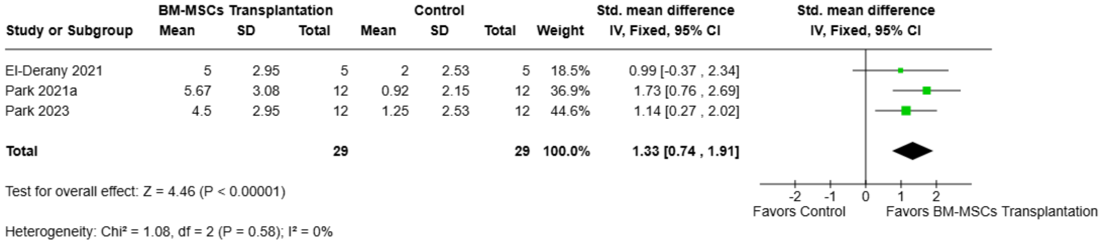


**Supplemental Fig. 27. Forest plot of the pooled effect of BM-MSC transplantation on the number of offspring.** Forest plot showing the standardized mean difference (SMD) in the number of offspring between animals treated with bone marrow–derived mesenchymal stem cell (BM-MSC) transplantation and control groups. Effect estimates were pooled using a fixed-effect inverse-variance model and are presented with 95% confidence intervals (CIs). Individual study estimates are shown as squares, with the size of each square reflecting the study weight, and the pooled effect estimate is shown as a diamond. BM-MSC transplantation was associated with a significantly higher number of offspring compared with controls (SMD = 1.33, 95% CI 0.74 to 1.91; P < 0.00001). No statistical heterogeneity was observed across studies ($I^2$ = 0%). Values to the right of the line of no effect favor BM-MSC transplantation.

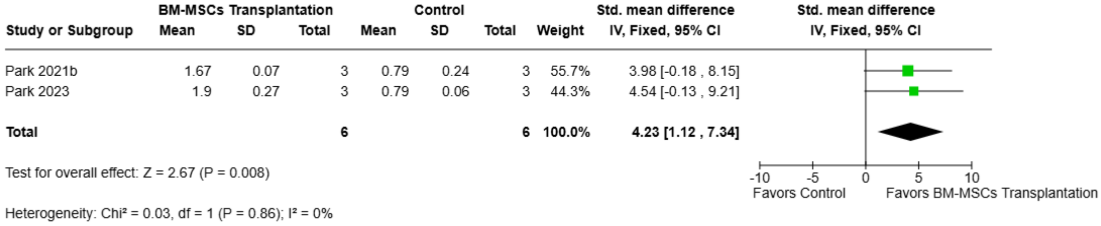


**Supplemental Fig. 28. Forest plot of the pooled effect of BM-MSC transplantation on ovarian size.** Forest plot showing the standardized mean difference (SMD) in ovarian size between animals treated with bone marrow–derived mesenchymal stem cell (BM-MSC)

transplantation and control groups. Effect estimates were pooled using a fixed-effect inverse-variance model and are presented with 95% confidence intervals (CIs). Individual study estimates are shown as squares, with the size of each square reflecting the study weight, and the pooled effect estimate is shown as a diamond. BM-MSC transplantation was associated with a significantly larger ovarian size compared with controls (SMD = 4.23, 95% CI 1.12 to 7.34; P = 0.008). No statistical heterogeneity was observed across studies (I² = 0%). Values to the right of the line of no effect favor BM-MSC transplantation.

| Study or Subgroup | BM-MSCs Transplantation Mean | SD | Total | Control Mean | SD | Total | Weight | Std. mean difference IV, Fixed, 95% CI |
|---|---|---|---|---|---|---|---|---|
| Chen 2018 | 21.81 | 1.17 | 5 | 18.19 | 0.1 | 5 | 6.0% | 3.94 [1.40 , 6.48] |
| El-Derany 2021 | 0.07 | 0.02 | 8 | 0.04 | 0.01 | 8 | 26.5% | 1.79 [0.58 , 3.01] |
| Fu 2017 | 21.85 | 1.81 | 5 | 17.72 | 1.29 | 5 | 11.7% | 2.37 [0.55 , 4.20] |
| Lu 2024 | 75.72 | 13.02 | 9 | 62.7 | 6.75 | 8 | 35.2% | 1.17 [0.12 , 2.22] |
| Mohamed 2018 | 25.89 | 2.33 | 6 | 19.93 | 2.41 | 6 | 15.2% | 2.32 [0.72 , 3.92] |
| Nabil Salama 2024 | 64.51 | 10.96 | 6 | 21.34 | 2.3 | 6 | 5.3% | 5.03 [2.33 , 7.74] |
| **Total** | | | **39** | | | **38** | **100.0%** | **2.02 [1.40 , 2.65]** |

Test for overall effect: Z = 6.35 (P < 0.00001)

Heterogeneity: Chi² = 9.88, df = 5 (P = 0.08); I² = 49%

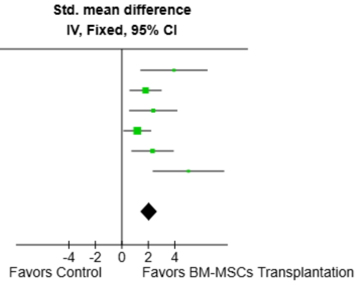


**Supplemental Fig. 29. Forest plot of the pooled effect of BM-MSC transplantation on ovarian weight.** Forest plot showing the standardized mean difference (SMD) in ovarian weight between animals treated with bone marrow–derived mesenchymal stem cell (BM-MSC) transplantation and control groups. Effect estimates were pooled using a fixed-effect inverse-variance model and are presented with 95% confidence intervals (CIs). Individual study estimates are shown as squares, with the size of each square reflecting the study weight, and the pooled effect estimate is shown as a diamond. BM-MSC transplantation was associated with a significantly higher ovarian weight compared with controls (SMD = 2.02, 95% CI 1.40 to 2.65; P < 0.00001). Moderate heterogeneity was observed across studies (I² = 49%). Values to the right of the line of no effect favor BM-MSC transplantation.

| Study or Subgroup | BM-MSCs Transplantation Mean | SD | Total | Control Mean | SD | Total | Weight | Std. mean difference IV, Random, 95% CI |
|---|---|---|---|---|---|---|---|---|
| Atwa 2025 | 164.7 | 10.05 | 7 | 216.3 | 11.91 | 7 | 20.4% | -4.38 [-6.57 , -2.20] |
| Bahrehbar 2020 | 17.56 | 2.37 | 5 | 16.24 | 1.79 | 5 | 23.7% | 0.57 [-0.71 , 1.85] |
| El-Derany 2021 | 92.13 | 12.29 | 8 | 72.8 | 13.11 | 8 | 24.1% | 1.44 [0.30 , 2.57] |
| Maqbool 2022 | 313.33 | 6.46 | 5 | 253.33 | 2.89 | 5 | 8.1% | 10.83 [4.61 , 17.05] |
| Mohamed 2018 | 23.46 | 0.62 | 6 | 22.72 | 0.63 | 6 | 23.7% | 1.09 [-0.16 , 2.34] |
| **Total (Wald[a])** | | | **31** | | | **31** | **100.0%** | **0.73 [-1.42 , 2.87]** |

Test for overall effect: Z = 0.67 (P = 0.51)

Heterogeneity: Tau² (DL[b]) = 4.64; Chi² = 33.05, df = 4 (P < 0.00001); I² = 88%

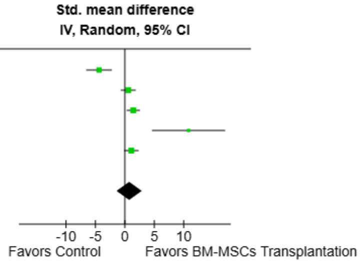


**Footnotes**
[a]CI calculated by Wald-type method.
[b]Tau² calculated by DerSimonian and Laird method.

**Supplemental Fig. 30. Forest plot of the pooled effect of BM-MSC transplantation on body weight.** Forest plot showing the standardized mean difference (SMD) in body weight between animals treated with bone marrow–derived mesenchymal stem cell (BM-MSC) transplantation and control groups. Effect estimates were pooled using a random-effects inverse-variance model and are presented with 95% confidence intervals (CIs). Individual study estimates are shown as squares, with the size of each square reflecting the study weight, and the pooled effect estimate is shown as a diamond. BM-MSC transplantation was not associated with a significant difference in body weight compared with controls (SMD = 0.73, 95% CI −1.42 to 2.87; P = 0.51). Considerable heterogeneity was observed across studies ($I^2$ = 88%). Values to the right of the line of no effect favor BM-MSC transplantation.

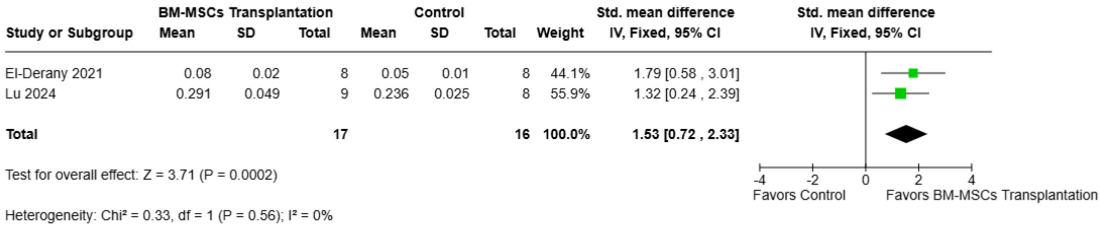


**Supplemental Fig. 31. Forest plot of the pooled effect of BM-MSC transplantation on ovarian weight index.** Forest plot showing the standardized mean difference (SMD) in ovarian weight index between animals treated with bone marrow–derived mesenchymal stem cell (BM-MSC) transplantation and control groups. Effect estimates were pooled using a fixed-effect inverse-variance model and are presented with 95% confidence intervals (CIs). Individual study estimates are shown as squares, with the size of each square reflecting the study weight, and the pooled effect estimate is shown as a diamond. BM-MSC transplantation was associated with a significantly higher ovarian weight index compared with controls (SMD = 1.53, 95% CI 0.72 to 2.33; P = 0.0002). No statistical heterogeneity was observed across studies ($I^2$ = 0%). Values to the right of the line of no effect favor BM-MSC transplantation.

#### BM-MSCs Secretome (Acellular therapy)

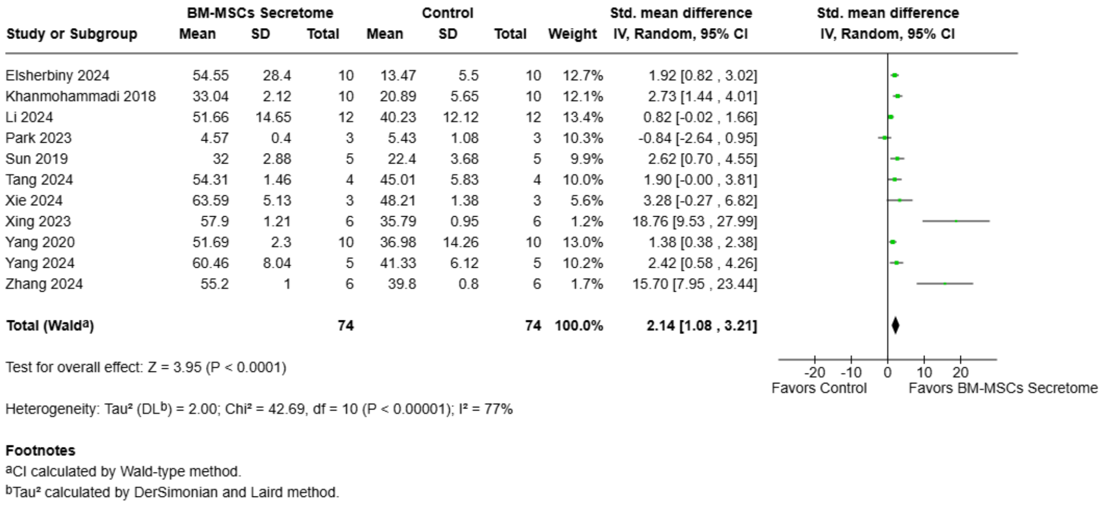


**Supplemental Fig. 32. Forest plot of the pooled effect of BM-MSC secretome on serum estradiol levels.** Forest plot showing the standardized mean difference (SMD) in serum estradiol (E2) levels between animals treated with bone marrow–derived mesenchymal stem cell (BM-MSC) secretome and control groups. Effect estimates were pooled using a random-effects inverse-variance model and are presented with 95% confidence intervals (CIs). Individual study estimates are shown as squares, with the size of each square reflecting the study weight, and the pooled effect estimate is shown as a diamond. BM-MSC secretome was associated with significantly higher serum E2 levels compared with controls (SMD = 2.14, 95% CI 1.08 to 3.21; P < 0.0001). Substantial heterogeneity was observed across studies ($I^2$ = 77%). Values to the right of the line of no effect favor BM-MSC secretome.

| Study or Subgroup | BM-MSCs Secretome Mean | SD | Total | Control Mean | SD | Total | Weight | Std. mean difference IV, Random, 95% CI |
|---|---|---|---|---|---|---|---|---|
| Elsherbiny 2024 | 46.41 | 7.68 | 10 | 98.36 | 126.71 | 10 | 12.9% | -0.55 [-1.45 , 0.34] |
| Khanmohammadi 2018 | 7.65 | 0.45 | 10 | 12.58 | 0.89 | 10 | 10.3% | -6.70 [-9.17 , -4.22] |
| Li 2024 | 15.29 | 2.73 | 12 | 18.26 | 3.91 | 12 | 13.0% | -0.85 [-1.69 , -0.01] |
| Park 2023 | 31.05 | 3.74 | 3 | 26.98 | 9.04 | 3 | 11.8% | 0.47 [-1.19 , 2.13] |
| Tang 2024 | 12.84 | 1.13 | 4 | 15.18 | 0.68 | 4 | 11.1% | -2.18 [-4.22 , -0.14] |
| Xie 2024 | 66.22 | 5.74 | 3 | 89.2 | 4.26 | 5 | 8.9% | -4.16 [-7.36 , -0.96] |
| Xing 2023 | 14.38 | 0.24 | 6 | 19.44 | 0.33 | 6 | 3.2% | -16.19 [-24.17 , -8.21] |
| Yang 2020 | 14.93 | 1.99 | 10 | 12.58 | 0.89 | 10 | 12.8% | 1.46 [0.45 , 2.47] |
| Yang 2024 | 63.92 | 6.25 | 5 | 89.2 | 4.26 | 5 | 9.8% | -4.27 [-6.97 , -1.56] |
| Zhang 2024 | 9.74 | 0.77 | 6 | 17.21 | 0.67 | 6 | 6.2% | -9.55 [-14.35 , -4.75] |
| **Total (Wald[a])** | | | **69** | | | **71** | **100.0%** | **-2.78 [-4.42 , -1.14]** |

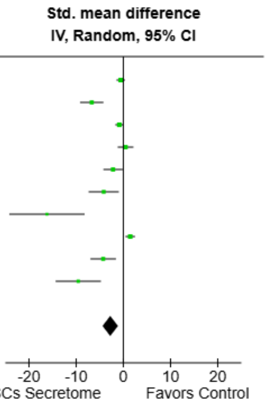


Test for overall effect: Z = 3.32 (P = 0.0009)

Heterogeneity: Tau$^2$ (DL[b]) = 5.23; Chi$^2$ = 82.84, df = 9 (P < 0.00001); $I^2$ = 89%

**Footnotes**

[a]CI calculated by Wald-type method.

[b]Tau$^2$ calculated by DerSimonian and Laird method.

**Supplemental Fig. 33. Forest plot of the pooled effect of BM-MSC secretome on serum follicle-stimulating hormone levels.** Forest plot showing the standardized mean difference (SMD) in serum follicle-stimulating hormone (FSH) levels between animals treated with bone marrow-derived mesenchymal stem cell (BM-MSC) secretome and control groups. Effect estimates were pooled using a random-effects inverse-variance model and are presented with 95% confidence intervals (CI). Individual study estimates are shown as squares, with the size of each square reflecting the study weight, and the pooled effect estimate is shown as a diamond. BM-MSC secretome was associated with significantly lower serum FSH levels compared with controls (SMD = -2.78, 95% CI -4.42 to -1.14; P = 0.0009). Considerable heterogeneity was observed across studies ($I^2$ = 89%). Values to the left of the line of no effect favor BM-MSC secretome.

| Study or Subgroup | BM-MSCs Secretome Mean | SD | Total | Control Mean | SD | Total | Weight | Std. mean difference IV, Random, 95% CI |
|---|---|---|---|---|---|---|---|---|
| Elsherbiny 2024 | 8.05 | 3.23 | 10 | 2.71 | 2.02 | 10 | 15.5% | 1.90 [0.80 , 2.99] |
| Li 2024 | 143.99 | 21.75 | 12 | 130.09 | 12.82 | 12 | 16.4% | 0.75 [-0.08 , 1.59] |
| Park 2023 | 3.24 | 0.01 | 3 | 4.01 | 1.53 | 3 | 13.1% | -0.57 [-2.26 , 1.12] |
| Tang 2024 | 2224 | 176 | 4 | 1578 | 214 | 4 | 10.3% | 2.87 [0.46 , 5.28] |
| Xie 2024 | 538.25 | 25.81 | 3 | 440.55 | 40.56 | 3 | 9.2% | 2.30 [-0.44 , 5.04] |
| Xing 2023 | 146.2 | 0.7 | 6 | 129.06 | 0.88 | 6 | 1.4% | 19.90 [10.12 , 29.68] |
| Yang 2020 | 150.25 | 9.49 | 10 | 131 | 11.86 | 10 | 15.6% | 1.72 [0.66 , 2.78] |
| Yang 2024 | 452.38 | 42.33 | 5 | 380.95 | 76.72 | 5 | 14.4% | 1.04 [-0.33 , 2.41] |
| Zhang 2024 | 151.03 | 2.07 | 6 | 131.38 | 1.38 | 6 | 4.1% | 10.31 [5.15 , 15.47] |
| **Total (Wald[a])** | | | **59** | | | **59** | **100.0%** | **1.97 [0.77 , 3.16]** |

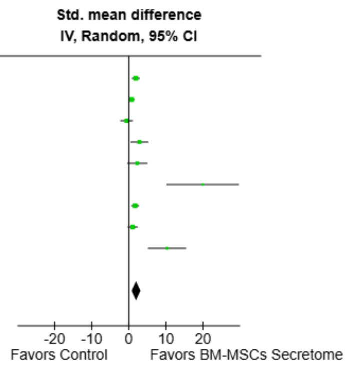


Test for overall effect: Z = 3.23 (P = 0.001)

Heterogeneity: Tau² (DL[b]) = 2.08; Chi² = 35.93, df = 8 (P < 0.0001); I² = 78%

**Footnotes**
[a]CI calculated by Wald-type method.
[b]Tau² calculated by DerSimonian and Laird method.

**Supplemental Fig. 34. Forest plot of the pooled effect of BM-MSC secretome on serum anti-Müllerian hormone levels.** Forest plot showing the standardized mean difference (SMD) in serum anti-Müllerian hormone (AMH) levels between animals treated with bone marrow–derived mesenchymal stem cell (BM-MSC) secretome and control groups. Effect estimates were pooled using a random-effects inverse-variance model and are presented with 95% confidence intervals (CI). Individual study estimates are shown as squares, with the size of each square reflecting the study weight, and the pooled effect estimate is shown as a diamond. BM-MSC secretome was associated with significantly higher serum AMH levels compared with controls (SMD = 1.97, 95% CI 0.77 to 3.16; P = 0.001). Substantial heterogeneity was observed across studies (I² = 78%). Values to the right of the line of no effect favor BM-MSC secretome.

| Study or Subgroup | BM-MSCs Secretome Mean | SD | Total | Control Mean | SD | Total | Weight | Mean difference IV, Random, 95% CI |
|---|---|---|---|---|---|---|---|---|
| Elsherbiny 2024 | 8.12 | 6.86 | 10 | 28.18 | 26.56 | 10 | 0.1% | -20.06 [-37.06 , -3.06] |
| Li 2024 | 6.7 | 2.32 | 12 | 8.67 | 3.75 | 12 | 4.2% | -1.97 [-4.46 , 0.52] |
| Xing 2023 | 6.8 | 0.06 | 6 | 9.15 | 0.06 | 6 | 40.9% | -2.35 [-2.42 , -2.28] |
| Yang 2020 | 6.1 | 1.08 | 10 | 10.92 | 1.42 | 10 | 15.0% | -4.82 [-5.93 , -3.71] |
| Zhang 2024 | 5.96 | 0.13 | 6 | 8.81 | 0.13 | 6 | 39.9% | -2.85 [-3.00 , -2.70] |
| **Total (Wald[a])** | | | **44** | | | **44** | **100.0%** | **-2.92 [-3.46 , -2.38]** |

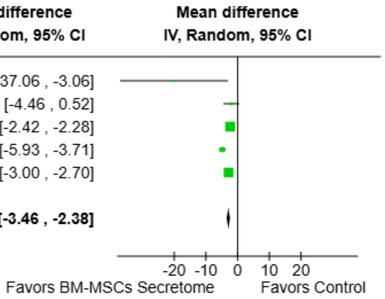


Test for overall effect: Z = 10.66 (P < 0.00001)

Heterogeneity: Tau² (DL[b]) = 0.18; Chi² = 58.62, df = 4 (P < 0.00001); I² = 93%

**Footnotes**
[a]CI calculated by Wald-type method.
[b]Tau² calculated by DerSimonian and Laird method.

**Supplemental Fig. 35. Forest plot of the pooled effect of BM-MSC secretome on serum luteinizing hormone levels.** Forest plot showing the mean difference (MD) in serum luteinizing hormone (LH) levels between animals treated with bone marrow–derived mesenchymal stem cell (BM-MSC) secretome and control groups. Effect estimates were pooled using a random-effects inverse-variance model and are presented with 95% confidence intervals (CI). Individual study estimates are shown as squares, with the size of each square reflecting the study weight, and the pooled effect estimate is shown as a diamond. BM-MSC secretome was associated with significantly lower serum LH levels compared with controls (MD = −2.92, 95% CI −3.46 to −2.38; P < 0.00001). Considerable heterogeneity was observed across studies (I² = 93%). Values to the left of the line of no effect favor BM−MSC secretome.

| Study or Subgroup | BM-MSCs Secretome Mean | SD | Total | Control Mean | SD | Total | Weight | Std. mean difference IV, Fixed, 95% CI |
|---|---|---|---|---|---|---|---|---|
| Khanmohammadi 2018 | 35.33 | 10.17 | 10 | 21.5 | 4.33 | 10 | 34.5% | 1.69 [0.64 , 2.75] |
| Tang 2024 | 176.4 | 12.4 | 4 | 104.3 | 12.4 | 4 | 2.7% | 5.06 [1.31 , 8.80] |
| Xie 2024 | 4.74 | 0.56 | 5 | 3.28 | 0.3 | 5 | 9.0% | 2.94 [0.87 , 5.00] |
| Xing 2023 | 14.23 | 1.29 | 6 | 5.98 | 1.55 | 6 | 4.7% | 5.34 [2.50 , 8.18] |
| Yang 2020 | 16.04 | 2.06 | 10 | 6.03 | 3.16 | 10 | 16.6% | 3.59 [2.07 , 5.12] |
| Yang 2024 | 4.84 | 1.14 | 5 | 3.05 | 1.03 | 5 | 17.1% | 1.49 [-0.01 , 2.98] |
| Zhang 2024 | 5.97 | 0.89 | 6 | 2.98 | 1.48 | 6 | 15.3% | 2.26 [0.68 , 3.84] |
| **Total** | | | **46** | | | **46** | **100.0%** | **2.44 [1.82 , 3.06]** |

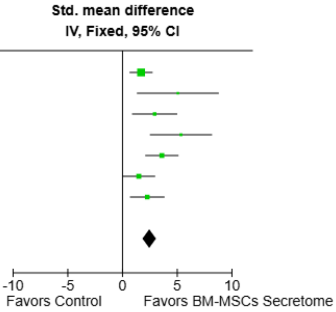


Test for overall effect: Z = 7.72 (P < 0.00001)

Heterogeneity: Chi² = 11.83, df = 6 (P = 0.07); I² = 49%

**Supplemental Fig. 36. Forest plot of the pooled effect of BM-MSC secretome on primordial follicle count.** Forest plot showing the standardized mean difference (SMD) in primordial follicle count between animals treated with bone marrow–derived mesenchymal stem cell (BM-MSC) secretome and control groups. Effect estimates were pooled using a fixed-effect inverse-variance model and are presented with 95% confidence intervals (CI). Individual study estimates are shown as squares, with the size of each square reflecting the study weight, and the pooled effect estimate is shown as a diamond. BM-MSC secretome was associated with a significantly higher primordial follicle count compared with controls (SMD = 2.44, 95% CI 1.82 to 3.06; P < 0.00001). Moderate heterogeneity was observed across studies ($I^2$ = 49%). Values to the right of the line of no effect favor BM-MSC secretome.

| Study or Subgroup | BM-MSCs Secretome Mean | SD | Total | Control Mean | SD | Total | Weight | Std. mean difference IV, Random, 95% CI |
|---|---|---|---|---|---|---|---|---|
| Khanmohammadi 2018 | 30 | 6.67 | 10 | 22.5 | 9.67 | 10 | 22.8% | 0.86 [-0.06 , 1.79] |
| Tang 2024 | 112.4 | 13.5 | 4 | 65.2 | 9.45 | 4 | 7.5% | 3.52 [0.73 , 6.31] |
| Xie 2024 | 4.02 | 0.8 | 5 | 0.98 | 0.48 | 5 | 8.1% | 4.16 [1.51 , 6.81] |
| Xing 2023 | 11.04 | 1.26 | 6 | 5.29 | 1.73 | 6 | 11.5% | 3.51 [1.45 , 5.56] |
| Yang 2020 | 11.01 | 3.32 | 10 | 5.35 | 3.73 | 10 | 21.5% | 1.54 [0.51 , 2.56] |
| Yang 2024 | 4.12 | 0.72 | 5 | 2.68 | 0.65 | 5 | 14.9% | 1.90 [0.26 , 3.53] |
| Zhang 2024 | 5.11 | 0.8 | 6 | 2.58 | 0.89 | 6 | 13.7% | 2.76 [1.00 , 4.52] |
| **Total (Wald[a])** | | | **46** | | | **46** | **100.0%** | **2.19 [1.31 , 3.08]** |

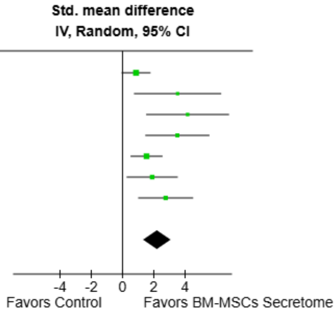


Test for overall effect: Z = 4.88 (P < 0.00001)

Heterogeneity: Tau² (DL[b]) = 0.67; Chi² = 12.50, df = 6 (P = 0.05); I² = 52%

**Footnotes**
[a]CI calculated by Wald-type method.
[b]Tau² calculated by DerSimonian and Laird method.

**Supplemental Fig. 37. Forest plot of the pooled effect of BM-MSC secretome on primary follicle count.** Forest plot showing the standardized mean difference (SMD) in primary follicle count between animals treated with bone marrow–derived mesenchymal stem cell (BM-MSC) secretome and control groups. Effect estimates were pooled using a random-effects inverse-variance model and are presented with 95% confidence intervals (CI). Individual study estimates are shown as squares, with the size of each square reflecting the study weight, and the pooled effect estimate is shown as a diamond. BM-MSC secretome was associated with a significantly higher primary follicle count compared with controls (SMD = 2.19, 95% CI 1.31 to 3.08; P < 0.00001). Significant heterogeneity was observed across studies ($I^2$ = 52%). Values to the right of the line of no effect favor BM-MSC secretome.

| Study or Subgroup | BM-MSCs Secretome Mean | SD | Total | Control Mean | SD | Total | Weight | Std. mean difference IV, Random, 95% CI |
|---|---|---|---|---|---|---|---|---|
| Khanmohammadi 2018 | 28.67 | 6 | 10 | 21.67 | 6.17 | 10 | 21.4% | 1.10 [0.15 , 2.06] |
| Tang 2024 | 129.83 | 5.68 | 4 | 43.56 | 6.82 | 4 | 1.3% | 11.95 [3.62 , 20.29] |
| Xie 2024 | 2.23 | 0.29 | 5 | 0.7 | 0.45 | 5 | 10.3% | 3.65 [1.25 , 6.05] |
| Xing 2023 | 8.31 | 1.77 | 6 | 2.97 | 1.33 | 6 | 13.3% | 3.15 [1.24 , 5.06] |
| Yang 2020 | 9.02 | 3.32 | 10 | 3.98 | 3.35 | 10 | 20.9% | 1.45 [0.44 , 2.46] |
| Yang 2024 | 3.75 | 1.6 | 5 | 2.68 | 0.82 | 5 | 18.2% | 0.76 [-0.55 , 2.07] |
| Zhang 2024 | 4.09 | 0.8 | 6 | 1.72 | 0.83 | 6 | 14.6% | 2.68 [0.95 , 4.42] |
| **Total (Wald[a])** | | | **46** | | | **46** | **100.0%** | **2.02 [1.04 , 3.00]** |

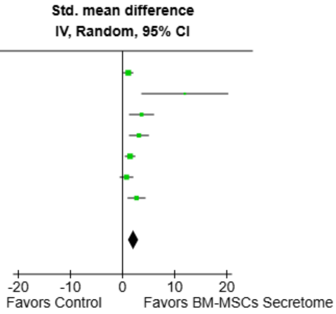


Test for overall effect: Z = 4.05 (P < 0.0001)

Heterogeneity: Tau² (DL[b]) = 0.92; Chi² = 15.46, df = 6 (P = 0.02); I² = 61%

**Footnotes**
[a]CI calculated by Wald-type method.
[b]Tau² calculated by DerSimonian and Laird method.

**Supplemental Fig. 38. Forest plot of the pooled effect of BM-MSC secretome on secondary follicle count.** Forest plot showing the standardized mean difference (SMD) in secondary follicle count between animals treated with bone marrow–derived mesenchymal stem cell (BM-MSC) secretome and control groups. Effect estimates were pooled using a random-effects inverse-variance model and are presented with 95% confidence intervals (CI). Individual study estimates are shown as squares, with the size of each square reflecting the study weight, and the pooled effect estimate is shown as a diamond. BM-MSC secretome was associated with a significantly higher secondary follicle count compared with controls (SMD = 2.02, 95% CI 1.04 to 3.00; P < 0.0001). Significant heterogeneity was observed across studies (I² = 61%). Values to the right of the line of no effect favor BM-MSC secretome.

| Study or Subgroup | BM-MSCs Secretome Mean | SD | Total | Control Mean | SD | Total | Weight | Std. mean difference IV, Random, 95% CI |
|---|---|---|---|---|---|---|---|---|
| Elsherbiny 2024 | 9.23 | 2.12 | 10 | 1.25 | 2.02 | 10 | 29.1% | 3.69 [2.14 , 5.24] |
| Nabil Salama 2024 | 144.96 | 7.88 | 6 | 16.28 | 6.3 | 6 | 6.4% | 16.65 [8.44 , 24.86] |
| Park 2023 | 15.47 | 3.29 | 3 | 5.43 | 1.56 | 3 | 19.4% | 3.12 [-0.29 , 6.53] |
| Tang 2024 | 428.57 | 40.1 | 4 | 218.05 | 25.06 | 4 | 16.7% | 5.47 [1.46 , 9.49] |
| Yang 2024 | 13.64 | 2.27 | 5 | 8.9 | 1.83 | 5 | 28.4% | 2.08 [0.37 , 3.78] |
| **Total (Wald[a])** | | | **28** | | | **28** | **100.0%** | **4.25 [1.94 , 6.57]** |

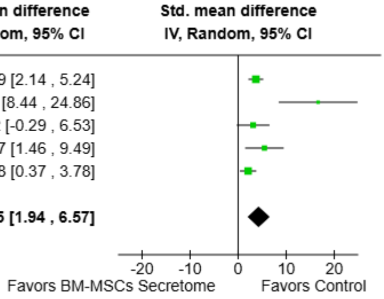


Test for overall effect: Z = 3.60 (P = 0.0003)

Heterogeneity: Tau² (DL[b]) = 4.16; Chi² = 13.52, df = 4 (P = 0.009); I² = 70%

**Footnotes**
[a]CI calculated by Wald-type method.
[b]Tau² calculated by DerSimonian and Laird method.

**Supplemental Fig. 39. Forest plot of the pooled effect of BM-MSC secretome on total follicle count.** Forest plot showing the standardized mean difference (SMD) in total follicle count between animals treated with bone marrow–derived mesenchymal stem cell (BM-MSC) secretome and control groups. Effect estimates were pooled using a random-effects inverse-variance model and are presented with 95% confidence intervals (CI). Individual study estimates are shown as squares, with the size of each square reflecting the study weight, and the pooled effect estimate is shown as a diamond. BM-MSC secretome was associated with a significantly higher total follicle count compared with controls (SMD = 4.25, 95% CI 1.94 to 6.57; P = 0.0003). Significant heterogeneity was observed across studies (I² = 70%). Values to the right of the line of no effect favor BM-MSC secretome.

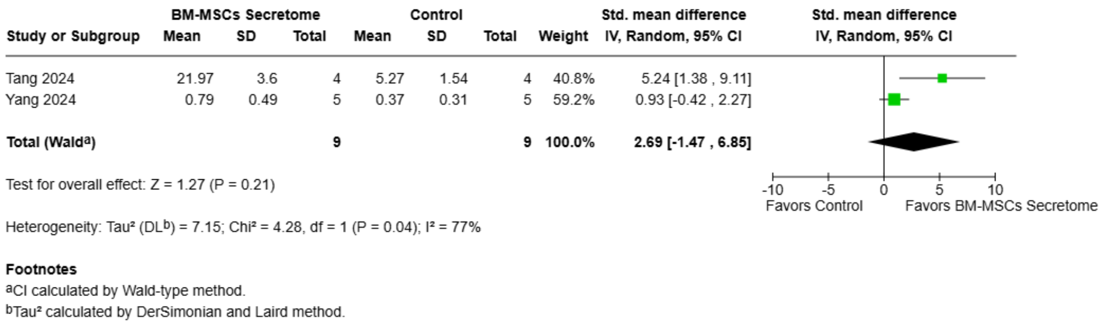

| Study or Subgroup | BM-MSCs Secretome Mean | SD | Total | Control Mean | SD | Total | Weight | Std. mean difference IV, Random, 95% CI |
|---|---|---|---|---|---|---|---|---|
| Tang 2024 | 21.97 | 3.6 | 4 | 5.27 | 1.54 | 4 | 40.8% | 5.24 [1.38 , 9.11] |
| Yang 2024 | 0.79 | 0.49 | 5 | 0.37 | 0.31 | 5 | 59.2% | 0.93 [-0.42 , 2.27] |
| **Total (Wald[a])** | | | **9** | | | **9** | **100.0%** | **2.69 [-1.47 , 6.85]** |

Test for overall effect: Z = 1.27 (P = 0.21)

Heterogeneity: Tau² (DL[b]) = 7.15; Chi² = 4.28, df = 1 (P = 0.04); I² = 77%

**Footnotes**

[a]CI calculated by Wald-type method.

[b]Tau² calculated by DerSimonian and Laird method.



**Supplemental Fig. 40. Forest plot of the pooled effect of BM-MSC secretome on mature follicle count.** Forest plot showing the standardized mean difference (SMD) in mature follicle count between animals treated with bone marrow-derived mesenchymal stem cell (BM-MSC) secretome and control groups. Effect estimates were pooled using a random-effects inverse-variance model and are presented with 95% confidence intervals (CI). Individual study estimates are shown as squares, with the size of each square reflecting the study weight, and the pooled effect estimate is shown as a diamond. BM-MSC secretome was not associated with a significant difference in mature follicle count compared with controls (SMD = 2.69, 95% CI −1.47 to 6.85; P = 0.21). Significant heterogeneity was observed across studies ($I^2$ = 77%). Values to the right of the line of no effect favor BM-MSC secretome.

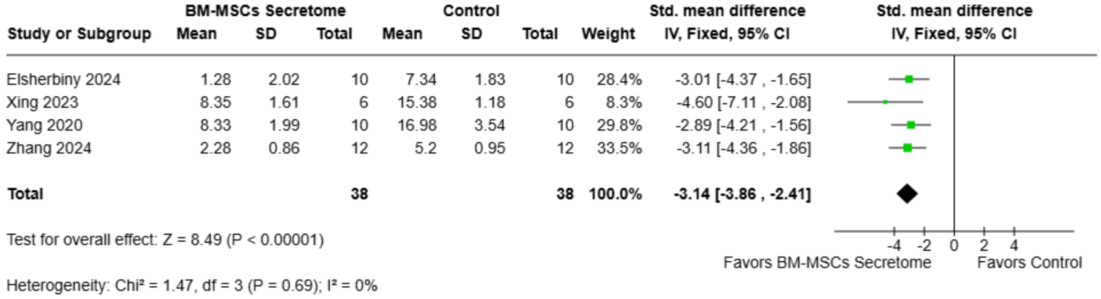

| Study or Subgroup | BM-MSCs Secretome Mean | SD | Total | Control Mean | SD | Total | Weight | Std. mean difference IV, Fixed, 95% CI |
|---|---|---|---|---|---|---|---|---|
| Elsherbiny 2024 | 1.28 | 2.02 | 10 | 7.34 | 1.83 | 10 | 28.4% | -3.01 [-4.37 , -1.65] |
| Xing 2023 | 8.35 | 1.61 | 6 | 15.38 | 1.18 | 6 | 8.3% | -4.60 [-7.11 , -2.08] |
| Yang 2020 | 8.33 | 1.99 | 10 | 16.98 | 3.54 | 10 | 29.8% | -2.89 [-4.21 , -1.56] |
| Zhang 2024 | 2.28 | 0.86 | 12 | 5.2 | 0.95 | 12 | 33.5% | -3.11 [-4.36 , -1.86] |
| **Total** | | | **38** | | | **38** | **100.0%** | **-3.14 [-3.86 , -2.41]** |

Test for overall effect: Z = 8.49 (P < 0.00001)

Heterogeneity: Chi² = 1.47, df = 3 (P = 0.69); I² = 0%



**Supplemental Fig. 41. Forest plot of the pooled effect of BM-MSC secretome on atretic follicle count.** Forest plot showing the standardized mean difference (SMD) in atretic follicle count between animals treated with bone marrow-derived mesenchymal stem cell (BM-MSC) secretome and control groups. Effect estimates were pooled using a fixed-effect inverse-variance model and are presented with 95% confidence intervals (CI). Individual study estimates are shown as squares, with the size of each square reflecting the study weight, and the pooled effect estimate is shown as a diamond. BM-MSC secretome was associated with a significantly lower atretic follicle count compared with controls (SMD = −3.14, 95% CI −3.86 to −2.41; P < 0.00001). No significant heterogeneity was observed across studies

(I² = 0%). Values to the left of the line of no effect favor BM-MSC secretome.

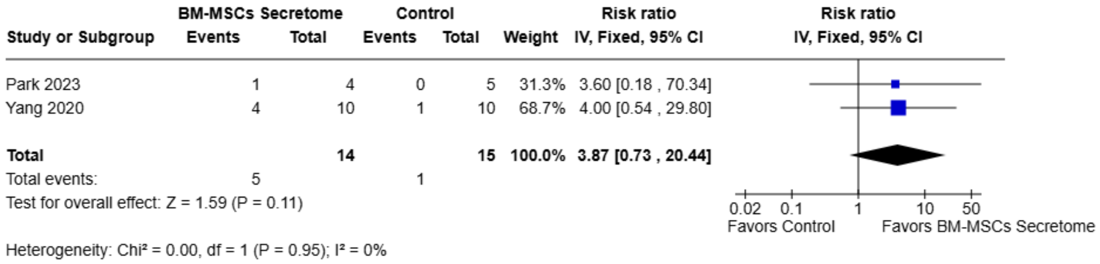


**Supplemental Fig. 42. Forest plot of the pooled effect of BM-MSC secretome on normal estrous cycle.** Forest plot showing the risk ratio (RR) for the presence of a normal estrous cycle between animals treated with bone marrow–derived mesenchymal stem cell (BM-MSC) secretome and control groups. Effect estimates were pooled using a fixed-effect inverse-variance model and are presented with 95% confidence intervals (CI). Individual study estimates are shown as squares, with the size of each square reflecting the study weight, and the pooled effect estimate is shown as a diamond. BM-MSC secretome was not associated with a significant difference in normal estrous cycle recovery compared with controls (RR = 3.87, 95% CI 0.73 to 20.44; P = 0.11). No significant heterogeneity was observed across studies (I² = 0%). Values to the right of the line of no effect favor BM-MSC secretome.

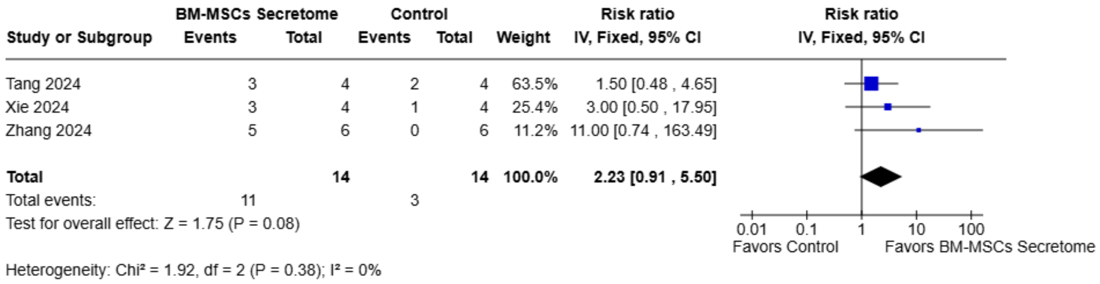


**Supplemental Fig. 43. Forest plot of the pooled effect of BM-MSC secretome on pregnancy occurrence.** Forest plot showing the risk ratio (RR) for pregnancy occurrence between animals treated with bone marrow–derived mesenchymal stem cell (BM-MSC) secretome and control groups. Effect estimates were pooled using a fixed-effect inverse-variance model and are presented with 95% confidence intervals (CI). Individual study estimates are shown as squares, with the size of each square reflecting the study weight, and the pooled effect estimate is shown as a diamond. BM-MSC secretome was not associated with a significant difference in pregnancy occurrence compared with controls (RR = 2.23, 95% CI 0.91 to 5.50; P = 0.08). No significant heterogeneity was observed across studies (I² = 0%). Values to the right of the line of no effect favor BM-MSC secretome.

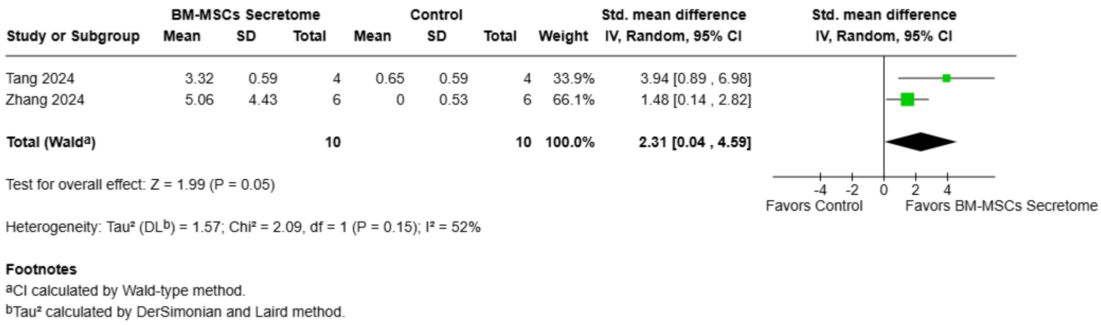


**Supplemental Fig. 44. Forest plot of the pooled effect of BM-MSC secretome on the number of offspring.** Forest plot showing the standardized mean difference (SMD) in the number of offspring between animals treated with bone marrow–derived mesenchymal stem cell (BM-MSC) secretome and control groups. Effect estimates were pooled using a random-effects inverse-variance model and are presented with 95% confidence intervals (CI). Individual study estimates are shown as squares, with the size of each square reflecting the study weight, and the pooled effect estimate is shown as a diamond. BM-MSC secretome was associated with a significantly higher number of offspring compared with controls (SMD = 2.31, 95% CI 0.04 to 4.59; P = 0.05). Significant heterogeneity was observed across studies ($I^2$ = 52%). Values to the right of the line of no effect favor BM-MSC secretome.

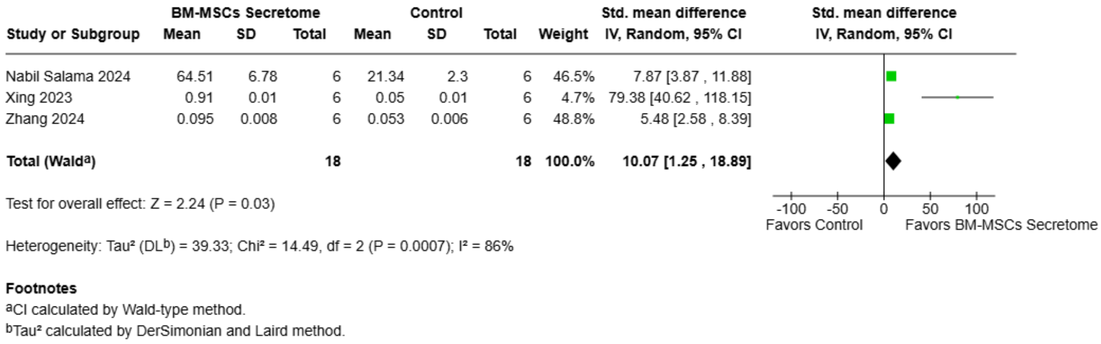


**Supplemental Fig. 45. Forest plot of the pooled effect of BM-MSC secretome on ovarian weight.** Forest plot showing the standardized mean difference (SMD) in ovarian weight between animals treated with bone marrow–derived mesenchymal stem cell (BM-MSC) secretome and control groups. Effect estimates were pooled using a random-effects inverse-variance model and are presented with 95% confidence intervals (CI). Individual study estimates are shown as squares, with the size of each square reflecting the study weight, and the pooled effect estimate is shown as a diamond. BM-MSC secretome was associated with significantly higher ovarian weight compared with controls (SMD = 10.07, 95% CI 1.25 to 18.89; P = 0.03). Significant heterogeneity was observed across studies ($I^2$ = 86%). Values to the right of the line of no effect favor BM-MSC secretome.

#### BM-MSCs miR-21 Exosomes (Acellular therapy)

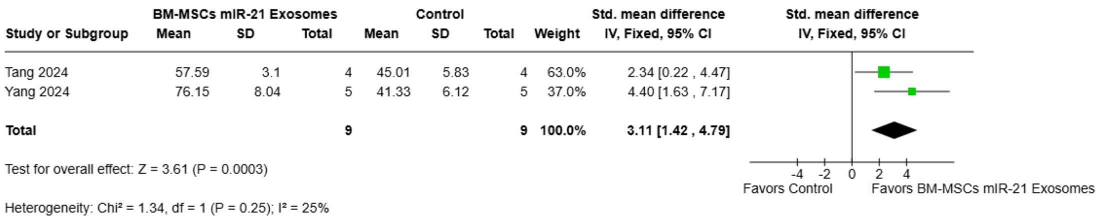


**Supplemental Fig. 46. Forest plot of the pooled effect of BM-MSC miR-21 exosomes on serum estradiol levels.** Forest plot showing the standardized mean difference (SMD) in serum estradiol (E2) levels between animals treated with bone marrow–derived mesenchymal stem cell (BM-MSC) miR-21 exosomes and control groups. Effect estimates were pooled using a fixed-effect inverse-variance model and are presented with 95% confidence intervals (CI). Individual study estimates are shown as squares, with the size of each square reflecting the study weight, and the pooled effect estimate is shown as a diamond. BM-MSC miR-21 exosomes were associated with significantly higher serum E2 levels compared with controls (SMD = 3.11, 95% CI 1.42 to 4.79; $P = 0.0003$). No significant heterogeneity was observed across studies ($I^2 = 25\%$). Values to the right of the line of no effect favor BM-MSC miR-21 exosomes.

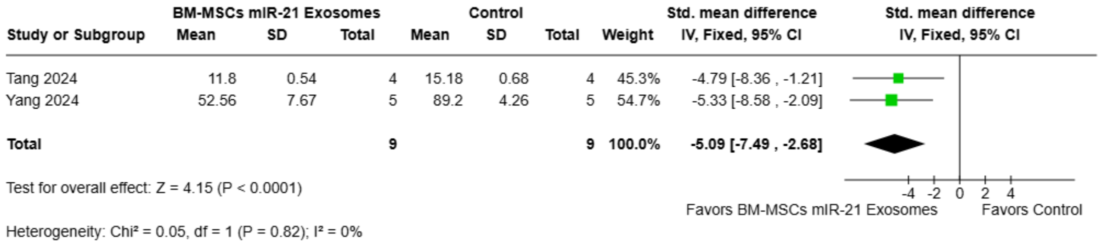


**Supplemental Fig. 47. Forest plot of the pooled effect of BM-MSC miR-21 exosomes on serum follicle-stimulating hormone levels.** Forest plot showing the standardized mean difference (SMD) in serum follicle-stimulating hormone (FSH) levels between animals treated with bone marrow-derived mesenchymal stem cell (BM-MSC) miR-21 exosomes and control groups. Effect estimates were pooled using a fixed-effect inverse-variance model and are presented with 95% confidence intervals (CI). Individual study estimates are shown as squares, with the size of each square reflecting the study weight, and the pooled effect estimate is shown as a diamond. BM-MSC miR-21 exosomes were associated with significantly lower serum FSH levels compared with controls (SMD = −5.09, 95% CI −7.49 to −2.68; $P < 0.0001$). No significant heterogeneity was observed across studies ($I^2 = 0\%$). Values to the left of the line of no effect favor BM-MSC miR-21 exosomes.

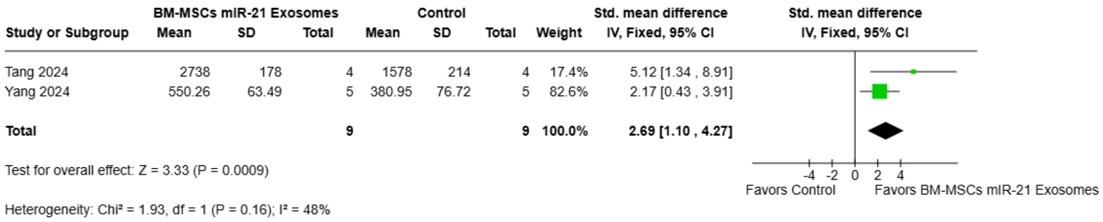


**Supplemental Fig. 48. Forest plot of the pooled effect of BM-MSC miR-21 exosomes on serum anti-Müllerian hormone levels.** Forest plot showing the standardized mean difference (SMD) in serum anti-Müllerian hormone (AMH) levels between animals treated with bone marrow–derived mesenchymal stem cell (BM-MSC) miR-21 exosomes and control groups. Effect estimates were pooled using a fixed-effect inverse-variance model and are presented with 95% confidence intervals (CI). Individual study estimates are shown as squares, with the size of each square reflecting the study weight, and the pooled effect estimate is shown as a diamond. BM-MSC miR-21 exosomes were associated with significantly higher serum AMH levels compared with controls (SMD = 2.69, 95% CI 1.10 to 4.27; P = 0.0009). No significant heterogeneity was observed across studies ($I^2$ = 48%). Values to the right of the line of no effect favor BM-MSC miR-21 exosomes.

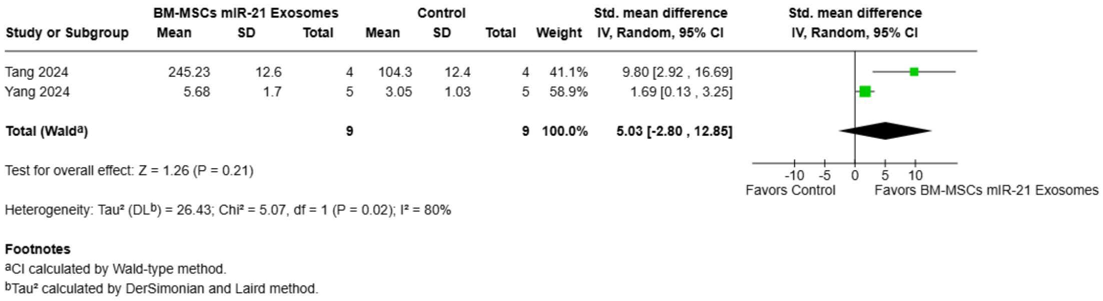


**Supplemental Fig. 49. Forest plot of the pooled effect of BM-MSC miR-21 exosomes on primordial follicle count.** Forest plot showing the standardized mean difference (SMD) in primordial follicle count between animals treated with bone marrow–derived mesenchymal stem cell (BM-MSC) miR-21 exosomes and control groups. Effect estimates were pooled using a random-effects inverse-variance model and are presented with 95% confidence intervals (CI). Individual study estimates are shown as squares, with the size of each square reflecting the study weight, and the pooled effect estimate is shown as a diamond. BM-MSC miR-21 exosomes were not associated with a significant difference in primordial follicle count compared with controls (SMD = 5.03, 95% CI −2.80 to 12.85; P = 0.21). Significant heterogeneity was observed across studies ($I^2$ = 80%). Values to the right of the line of no effect favor BM-MSC miR-21 exosomes.

| Study or Subgroup | BM-MSCs miR-21 Exosomes Mean | SD | Total | Control Mean | SD | Total | Weight | Std. mean difference IV, Fixed, 95% CI |
|---|---|---|---|---|---|---|---|---|
| Tang 2024 | 167.4 | 12.9 | 4 | 65.2 | 9.45 | 4 | 20.0% | 7.86 [2.28 , 13.44] |
| Yang 2024 | 5.78 | 0.61 | 5 | 2.68 | 0.65 | 5 | 80.0% | 4.44 [1.65 , 7.23] |
| **Total** | | | **9** | | | **9** | **100.0%** | **5.13 [2.63 , 7.62]** |

Test for overall effect: Z = 4.02 (P < 0.0001)

Heterogeneity: $Chi^2 = 1.15$, df = 1 (P = 0.28); $I^2 = 13\%$

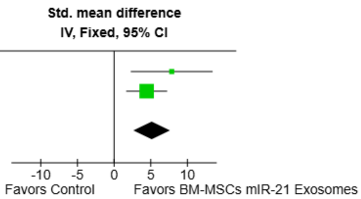


**Supplemental Fig. 50. Forest plot of the pooled effect of BM-MSC miR-21 exosomes on primary follicle count.** Forest plot showing the standardized mean difference (SMD) in primary follicle count between animals treated with bone marrow–derived mesenchymal stem cell (BM-MSC) miR-21 exosomes and control groups. Effect estimates were pooled using a fixed-effect inverse-variance model and are presented with 95% confidence intervals (CI). Individual study estimates are shown as squares, with the size of each square reflecting the study weight, and the pooled effect estimate is shown as a diamond. BM-MSC miR-21 exosomes were associated with a significantly higher primary follicle count compared with controls (SMD = 5.13, 95% CI 2.63 to 7.62; P < 0.0001). No significant heterogeneity was observed across studies ($I^2 = 13\%$). Values to the right of the line of no effect favor BM-MSC miR-21 exosomes.

| Study or Subgroup | BM-MSCs miR-21 Exosomes Mean | SD | Total | Control Mean | SD | Total | Weight | Std. mean difference IV, Random, 95% CI |
|---|---|---|---|---|---|---|---|---|
| Tang 2024 | 117.42 | 7.58 | 4 | 43.56 | 6.82 | 4 | 40.3% | 8.91 [2.63 , 15.19] |
| Yang 2024 | 4.28 | 0.7 | 5 | 2.68 | 0.82 | 5 | 59.7% | 1.90 [0.26 , 3.53] |
| **Total (Wald[a])** | | | **9** | | | **9** | **100.0%** | **4.72 [-2.02 , 11.46]** |

Test for overall effect: Z = 1.37 (P = 0.17)

Heterogeneity: $Tau^2$ (DL[b]) = 19.10; $Chi^2 = 4.48$, df = 1 (P = 0.03); $I^2 = 78\%$

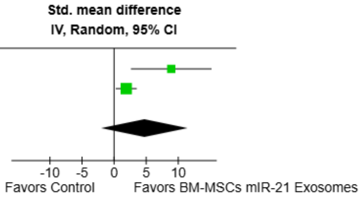


**Footnotes**
[a]CI calculated by Wald-type method.
[b]$Tau^2$ calculated by DerSimonian and Laird method.

**Supplemental Fig. 51. Forest plot of the pooled effect of BM-MSC miR-21 exosomes on secondary follicle count.** Forest plot showing the standardized mean difference (SMD) in secondary follicle count between animals treated with bone marrow–derived mesenchymal stem cell (BM-MSC) miR-21 exosomes and control groups. Effect estimates were pooled using a random-effects inverse-variance model and are presented with 95% confidence intervals (CI). Individual study estimates are shown as squares, with the size of each square reflecting the study weight, and the pooled effect estimate is shown as a diamond. BM-MSC miR-21 exosomes were not associated with a significant difference in secondary follicle count compared with controls (SMD = 4.72, 95% CI −2.02 to 11.46; P = 0.17). Significant heterogeneity was observed across studies ($I^2 = 78\%$). Values to the right of the line of no effect favor BM-MSC miR-21 exosomes.

| | BM-MSCs miR-21 Exosomes | | | Control | | | | Std. mean difference |
|---|---|---|---|---|---|---|---|---|
| Study or Subgroup | Mean | SD | Total | Mean | SD | Total | Weight | IV, Fixed, 95% CI |
| Tang 2024 | 28.15 | 9.25 | 4 | 5.27 | 1.54 | 4 | 45.0% | 3.00 [0.51 , 5.49] |
| Yang 2024 | 1.59 | 0.35 | 5 | 0.37 | 0.31 | 5 | 55.0% | 3.33 [1.08 , 5.58] |
| **Total** | | | **9** | | | **9** | **100.0%** | **3.18 [1.52 , 4.85]** |

Test for overall effect: Z = 3.74 (P = 0.0002)

Heterogeneity: Chi² = 0.04, df = 1 (P = 0.85); I² = 0%

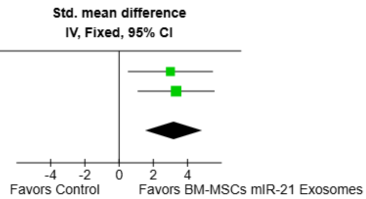


**Supplemental Fig. 52. Forest plot of the pooled effect of BM-MSC miR-21 exosomes on mature follicle count.** Forest plot showing the standardized mean difference (SMD) in mature follicle count between animals treated with bone marrow–derived mesenchymal stem cell (BM-MSC) miR-21 exosomes and control groups. Effect estimates were pooled using a fixed-effect inverse-variance model and are presented with 95% confidence intervals (CI). Individual study estimates are shown as squares, with the size of each square reflecting the study weight, and the pooled effect estimate is shown as a diamond. BM-MSC miR-21 exosomes were associated with a significantly higher mature follicle count compared with controls (SMD = 3.18, 95% CI 1.52 to 4.85; P = 0.0002). No significant heterogeneity was observed across studies (I² = 0%). Values to the right of the line of no effect favor BM-MSC miR-21 exosomes..

| | BM-MSCs miR-21 Exosomes | | | Control | | | | Std. mean difference |
|---|---|---|---|---|---|---|---|---|
| Study or Subgroup | Mean | SD | Total | Mean | SD | Total | Weight | IV, Random, 95% CI |
| Tang 2024 | 566.42 | 32.58 | 4 | 218.05 | 25.06 | 4 | 33.6% | 10.42 [3.12 , 17.72] |
| Yang 2024 | 17.36 | 1.64 | 5 | 8.9 | 1.83 | 5 | 66.4% | 4.40 [1.63 , 7.17] |
| **Total (Wald[a])** | | | **9** | | | **9** | **100.0%** | **6.42 [0.84 , 12.00]** |

Test for overall effect: Z = 2.26 (P = 0.02)

Heterogeneity: Tau² (REML[b]) = 10.21; Chi² = 2.29, df = 1 (P = 0.13); I² = 56%

**Footnotes**
[a]CI calculated by Wald-type method.
[b]Tau² calculated by Restricted Maximum-Likelihood method.

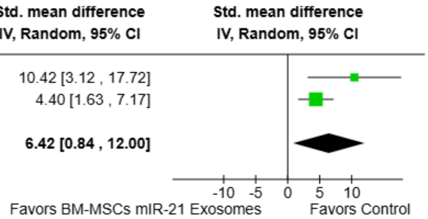


**Supplemental Fig. 53. Forest plot of the pooled effect of BM-MSC miR-21 exosomes on total follicle count.** Forest plot showing the standardized mean difference (SMD) in total follicle count between animals treated with bone marrow–derived mesenchymal stem cell (BM-MSC) miR-21 exosomes and control groups. Effect estimates were pooled using a random-effects inverse-variance model and are presented with 95% confidence intervals (CI). Individual study estimates are shown as squares, with the size of each square reflecting the study weight, and the pooled effect estimate is shown as a diamond. BM-MSC miR-21 exosomes were associated with a significantly higher total follicle count compared with controls (SMD = 6.42, 95% CI 0.84 to 12.00; P = 0.02). Significant heterogeneity was observed across studies (I² = 56%). Values to the right of the line of no effect favor BM-MSC miR-21 exosomes.

#### Funnel Plots

**Supplemental Fig. 54. Funnel Plot of BM-MSCs Therapy Analysis.** a. Estradiol **(**E2), b. (FSH), c. Anti-Müllerian Hormone (AMH), d. Primordial follicles, e. Primary Follicles, f. Secondary follicles, g. Antral Follicles, h. Total Follicles,

a. Estradiol (E2)

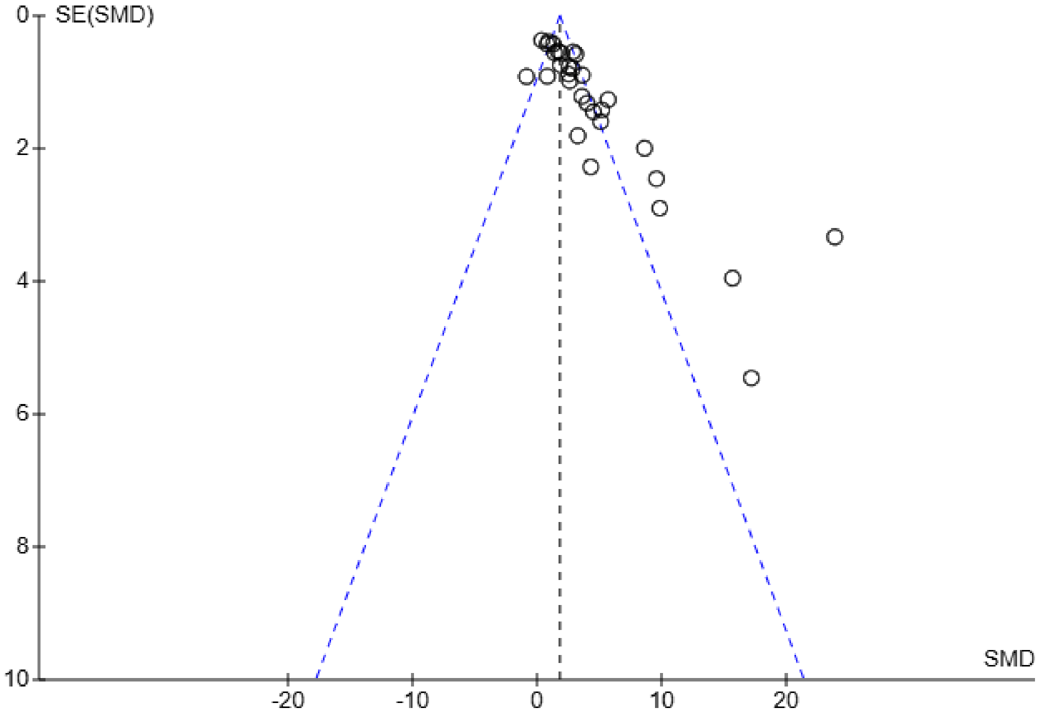


b. FSH

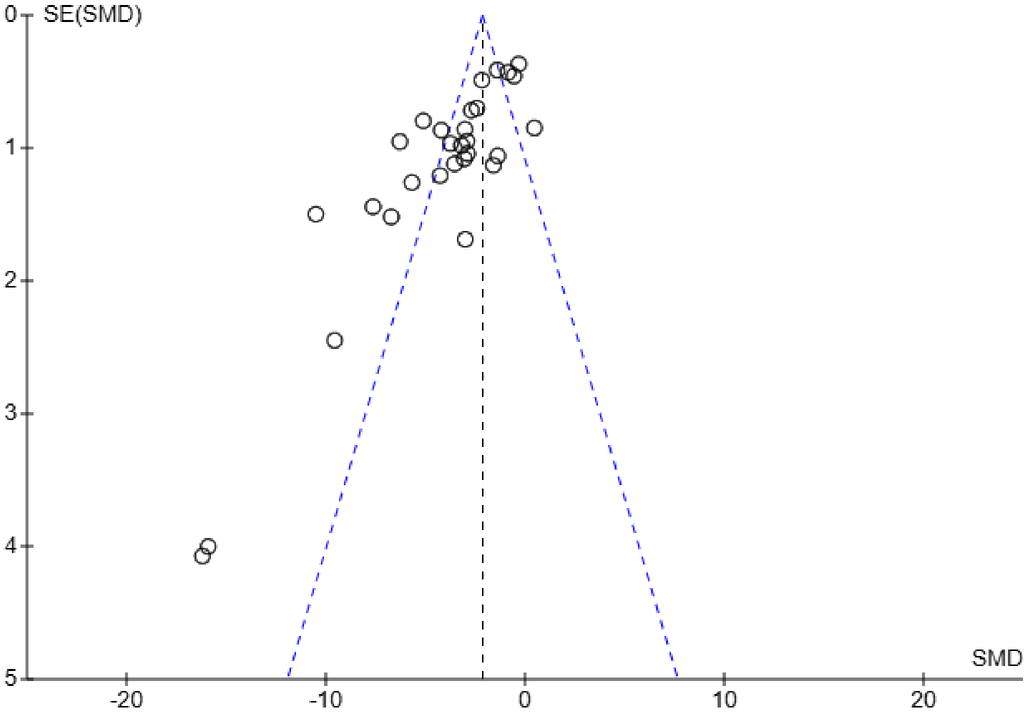


c. AMH

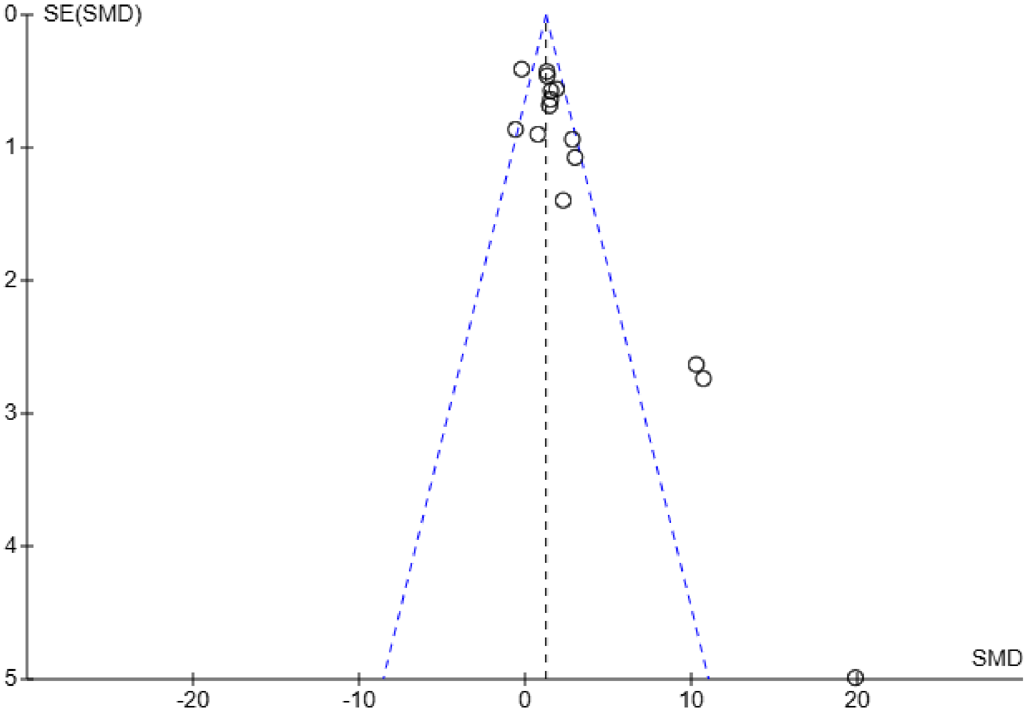


d. Primordial Follicles

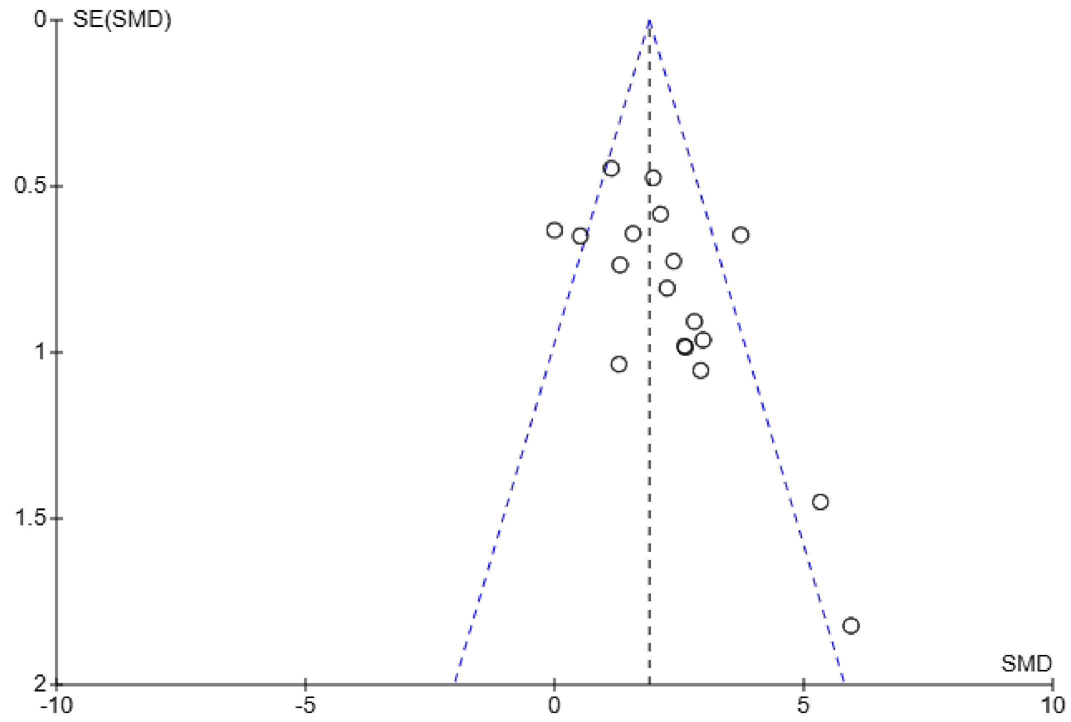


e. Primary Follicles

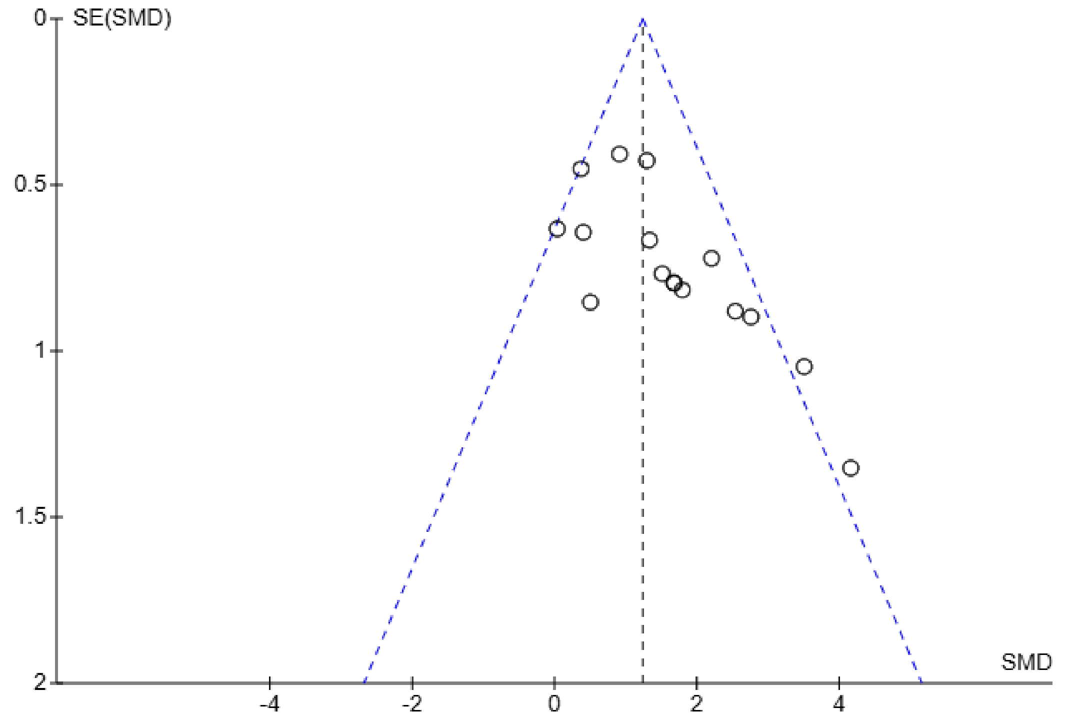


f. Secondary Follicles

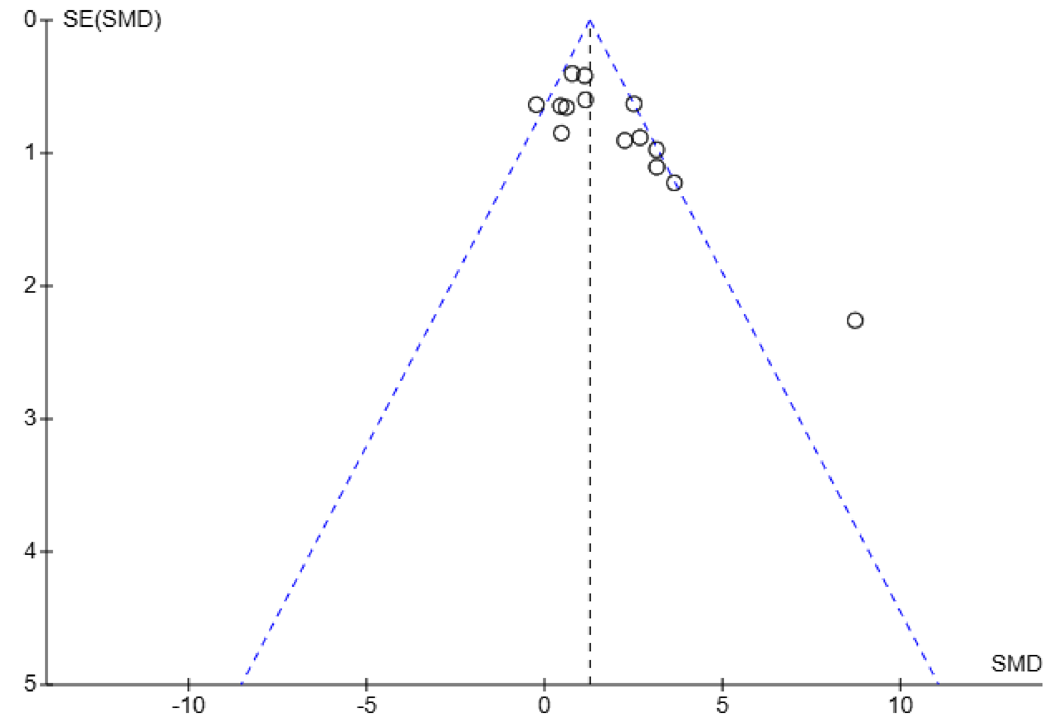

g. Antral Follicles

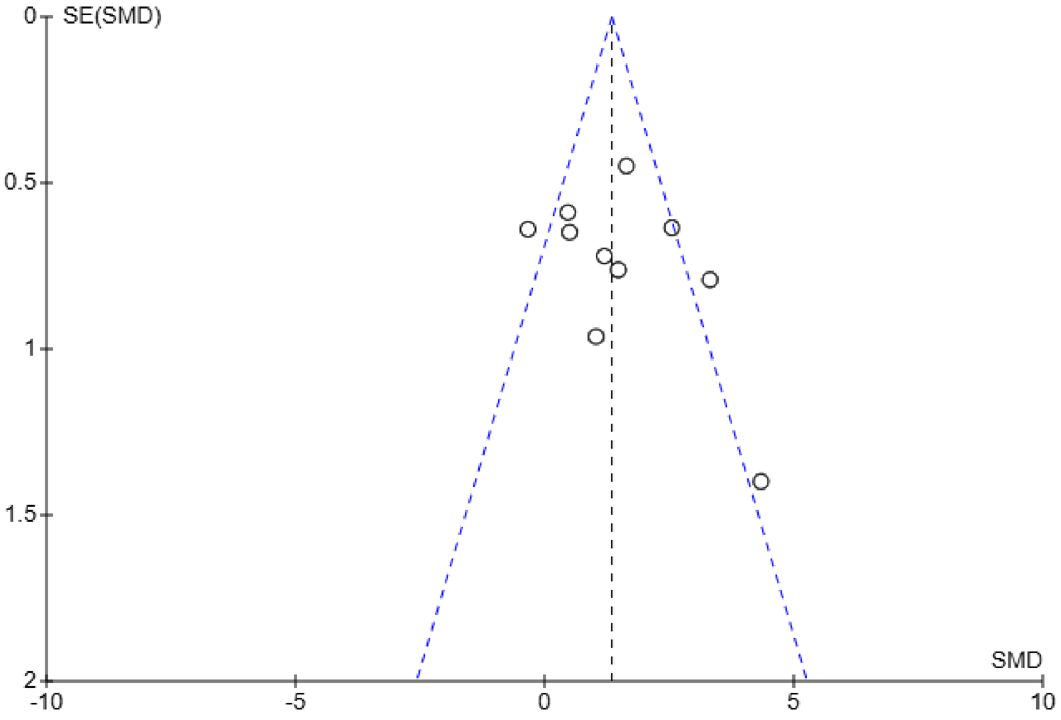

h. Total Follicles

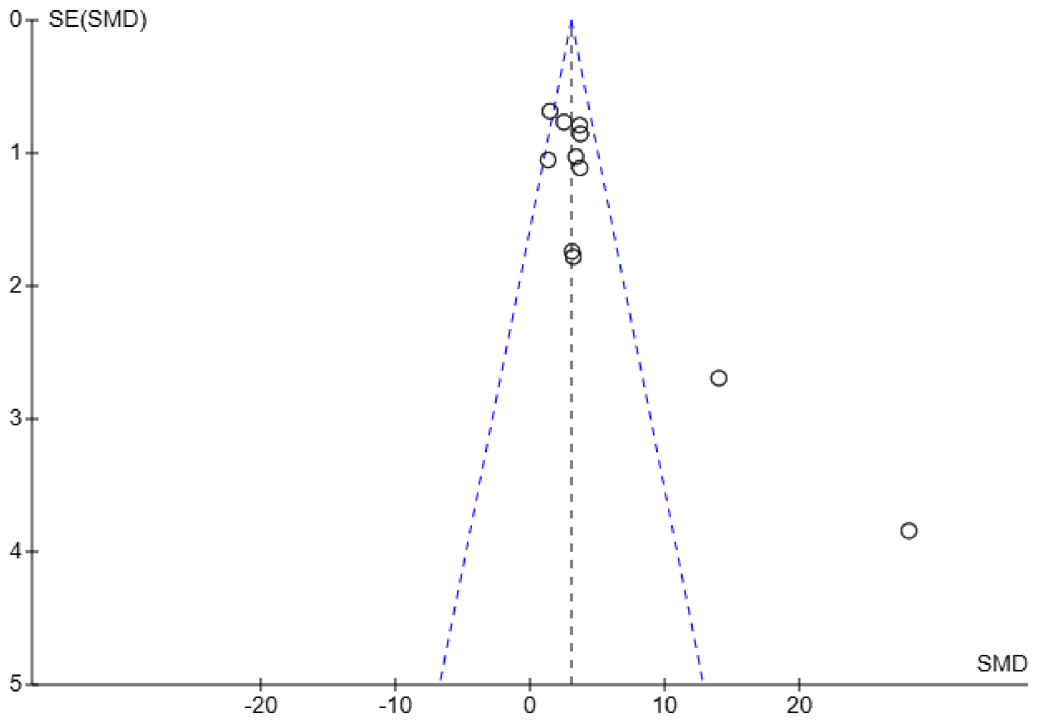

SE(SMD)
0
1
2
3
4
5
SMD
-20
-10
0
10
20

Sensitivity analysis assessing factors contributing to heterogeneity.

| Study or Subgroup | BM-MSCs Therapy Mean | SD | Total | Control Mean | SD | Total | Weight | Std. mean difference IV, Random, 95% CI |
|---|---|---|---|---|---|---|---|---|
| **6.1.1 Intraovarian** | | | | | | | | |
| Chen 2018[a] | 35.96 | 1.98 | 15 | 29.56 | 2.29 | 15 | 4.4% | 2.91 [1.84 , 3.97] |
| Fu 2008[a] | 38.13 | 38.42 | 15 | 24.7 | 28.93 | 15 | 4.7% | 0.38 [-0.34 , 1.11] |
| Fu 2017[a] | 34.92 | 9.85 | 15 | 26.72 | 7 | 15 | 4.6% | 0.93 [0.17 , 1.69] |
| Khanmohammadi 2018[b] | 34.45 | 3.33 | 20 | 20.89 | 5.65 | 10 | 4.4% | 3.13 [1.99 , 4.27] |
| Ma 2023[a] | 166.47 | 31.77 | 5 | 53.53 | 2.35 | 5 | 2.8% | 4.53 [1.69 , 7.36] |
| Maqbool 2022[a] | 13 | 2.24 | 5 | 5.67 | 1.3 | 5 | 3.2% | 3.62 [1.23 , 6.00] |
| Mohamed 2018[a] | 13.47 | 2.99 | 6 | 6.97 | 3.39 | 6 | 4.1% | 1.88 [0.42 , 3.33] |
| Sameni 2019[a] | 34 | 5.06 | 10 | 23.98 | 6.32 | 10 | 4.4% | 1.68 [0.63 , 2.73] |
| Xie 2024[c] | 63.59 | 5.13 | 3 | 48.21 | 1.38 | 3 | 2.3% | 3.28 [-0.27 , 6.82] |
| Yamchi 2023[a] | 4.45 | 0.7 | 3 | 1.78 | 0.01 | 3 | 1.7% | 4.31 [-0.15 , 8.78] |
| Yang 2024[d] | 68.31 | 11.22 | 10 | 41.33 | 6.12 | 5 | 4.0% | 2.56 [1.04 , 4.07] |
| **Subtotal (Wald[e])** | | | **107** | | | **92** | **40.7%** | **2.23 [1.43 , 3.04]** |
| Test for overall effect: Z = 5.44 (P < 0.00001) | | | | | | | | |
| Heterogeneity: Tau$^2$ (DL[f]) = 1.12; Chi$^2$ = 37.24, df = 10 (P < 0.0001); $I^2$ = 73% | | | | | | | | |
| **6.1.2 Intraperitoneal** | | | | | | | | |
| Elsherbiny 2024[c] | 54.55 | 28.4 | 10 | 13.47 | 5.5 | 10 | 4.4% | 1.92 [0.82 , 3.02] |
| Tang 2024[d] | 55.95 | 2.84 | 8 | 45.01 | 5.83 | 4 | 3.8% | 2.54 [0.81 , 4.26] |
| Yang 2020[b] | 50.37 | 7.31 | 20 | 36.98 | 14.26 | 10 | 4.6% | 1.29 [0.46 , 2.13] |
| **Subtotal (Wald[e])** | | | **38** | | | **24** | **12.8%** | **1.65 [1.03 , 2.28]** |
| Test for overall effect: Z = 5.22 (P < 0.00001) | | | | | | | | |
| Heterogeneity: Tau$^2$ (DL[f]) = 0.00; Chi$^2$ = 1.96, df = 2 (P = 0.38); $I^2$ = 0% | | | | | | | | |
| **6.1.3 Intravenous** | | | | | | | | |
| Abd-Allah 2013[a] | 47.03 | 0.91 | 15 | 21 | 1.19 | 15 | 1.0% | 23.91 [17.38 , 30.44] |
| Afifi 2013[a] | 23 | 3.39 | 8 | 14 | 2.36 | 7 | 4.0% | 2.86 [1.30 , 4.43] |
| Atwa 2025[a] | 36.2 | 1.32 | 7 | 16.9 | 2.65 | 7 | 2.0% | 8.63 [4.72 , 12.54] |
| Badawy 2017[a] | 249.41 | 6.91 | 5 | 161.22 | 9.11 | 5 | 1.2% | 9.85 [4.17 , 15.54] |
| Bahrehbar 2020[a] | 74.35 | 10.62 | 5 | 32.54 | 7.96 | 5 | 3.0% | 4.02 [1.44 , 6.61] |
| Bao 2018[a] | 26.4 | 2.55 | 8 | 17.6 | 1.98 | 8 | 3.8% | 3.64 [1.89 , 5.40] |
| El-Derany 2021[a] | 5.73 | 0.18 | 8 | 4.45 | 0.24 | 8 | 3.1% | 5.70 [3.23 , 8.18] |
| Gabr 2016[a] | 43.55 | 1.94 | 6 | 29.88 | 2.83 | 6 | 2.9% | 5.20 [2.42 , 7.98] |
| Li 2024[c] | 51.66 | 14.65 | 12 | 40.23 | 12.12 | 12 | 4.6% | 0.82 [-0.02 , 1.66] |
| Liu 2014[a] | 52.64 | 3.53 | 5 | 35.85 | 2.28 | 5 | 2.6% | 5.10 [1.97 , 8.23] |
| Lu 2024[a] | 7.62 | 2.06 | 9 | 5.12 | 1.07 | 8 | 4.4% | 1.42 [0.32 , 2.52] |
| Nabil Salama 2024[a] | 117.89 | 7.78 | 6 | 55.05 | 3.56 | 6 | 1.6% | 9.59 [4.77 , 14.40] |
| Park 2023[a] | 5.75 | 2 | 3 | 4.18 | 0.79 | 3 | 3.8% | 0.83 [-0.96 , 2.61] |
| Park 2023[c] | 4.57 | 0.4 | 3 | 5.43 | 1.08 | 3 | 3.8% | -0.84 [-2.64 , 0.95] |
| Sun 2019[c] | 32 | 2.88 | 5 | 22.4 | 3.68 | 5 | 3.6% | 2.62 [0.70 , 4.55] |
| Xing 2023[c] | 57.9 | 1.21 | 6 | 35.79 | 0.95 | 3 | 0.4% | 17.21 [6.52 , 27.91] |
| Zhang 2024[c] | 55.2 | 1 | 6 | 39.8 | 0.8 | 6 | 0.7% | 15.70 [7.95 , 23.44] |
| **Subtotal (Wald[e])** | | | **117** | | | **112** | **46.5%** | **4.96 [3.39 , 6.53]** |
| Test for overall effect: Z = 6.19 (P < 0.00001) | | | | | | | | |
| Heterogeneity: Tau$^2$ (DL[f]) = 7.94; Chi$^2$ = 134.50, df = 16 (P < 0.00001); $I^2$ = 88% | | | | | | | | |
| **Total (Wald[e])** | | | **262** | | | **228** | **100.0%** | **3.11 [2.38 , 3.84]** |

Test for overall effect: Z = 8.39 (P < 0.00001)
Test for subgroup differences: Chi$^2$ = 14.81, df = 2 (P = 0.0006), $I^2$ = 86.5%
Heterogeneity: Tau$^2$ (DL[f]) = 2.81; Chi$^2$ = 177.98, df = 30 (P < 0.00001); $I^2$ = 83%

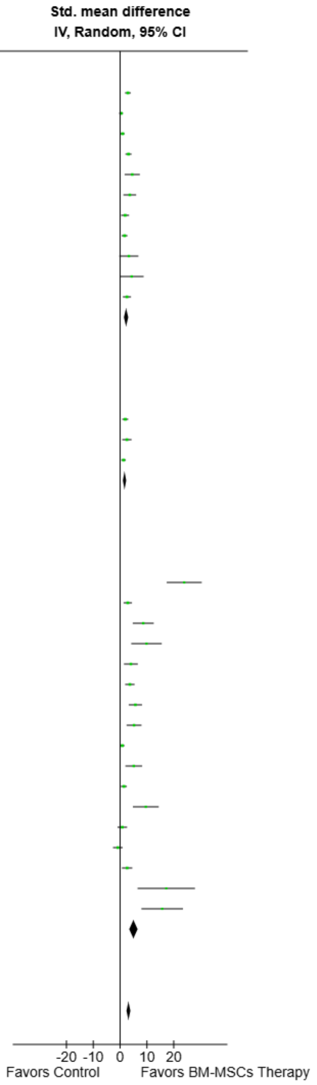


**Footnotes**
[a]BM-MSCs Transplantation
[b]BM-MSCs Transplantation and Secretome
[c]BM-MSCs Secretome
[d]BM-MSCs Secretome and miR-21 Exosomes
[e]CI calculated by Wald-type method.
[f]Tau$^2$ calculated by DerSimonian and Laird method.

**Supplemental Fig. 55. Subgroup analysis of BM-MSC Therapy on serum estradiol (E2) levels by route of administration.**

| Study or Subgroup | BM-MSCs Therapy Mean | SD | Total | Control Mean | SD | Total | Weight | Std. mean difference IV, Random, 95% CI |
|---|---|---|---|---|---|---|---|---|
| **6.2.1 Mice** | | | | | | | | |
| Badawy 2017[a] | 249.41 | 6.91 | 5 | 161.22 | 9.11 | 5 | 1.2% | 9.85 [4.17 , 15.54] |
| Bahrehbar 2020[a] | 74.35 | 10.62 | 5 | 32.54 | 7.96 | 5 | 3.0% | 4.02 [1.44 , 6.61] |
| Bao 2018[a] | 26.4 | 2.55 | 8 | 17.6 | 1.98 | 8 | 3.8% | 3.64 [1.89 , 5.40] |
| Ma 2023[a] | 166.47 | 31.77 | 5 | 53.53 | 2.35 | 5 | 2.8% | 4.53 [1.69 , 7.36] |
| Mohamed 2018[a] | 13.47 | 2.99 | 6 | 6.97 | 3.39 | 6 | 4.1% | 1.88 [0.42 , 3.33] |
| Nabil Salama 2024[a] | 117.89 | 7.78 | 6 | 55.05 | 3.56 | 6 | 1.6% | 9.59 [4.77 , 14.40] |
| Park 2023[b] | 4.57 | 0.4 | 3 | 5.43 | 1.08 | 3 | 3.8% | -0.84 [-2.64 , 0.95] |
| Park 2023[a] | 5.75 | 2 | 3 | 4.18 | 0.79 | 3 | 3.8% | 0.83 [-0.96 , 2.61] |
| Sun 2019[b] | 32 | 2.88 | 5 | 22.4 | 3.68 | 5 | 3.6% | 2.62 [0.70 , 4.55] |
| Xie 2024[b] | 63.59 | 5.13 | 3 | 48.21 | 1.38 | 3 | 2.3% | 3.28 [-0.27 , 6.82] |
| Yang 2024[c] | 68.31 | 11.22 | 10 | 41.33 | 6.12 | 5 | 4.0% | 2.56 [1.04 , 4.07] |
| **Subtotal (Wald[d])** | | | **59** | | | **54** | **34.0%** | **3.00 [1.70 , 4.30]** |
| Test for overall effect: Z = 4.52 (P < 0.00001) | | | | | | | | |
| Heterogeneity: Tau² (DL[e]) = 3.17; Chi² = 37.11, df = 10 (P < 0.0001); I² = 73% | | | | | | | | |
| **6.2.2 Rabbit** | | | | | | | | |
| Abd-Allah 2013[a] | 47.03 | 0.91 | 15 | 21 | 1.19 | 15 | 1.0% | 23.91 [17.38 , 30.44] |
| **Subtotal** | | | **15** | | | **15** | **1.0%** | **23.91 [17.38 , 30.44]** |
| Test for overall effect: Z = 7.18 (P < 0.00001) | | | | | | | | |
| Heterogeneity: Not applicable | | | | | | | | |
| **6.2.3 Rats** | | | | | | | | |
| Afifi 2013[a] | 23 | 3.39 | 8 | 14 | 2.36 | 7 | 4.0% | 2.86 [1.30 , 4.43] |
| Atwa 2025[a] | 36.2 | 1.32 | 7 | 16.9 | 2.65 | 7 | 2.0% | 8.63 [4.72 , 12.54] |
| Chen 2018[a] | 35.96 | 1.98 | 15 | 29.56 | 2.29 | 15 | 4.4% | 2.91 [1.84 , 3.97] |
| El-Derany 2021[a] | 5.73 | 0.18 | 8 | 4.45 | 0.24 | 8 | 3.1% | 5.70 [3.23 , 8.18] |
| Elsherbiny 2024[b] | 54.55 | 28.4 | 10 | 13.47 | 5.5 | 10 | 4.4% | 1.92 [0.82 , 3.02] |
| Fu 2008[a] | 38.13 | 38.42 | 15 | 24.7 | 28.93 | 15 | 4.7% | 0.38 [-0.34 , 1.11] |
| Fu 2017[a] | 34.92 | 9.85 | 15 | 26.72 | 7 | 15 | 4.6% | 0.93 [0.17 , 1.69] |
| Gabr 2016[a] | 43.55 | 1.94 | 6 | 29.88 | 2.83 | 6 | 2.9% | 5.20 [2.42 , 7.98] |
| Khanmohammadi 2018[f] | 34.45 | 3.33 | 20 | 20.89 | 5.65 | 10 | 4.4% | 3.13 [1.99 , 4.27] |
| Li 2024[b] | 51.66 | 14.65 | 12 | 40.23 | 12.12 | 12 | 4.6% | 0.82 [-0.02 , 1.66] |
| Liu 2014[a] | 52.64 | 3.53 | 5 | 35.85 | 2.28 | 5 | 2.6% | 5.10 [1.97 , 8.23] |
| Lu 2024[a] | 7.62 | 2.06 | 9 | 5.12 | 1.07 | 8 | 4.4% | 1.42 [0.32 , 2.52] |
| Maqbool 2022[a] | 13 | 2.24 | 5 | 5.67 | 1.3 | 5 | 3.2% | 3.62 [1.23 , 6.00] |
| Sameni 2019[a] | 34 | 5.06 | 10 | 23.98 | 6.32 | 10 | 4.4% | 1.68 [0.63 , 2.73] |
| Tang 2024[c] | 55.95 | 2.84 | 8 | 45.01 | 5.83 | 4 | 3.8% | 2.54 [0.81 , 4.26] |
| Xing 2023[b] | 57.9 | 1.21 | 6 | 35.79 | 0.95 | 3 | 0.4% | 17.21 [6.52 , 27.91] |
| Yamchi 2023[a] | 4.45 | 0.7 | 3 | 1.78 | 0.01 | 3 | 1.7% | 4.31 [-0.15 , 8.78] |
| Yang 2020[f] | 50.37 | 7.31 | 20 | 36.98 | 14.26 | 10 | 4.6% | 1.29 [0.46 , 2.13] |
| Zhang 2024[b] | 55.2 | 1 | 6 | 39.8 | 0.8 | 6 | 0.7% | 15.70 [7.95 , 23.44] |
| **Subtotal (Wald[d])** | | | **188** | | | **159** | **65.0%** | **2.68 [1.93 , 3.44]** |
| Test for overall effect: Z = 6.97 (P < 0.00001) | | | | | | | | |
| Heterogeneity: Tau² (DL[e]) = 1.79; Chi² = 93.31, df = 18 (P < 0.00001); I² = 81% | | | | | | | | |
| **Total (Wald[d])** | | | **262** | | | **228** | **100.0%** | **3.11 [2.38 , 3.84]** |

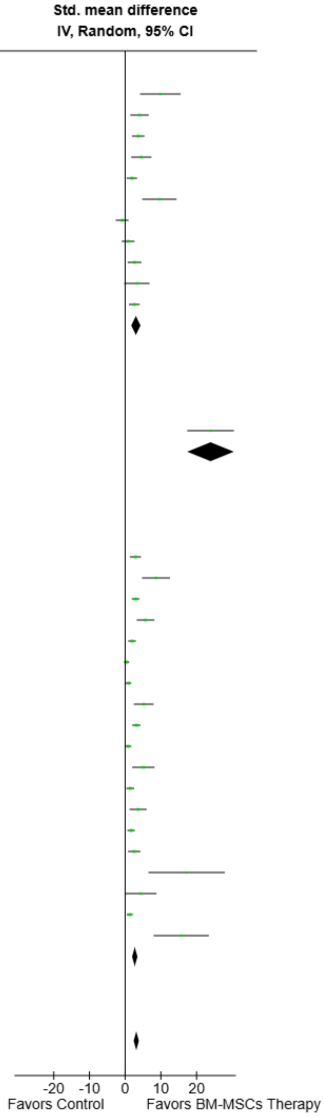


Test for overall effect: Z = 8.39 (P < 0.00001)
Test for subgroup differences: Chi² = 40.05, df = 2 (P < 0.00001), I² = 95.0%
Heterogeneity: Tau² (DL[e]) = 2.81; Chi² = 177.98, df = 30 (P < 0.00001); I² = 83%

**Footnotes**
[a]BM-MSCs Transplantation
[b]BM-MSCs Secretome
[c]BM-MSCs Secretome and miR-21 Exosomes
[d]CI calculated by Wald-type method.
[e]Tau² calculated by DerSimonian and Laird method.
[f]BM-MSCs Transplantation and Secretome

**Supplemental Fig. 56. Subgroup analysis of BM-MSC Therapy on serum estradiol (E2) levels by animal model.**

| Study or Subgroup | BM-MSCs Therapy Mean | SD | Total | Control Mean | SD | Total | Weight | Std. mean difference IV, Random, 95% CI |
|---|---|---|---|---|---|---|---|---|
| **6.3.1 Enzyme-Linked Immunosorbent Assay (ELISA)** | | | | | | | | |
| Abd-Allah 2013[a] | 47.03 | 0.91 | 15 | 21 | 1.19 | 15 | 1.0% | 23.91 [17.38 , 30.44] |
| Afifi 2013[a] | 23 | 3.39 | 8 | 14 | 2.36 | 7 | 4.0% | 2.86 [1.30 , 4.43] |
| Atwa 2025[a] | 36.2 | 1.32 | 7 | 16.9 | 2.65 | 7 | 2.0% | 8.63 [4.72 , 12.54] |
| Bahrehbar 2020[a] | 74.35 | 10.62 | 5 | 32.54 | 7.96 | 5 | 3.0% | 4.02 [1.44 , 6.61] |
| El-Derany 2021[a] | 5.73 | 0.18 | 8 | 4.45 | 0.24 | 8 | 3.1% | 5.70 [3.23 , 8.18] |
| Elsherbiny 2024[b] | 54.55 | 28.4 | 10 | 13.47 | 5.5 | 10 | 4.4% | 1.92 [0.82 , 3.02] |
| Gabr 2016[a] | 43.55 | 1.94 | 6 | 29.88 | 2.83 | 6 | 2.9% | 5.20 [2.42 , 7.98] |
| Khanmohammadi 2018[c] | 34.45 | 3.33 | 20 | 20.89 | 5.65 | 10 | 4.4% | 3.13 [1.99 , 4.27] |
| Lu 2024[a] | 7.62 | 2.06 | 9 | 5.12 | 1.07 | 8 | 4.4% | 1.42 [0.32 , 2.52] |
| Maqbool 2022[a] | 13 | 2.24 | 5 | 5.67 | 1.3 | 5 | 3.2% | 3.62 [1.23 , 6.00] |
| Mohamed 2018[a] | 13.47 | 2.99 | 6 | 6.97 | 3.39 | 6 | 4.1% | 1.88 [0.42 , 3.33] |
| Nabil Salama 2024[a] | 117.89 | 7.78 | 6 | 55.05 | 3.56 | 6 | 1.6% | 9.59 [4.77 , 14.40] |
| Park 2023[a] | 5.75 | 2 | 3 | 4.18 | 0.79 | 3 | 3.8% | 0.83 [-0.96 , 2.61] |
| Park 2023[b] | 4.57 | 0.4 | 3 | 5.43 | 1.08 | 3 | 3.8% | -0.84 [-2.64 , 0.95] |
| Sameni 2019[a] | 34 | 5.06 | 10 | 23.98 | 6.32 | 10 | 4.4% | 1.68 [0.63 , 2.73] |
| Sun 2019[b] | 32 | 2.88 | 5 | 22.4 | 3.68 | 5 | 3.6% | 2.62 [0.70 , 4.55] |
| Tang 2024[d] | 55.95 | 2.84 | 8 | 45.01 | 5.83 | 4 | 3.8% | 2.54 [0.81 , 4.26] |
| Xie 2024[b] | 63.59 | 5.13 | 3 | 48.21 | 1.38 | 3 | 2.3% | 3.28 [-0.27 , 6.82] |
| Xing 2023[b] | 57.9 | 1.21 | 6 | 35.79 | 0.95 | 3 | 0.4% | 17.21 [6.52 , 27.91] |
| Yamchi 2023[a] | 4.45 | 0.7 | 3 | 1.78 | 0.01 | 3 | 1.7% | 4.31 [-0.15 , 8.78] |
| Yang 2020[c] | 50.37 | 7.31 | 20 | 36.98 | 14.26 | 10 | 4.6% | 1.29 [0.46 , 2.13] |
| Yang 2024[d] | 68.31 | 11.22 | 10 | 41.33 | 6.12 | 5 | 4.0% | 2.56 [1.04 , 4.07] |
| Zhang 2024[b] | 55.2 | 1 | 6 | 39.8 | 0.8 | 6 | 0.7% | 15.70 [7.95 , 23.44] |
| **Subtotal (Wald[e])** | | | **182** | | | **148** | **71.3%** | **3.44 [2.50 , 4.37]** |
| Test for overall effect: Z = 7.21 (P < 0.00001) | | | | | | | | |
| Heterogeneity: $Tau^2$ (DL[f]) = 3.49; $Chi^2$ = 122.59, df = 22 (P < 0.00001); $I^2$ = 82% | | | | | | | | |
| **6.3.2 Chemiluminescence Immunoassay (CLIA)** | | | | | | | | |
| Chen 2018[a] | 35.96 | 1.98 | 15 | 29.56 | 2.29 | 15 | 4.4% | 2.91 [1.84 , 3.97] |
| Fu 2008[a] | 38.13 | 38.42 | 15 | 24.7 | 28.93 | 15 | 4.7% | 0.38 [-0.34 , 1.11] |
| Fu 2017[a] | 34.92 | 9.85 | 15 | 26.72 | 7 | 15 | 4.6% | 0.93 [0.17 , 1.69] |
| Liu 2014[a] | 52.64 | 3.53 | 5 | 35.85 | 2.28 | 5 | 2.6% | 5.10 [1.97 , 8.23] |
| Ma 2023[a] | 166.47 | 31.77 | 5 | 53.53 | 2.35 | 5 | 2.8% | 4.53 [1.69 , 7.36] |
| **Subtotal (Wald[e])** | | | **55** | | | **55** | **19.1%** | **2.21 [0.82 , 3.61]** |
| Test for overall effect: Z = 3.10 (P = 0.002) | | | | | | | | |
| Heterogeneity: $Tau^2$ (DL[f]) = 1.84; $Chi^2$ = 26.44, df = 4 (P < 0.0001); $I^2$ = 85% | | | | | | | | |
| **6.3.3 Radioimmunoassay (RIA)** | | | | | | | | |
| Badawy 2017[a] | 249.41 | 6.91 | 5 | 161.22 | 9.11 | 5 | 1.2% | 9.85 [4.17 , 15.54] |
| Bao 2018[a] | 26.4 | 2.55 | 8 | 17.6 | 1.98 | 8 | 3.8% | 3.64 [1.89 , 5.40] |
| Li 2024[b] | 51.66 | 14.65 | 12 | 40.23 | 12.12 | 12 | 4.6% | 0.82 [-0.02 , 1.66] |
| **Subtotal (Wald[e])** | | | **25** | | | **25** | **9.6%** | **3.63 [0.33 , 6.93]** |
| Test for overall effect: Z = 2.16 (P = 0.03) | | | | | | | | |
| Heterogeneity: $Tau^2$ (DL[f]) = 6.53; $Chi^2$ = 16.55, df = 2 (P = 0.0003); $I^2$ = 88% | | | | | | | | |
| **Total (Wald[e])** | | | **262** | | | **228** | **100.0%** | **3.11 [2.38 , 3.84]** |

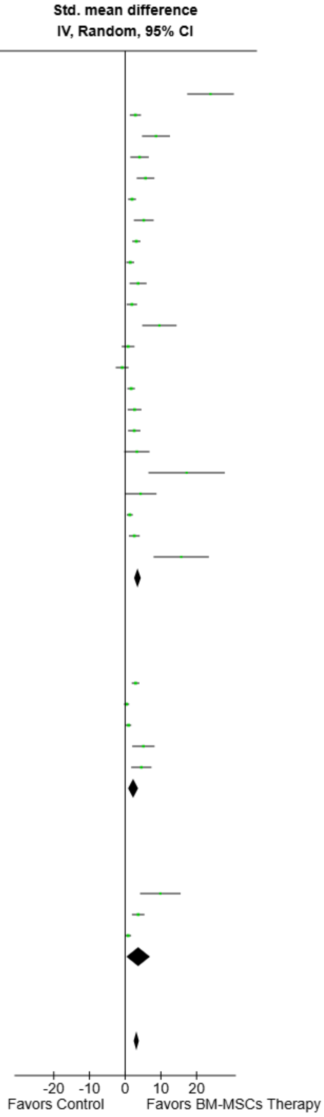


Test for overall effect: Z = 8.39 (P < 0.00001)
Test for subgroup differences: $Chi^2$ = 2.14, df = 2 (P = 0.34), $I^2$ = 6.6%
Heterogeneity: $Tau^2$ (DL[f]) = 2.81; $Chi^2$ = 177.98, df = 30 (P < 0.00001); $I^2$ = 83%

**Footnotes**
[a]BM-MSCs Transplantation
[b]BM-MSCs Secretome
[c]BM-MSCs Transplantation and Secretome
[d]BM-MSCs Secretome and mIR-21 Exosomes
[e]CI calculated by Wald-type method.
[f]$Tau^2$ calculated by DerSimonian and Laird method.

**Supplemental Fig. 57. Subgroup analysis of BM-MSC Therapy on serum estradiol (E2) levels by hormonal measurement technique.**

| Study or Subgroup | BM-MSCs Therapy Mean | SD | Total | Control Mean | SD | Total | Weight | Std. mean difference IV, Random, 95% CI |
|---|---|---|---|---|---|---|---|---|
| **6.4.1 Autoimmune** | | | | | | | | |
| El-Derany 2021[a] | 5.73 | 0.18 | 8 | 4.45 | 0.24 | 8 | 3.1% | 5.70 [3.23 , 8.18] |
| Li 2024[b] | 51.66 | 14.65 | 12 | 40.23 | 12.12 | 12 | 4.6% | 0.82 [-0.02 , 1.66] |
| Ma 2023[a] | 166.47 | 31.77 | 5 | 53.53 | 2.35 | 5 | 2.8% | 4.53 [1.69 , 7.36] |
| Xie 2024[b] | 63.59 | 5.13 | 3 | 48.21 | 1.38 | 3 | 2.3% | 3.28 [-0.27 , 6.82] |
| Yang 2024[c] | 68.31 | 11.22 | 10 | 41.33 | 6.12 | 5 | 4.0% | 2.56 [1.04 , 4.07] |
| **Subtotal (Wald[d])** | | | **38** | | | **33** | **16.8%** | **3.14 [1.23 , 5.04]** |
| Test for overall effect: Z = 3.22 (P = 0.001) | | | | | | | | |
| Heterogeneity: Tau² (DL[e]) = 3.43; Chi² = 19.94, df = 4 (P = 0.0005); I² = 80% | | | | | | | | |
| **6.4.2 Chemotherapy (Cisplatin)** | | | | | | | | |
| Nabil Salama 2024[a] | 117.89 | 7.78 | 6 | 55.05 | 3.56 | 6 | 1.6% | 9.59 [4.77 , 14.40] |
| Sun 2019[b] | 32 | 2.88 | 5 | 22.4 | 3.68 | 5 | 3.6% | 2.62 [0.70 , 4.55] |
| Tang 2024[c] | 55.95 | 2.84 | 8 | 45.01 | 5.83 | 4 | 3.8% | 2.54 [0.81 , 4.26] |
| **Subtotal (Wald[d])** | | | **19** | | | **15** | **9.0%** | **3.96 [1.20 , 6.71]** |
| Test for overall effect: Z = 2.81 (P = 0.005) | | | | | | | | |
| Heterogeneity: Tau² (DL[e]) = 4.06; Chi² = 7.61, df = 2 (P = 0.02); I² = 74% | | | | | | | | |
| **6.4.3 Chemotherapy (Cyclophosphamide)** | | | | | | | | |
| Abd-Allah 2013[a] | 47.03 | 0.91 | 15 | 21 | 1.19 | 15 | 1.0% | 23.91 [17.38 , 30.44] |
| Afifi 2013[a] | 23 | 3.39 | 8 | 14 | 2.36 | 7 | 4.0% | 2.86 [1.30 , 4.43] |
| Atwa 2025[a] | 36.2 | 1.32 | 7 | 16.9 | 2.65 | 7 | 2.0% | 8.63 [4.72 , 12.54] |
| Badawy 2017[a] | 249.41 | 6.91 | 5 | 161.22 | 9.11 | 5 | 1.2% | 9.85 [4.17 , 15.54] |
| Bahrehbar 2020[a] | 74.35 | 10.62 | 5 | 32.54 | 7.96 | 5 | 3.0% | 4.02 [1.44 , 6.61] |
| Bao 2018[a] | 26.4 | 2.55 | 8 | 17.6 | 1.98 | 8 | 3.8% | 3.64 [1.89 , 5.40] |
| Chen 2018[a] | 35.96 | 1.98 | 15 | 29.56 | 2.29 | 15 | 4.4% | 2.91 [1.84 , 3.97] |
| Elsherbiny 2024[b] | 54.55 | 28.4 | 10 | 13.47 | 5.5 | 10 | 4.4% | 1.92 [0.82 , 3.02] |
| Fu 2008[a] | 38.13 | 38.42 | 15 | 24.7 | 28.93 | 15 | 4.7% | 0.38 [-0.34 , 1.11] |
| Fu 2017[a] | 34.92 | 9.85 | 15 | 26.72 | 7 | 15 | 4.6% | 0.93 [0.17 , 1.69] |
| Gabr 2016[a] | 43.55 | 1.94 | 6 | 29.88 | 2.83 | 6 | 2.9% | 5.20 [2.42 , 7.98] |
| Khanmohammadi 2018[f] | 34.45 | 3.33 | 20 | 20.89 | 5.65 | 10 | 4.4% | 3.13 [1.99 , 4.27] |
| Lu 2024[a] | 7.62 | 2.06 | 9 | 5.12 | 1.07 | 8 | 4.4% | 1.42 [0.32 , 2.52] |
| Maqbool 2022[a] | 13 | 2.24 | 5 | 5.67 | 1.3 | 5 | 3.2% | 3.62 [1.23 , 6.00] |
| Mohamed 2018[a] | 13.47 | 2.99 | 6 | 6.97 | 3.39 | 6 | 4.1% | 1.88 [0.42 , 3.33] |
| Park 2023[a] | 5.75 | 2 | 3 | 4.18 | 0.79 | 3 | 3.8% | 0.83 [-0.96 , 2.61] |
| Park 2023[b] | 4.57 | 0.4 | 3 | 5.43 | 1.08 | 3 | 3.8% | -0.84 [-2.64 , 0.95] |
| Sameni 2019[a] | 34 | 5.06 | 10 | 23.98 | 6.32 | 10 | 4.4% | 1.68 [0.63 , 2.73] |
| Xing 2023[b] | 57.9 | 1.21 | 6 | 35.79 | 0.95 | 3 | 0.4% | 17.21 [6.52 , 27.91] |
| Yamchi 2023[a] | 4.45 | 0.7 | 3 | 1.78 | 0.01 | 3 | 1.7% | 4.31 [-0.15 , 8.78] |
| Yang 2020[f] | 50.37 | 7.31 | 20 | 36.98 | 14.26 | 10 | 4.6% | 1.29 [0.46 , 2.13] |
| Zhang 2024[b] | 55.2 | 1 | 6 | 39.8 | 0.8 | 6 | 0.7% | 15.70 [7.95 , 23.44] |
| **Subtotal (Wald[d])** | | | **200** | | | **175** | **71.6%** | **2.99 [2.11 , 3.87]** |
| Test for overall effect: Z = 6.68 (P < 0.00001) | | | | | | | | |
| Heterogeneity: Tau² (DL[e]) = 2.96; Chi² = 142.33, df = 21 (P < 0.00001); I² = 85% | | | | | | | | |
| **6.4.4 Chemotherapy (Cyclophosphamide + Cisplatin)** | | | | | | | | |
| Liu 2014[a] | 52.64 | 3.53 | 5 | 35.85 | 2.28 | 5 | 2.6% | 5.10 [1.97 , 8.23] |
| **Subtotal** | | | **5** | | | **5** | **2.6%** | **5.10 [1.97 , 8.23]** |
| Test for overall effect: Z = 3.20 (P = 0.001) | | | | | | | | |
| Heterogeneity: Not applicable | | | | | | | | |
| **Total (Wald[d])** | | | **262** | | | **228** | **100.0%** | **3.11 [2.38 , 3.84]** |

Test for overall effect: Z = 8.39 (P < 0.00001)
Test for subgroup differences: Chi² = 1.93, df = 3 (P = 0.59), I² = 0%
Heterogeneity: Tau² (DL[e]) = 2.81; Chi² = 177.98, df = 30 (P < 0.00001); I² = 83%

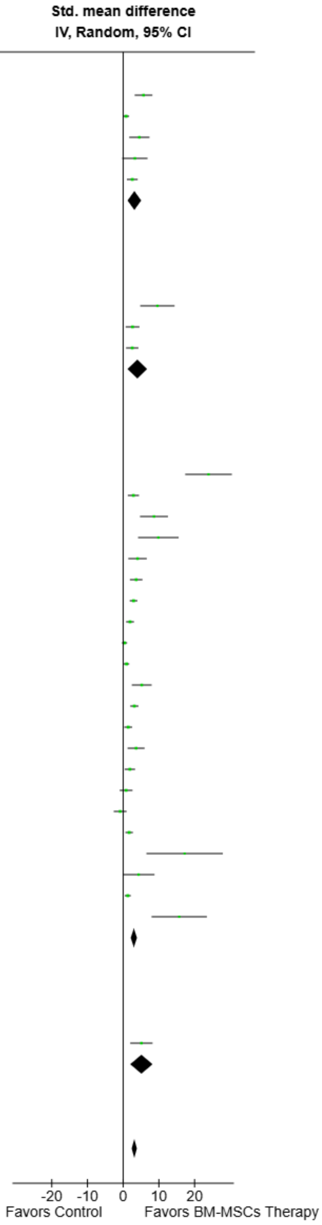


**Footnotes**
[a]BM-MSCs Transplantation
[b]BM-MSCs Secretome
[c]BM-MSCs Secretome and mIR-21 Exosomes
[d]CI calculated by Wald-type method.
[e]Tau² calculated by DerSimonian and Laird method.
[f]BM-MSCs Transplantation and Secretome

**Supplemental Fig. 58. Subgroup analysis of BM-MSC Therapy on serum estradiol (E2) levels by POI induction model.**

| Study or Subgroup | BM-MSCs Therapy Mean | SD | Total | Control Mean | SD | Total | Weight | Std. mean difference IV, Random, 95% CI |
|---|---|---|---|---|---|---|---|---|
| **6.5.1 Intraovarian** | | | | | | | | |
| Chen 2018[a] | 15.57 | 0.97 | 15 | 22.64 | 1.64 | 15 | 4.0% | -5.11 [-6.67 , -3.55] |
| Fu 2008[a] | 8.7 | 18.13 | 15 | 14.7 | 18.75 | 15 | 4.5% | -0.32 [-1.04 , 0.40] |
| Fu 2017[a] | 18.04 | 3.29 | 15 | 22.96 | 3.58 | 15 | 4.4% | -1.39 [-2.20 , -0.58] |
| Khanmohammadi 2018[b] | 7.39 | 0.76 | 20 | 12.58 | 0.89 | 10 | 3.7% | -6.28 [-8.15 , -4.41] |
| Ma 2023[a] | 73.88 | 3.78 | 5 | 92.17 | 1.64 | 15 | 3.0% | -7.63 [-10.46 , -4.81] |
| Maqbool 2022[a] | 10.33 | 1.27 | 5 | 14.67 | 1.3 | 5 | 3.5% | -3.05 [-5.17 , -0.93] |
| Mohamed 2018[a] | 10 | 1.21 | 6 | 18.67 | 3.34 | 6 | 3.7% | -3.19 [-5.11 , -1.26] |
| Sameni 2019[a] | 7.77 | 0.65 | 10 | 11.19 | 0.89 | 10 | 3.9% | -4.20 [-5.90 , -2.51] |
| Xie 2024[c] | 66.22 | 5.74 | 3 | 84.78 | 4.01 | 3 | 2.7% | -3.00 [-6.31 , 0.31] |
| Yamchi 2023[a] | 2.93 | 0.13 | 3 | 3.16 | 0.1 | 3 | 3.5% | -1.59 [-3.80 , 0.63] |
| Yang 2024[d] | 58.24 | 8.91 | 10 | 89.2 | 4.26 | 5 | 3.7% | -3.74 [-5.64 , -1.85] |
| **Subtotal (Wald[e])** | | | **107** | | | **102** | **40.5%** | **-3.48 [-4.85 , -2.11]** |
| Test for overall effect: Z = 4.98 (P < 0.00001) | | | | | | | | |
| Heterogeneity: Tau² (DL[f]) = 4.38; Chi² = 85.93, df = 10 (P < 0.00001); I² = 88% | | | | | | | | |
| **6.5.2 Intraperitoneal** | | | | | | | | |
| Elsherbiny 2024[c] | 46.41 | 7.68 | 10 | 98.36 | 126.71 | 10 | 4.4% | -0.55 [-1.45 , 0.34] |
| Tang 2024[d] | 12.32 | 0.99 | 8 | 15.18 | 0.68 | 4 | 3.7% | -2.91 [-4.77 , -1.05] |
| Yang 2020[b] | 14.96 | 2.15 | 20 | 19.01 | 0.76 | 10 | 4.3% | -2.16 [-3.12 , -1.20] |
| **Subtotal (Wald[e])** | | | **38** | | | **24** | **12.5%** | **-1.73 [-3.10 , -0.37]** |
| Test for overall effect: Z = 2.49 (P = 0.01) | | | | | | | | |
| Heterogeneity: Tau² (DL[f]) = 1.06; Chi² = 8.29, df = 2 (P = 0.02); I² = 76% | | | | | | | | |
| **6.5.3 Intravenous** | | | | | | | | |
| Abd-Allah 2013 | 5.9 | 0.9 | 15 | 19.9 | 1.6 | 15 | 2.9% | -10.49 [-13.43 , -7.56] |
| Afifi 2013 | 7.7 | 0.21 | 8 | 29.3 | 4.45 | 7 | 2.9% | -6.72 [-9.69 , -3.74] |
| Atwa 2025[a] | 0.5 | 0.11 | 7 | 0.81 | 0.08 | 7 | 3.9% | -3.02 [-4.70 , -1.33] |
| Badawy 2017[a] | 0.34 | 0.07 | 5 | 0.59 | 0.06 | 6 | 3.5% | -3.54 [-5.73 , -1.34] |
| Bahrehbar 2020[a] | 6.61 | 1.77 | 5 | 16.77 | 4.14 | 5 | 3.6% | -2.88 [-4.92 , -0.84] |
| Bao 2018[a] | 5.41 | 2.46 | 8 | 14.44 | 4.38 | 8 | 4.1% | -2.40 [-3.77 , -1.03] |
| El-Derany 2021[a] | 0.39 | 0.08 | 8 | 0.77 | 0.04 | 8 | 3.3% | -5.68 [-8.15 , -3.21] |
| Gabr 2016[a] | 13.12 | 1.41 | 6 | 20.08 | 1.6 | 6 | 3.4% | -4.26 [-6.63 , -1.89] |
| Li 2024 | 15.29 | 2.73 | 12 | 18.26 | 3.91 | 12 | 4.4% | -0.85 [-1.69 , -0.01] |
| Lu 2024[a] | 11.86 | 4.3 | 9 | 28.79 | 7.43 | 8 | 4.1% | -2.69 [-4.10 , -1.29] |
| Nabil Salama 2024[a] | 58.45 | 3.8 | 6 | 107.6 | 1.36 | 6 | 0.9% | -15.90 [-23.74 , -8.05] |
| Park 2023[c] | 31.05 | 3.74 | 3 | 26.98 | 9.04 | 3 | 3.9% | 0.47 [-1.19 , 2.13] |
| Park 2023[a] | 14.39 | 4.88 | 3 | 23.08 | 5.26 | 3 | 3.6% | -1.37 [-3.45 , 0.71] |
| Xing 2023[c] | 14.38 | 0.24 | 6 | 19.44 | 0.33 | 6 | 0.9% | -16.19 [-24.17 , -8.21] |
| Zhang 2024[c] | 9.74 | 0.77 | 6 | 17.21 | 0.67 | 6 | 1.8% | -9.55 [-14.35 , -4.75] |
| **Subtotal (Wald[e])** | | | **107** | | | **106** | **47.0%** | **-4.32 [-5.77 , -2.86]** |
| Test for overall effect: Z = 5.82 (P < 0.00001) | | | | | | | | |
| Heterogeneity: Tau² (DL[f]) = 6.21; Chi² = 105.00, df = 14 (P < 0.00001); I² = 87% | | | | | | | | |
| **Total (Wald[e])** | | | **252** | | | **232** | **100.0%** | **-3.54 [-4.37 , -2.71]** |

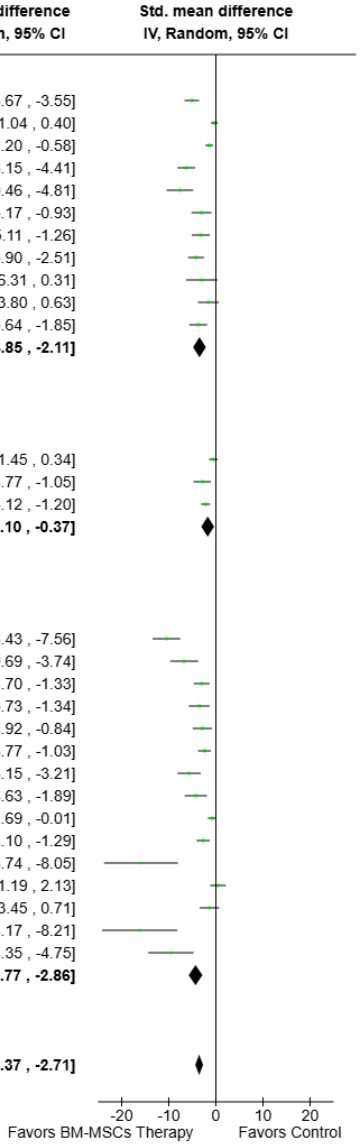


Test for overall effect: Z = 8.35 (P < 0.00001)
Test for subgroup differences: Chi² = 6.84, df = 2 (P = 0.03), I² = 70.8%
Heterogeneity: Tau² (DL[f]) = 3.91; Chi² = 205.41, df = 28 (P < 0.00001); I² = 86%

**Footnotes**
[a]BM-MSCs Transplantation
[b]BM-MSCs Transplantation and Secretome
[c]BM-MSCs Secretome
[d]BM-MSCs Secretome and miR-21 Exosomes
[e]CI calculated by Wald-type method.
[f]Tau² calculated by DerSimonian and Laird method.

**Supplemental Fig. 59. Subgroup analysis of BM-MSC Therapy on serum follicle-stimulating hormone (FSH) levels by route of administration.**

| Study or Subgroup | BM-MSCs Therapy Mean | SD | Total | Control Mean | SD | Total | Weight | Std. mean difference IV, Random, 95% CI |
|---|---|---|---|---|---|---|---|---|
| **6.6.1 Mice** | | | | | | | | |
| Badawy 2017[a] | 0.34 | 0.07 | 5 | 0.59 | 0.06 | 6 | 3.5% | -3.54 [-5.73 , -1.34] |
| Bahrehbar 2020[a] | 6.61 | 1.77 | 5 | 16.77 | 4.14 | 5 | 3.6% | -2.88 [-4.92 , -0.84] |
| Bao 2018[a] | 5.41 | 2.46 | 8 | 14.44 | 4.38 | 8 | 4.1% | -2.40 [-3.77 , -1.03] |
| Ma 2023[a] | 73.88 | 3.78 | 5 | 92.17 | 1.64 | 15 | 3.0% | -7.63 [-10.46 , -4.81] |
| Mohamed 2018[a] | 10 | 1.21 | 6 | 18.67 | 3.34 | 6 | 3.7% | -3.19 [-5.11 , -1.26] |
| Nabil Salama 2024[a] | 58.45 | 3.8 | 6 | 107.6 | 1.36 | 6 | 0.9% | -15.90 [-23.74 , -8.05] |
| Park 2023[b] | 31.05 | 3.74 | 3 | 26.98 | 9.04 | 3 | 3.9% | 0.47 [-1.19 , 2.13] |
| Park 2023[a] | 14.39 | 4.88 | 3 | 23.08 | 5.26 | 3 | 3.6% | -1.37 [-3.45 , 0.71] |
| Xie 2024[b] | 66.22 | 5.74 | 3 | 84.78 | 4.01 | 3 | 2.7% | -3.00 [-6.31 , 0.31] |
| Yang 2024[c] | 58.24 | 8.91 | 10 | 89.2 | 4.26 | 5 | 3.7% | -3.74 [-5.64 , -1.85] |
| **Subtotal (Wald[d])** | | | **54** | | | **60** | **32.6%** | **-3.27 [-4.74 , -1.81]** |
| Test for overall effect: Z = 4.37 (P < 0.0001) | | | | | | | | |
| Heterogeneity: Tau² (DL[e]) = 4.00; Chi² = 40.30, df = 9 (P < 0.00001); I² = 78% | | | | | | | | |
| **6.6.2 Rabbit** | | | | | | | | |
| Abd-Allah 2013[a] | 5.9 | 0.9 | 15 | 19.9 | 1.6 | 15 | 2.9% | -10.49 [-13.43 , -7.56] |
| **Subtotal** | | | **15** | | | **15** | **2.9%** | **-10.49 [-13.43 , -7.56]** |
| Test for overall effect: Z = 7.00 (P < 0.00001) | | | | | | | | |
| Heterogeneity: Not applicable | | | | | | | | |
| **6.6.3 Rats** | | | | | | | | |
| Afifi 2013[a] | 7.7 | 0.21 | 8 | 29.3 | 4.45 | 7 | 2.9% | -6.72 [-9.69 , -3.74] |
| Atwa 2025[a] | 0.5 | 0.11 | 7 | 0.81 | 0.08 | 7 | 3.9% | -3.02 [-4.70 , -1.33] |
| Chen 2018[a] | 15.57 | 0.97 | 15 | 22.64 | 1.64 | 15 | 4.0% | -5.11 [-6.67 , -3.55] |
| El-Derany 2021[a] | 0.39 | 0.08 | 8 | 0.77 | 0.04 | 8 | 3.3% | -5.68 [-8.15 , -3.21] |
| Elsherbiny 2024[b] | 46.41 | 7.68 | 10 | 98.36 | 126.71 | 10 | 4.4% | -0.55 [-1.45 , 0.34] |
| Fu 2008[a] | 8.7 | 18.13 | 15 | 14.7 | 18.75 | 15 | 4.5% | -0.32 [-1.04 , 0.40] |
| Fu 2017[a] | 18.04 | 3.29 | 15 | 22.96 | 3.58 | 15 | 4.4% | -1.39 [-2.20 , -0.58] |
| Gabr 2016[a] | 13.12 | 1.41 | 6 | 20.08 | 1.6 | 6 | 3.4% | -4.26 [-6.63 , -1.89] |
| Khanmohammadi 2018[f] | 7.39 | 0.76 | 20 | 12.58 | 0.89 | 10 | 3.7% | -6.28 [-8.15 , -4.41] |
| Li 2024[b] | 15.29 | 2.73 | 12 | 18.26 | 3.91 | 12 | 4.4% | -0.85 [-1.69 , -0.01] |
| Lu 2024[a] | 11.86 | 4.3 | 9 | 28.79 | 7.43 | 8 | 4.1% | -2.69 [-4.10 , -1.29] |
| Maqbool 2022[a] | 10.33 | 1.27 | 5 | 14.67 | 1.3 | 5 | 3.5% | -3.05 [-5.17 , -0.93] |
| Sameni 2019[a] | 7.77 | 0.65 | 10 | 11.19 | 0.89 | 10 | 3.9% | -4.20 [-5.90 , -2.51] |
| Tang 2024[c] | 12.32 | 0.99 | 8 | 15.18 | 0.68 | 4 | 3.7% | -2.91 [-4.77 , -1.05] |
| Xing 2023[b] | 14.38 | 0.24 | 6 | 19.44 | 0.33 | 6 | 0.9% | -16.19 [-24.17 , -8.21] |
| Yamchi 2023[a] | 2.93 | 0.13 | 3 | 3.16 | 0.1 | 3 | 3.5% | -1.59 [-3.80 , 0.63] |
| Yang 2020[f] | 14.96 | 2.15 | 20 | 19.01 | 0.76 | 10 | 4.3% | -2.16 [-3.12 , -1.20] |
| Zhang 2024[b] | 9.74 | 0.77 | 6 | 17.21 | 0.67 | 6 | 1.8% | -9.55 [-14.35 , -4.75] |
| **Subtotal (Wald[d])** | | | **183** | | | **157** | **64.4%** | **-3.30 [-4.25 , -2.34]** |
| Test for overall effect: Z = 6.78 (P < 0.00001) | | | | | | | | |
| Heterogeneity: Tau² (DL[e]) = 3.19; Chi² = 129.59, df = 17 (P < 0.00001); I² = 87% | | | | | | | | |
| **Total (Wald[d])** | | | **252** | | | **232** | **100.0%** | **-3.54 [-4.37 , -2.71]** |

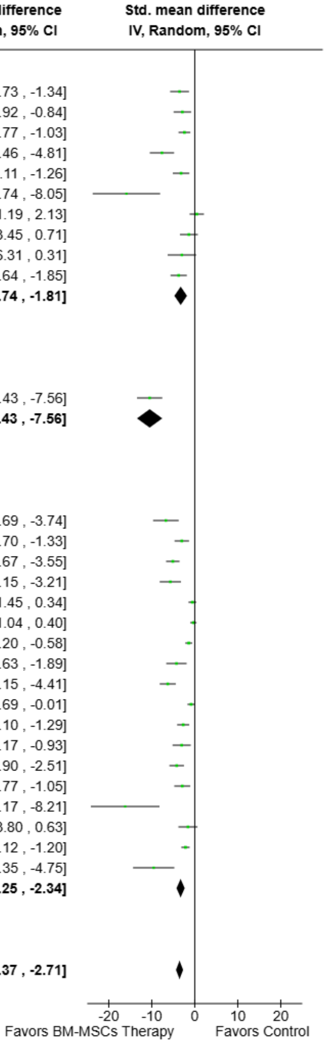


Test for overall effect: Z = 8.35 (P < 0.00001)
Test for subgroup differences: Chi² = 21.51, df = 2 (P < 0.0001), I² = 90.7%
Heterogeneity: Tau² (DL[e]) = 3.91; Chi² = 205.41, df = 28 (P < 0.00001); I² = 86%

**Footnotes**
[a]BM-MSCs Transplantation
[b]BM-MSCs Secretome
[c]BM-MSCs Secretome and mIR-21 Exosomes
[d]CI calculated by Wald-type method.
[e]Tau² calculated by DerSimonian and Laird method.
[f]BM-MSCs Transplantation and Secretome

**Supplemental Fig. 60. Subgroup analysis of BM-MSC Therapy on serum follicle-stimulating hormone (FSH) levels by animal model.**

| Study or Subgroup | BM-MSCs Therapy Mean | SD | Total | Control Mean | SD | Total | Weight | Std. mean difference IV, Random, 95% CI |
|---|---|---|---|---|---|---|---|---|
| **6.7.1 Autoimmune** | | | | | | | | |
| Ma 2023[a] | 73.88 | 3.78 | 5 | 92.17 | 1.64 | 15 | 3.0% | -7.63 [-10.46 , -4.81] |
| Xie 2024[b] | 66.22 | 5.74 | 3 | 84.78 | 4.01 | 3 | 2.7% | -3.00 [-6.31 , 0.31] |
| Yang 2024[c] | 58.24 | 8.91 | 10 | 89.2 | 4.26 | 5 | 3.7% | -3.74 [-5.64 , -1.85] |
| **Subtotal (Wald[d])** | | | **18** | | | **23** | **9.4%** | **-4.78 [-7.44 , -2.12]** |
| Test for overall effect: Z = 3.52 (P = 0.0004) | | | | | | | | |
| Heterogeneity: Tau² (DL[e]) = 3.69; Chi² = 6.10, df = 2 (P = 0.05); I² = 67% | | | | | | | | |
| **6.7.2 Chemotherapy (Cisplatin)** | | | | | | | | |
| Nabil Salama 2024[a] | 58.45 | 3.8 | 6 | 107.6 | 1.36 | 6 | 0.9% | -15.90 [-23.74 , -8.05] |
| Tang 2024[c] | 12.32 | 0.99 | 8 | 15.18 | 0.68 | 4 | 3.7% | -2.91 [-4.77 , -1.05] |
| **Subtotal (Wald[d])** | | | **14** | | | **10** | **4.6%** | **-8.82 [-21.50 , 3.86]** |
| Test for overall effect: Z = 1.36 (P = 0.17) | | | | | | | | |
| Heterogeneity: Tau² (DL[e]) = 75.92; Chi² = 9.98, df = 1 (P = 0.002); I² = 90% | | | | | | | | |
| **6.7.3 Chemotherapy (Cyclophosphamide)** | | | | | | | | |
| Abd-Allah 2013[a] | 5.9 | 0.9 | 15 | 19.9 | 1.6 | 15 | 2.9% | -10.49 [-13.43 , -7.56] |
| Afifi 2013[a] | 7.7 | 0.21 | 8 | 29.3 | 4.45 | 7 | 2.9% | -6.72 [-9.69 , -3.74] |
| Atwa 2025[a] | 0.5 | 0.11 | 7 | 0.81 | 0.08 | 7 | 3.9% | -3.02 [-4.70 , -1.33] |
| Badawy 2017[a] | 0.34 | 0.07 | 5 | 0.59 | 0.06 | 6 | 3.5% | -3.54 [-5.73 , -1.34] |
| Bahrehbar 2020[a] | 6.61 | 1.77 | 5 | 16.77 | 4.14 | 5 | 3.6% | -2.88 [-4.92 , -0.84] |
| Bao 2018[a] | 5.41 | 2.46 | 8 | 14.44 | 4.38 | 8 | 4.1% | -2.40 [-3.77 , -1.03] |
| Chen 2018[a] | 15.57 | 0.97 | 15 | 22.64 | 1.64 | 15 | 4.0% | -5.11 [-6.67 , -3.55] |
| Elsherbiny 2024[b] | 46.41 | 7.68 | 10 | 98.36 | 126.71 | 10 | 4.4% | -0.55 [-1.45 , 0.34] |
| Fu 2008[a] | 8.7 | 18.13 | 15 | 14.7 | 18.75 | 15 | 4.5% | -0.32 [-1.04 , 0.40] |
| Fu 2017[a] | 18.04 | 3.29 | 15 | 22.96 | 3.58 | 15 | 4.4% | -1.39 [-2.20 , -0.58] |
| Gabr 2016[a] | 13.12 | 1.41 | 6 | 20.08 | 1.6 | 6 | 3.4% | -4.26 [-6.63 , -1.89] |
| Khanmohammadi 2018[f] | 7.39 | 0.76 | 20 | 12.58 | 0.89 | 10 | 3.7% | -6.28 [-8.15 , -4.41] |
| Li 2024[b] | 15.29 | 2.73 | 12 | 18.26 | 3.91 | 12 | 4.4% | -0.85 [-1.69 , -0.01] |
| Lu 2024[a] | 11.86 | 4.3 | 9 | 28.79 | 7.43 | 8 | 4.1% | -2.69 [-4.10 , -1.29] |
| Maqbool 2022[a] | 10.33 | 1.27 | 5 | 14.67 | 1.3 | 5 | 3.5% | -3.05 [-5.17 , -0.93] |
| Mohamed 2018[a] | 10 | 1.21 | 6 | 18.67 | 3.34 | 6 | 3.7% | -3.19 [-5.11 , -1.26] |
| Park 2023[b] | 31.05 | 3.74 | 3 | 26.98 | 9.04 | 3 | 3.9% | 0.47 [-1.19 , 2.13] |
| Park 2023[a] | 14.39 | 4.88 | 3 | 23.08 | 5.26 | 3 | 3.6% | -1.37 [-3.45 , 0.71] |
| Sameni 2019[a] | 7.77 | 0.65 | 10 | 11.19 | 0.89 | 10 | 3.9% | -4.20 [-5.90 , -2.51] |
| Xing 2023[b] | 14.38 | 0.24 | 6 | 19.44 | 0.33 | 6 | 0.9% | -16.19 [-24.17 , -8.21] |
| Yamchi 2023[a] | 2.93 | 0.13 | 3 | 3.16 | 0.1 | 3 | 3.5% | -1.59 [-3.80 , 0.63] |
| Yang 2020[f] | 14.96 | 2.15 | 20 | 19.01 | 0.76 | 10 | 4.3% | -2.16 [-3.12 , -1.20] |
| Zhang 2024[b] | 9.74 | 0.77 | 6 | 17.21 | 0.67 | 6 | 1.8% | -9.55 [-14.35 , -4.75] |
| **Subtotal (Wald[d])** | | | **212** | | | **191** | **82.7%** | **-3.17 [-4.04 , -2.30]** |
| Test for overall effect: Z = 7.14 (P < 0.00001) | | | | | | | | |
| Heterogeneity: Tau² (DL[e]) = 3.46; Chi² = 165.61, df = 22 (P < 0.00001); I² = 87% | | | | | | | | |
| **6.7.4 Radiotherapy** | | | | | | | | |
| El-Derany 2021[a] | 0.39 | 0.08 | 8 | 0.77 | 0.04 | 8 | 3.3% | -5.68 [-8.15 , -3.21] |
| **Subtotal** | | | **8** | | | **8** | **3.3%** | **-5.68 [-8.15 , -3.21]** |
| Test for overall effect: Z = 4.51 (P < 0.00001) | | | | | | | | |
| Heterogeneity: Not applicable | | | | | | | | |
| **Total (Wald[d])** | | | **252** | | | **232** | **100.0%** | **-3.54 [-4.37 , -2.71]** |

Test for overall effect: Z = 8.35 (P < 0.00001)
Test for subgroup differences: Chi² = 5.05, df = 3 (P = 0.17), I² = 40.6%
Heterogeneity: Tau² (DL[e]) = 3.91; Chi² = 205.41, df = 28 (P < 0.00001); I² = 86%

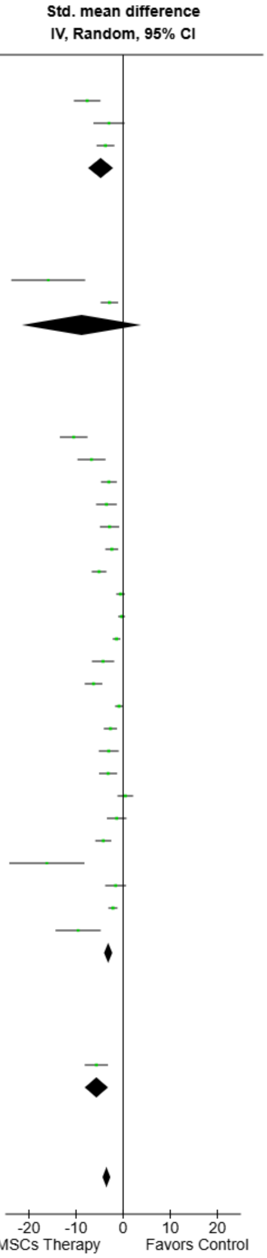


**Footnotes**
[a]BM-MSCs Transplantation
[b]BM-MSCs Secretome
[c]BM-MSCs Secretome and miR-21 Exosomes
[d]CI calculated by Wald-type method.
[e]Tau² calculated by DerSimonian and Laird method.
[f]BM-MSCs Transplantation and Secretome

**Supplemental Fig. 61. Subgroup analysis of BM-MSC Therapy on serum follicle-stimulating hormone (FSH) levels by POI induction model.**

| Study or Subgroup | BM-MSCs Therapy Mean | SD | Total | Control Mean | SD | Total | Weight | Std. mean difference IV, Random, 95% CI |
|---|---|---|---|---|---|---|---|---|
| **6.8.1 Intraovarian** | | | | | | | | |
| Maqbool 2022[a] | 29 | 8.07 | 5 | 9.67 | 1.3 | 5 | 6.3% | 3.02 [0.92 , 5.13] |
| Mohamed 2018[a] | 36.24 | 16.02 | 6 | 8.42 | 18.74 | 6 | 8.2% | 1.47 [0.13 , 2.81] |
| Xie 2024[b] | 538.25 | 25.81 | 3 | 440.55 | 40.56 | 3 | 4.9% | 2.30 [-0.44 , 5.04] |
| Yang 2024[c] | 501.32 | 72.45 | 10 | 380.95 | 76.72 | 5 | 8.5% | 1.54 [0.29 , 2.79] |
| **Subtotal (Wald[d])** | | | **24** | | | **19** | **27.9%** | **1.79 [0.99 , 2.60]** |
| Test for overall effect: Z = 4.39 (P < 0.0001) | | | | | | | | |
| Heterogeneity: Tau$^2$ (DL[e]) = 0.00; Chi$^2$ = 1.82, df = 3 (P = 0.61); I$^2$ = 0% | | | | | | | | |
| **6.8.2 Intraperitoneal** | | | | | | | | |
| Besikcioglu 2019[a] | 0.77 | 0.27 | 12 | 0.82 | 0.2 | 12 | 9.5% | -0.20 [-1.01 , 0.60] |
| Elsherbiny 2024[b] | 8.05 | 3.23 | 10 | 2.71 | 2.02 | 10 | 8.9% | 1.90 [0.80 , 2.99] |
| Tang 2024[c] | 2481 | 319.9 | 8 | 1578.38 | 214 | 4 | 6.9% | 2.85 [1.01 , 4.69] |
| Yang 2020[f] | 148 | 12.9 | 20 | 131 | 11.86 | 10 | 9.5% | 1.32 [0.48 , 2.15] |
| **Subtotal (Wald[d])** | | | **50** | | | **36** | **34.8%** | **1.32 [0.12 , 2.52]** |
| Test for overall effect: Z = 2.15 (P = 0.03) | | | | | | | | |
| Heterogeneity: Tau$^2$ (DL[e]) = 1.16; Chi$^2$ = 15.65, df = 3 (P = 0.001); I$^2$ = 81% | | | | | | | | |
| **6.8.3 Intravenous** | | | | | | | | |
| Li 2024[b] | 143.99 | 6.28 | 12 | 130.09 | 12.82 | 12 | 9.3% | 1.33 [0.43 , 2.23] |
| Lu 2024[a] | 378.88 | 49.6 | 9 | 276.4 | 74.53 | 8 | 8.8% | 1.56 [0.43 , 2.68] |
| Nabil Salama 2024[a] | 8.52 | 0.78 | 6 | 1.93 | 0.18 | 6 | 2.0% | 10.75 [5.38 , 16.11] |
| Park 2023[a] | 4.78 | 2.19 | 3 | 3.29 | 0.08 | 3 | 7.1% | 0.77 [-1.00 , 2.53] |
| Park 2023[b] | 3.24 | 0.01 | 3 | 4.01 | 1.53 | 3 | 7.3% | -0.57 [-2.26 , 1.12] |
| Xing 2023[b] | 146.2 | 0.7 | 6 | 129.06 | 0.88 | 6 | 0.7% | 19.90 [10.12 , 29.68] |
| Zhang 2024[b] | 151.03 | 2.07 | 6 | 131.38 | 1.38 | 6 | 2.1% | 10.31 [5.15 , 15.47] |
| **Subtotal (Wald[d])** | | | **45** | | | **44** | **37.3%** | **3.21 [1.18 , 5.24]** |
| Test for overall effect: Z = 3.09 (P = 0.002) | | | | | | | | |
| Heterogeneity: Tau$^2$ (DL[e]) = 4.96; Chi$^2$ = 42.65, df = 6 (P < 0.00001); I$^2$ = 86% | | | | | | | | |
| **Total (Wald[d])** | | | **119** | | | **99** | **100.0%** | **1.86 [1.03 , 2.69]** |

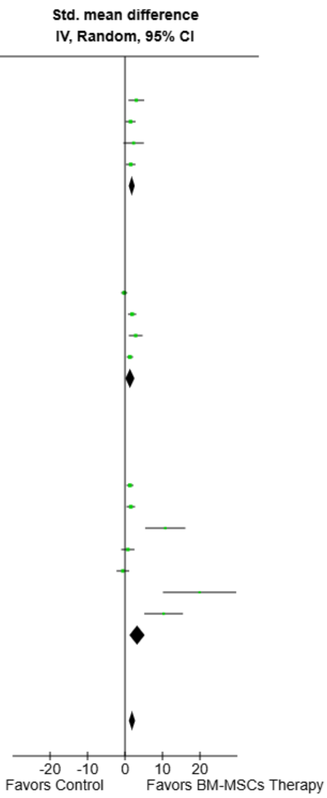


Test for overall effect: Z = 4.38 (P < 0.0001)
Test for subgroup differences: Chi$^2$ = 2.47, df = 2 (P = 0.29), I$^2$ = 18.9%
Heterogeneity: Tau$^2$ (DL[e]) = 1.73; Chi$^2$ = 63.35, df = 14 (P < 0.00001); I$^2$ = 78%

**Footnotes**

[a]BM-MSCs Transplantation
[b]BM-MSCs Secretome
[c]BM-MSCs Secretome and mIR-21 Exosomes
[d]CI calculated by Wald-type method.
[e]Tau$^2$ calculated by DerSimonian and Laird method.
[f]BM-MSCs Transplantation and Secretome

**Supplemental Fig. 62. Subgroup analysis of BM-MSC Therapy on serum anti-müllerian hormone (AMH) levels by route of administration.**

| Study or Subgroup | BM-MSCs Therapy Mean | SD | Total | Control Mean | SD | Total | Weight | Std. mean difference IV, Random, 95% CI |
|---|---|---|---|---|---|---|---|---|
| **6.9.1 Mice** | | | | | | | | |
| Mohamed 2018[a] | 36.24 | 16.02 | 6 | 8.42 | 18.74 | 6 | 8.2% | 1.47 [0.13 , 2.81] |
| Nabil Salama 2024[a] | 8.52 | 0.78 | 6 | 1.93 | 0.18 | 6 | 2.0% | 10.75 [5.38 , 16.11] |
| Park 2023[b] | 3.24 | 0.01 | 3 | 4.01 | 1.53 | 3 | 7.3% | -0.57 [-2.26 , 1.12] |
| Park 2023[a] | 4.78 | 2.19 | 3 | 3.29 | 0.08 | 3 | 7.1% | 0.77 [-1.00 , 2.53] |
| Xie 2024[b] | 538.25 | 25.81 | 3 | 440.55 | 40.56 | 3 | 4.9% | 2.30 [-0.44 , 5.04] |
| Yang 2024[c] | 501.32 | 72.45 | 10 | 380.95 | 76.72 | 5 | 8.5% | 1.54 [0.29 , 2.79] |
| **Subtotal (Wald[d])** | | | **31** | | | **26** | **38.0%** | **1.60 [0.15 , 3.05]** |
| Test for overall effect: Z = 2.17 (P = 0.03) | | | | | | | | |
| Heterogeneity: Tau² (DL[e]) = 2.10; Chi² = 17.63, df = 5 (P = 0.003); I² = 72% | | | | | | | | |
| **6.9.2 Rats** | | | | | | | | |
| Besikcioglu 2019[a] | 0.77 | 0.27 | 12 | 0.82 | 0.2 | 12 | 9.5% | -0.20 [-1.01 , 0.60] |
| Elsherbiny 2024[b] | 8.05 | 3.23 | 10 | 2.71 | 2.02 | 10 | 8.9% | 1.90 [0.80 , 2.99] |
| Li 2024[b] | 143.99 | 6.28 | 12 | 130.09 | 12.82 | 12 | 9.3% | 1.33 [0.43 , 2.23] |
| Lu 2024[a] | 378.88 | 49.6 | 9 | 276.4 | 74.53 | 8 | 8.8% | 1.56 [0.43 , 2.68] |
| Maqbool 2022[a] | 29 | 8.07 | 5 | 9.67 | 1.3 | 5 | 6.3% | 3.02 [0.92 , 5.13] |
| Tang 2024[c] | 2481 | 319.9 | 8 | 1578.38 | 214 | 4 | 6.9% | 2.85 [1.01 , 4.69] |
| Xing 2023[b] | 146.2 | 0.7 | 6 | 129.06 | 0.88 | 6 | 0.7% | 19.90 [10.12 , 29.68] |
| Yang 2020[f] | 148 | 12.9 | 20 | 131 | 11.86 | 10 | 9.5% | 1.32 [0.48 , 2.15] |
| Zhang 2024[b] | 151.03 | 2.07 | 6 | 131.38 | 1.38 | 6 | 2.1% | 10.31 [5.15 , 15.47] |
| **Subtotal (Wald[d])** | | | **88** | | | **73** | **62.0%** | **2.09 [0.99 , 3.18]** |
| Test for overall effect: Z = 3.74 (P = 0.0002) | | | | | | | | |
| Heterogeneity: Tau² (DL[e]) = 1.87; Chi² = 45.72, df = 8 (P < 0.00001); I² = 83% | | | | | | | | |
| **Total (Wald[d])** | | | **119** | | | **99** | **100.0%** | **1.86 [1.03 , 2.69]** |

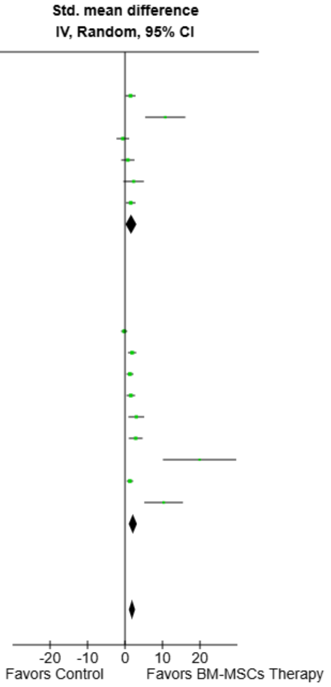


Test for overall effect: Z = 4.38 (P < 0.0001)
Test for subgroup differences: Chi² = 0.27, df = 1 (P = 0.60), I² = 0%
Heterogeneity: Tau² (DL[e]) = 1.73; Chi² = 63.35, df = 14 (P < 0.00001); I² = 78%

**Footnotes**
[a]BM-MSCs Transplantation
[b]BM-MSCs Secretome
[c]BM-MSCs Secretome and mIR-21 Exosomes
[d]CI calculated by Wald-type method.
[e]Tau² calculated by DerSimonian and Laird method.
[f]BM-MSCs Transplantation and Secretome

**Supplemental Fig. 63. Subgroup analysis of BM-MSC Therapy on serum anti-müllerian hormone (AMH) levels by animal model.**

| Study or Subgroup | BM-MSCs Therapy Mean | SD | Total | Control Mean | SD | Total | Weight | Std. mean difference IV, Random, 95% CI |
|---|---|---|---|---|---|---|---|---|
| **6.10.1 Autoimmune** | | | | | | | | |
| Xie 2024[a] | 538.25 | 25.81 | 3 | 440.55 | 40.56 | 3 | 4.9% | 2.30 [-0.44 , 5.04] |
| Yang 2024[b] | 501.32 | 72.45 | 10 | 380.95 | 76.72 | 5 | 8.5% | 1.54 [0.29 , 2.79] |
| **Subtotal (Wald[c])** | | | **13** | | | **8** | **13.4%** | **1.67 [0.53 , 2.80]** |
| Test for overall effect: Z = 2.87 (P = 0.004) | | | | | | | | |
| Heterogeneity: Tau² (DL[d]) = 0.00; Chi² = 0.25, df = 1 (P = 0.62); I² = 0% | | | | | | | | |
| **6.10.2 Chemotherapy (Cisplatin)** | | | | | | | | |
| Nabil Salama 2024[e] | 8.52 | 0.78 | 6 | 1.93 | 0.18 | 6 | 2.0% | 10.75 [5.38 , 16.11] |
| Tang 2024[b] | 2481 | 319.9 | 8 | 1578.38 | 214 | 4 | 6.9% | 2.85 [1.01 , 4.69] |
| **Subtotal (Wald[c])** | | | **14** | | | **10** | **8.9%** | **6.38 [-1.31 , 14.07]** |
| Test for overall effect: Z = 1.63 (P = 0.10) | | | | | | | | |
| Heterogeneity: Tau² (DL[d]) = 26.98; Chi² = 7.44, df = 1 (P = 0.006); I² = 87% | | | | | | | | |
| **6.10.3 Chemotherapy (Cyclophosphamide)** | | | | | | | | |
| Besikcioglu 2019[e] | 0.77 | 0.27 | 12 | 0.82 | 0.2 | 12 | 9.5% | -0.20 [-1.01 , 0.60] |
| Elsherbiny 2024[a] | 8.05 | 3.23 | 10 | 2.71 | 2.02 | 10 | 8.9% | 1.90 [0.80 , 2.99] |
| Li 2024[a] | 143.99 | 6.28 | 12 | 130.09 | 12.82 | 12 | 9.3% | 1.33 [0.43 , 2.23] |
| Lu 2024[e] | 378.88 | 49.6 | 9 | 276.4 | 74.53 | 8 | 8.8% | 1.56 [0.43 , 2.68] |
| Maqbool 2022[e] | 29 | 8.07 | 5 | 9.67 | 1.3 | 5 | 6.3% | 3.02 [0.92 , 5.13] |
| Mohamed 2018[e] | 36.24 | 16.02 | 6 | 8.42 | 18.74 | 6 | 8.2% | 1.47 [0.13 , 2.81] |
| Park 2023[a] | 3.24 | 0.01 | 3 | 4.01 | 1.53 | 3 | 7.3% | -0.57 [-2.26 , 1.12] |
| Park 2023[e] | 4.78 | 2.19 | 3 | 3.29 | 0.08 | 3 | 7.1% | 0.77 [-1.00 , 2.53] |
| Xing 2023[a] | 146.2 | 0.7 | 6 | 129.06 | 0.88 | 6 | 0.7% | 19.90 [10.12 , 29.68] |
| Yang 2020[f] | 148 | 12.9 | 20 | 131 | 11.86 | 10 | 9.5% | 1.32 [0.48 , 2.15] |
| Zhang 2024[a] | 151.03 | 2.07 | 6 | 131.38 | 1.38 | 6 | 2.1% | 10.31 [5.15 , 15.47] |
| **Subtotal (Wald[c])** | | | **92** | | | **81** | **77.7%** | **1.52 [0.62 , 2.43]** |
| Test for overall effect: Z = 3.30 (P = 0.0010) | | | | | | | | |
| Heterogeneity: Tau² (DL[d]) = 1.54; Chi² = 47.11, df = 10 (P < 0.00001); I² = 79% | | | | | | | | |
| **Total (Wald[c])** | | | **119** | | | **99** | **100.0%** | **1.86 [1.03 , 2.69]** |

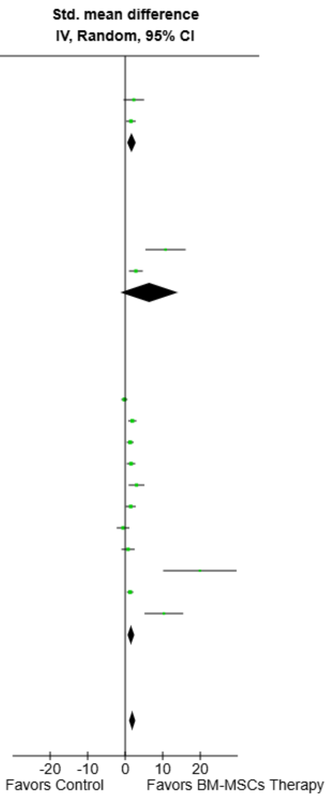


Test for overall effect: Z = 4.38 (P < 0.0001)
Test for subgroup differences: Chi² = 1.52, df = 2 (P = 0.47), I² = 0%
Heterogeneity: Tau² (DL[d]) = 1.73; Chi² = 63.35, df = 14 (P < 0.00001); I² = 78%

**Footnotes**
[a]BM-MSCs Secretome
[b]BM-MSCs Secretome and miR-21 Exosomes
[c]CI calculated by Wald-type method.
[d]Tau² calculated by DerSimonian and Laird method.
[e]BM-MSCs Transplantation
[f]BM-MSCs Transplantation and Secretome

**Supplemental Fig. 64. Subgroup analysis of BM-MSC Therapy on serum anti-müllerian hormone (AMH) levels by POI induction model.**

| Study or Subgroup | BM-MSCs Therapy Mean | SD | Total | Control Mean | SD | Total | Weight | Std. mean difference IV, Random, 95% CI |
|---|---|---|---|---|---|---|---|---|
| **6.11.1 Intraovarian** | | | | | | | | |
| Chen 2018[a] | 383.54 | 5.7 | 5 | 360.76 | 9.49 | 5 | 4.6% | 2.63 [0.70 , 4.56] |
| Fu 2008[a] | 383.57 | 32.27 | 5 | 364.33 | 34.95 | 5 | 6.7% | 0.52 [-0.76 , 1.79] |
| Fu 2017[a] | 357.32 | 19.51 | 5 | 357.32 | 12 | 5 | 6.8% | 0.00 [-1.24 , 1.24] |
| Khanmohammadi 2018[b] | 36.42 | 8.38 | 20 | 21.5 | 4.33 | 10 | 8.0% | 1.98 [1.05 , 2.91] |
| Ma 2023[a] | 10.04 | 1.81 | 5 | 1.52 | 0.26 | 5 | 1.9% | 5.95 [2.38 , 9.52] |
| Park 2021b[a] | 4.34 | 2.13 | 3 | 1.54 | 1.21 | 3 | 4.3% | 1.29 [-0.74 , 3.32] |
| Sameni 2019[a] | 32.94 | 4.92 | 10 | 23.83 | 3.07 | 10 | 7.1% | 2.13 [0.98 , 3.27] |
| Xie 2024[c] | 4.74 | 0.56 | 5 | 3.28 | 0.3 | 5 | 4.2% | 2.94 [0.87 , 5.00] |
| Yang 2024[d] | 5.26 | 1.43 | 10 | 3.05 | 1.03 | 5 | 6.7% | 1.58 [0.32 , 2.83] |
| **Subtotal (Wald[e])** | | | **68** | | | **53** | **50.3%** | **1.72 [0.95 , 2.48]** |
| Test for overall effect: Z = 4.37 (P < 0.0001) | | | | | | | | |
| Heterogeneity: Tau² (DL[f]) = 0.74; Chi² = 19.13, df = 8 (P = 0.01); I² = 58% | | | | | | | | |
| **6.11.2 Intraperitoneal** | | | | | | | | |
| Besikcioglu 2019[a] | 26.18 | 5.38 | 12 | 19.91 | 5.24 | 12 | 8.2% | 1.14 [0.27 , 2.01] |
| Tang 2024[d] | 210.82 | 38.57 | 8 | 104.3 | 12.4 | 4 | 4.7% | 2.98 [1.10 , 4.87] |
| Yang 2020[b] | 15.17 | 1.9 | 20 | 6.03 | 3.16 | 10 | 6.7% | 3.74 [2.47 , 5.01] |
| **Subtotal (Wald[e])** | | | **40** | | | **26** | **19.6%** | **2.55 [0.73 , 4.37]** |
| Test for overall effect: Z = 2.75 (P = 0.006) | | | | | | | | |
| Heterogeneity: Tau² (DL[f]) = 2.10; Chi² = 11.89, df = 2 (P = 0.003); I² = 83% | | | | | | | | |
| **6.11.3 Intravenous** | | | | | | | | |
| Afifi 2013[a] | 30.2 | 10.4 | 8 | 10.3 | 2.6 | 7 | 6.1% | 2.39 [0.97 , 3.81] |
| Bahrehbar 2020[a] | 77.95 | 15.25 | 5 | 59.01 | 10.3 | 5 | 6.1% | 1.31 [-0.13 , 2.76] |
| El-Derany 2021[a] | 10.24 | 1.99 | 5 | 4.84 | 1.74 | 5 | 4.6% | 2.61 [0.69 , 4.53] |
| Lu 2024[a] | 65.02 | 11.61 | 6 | 26.01 | 13.93 | 6 | 5.0% | 2.81 [1.03 , 4.59] |
| Xing 2023[c] | 14.23 | 1.29 | 6 | 5.98 | 1.55 | 6 | 2.8% | 5.34 [2.50 , 8.18] |
| Zhang 2024[c] | 5.97 | 0.89 | 6 | 2.98 | 1.48 | 6 | 5.6% | 2.26 [0.68 , 3.84] |
| **Subtotal (Wald[e])** | | | **36** | | | **35** | **30.1%** | **2.44 [1.64 , 3.25]** |
| Test for overall effect: Z = 5.94 (P < 0.00001) | | | | | | | | |
| Heterogeneity: Tau² (DL[f]) = 0.24; Chi² = 6.56, df = 5 (P = 0.26); I² = 24% | | | | | | | | |
| **Total (Wald[e])** | | | **144** | | | **114** | **100.0%** | **2.11 [1.56 , 2.66]** |

Test for overall effect: Z = 7.47 (P < 0.00001)
Test for subgroup differences: Chi² = 1.90, df = 2 (P = 0.39), I² = 0%
Heterogeneity: Tau² (DL[f]) = 0.77; Chi² = 41.48, df = 17 (P = 0.0008); I² = 59%

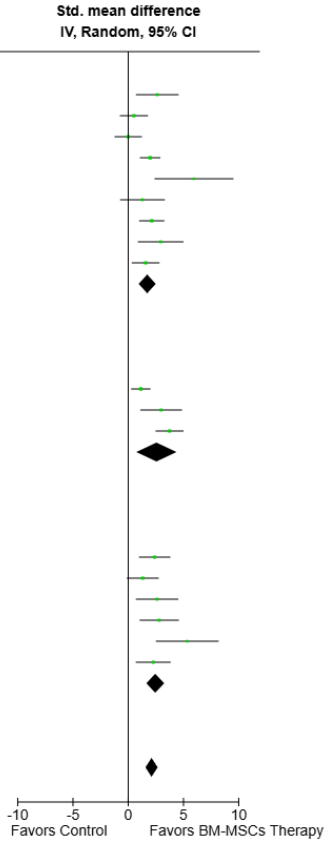


**Footnotes**
[a]BM-MSCs Transplantation
[b]BM-MSCs Transplantation and Secretome
[c]BM-MSCs Secretome
[d]BM-MSCs Secretome and mIR-21 Exosomes
[e]CI calculated by Wald-type method.
[f]Tau² calculated by DerSimonian and Laird method.

**Supplemental Fig. 65. Subgroup analysis of BM-MSC Therapy on primordial follicle count by route of administration.**

| Study or Subgroup | BM-MSCs Therapy Mean | SD | Total | Control Mean | SD | Total | Weight | Std. mean difference IV, Random, 95% CI |
|---|---|---|---|---|---|---|---|---|
| **6.13.1 Autoimmune** | | | | | | | | |
| Ma 2023[a] | 10.04 | 1.81 | 5 | 1.52 | 0.26 | 5 | 1.9% | 5.95 [2.38 , 9.52] |
| Xie 2024[b] | 4.74 | 0.56 | 5 | 3.28 | 0.3 | 5 | 4.2% | 2.94 [0.87 , 5.00] |
| Yang 2024[c] | 5.26 | 1.43 | 10 | 3.05 | 1.03 | 5 | 6.7% | 1.58 [0.32 , 2.83] |
| **Subtotal (Wald[d])** | | | **20** | | | **15** | **12.9%** | **2.95 [0.88 , 5.01]** |
| Test for overall effect: Z = 2.80 (P = 0.005) | | | | | | | | |
| Heterogeneity: Tau² (DL[e]) = 2.08; Chi² = 5.65, df = 2 (P = 0.06); I² = 65% | | | | | | | | |
| **6.13.2 Chemotherapy (Cisplatin)** | | | | | | | | |
| Tang 2024[c] | 210.82 | 38.57 | 8 | 104.3 | 12.4 | 4 | 4.7% | 2.98 [1.10 , 4.87] |
| **Subtotal** | | | **8** | | | **4** | **4.7%** | **2.98 [1.10 , 4.87]** |
| Test for overall effect: Z = 3.10 (P = 0.002) | | | | | | | | |
| Heterogeneity: Not applicable | | | | | | | | |
| **6.13.3 Chemotherapy (Cyclophosphamide)** | | | | | | | | |
| Afifi 2013[a] | 30.2 | 10.4 | 8 | 10.3 | 2.6 | 7 | 6.1% | 2.39 [0.97 , 3.81] |
| Bahrehbar 2020[a] | 77.95 | 15.25 | 5 | 59.01 | 10.3 | 5 | 6.1% | 1.31 [-0.13 , 2.76] |
| Besikcioglu 2019[a] | 26.18 | 5.38 | 12 | 19.91 | 5.24 | 12 | 8.2% | 1.14 [0.27 , 2.01] |
| Chen 2018[a] | 383.54 | 5.7 | 5 | 360.76 | 9.49 | 5 | 4.6% | 2.63 [0.70 , 4.56] |
| Fu 2008[a] | 383.57 | 32.27 | 5 | 364.33 | 34.95 | 5 | 6.7% | 0.52 [-0.76 , 1.79] |
| Fu 2017[a] | 357.32 | 19.51 | 5 | 357.32 | 12 | 5 | 6.8% | 0.00 [-1.24 , 1.24] |
| Khanmohammadi 2018[f] | 36.42 | 8.38 | 20 | 21.5 | 4.33 | 10 | 8.0% | 1.98 [1.05 , 2.91] |
| Lu 2024[a] | 65.02 | 11.61 | 6 | 26.01 | 13.93 | 6 | 5.0% | 2.81 [1.03 , 4.59] |
| Park 2021b[a] | 4.34 | 2.13 | 3 | 1.54 | 1.21 | 3 | 4.3% | 1.29 [-0.74 , 3.32] |
| Sameni 2019[a] | 32.94 | 4.92 | 10 | 23.83 | 3.07 | 10 | 7.1% | 2.13 [0.98 , 3.27] |
| Xing 2023[b] | 14.23 | 1.29 | 6 | 5.98 | 1.55 | 6 | 2.8% | 5.34 [2.50 , 8.18] |
| Yang 2020[f] | 15.17 | 1.9 | 20 | 6.03 | 3.16 | 10 | 6.7% | 3.74 [2.47 , 5.01] |
| Zhang 2024[b] | 5.97 | 0.89 | 6 | 2.98 | 1.48 | 6 | 5.6% | 2.26 [0.68 , 3.84] |
| **Subtotal (Wald[d])** | | | **111** | | | **90** | **77.8%** | **1.94 [1.30 , 2.58]** |
| Test for overall effect: Z = 5.93 (P < 0.00001) | | | | | | | | |
| Heterogeneity: Tau² (DL[e]) = 0.83; Chi² = 33.20, df = 12 (P = 0.0009); I² = 64% | | | | | | | | |
| **6.13.4 Radiotherapy** | | | | | | | | |
| El-Derany 2021[a] | 10.24 | 1.99 | 5 | 4.84 | 1.74 | 5 | 4.6% | 2.61 [0.69 , 4.53] |
| **Subtotal** | | | **5** | | | **5** | **4.6%** | **2.61 [0.69 , 4.53]** |
| Test for overall effect: Z = 2.66 (P = 0.008) | | | | | | | | |
| Heterogeneity: Not applicable | | | | | | | | |
| **Total (Wald[d])** | | | **144** | | | **114** | **100.0%** | **2.11 [1.56 , 2.66]** |

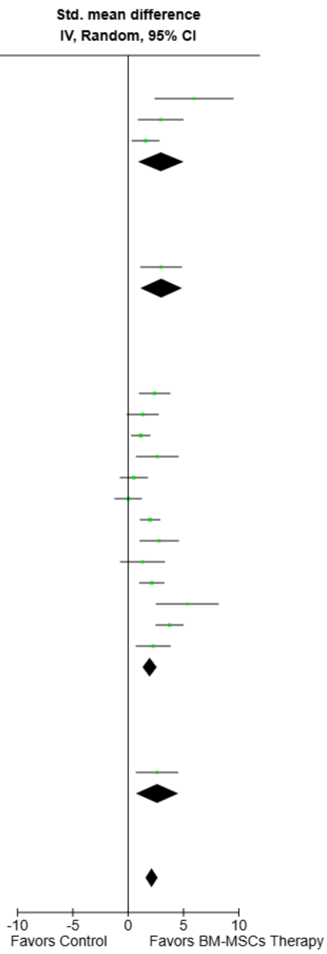


Test for overall effect: Z = 7.47 (P < 0.00001)
Test for subgroup differences: Chi² = 1.97, df = 3 (P = 0.58), I² = 0%
Heterogeneity: Tau² (DL[e]) = 0.77; Chi² = 41.48, df = 17 (P = 0.0008); I² = 59%

**Footnotes**
[a]BM-MSCs Transplantation
[b]BM-MSCs Secretome
[c]BM-MSCs Secretome and mIR-21 Exosomes
[d]CI calculated by Wald-type method.
[e]Tau² calculated by DerSimonian and Laird method.
[f]BM-MSCs Transplantation and Secretome

**Supplemental Fig. 66. Subgroup analysis of BM-MSC Therapy on primordial follicle count by animal model.**

| Study or Subgroup | BM-MSCs Therapy Mean | SD | Total | Control Mean | SD | Total | Weight | Std. mean difference IV, Random, 95% CI |
|---|---|---|---|---|---|---|---|---|
| **6.12.1 Mice** | | | | | | | | |
| Bahrehbar 2020[a] | 77.95 | 15.25 | 5 | 59.01 | 10.3 | 5 | 6.1% | 1.31 [-0.13 , 2.76] |
| Ma 2023[a] | 10.04 | 1.81 | 5 | 1.52 | 0.26 | 5 | 1.9% | 5.95 [2.38 , 9.52] |
| Park 2021b[a] | 4.34 | 2.13 | 3 | 1.54 | 1.21 | 3 | 4.3% | 1.29 [-0.74 , 3.32] |
| Xie 2024[b] | 4.74 | 0.56 | 5 | 3.28 | 0.3 | 5 | 4.2% | 2.94 [0.87 , 5.00] |
| Yang 2024[c] | 5.26 | 1.43 | 10 | 3.05 | 1.03 | 5 | 6.7% | 1.58 [0.32 , 2.83] |
| **Subtotal (Wald[d])** | | | **28** | | | **23** | **23.3%** | **2.05 [0.94 , 3.15]** |
| Test for overall effect: Z = 3.63 (P = 0.0003) | | | | | | | | |
| Heterogeneity: Tau² (DL[e]) = 0.67; Chi² = 7.12, df = 4 (P = 0.13); I² = 44% | | | | | | | | |
| **6.12.2 Rats** | | | | | | | | |
| Afifi 2013[a] | 30.2 | 10.4 | 8 | 10.3 | 2.6 | 7 | 6.1% | 2.39 [0.97 , 3.81] |
| Besikcioglu 2019[a] | 26.18 | 5.38 | 12 | 19.91 | 5.24 | 12 | 8.2% | 1.14 [0.27 , 2.01] |
| Chen 2018[a] | 383.54 | 5.7 | 5 | 360.76 | 9.49 | 5 | 4.6% | 2.63 [0.70 , 4.56] |
| El-Derany 2021[a] | 10.24 | 1.99 | 5 | 4.84 | 1.74 | 5 | 4.6% | 2.61 [0.69 , 4.53] |
| Fu 2008[a] | 383.57 | 32.27 | 5 | 364.33 | 34.95 | 5 | 6.7% | 0.52 [-0.76 , 1.79] |
| Fu 2017[a] | 357.32 | 19.51 | 5 | 357.32 | 12 | 5 | 6.8% | 0.00 [-1.24 , 1.24] |
| Khanmohammadi 2018[f] | 36.42 | 8.38 | 20 | 21.5 | 4.33 | 10 | 8.0% | 1.98 [1.05 , 2.91] |
| Lu 2024[a] | 65.02 | 11.61 | 6 | 26.01 | 13.93 | 6 | 5.0% | 2.81 [1.03 , 4.59] |
| Sameni 2019[a] | 32.94 | 4.92 | 10 | 23.83 | 3.07 | 10 | 7.1% | 2.13 [0.98 , 3.27] |
| Tang 2024[c] | 210.82 | 38.57 | 8 | 104.3 | 12.4 | 4 | 4.7% | 2.98 [1.10 , 4.87] |
| Xing 2023[g] | 14.23 | 1.29 | 6 | 5.98 | 1.55 | 6 | 2.8% | 5.34 [2.50 , 8.18] |
| Yang 2020[f] | 15.17 | 1.9 | 20 | 6.03 | 3.16 | 10 | 6.7% | 3.74 [2.47 , 5.01] |
| Zhang 2024[g] | 5.97 | 0.89 | 6 | 2.98 | 1.48 | 6 | 5.6% | 2.26 [0.68 , 3.84] |
| **Subtotal (Wald[d])** | | | **116** | | | **91** | **76.7%** | **2.13 [1.47 , 2.80]** |
| Test for overall effect: Z = 6.33 (P < 0.00001) | | | | | | | | |
| Heterogeneity: Tau² (DL[e]) = 0.89; Chi² = 34.34, df = 12 (P = 0.0006); I² = 65% | | | | | | | | |
| **Total (Wald[d])** | | | **144** | | | **114** | **100.0%** | **2.11 [1.56 , 2.66]** |

Test for overall effect: Z = 7.47 (P < 0.00001)
Test for subgroup differences: Chi² = 0.02, df = 1 (P = 0.90), I² = 0%
Heterogeneity: Tau² (DL[e]) = 0.77; Chi² = 41.48, df = 17 (P = 0.0008); I² = 59%

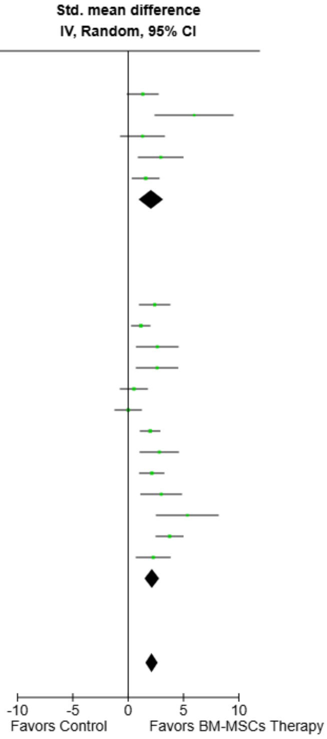


**Footnotes**
[a]BM-MSCs Transplantation
[b]BM-MSCs Secretome
[c]BM-MSCs Secretome and mIR-21 Exosomes
[d]CI calculated by Wald-type method.
[e]Tau² calculated by DerSimonian and Laird method.
[f]BM-MSCs Transplantation and Secretome
[g]BM-MSCs Secretome

**Supplemental Fig. 67. Subgroup analysis of BM-MSC Therapy on primordial follicle count by POI induction model.**

| Study or Subgroup | BM-MSCs Therapy Mean | SD | Total | Control Mean | SD | Total | Weight | Std. mean difference IV, Fixed, 95% CI |
|---|---|---|---|---|---|---|---|---|
| **6.14.1 Intraovarian** | | | | | | | | |
| Chen 2018[a] | 122.5 | 4.16 | 5 | 115 | 3.33 | 5 | 4.0% | 1.80 [0.20 , 3.40] |
| Fu 2008[a] | 132.41 | 29.92 | 5 | 119.02 | 29.94 | 5 | 6.5% | 0.40 [-0.86 , 1.66] |
| Fu 2017[a] | 117.09 | 14.33 | 5 | 116.6 | 7.41 | 5 | 6.7% | 0.04 [-1.20 , 1.28] |
| Khanmohammadi 2018[b] | 29.83 | 6.74 | 20 | 22.5 | 9.67 | 10 | 16.2% | 0.91 [0.12 , 1.71] |
| Ma 2023[a] | 3.48 | 1 | 5 | 2.06 | 0.4 | 5 | 4.2% | 1.68 [0.12 , 3.24] |
| Park 2021b[a] | 3.93 | 1.65 | 3 | 2.89 | 1.65 | 3 | 3.7% | 0.50 [-1.17 , 2.18] |
| Sameni 2019[a] | 22.77 | 3.17 | 10 | 20.85 | 6.24 | 10 | 13.2% | 0.37 [-0.51 , 1.26] |
| Xie 2024[c] | 4.02 | 0.8 | 5 | 0.98 | 0.48 | 5 | 1.5% | 4.16 [1.51 , 6.81] |
| Yang 2024[d] | 4.95 | 1.08 | 10 | 2.68 | 0.65 | 5 | 5.2% | 2.21 [0.79 , 3.62] |
| **Subtotal** | | | **68** | | | **53** | **61.1%** | **0.92 [0.51 , 1.33]** |
| Test for overall effect: Z = 4.39 (P < 0.0001) | | | | | | | | |
| Heterogeneity: Chi² = 15.30, df = 8 (P = 0.05); I² = 48% | | | | | | | | |
| **6.14.2 Intraperitoneal** | | | | | | | | |
| Tang 2024[d] | 139.9 | 31.84 | 8 | 65.2 | 9.45 | 4 | 3.5% | 2.54 [0.81 , 4.27] |
| Yang 2020[b] | 10.02 | 3.39 | 20 | 5.35 | 3.73 | 10 | 14.7% | 1.30 [0.46 , 2.13] |
| **Subtotal** | | | **28** | | | **14** | **18.2%** | **1.53 [0.78 , 2.29]** |
| Test for overall effect: Z = 3.99 (P < 0.0001) | | | | | | | | |
| Heterogeneity: Chi² = 1.62, df = 1 (P = 0.20); I² = 38% | | | | | | | | |
| **6.14.3 Intravenous** | | | | | | | | |
| Bahrehbar 2020[a] | 51.41 | 7.62 | 5 | 32.57 | 12.21 | 5 | 4.3% | 1.67 [0.12 , 3.23] |
| El-Derany 2021[a] | 1.81 | 0.8 | 5 | 0.78 | 0.34 | 5 | 4.6% | 1.51 [0.01 , 3.02] |
| Lu 2024[a] | 20.77 | 10.46 | 6 | 7.85 | 7.08 | 6 | 6.1% | 1.34 [0.03 , 2.64] |
| Xing 2023[c] | 11.04 | 1.26 | 6 | 5.29 | 1.73 | 6 | 2.5% | 3.51 [1.45 , 5.56] |
| Zhang 2024[c] | 5.11 | 0.8 | 6 | 2.58 | 0.89 | 6 | 3.3% | 2.76 [1.00 , 4.52] |
| **Subtotal** | | | **28** | | | **28** | **20.7%** | **1.93 [1.22 , 2.64]** |
| Test for overall effect: Z = 5.36 (P < 0.00001) | | | | | | | | |
| Heterogeneity: Chi² = 4.32, df = 4 (P = 0.36); I² = 7% | | | | | | | | |
| **Total** | | | **124** | | | **95** | **100.0%** | **1.24 [0.92 , 1.56]** |

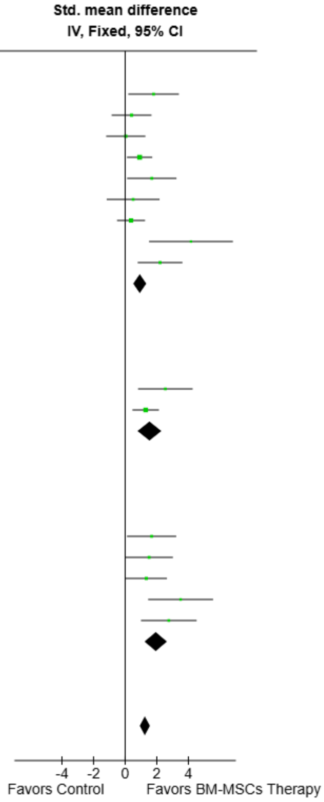


Test for overall effect: Z = 7.57 (P < 0.00001)
Test for subgroup differences: Chi² = 6.58, df = 2 (P = 0.04), I² = 69.6%
Heterogeneity: Chi² = 27.81, df = 15 (P = 0.02); I² = 46%

**Footnotes**
[a]BM-MSCs Transplantation
[b]BM-MSCs Transplantation and Secretome
[c]BM-MSCs Secretome
[d]BM-MSCs Secretome and mIR-21 Exosomes

**Supplemental Fig. 68. Subgroup analysis of BM-MSC Therapy on primary follicle count by route of administration.**

| Study or Subgroup | BM-MSCs Therapy Mean | SD | Total | Control Mean | SD | Total | Weight | Std. mean difference IV, Fixed, 95% CI |
|---|---|---|---|---|---|---|---|---|
| **6.15.1 Mice** | | | | | | | | |
| Bahrehbar 2020[a] | 51.41 | 7.62 | 5 | 32.57 | 12.21 | 5 | 4.3% | 1.67 [0.12 , 3.23] |
| Ma 2023[a] | 3.48 | 1 | 5 | 2.06 | 0.4 | 5 | 4.2% | 1.68 [0.12 , 3.24] |
| Park 2021b[a] | 3.93 | 1.65 | 3 | 2.89 | 1.65 | 3 | 3.7% | 0.50 [-1.17 , 2.18] |
| Xie 2024[b] | 4.02 | 0.8 | 5 | 0.98 | 0.48 | 5 | 1.5% | 4.16 [1.51 , 6.81] |
| Yang 2024[c] | 4.95 | 1.08 | 10 | 2.68 | 0.65 | 5 | 5.2% | 2.21 [0.79 , 3.62] |
| **Subtotal** | | | **28** | | | **23** | **18.8%** | **1.79 [1.05 , 2.53]** |
| Test for overall effect: Z = 4.73 (P < 0.00001) | | | | | | | | |
| Heterogeneity: Chi² = 5.72, df = 4 (P = 0.22); I² = 30% | | | | | | | | |
| **6.15.2 Rats** | | | | | | | | |
| Chen 2018[a] | 122.5 | 4.16 | 5 | 115 | 3.33 | 5 | 4.0% | 1.80 [0.20 , 3.40] |
| El-Derany 2021[a] | 1.81 | 0.8 | 5 | 0.78 | 0.34 | 5 | 4.6% | 1.51 [0.01 , 3.02] |
| Fu 2008[a] | 132.41 | 29.92 | 5 | 119.02 | 29.94 | 5 | 6.5% | 0.40 [-0.86 , 1.66] |
| Fu 2017[a] | 117.09 | 14.33 | 5 | 116.6 | 7.41 | 5 | 6.7% | 0.04 [-1.20 , 1.28] |
| Khanmohammadi 2018[d] | 29.83 | 6.74 | 20 | 22.5 | 9.67 | 10 | 16.2% | 0.91 [0.12 , 1.71] |
| Lu 2024[a] | 20.77 | 10.46 | 6 | 7.85 | 7.08 | 6 | 6.1% | 1.34 [0.03 , 2.64] |
| Sameni 2019[a] | 22.77 | 3.17 | 10 | 20.85 | 6.24 | 10 | 13.2% | 0.37 [-0.51 , 1.26] |
| Tang 2024[c] | 139.9 | 31.84 | 8 | 65.2 | 9.45 | 4 | 3.5% | 2.54 [0.81 , 4.27] |
| Xing 2023[b] | 11.04 | 1.26 | 6 | 5.29 | 1.73 | 6 | 2.5% | 3.51 [1.45 , 5.56] |
| Yang 2020[d] | 10.02 | 3.39 | 20 | 5.35 | 3.73 | 10 | 14.7% | 1.30 [0.46 , 2.13] |
| Zhang 2024[b] | 5.11 | 0.8 | 6 | 2.58 | 0.89 | 6 | 3.3% | 2.76 [1.00 , 4.52] |
| **Subtotal** | | | **96** | | | **72** | **81.2%** | **1.11 [0.76 , 1.47]** |
| Test for overall effect: Z = 6.13 (P < 0.00001) | | | | | | | | |
| Heterogeneity: Chi² = 19.52, df = 10 (P = 0.03); I² = 49% | | | | | | | | |
| **Total** | | | **124** | | | **95** | **100.0%** | **1.24 [0.92 , 1.56]** |

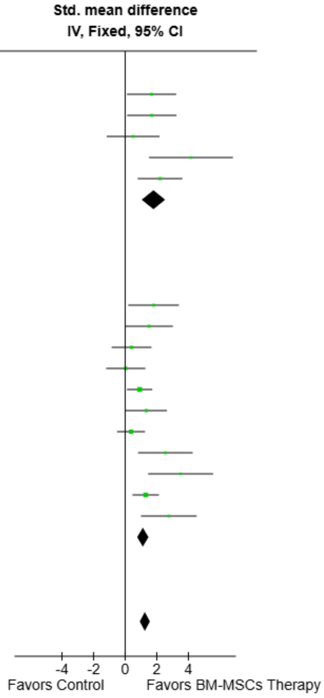


Test for overall effect: Z = 7.57 (P < 0.00001)
Test for subgroup differences: Chi² = 2.57, df = 1 (P = 0.11), I² = 61.0%
Heterogeneity: Chi² = 27.81, df = 15 (P = 0.02); I² = 46%

**Footnotes**
[a]BM-MSCs Transplantation
[b]BM-MSCs Secretome
[c]BM-MSCs Secretome and mIR-21 Exosomes
[d]BM-MSCs Transplantation and Secretome

**Supplemental Fig. 69. Subgroup analysis of BM-MSC Therapy on primary follicle count by animal model.**

| Study or Subgroup | BM-MSCs Therapy Mean | SD | Total | Control Mean | SD | Total | Weight | Std. mean difference IV, Fixed, 95% CI |
|---|---|---|---|---|---|---|---|---|
| **6.16.1 Autoimmune** | | | | | | | | |
| Ma 2023[a] | 3.48 | 1 | 5 | 2.06 | 0.4 | 5 | 4.2% | 1.68 [0.12 , 3.24] |
| Xie 2024[b] | 4.02 | 0.8 | 5 | 0.98 | 0.48 | 5 | 1.5% | 4.16 [1.51 , 6.81] |
| Yang 2024[c] | 4.95 | 1.08 | 10 | 2.68 | 0.65 | 5 | 5.2% | 2.21 [0.79 , 3.62] |
| **Subtotal** | | | **20** | | | **15** | **10.9%** | **2.27 [1.29 , 3.24]** |
| Test for overall effect: Z = 4.56 (P < 0.00001) | | | | | | | | |
| Heterogeneity: Chi² = 2.51, df = 2 (P = 0.29); I² = 20% | | | | | | | | |
| **6.16.2 Chemotherapy (Cisplatin)** | | | | | | | | |
| Tang 2024[c] | 139.9 | 31.84 | 8 | 65.2 | 9.45 | 4 | 3.5% | 2.54 [0.81 , 4.27] |
| **Subtotal** | | | **8** | | | **4** | **3.5%** | **2.54 [0.81 , 4.27]** |
| Test for overall effect: Z = 2.89 (P = 0.004) | | | | | | | | |
| Heterogeneity: Not applicable | | | | | | | | |
| **6.16.3 Chemotherapy (Cyclophosphamide)** | | | | | | | | |
| Bahrehbar 2020[a] | 51.41 | 7.62 | 5 | 32.57 | 12.21 | 5 | 4.3% | 1.67 [0.12 , 3.23] |
| Chen 2018[a] | 122.5 | 4.16 | 5 | 115 | 3.33 | 5 | 4.0% | 1.80 [0.20 , 3.40] |
| Fu 2008[a] | 132.41 | 29.92 | 5 | 119.02 | 29.94 | 5 | 6.5% | 0.40 [-0.86 , 1.66] |
| Fu 2017[a] | 117.09 | 14.33 | 5 | 116.6 | 7.41 | 5 | 6.7% | 0.04 [-1.20 , 1.28] |
| Khanmohammadi 2018[d] | 29.83 | 6.74 | 20 | 22.5 | 9.67 | 10 | 16.2% | 0.91 [0.12 , 1.71] |
| Lu 2024[a] | 20.77 | 10.46 | 6 | 7.85 | 7.08 | 6 | 6.1% | 1.34 [0.03 , 2.64] |
| Park 2021b[a] | 3.93 | 1.65 | 3 | 2.89 | 1.65 | 3 | 3.7% | 0.50 [-1.17 , 2.18] |
| Sameni 2019[a] | 22.77 | 3.17 | 10 | 20.85 | 6.24 | 10 | 13.2% | 0.37 [-0.51 , 1.26] |
| Xing 2023[b] | 11.04 | 1.26 | 6 | 5.29 | 1.73 | 6 | 2.5% | 3.51 [1.45 , 5.56] |
| Yang 2020[d] | 10.02 | 3.39 | 20 | 5.35 | 3.73 | 10 | 14.7% | 1.30 [0.46 , 2.13] |
| Zhang 2024[b] | 5.11 | 0.8 | 6 | 2.58 | 0.89 | 6 | 3.3% | 2.76 [1.00 , 4.52] |
| **Subtotal** | | | **91** | | | **71** | **81.1%** | **1.03 [0.68 , 1.39]** |
| Test for overall effect: Z = 5.68 (P < 0.00001) | | | | | | | | |
| Heterogeneity: Chi² = 17.44, df = 10 (P = 0.07); I² = 43% | | | | | | | | |
| **6.16.4 Radiotherapy** | | | | | | | | |
| El-Derany 2021[a] | 1.81 | 0.8 | 5 | 0.78 | 0.34 | 5 | 4.6% | 1.51 [0.01 , 3.02] |
| **Subtotal** | | | **5** | | | **5** | **4.6%** | **1.51 [0.01 , 3.02]** |
| Test for overall effect: Z = 1.97 (P = 0.05) | | | | | | | | |
| Heterogeneity: Not applicable | | | | | | | | |
| **Total** | | | **124** | | | **95** | **100.0%** | **1.24 [0.92 , 1.56]** |

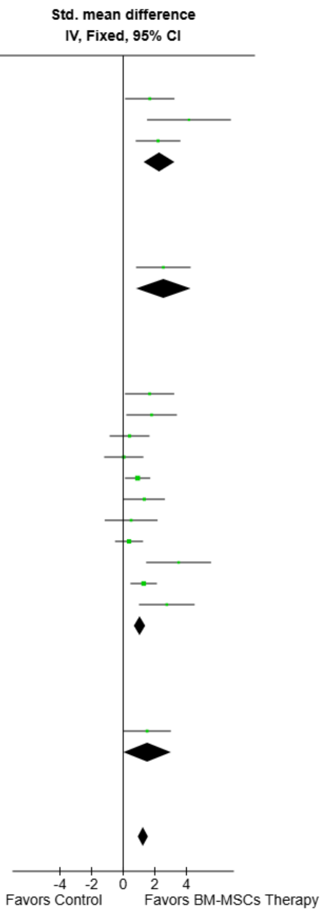


Test for overall effect: Z = 7.57 (P < 0.00001)
Test for subgroup differences: Chi² = 7.87, df = 3 (P = 0.05), I² = 61.9%
Heterogeneity: Chi² = 27.81, df = 15 (P = 0.02); I² = 46%

**Footnotes**
[a]BM-MSCs Transplantation
[b]BM-MSCs Secretome
[c]BM-MSCs Secretome and mIR-21 Exosomes
[d]BM-MSCs Transplantation and Secretome

**Supplemental Fig. 70. Subgroup analysis of BM-MSC Therapy on primary follicle count by POI induction model.**

| Study or Subgroup | BM-MSCs Therapy Mean | SD | Total | Control Mean | SD | Total | Weight | Std. mean difference IV, Random, 95% CI |
|---|---|---|---|---|---|---|---|---|
| **6.17.1 Intraovarian** | | | | | | | | |
| Chen 2018[a] | 51.18 | 2.03 | 5 | 44.11 | 2.02 | 5 | 5.3% | 3.15 [0.99 , 5.32] |
| Fu 2008[a] | 51.98 | 8.47 | 5 | 45.94 | 14.78 | 5 | 8.3% | 0.45 [-0.81 , 1.72] |
| Fu 2017[a] | 47.33 | 10.29 | 5 | 42.18 | 3.17 | 5 | 8.2% | 0.61 [-0.68 , 1.90] |
| Khanmohammadi 2018[b] | 28.34 | 5.53 | 20 | 21.67 | 6.17 | 10 | 10.1% | 1.13 [0.31 , 1.95] |
| Ma 2023[a] | 1.3 | 0.81 | 5 | 1.47 | 0.5 | 5 | 8.4% | -0.23 [-1.47 , 1.02] |
| Park 2021b[a] | 4.12 | 1.4 | 3 | 3.2 | 1.69 | 3 | 6.8% | 0.47 [-1.19 , 2.14] |
| Sameni 2019[a] | 30.79 | 3.17 | 10 | 17.88 | 6.19 | 10 | 8.4% | 2.51 [1.28 , 3.75] |
| Xie 2024[c] | 2.23 | 0.29 | 5 | 0.7 | 0.45 | 5 | 4.7% | 3.65 [1.25 , 6.05] |
| Yang 2024[d] | 4.02 | 1.2 | 10 | 2.68 | 0.82 | 5 | 8.7% | 1.15 [-0.03 , 2.33] |
| **Subtotal (Wald[e])** | | | **68** | | | **53** | **68.9%** | **1.22 [0.52 , 1.92]** |
| Test for overall effect: Z = 3.42 (P = 0.0006) | | | | | | | | |
| Heterogeneity: Tau$^2$ (DL[f]) = 0.63; Chi$^2$ = 19.30, df = 8 (P = 0.01); I$^2$ = 59% | | | | | | | | |
| **6.17.2 Intraperitoneal** | | | | | | | | |
| Tang 2024[d] | 123.63 | 9.08 | 8 | 43.56 | 6.82 | 4 | 1.9% | 8.73 [4.30 , 13.16] |
| Yang 2020[b] | 7.84 | 5.4 | 20 | 3.98 | 3.35 | 10 | 10.2% | 0.78 [-0.01 , 1.56] |
| **Subtotal (Wald[e])** | | | **28** | | | **14** | **12.1%** | **4.44 [-3.33 , 12.21]** |
| Test for overall effect: Z = 1.12 (P = 0.26) | | | | | | | | |
| Heterogeneity: Tau$^2$ (DL[f]) = 29.00; Chi$^2$ = 12.02, df = 1 (P = 0.0005); I$^2$ = 92% | | | | | | | | |
| **6.17.3 Intravenous** | | | | | | | | |
| El-Derany 2021[a] | 2.82 | 0.76 | 5 | 0.78 | 0.87 | 5 | 6.4% | 2.26 [0.48 , 4.03] |
| Xing 2023[c] | 8.31 | 1.77 | 6 | 2.97 | 1.33 | 6 | 6.0% | 3.15 [1.24 , 5.06] |
| Zhang 2024[c] | 4.09 | 0.8 | 6 | 1.72 | 0.83 | 6 | 6.6% | 2.68 [0.95 , 4.42] |
| **Subtotal (Wald[e])** | | | **17** | | | **17** | **19.0%** | **2.67 [1.64 , 3.71]** |
| Test for overall effect: Z = 5.04 (P < 0.00001) | | | | | | | | |
| Heterogeneity: Tau$^2$ (DL[f]) = 0.00; Chi$^2$ = 0.45, df = 2 (P = 0.80); I$^2$ = 0% | | | | | | | | |
| **Total (Wald[e])** | | | **113** | | | **84** | **100.0%** | **1.62 [0.95 , 2.28]** |

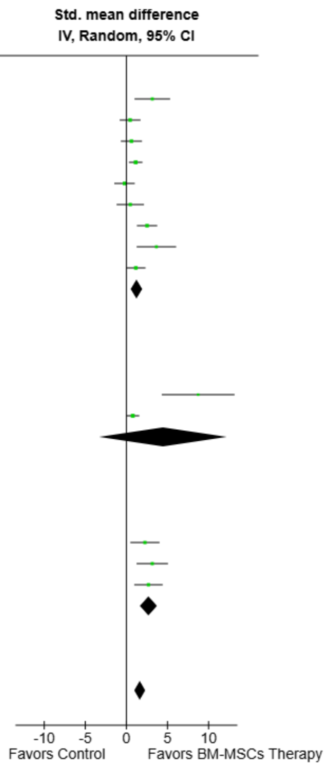


Test for overall effect: Z = 4.77 (P < 0.00001)
Test for subgroup differences: Chi$^2$ = 5.66, df = 2 (P = 0.06), I$^2$ = 64.6%
Heterogeneity: Tau$^2$ (DL[f]) = 0.97; Chi$^2$ = 39.66, df = 13 (P = 0.0002); I$^2$ = 67%

**Footnotes**
[a]BM-MSCs Transplantation
[b]BM-MSCs Transplantation and Secretome
[c]BM-MSCs Secretome
[d]BM-MSCs Secretome and mIR-21 Exosomes
[e]CI calculated by Wald-type method.
[f]Tau$^2$ calculated by DerSimonian and Laird method.

**Supplemental Fig. 71. Subgroup analysis of BM-MSC Therapy on secondary follicle count by route of administration.**

| Study or Subgroup | BM-MSCs Therapy Mean | SD | Total | Control Mean | SD | Total | Weight | Std. mean difference IV, Random, 95% CI |
|---|---|---|---|---|---|---|---|---|
| **6.18.1 Mice** | | | | | | | | |
| Ma 2023[a] | 1.3 | 0.81 | 5 | 1.47 | 0.5 | 5 | 8.4% | -0.23 [-1.47 , 1.02] |
| Park 2021b[a] | 4.12 | 1.4 | 3 | 3.2 | 1.69 | 3 | 6.8% | 0.47 [-1.19 , 2.14] |
| Xie 2024[b] | 2.23 | 0.29 | 5 | 0.7 | 0.45 | 5 | 4.7% | 3.65 [1.25 , 6.05] |
| Yang 2024[c] | 4.02 | 1.2 | 10 | 2.68 | 0.82 | 5 | 8.7% | 1.15 [-0.03 , 2.33] |
| **Subtotal (Wald[d])** | | | **23** | | | **18** | **28.5%** | **1.01 [-0.29 , 2.30]** |
| Test for overall effect: Z = 1.53 (P = 0.13) | | | | | | | | |
| Heterogeneity: Tau² (DL[e]) = 1.08; Chi² = 8.52, df = 3 (P = 0.04); I² = 65% | | | | | | | | |
| **6.18.2 Rats** | | | | | | | | |
| Chen 2018[a] | 51.18 | 2.03 | 5 | 44.11 | 2.02 | 5 | 5.3% | 3.15 [0.99 , 5.32] |
| El-Derany 2021[a] | 2.82 | 0.76 | 5 | 0.78 | 0.87 | 5 | 6.4% | 2.26 [0.48 , 4.03] |
| Fu 2008[a] | 51.98 | 8.47 | 5 | 45.94 | 14.78 | 5 | 8.3% | 0.45 [-0.81 , 1.72] |
| Fu 2017[a] | 47.33 | 10.29 | 5 | 42.18 | 3.17 | 5 | 8.2% | 0.61 [-0.68 , 1.90] |
| Khanmohammadi 2018[f] | 28.34 | 5.53 | 20 | 21.67 | 6.17 | 10 | 10.1% | 1.13 [0.31 , 1.95] |
| Sameni 2019[a] | 30.79 | 3.17 | 10 | 17.88 | 6.19 | 10 | 8.4% | 2.51 [1.28 , 3.75] |
| Tang 2024[c] | 123.63 | 9.08 | 8 | 43.56 | 6.82 | 4 | 1.9% | 8.73 [4.30 , 13.16] |
| Xing 2023[b] | 8.31 | 1.77 | 6 | 2.97 | 1.33 | 6 | 6.0% | 3.15 [1.24 , 5.06] |
| Yang 2020[f] | 7.84 | 5.4 | 20 | 3.98 | 3.35 | 10 | 10.2% | 0.78 [-0.01 , 1.56] |
| Zhang 2024[b] | 4.09 | 0.8 | 6 | 1.72 | 0.83 | 6 | 6.6% | 2.68 [0.95 , 4.42] |
| **Subtotal (Wald[d])** | | | **90** | | | **66** | **71.5%** | **1.88 [1.08 , 2.67]** |
| Test for overall effect: Z = 4.62 (P < 0.00001) | | | | | | | | |
| Heterogeneity: Tau² (DL[e]) = 1.00; Chi² = 28.81, df = 9 (P = 0.0007); I² = 69% | | | | | | | | |
| **Total (Wald[d])** | | | **113** | | | **84** | **100.0%** | **1.62 [0.95 , 2.28]** |

Test for overall effect: Z = 4.77 (P < 0.00001)
Test for subgroup differences: Chi² = 1.27, df = 1 (P = 0.26), I² = 21.0%
Heterogeneity: Tau² (DL[e]) = 0.97; Chi² = 39.66, df = 13 (P = 0.0002); I² = 67%

Std. mean difference IV, Random, 95% CI

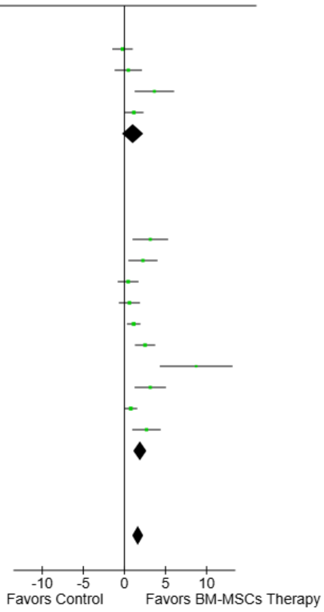


**Footnotes**
[a]BM-MSCs Transplantation
[b]BM-MSCs Secretome
[c]BM-MSCs Secretome and mIR-21 Exosomes
[d]CI calculated by Wald-type method.
[e]Tau² calculated by DerSimonian and Laird method.
[f]BM-MSCs Transplantation and Secretome

**Supplemental Fig. 72. Subgroup analysis of BM-MSC Therapy on secondary follicle count by animal model.**

| Study or Subgroup | BM-MSCs Therapy Mean | SD | Total | Control Mean | SD | Total | Weight | Std. mean difference IV, Random, 95% CI |
|---|---|---|---|---|---|---|---|---|
| **6.19.1 Autoimmune** | | | | | | | | |
| Ma 2023[a] | 1.3 | 0.81 | 5 | 1.47 | 0.5 | 5 | 8.4% | -0.23 [-1.47 , 1.02] |
| Xie 2024[b] | 2.23 | 0.29 | 5 | 0.7 | 0.45 | 5 | 4.7% | 3.65 [1.25 , 6.05] |
| Yang 2024[c] | 4.02 | 1.2 | 10 | 2.68 | 0.82 | 5 | 8.7% | 1.15 [-0.03 , 2.33] |
| **Subtotal (Wald[d])** | | | **20** | | | **15** | **21.7%** | **1.26 [-0.52 , 3.04]** |
| Test for overall effect: Z = 1.39 (P = 0.17) | | | | | | | | |
| Heterogeneity: Tau² (DL[e]) = 1.81; Chi² = 8.35, df = 2 (P = 0.02); I² = 76% | | | | | | | | |
| **6.19.2 Chemotherapy (Cisplatin)** | | | | | | | | |
| Tang 2024[c] | 123.63 | 9.08 | 8 | 43.56 | 6.82 | 4 | 1.9% | 8.73 [4.30 , 13.16] |
| **Subtotal** | | | **8** | | | **4** | **1.9%** | **8.73 [4.30 , 13.16]** |
| Test for overall effect: Z = 3.86 (P = 0.0001) | | | | | | | | |
| Heterogeneity: Not applicable | | | | | | | | |
| **6.19.3 Chemotherapy (Cyclophosphamide)** | | | | | | | | |
| Chen 2018[a] | 51.18 | 2.03 | 5 | 44.11 | 2.02 | 5 | 5.3% | 3.15 [0.99 , 5.32] |
| Fu 2008[a] | 51.98 | 8.47 | 5 | 45.94 | 14.78 | 5 | 8.3% | 0.45 [-0.81 , 1.72] |
| Fu 2017[a] | 47.33 | 10.29 | 5 | 42.18 | 3.17 | 5 | 8.2% | 0.61 [-0.68 , 1.90] |
| Khanmohammadi 2018[f] | 28.34 | 5.53 | 20 | 21.67 | 6.17 | 10 | 10.1% | 1.13 [0.31 , 1.95] |
| Park 2021b[a] | 4.12 | 1.4 | 3 | 3.2 | 1.69 | 3 | 6.8% | 0.47 [-1.19 , 2.14] |
| Sameni 2019[a] | 30.79 | 3.17 | 10 | 17.88 | 6.19 | 10 | 8.4% | 2.51 [1.28 , 3.75] |
| Xing 2023[b] | 8.31 | 1.77 | 6 | 2.97 | 1.33 | 6 | 6.0% | 3.15 [1.24 , 5.06] |
| Yang 2020[f] | 7.84 | 5.4 | 20 | 3.98 | 3.35 | 10 | 10.2% | 0.78 [-0.01 , 1.56] |
| Zhang 2024[b] | 4.09 | 0.8 | 6 | 1.72 | 0.83 | 6 | 6.6% | 2.68 [0.95 , 4.42] |
| **Subtotal (Wald[d])** | | | **80** | | | **60** | **69.9%** | **1.46 [0.80 , 2.11]** |
| Test for overall effect: Z = 4.36 (P < 0.0001) | | | | | | | | |
| Heterogeneity: Tau² (DL[e]) = 0.52; Chi² = 18.20, df = 8 (P = 0.02); I² = 56% | | | | | | | | |
| **6.19.4 Radiotherapy** | | | | | | | | |
| El-Derany 2021[a] | 2.82 | 0.76 | 5 | 0.78 | 0.87 | 5 | 6.4% | 2.26 [0.48 , 4.03] |
| **Subtotal** | | | **5** | | | **5** | **6.4%** | **2.26 [0.48 , 4.03]** |
| Test for overall effect: Z = 2.49 (P = 0.01) | | | | | | | | |
| Heterogeneity: Not applicable | | | | | | | | |
| **Total (Wald[d])** | | | **113** | | | **84** | **100.0%** | **1.62 [0.95 , 2.28]** |

Test for overall effect: Z = 4.77 (P < 0.00001)
Test for subgroup differences: Chi² = 10.79, df = 3 (P = 0.01), I² = 72.2%
Heterogeneity: Tau² (DL[e]) = 0.97; Chi² = 39.66, df = 13 (P = 0.0002); I² = 67%

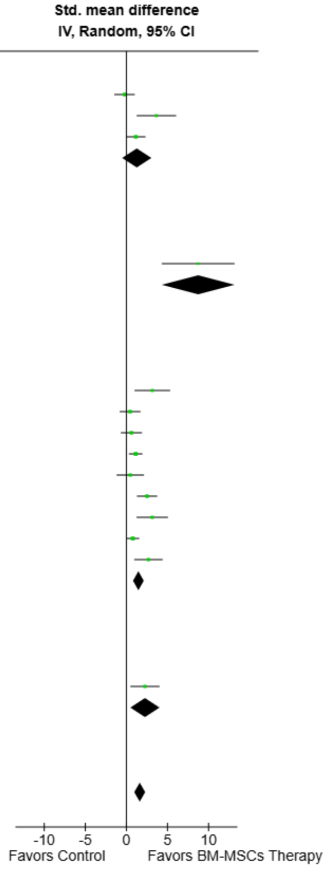


**Footnotes**
[a]BM-MSCs Transplantation
[b]BM-MSCs Secretome
[c]BM-MSCs Secretome and mIR-21 Exosomes
[d]CI calculated by Wald-type method.
[e]Tau² calculated by DerSimonian and Laird method.
[f]BM-MSCs Transplantation and Secretome

**Supplemental Fig. 73. Subgroup analysis of BM-MSC Therapy on secondary follicle count by POI induction model.**

| Study or Subgroup | BM-MSCs Therapy Events | BM-MSCs Therapy Total | Control Events | Control Total | Weight | Risk ratio M-H, Fixed, 95% CI |
|---|---|---|---|---|---|---|
| **6.20.1 Intraovarian** | | | | | | |
| Chen 2018[a] | 6 | 15 | 0 | 15 | 9.9% | 13.00 [0.80 , 212.02] |
| Fu 2008[a] | 6 | 20 | 0 | 20 | 9.9% | 13.00 [0.78 , 216.39] |
| Fu 2017[a] | 6 | 15 | 0 | 15 | 9.9% | 13.00 [0.80 , 212.02] |
| Ma 2023[a] | 4 | 6 | 0 | 5 | 10.7% | 7.71 [0.51 , 116.01] |
| **Subtotal** | | **56** | | **55** | **40.5%** | **11.60 [2.90 , 46.46]** |
| Total events: | 22 | | 0 | | | |
| Test for overall effect: Z = 3.46 (P = 0.0005) | | | | | | |
| Heterogeneity: Chi² = 0.11, df = 3 (P = 0.99); I² = 0% | | | | | | |
| **6.20.2 Intraperitoneal** | | | | | | |
| Yang 2020[b] | 7 | 20 | 1 | 10 | 26.5% | 3.50 [0.50 , 24.67] |
| **Subtotal** | | **20** | | **10** | **26.5%** | **3.50 [0.50 , 24.67]** |
| Total events: | 7 | | 1 | | | |
| Test for overall effect: Z = 1.26 (P = 0.21) | | | | | | |
| Heterogeneity: Not applicable | | | | | | |
| **6.20.3 Intravenous** | | | | | | |
| Lu 2024[a] | 7 | 9 | 0 | 8 | 10.5% | 13.50 [0.89 , 204.40] |
| Park 2023[a] | 3 | 14 | 0 | 6 | 13.5% | 3.27 [0.19 , 55.03] |
| Park 2023[c] | 1 | 4 | 0 | 5 | 9.0% | 3.60 [0.18 , 70.34] |
| **Subtotal** | | **27** | | **19** | **33.0%** | **6.60 [1.34 , 32.46]** |
| Total events: | 11 | | 0 | | | |
| Test for overall effect: Z = 2.32 (P = 0.02) | | | | | | |
| Heterogeneity: Chi² = 0.66, df = 2 (P = 0.72); I² = 0% | | | | | | |
| **Total** | | **103** | | **84** | **100.0%** | **7.80 [3.15 , 19.34]** |
| Total events: | 40 | | 1 | | | |
| Test for overall effect: Z = 4.44 (P < 0.00001) | | | | | | |
| Test for subgroup differences: Chi² = 0.99, df = 2 (P = 0.61), I² = 0% | | | | | | |
| Heterogeneity: Chi² = 1.81, df = 7 (P = 0.97); I² = 0% | | | | | | |

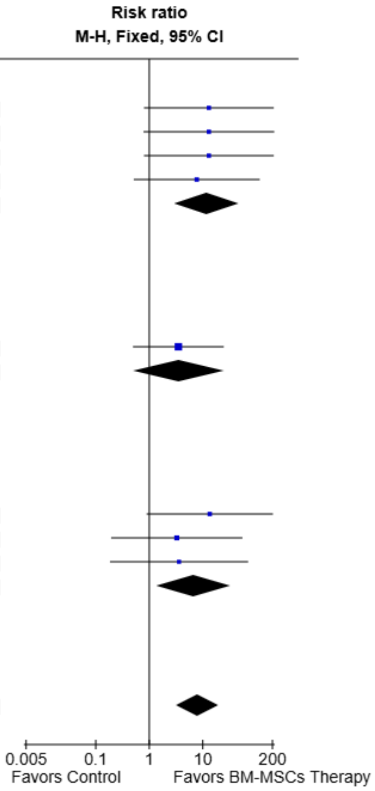


**Footnotes**

[a]BM-MSCs Transplantation

[b]BM-MSCs Transplantation and Secretome

[c]BM-MSCs Secretome

**Supplemental Fig. 74. Subgroup analysis of BM-MSC Therapy on normal estrous cycle by route of administration.**

| Study or Subgroup | BM-MSCs Therapy Events | BM-MSCs Therapy Total | Control Events | Control Total | Weight | Risk ratio M-H, Fixed, 95% CI |
|---|---|---|---|---|---|---|
| **6.21.1 Mice** | | | | | | |
| Ma 2023[a] | 4 | 6 | 0 | 5 | 10.7% | 7.71 [0.51 , 116.01] |
| Park 2023[a] | 3 | 14 | 0 | 6 | 13.5% | 3.27 [0.19 , 55.03] |
| Park 2023[b] | 1 | 4 | 0 | 5 | 9.0% | 3.60 [0.18 , 70.34] |
| **Subtotal** | | **24** | | **16** | **33.3%** | **4.79 [0.95 , 24.12]** |
| Total events: | 8 | | 0 | | | |
| Test for overall effect: Z = 1.90 (P = 0.06) | | | | | | |
| Heterogeneity: Chi² = 0.22, df = 2 (P = 0.89); I² = 0% | | | | | | |
| **6.21.2 Rats** | | | | | | |
| Chen 2018[a] | 6 | 15 | 0 | 15 | 9.9% | 13.00 [0.80 , 212.02] |
| Fu 2008[a] | 6 | 20 | 0 | 20 | 9.9% | 13.00 [0.78 , 216.39] |
| Fu 2017[a] | 6 | 15 | 0 | 15 | 9.9% | 13.00 [0.80 , 212.02] |
| Lu 2024[a] | 7 | 9 | 0 | 8 | 10.5% | 13.50 [0.89 , 204.40] |
| Yang 2020[c] | 7 | 20 | 1 | 10 | 26.5% | 3.50 [0.50 , 24.67] |
| **Subtotal** | | **79** | | **68** | **66.7%** | **9.31 [3.09 , 28.03]** |
| Total events: | 32 | | 1 | | | |
| Test for overall effect: Z = 3.97 (P < 0.0001) | | | | | | |
| Heterogeneity: Chi² = 1.20, df = 4 (P = 0.88); I² = 0% | | | | | | |
| **Total** | | **103** | | **84** | **100.0%** | **7.80 [3.15 , 19.34]** |
| Total events: | 40 | | 1 | | | |
| Test for overall effect: Z = 4.44 (P < 0.00001) | | | | | | |
| Test for subgroup differences: Chi² = 0.44, df = 1 (P = 0.51), I² = 0% | | | | | | |
| Heterogeneity: Chi² = 1.81, df = 7 (P = 0.97); I² = 0% | | | | | | |

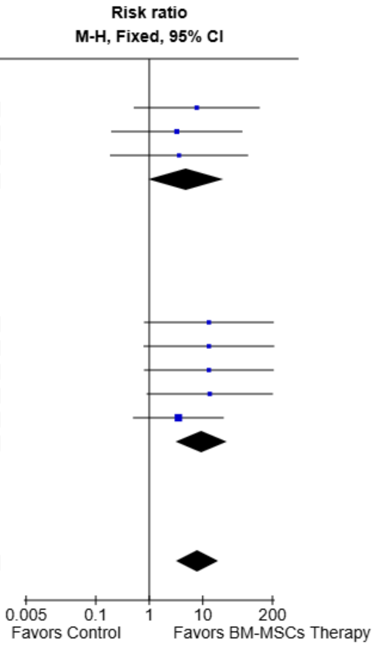


**Footnotes**
[a]BM-MSCs Transplantation
[b]BM-MSCs Secretome
[c]BM-MSCs Transplantation and Secretome

**Supplemental Fig. 75. Subgroup analysis of BM-MSC Therapy on normal estrous cycle by animal model.**

| Study or Subgroup | BM-MSCs Therapy Events | BM-MSCs Therapy Total | Control Events | Control Total | Weight | Risk ratio M-H, Fixed, 95% CI |
|---|---|---|---|---|---|---|
| **6.22.1 Autoimmune** | | | | | | |
| Ma 2023[a] | 4 | 6 | 0 | 5 | 10.7% | 7.71 [0.51 , 116.01] |
| **Subtotal** | | **6** | | **5** | **10.7%** | **7.71 [0.51 , 116.01]** |
| Total events: | 4 | | 0 | | | |
| Test for overall effect: Z = 1.48 (P = 0.14) | | | | | | |
| Heterogeneity: Not applicable | | | | | | |
| **6.22.2 Chemotherapy (Cyclophosphamide)** | | | | | | |
| Chen 2018[a] | 6 | 15 | 0 | 15 | 9.9% | 13.00 [0.80 , 212.02] |
| Fu 2008[a] | 6 | 20 | 0 | 20 | 9.9% | 13.00 [0.78 , 216.39] |
| Fu 2017[a] | 6 | 15 | 0 | 15 | 9.9% | 13.00 [0.80 , 212.02] |
| Lu 2024[a] | 7 | 9 | 0 | 8 | 10.5% | 13.50 [0.89 , 204.40] |
| Park 2023[a] | 3 | 14 | 0 | 6 | 13.5% | 3.27 [0.19 , 55.03] |
| Park 2023[b] | 1 | 4 | 0 | 5 | 9.0% | 3.60 [0.18 , 70.34] |
| Yang 2020[c] | 7 | 20 | 1 | 10 | 26.5% | 3.50 [0.50 , 24.67] |
| **Subtotal** | | **97** | | **79** | **89.3%** | **7.81 [2.98 , 20.46]** |
| Total events: | 36 | | 1 | | | |
| Test for overall effect: Z = 4.19 (P < 0.0001) | | | | | | |
| Heterogeneity: Chi² = 1.81, df = 6 (P = 0.94); I² = 0% | | | | | | |
| **Total** | | **103** | | **84** | **100.0%** | **7.80 [3.15 , 19.34]** |
| Total events: | 40 | | 1 | | | |
| Test for overall effect: Z = 4.44 (P < 0.00001) | | | | | | |
| Test for subgroup differences: Chi² = 0.00, df = 1 (P = 0.99), I² = 0% | | | | | | |
| Heterogeneity: Chi² = 1.81, df = 7 (P = 0.97); I² = 0% | | | | | | |

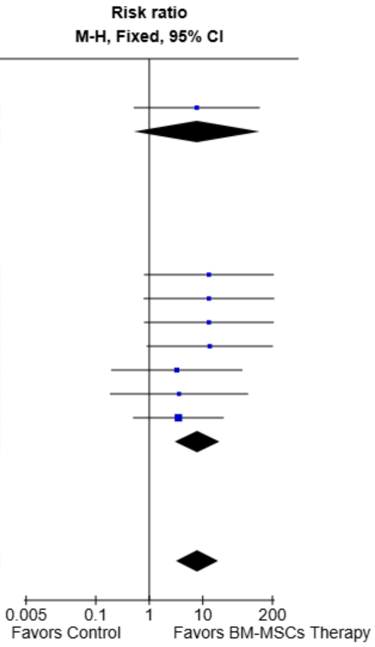


**Footnotes**
[a]BM-MSCs Transplantation
[b]BM-MSCs Secretome
[c]BM-MSCs Transplantation and Secretome

**Supplemental Fig. 76. Subgroup analysis of BM-MSC Therapy on normal estrous cycle by POI induction model.**

| Study or Subgroup | BM-MSCs Therapy Events | BM-MSCs Therapy Total | Control Events | Control Total | Weight | Risk ratio M-H, Fixed, 95% CI |
|---|---|---|---|---|---|---|
| **6.23.1 Intraovarian** | | | | | | |
| El-Derany 2021[a] | 3 | 5 | 1 | 5 | 9.0% | 3.00 [0.45 , 19.93] |
| Park 2021a[a] | 7 | 12 | 2 | 12 | 17.9% | 3.50 [0.91 , 13.53] |
| Xie 2024[b] | 3 | 4 | 1 | 4 | 9.0% | 3.00 [0.50 , 17.95] |
| **Subtotal** | | **21** | | **21** | **35.8%** | **3.25 [1.27 , 8.32]** |
| Total events: | 13 | | 4 | | | |
| Test for overall effect: Z = 2.46 (P = 0.01) | | | | | | |
| Heterogeneity: Chi² = 0.03, df = 2 (P = 0.99); I² = 0% | | | | | | |
| **6.23.2 Intraperitoneal** | | | | | | |
| Tang 2024[c] | 6 | 8 | 2 | 4 | 23.9% | 1.50 [0.52 , 4.32] |
| **Subtotal** | | **8** | | **4** | **23.9%** | **1.50 [0.52 , 4.32]** |
| Total events: | 6 | | 2 | | | |
| Test for overall effect: Z = 0.75 (P = 0.45) | | | | | | |
| Heterogeneity: Not applicable | | | | | | |
| **6.23.3 Intravenous** | | | | | | |
| Badawy 2017[a] | 5 | 5 | 0 | 5 | 4.5% | 11.00 [0.77 , 158.01] |
| Bahrehbar 2020[a] | 2 | 5 | 0 | 5 | 4.5% | 5.00 [0.30 , 83.69] |
| Park 2023[a] | 11 | 12 | 3 | 12 | 26.9% | 3.67 [1.36 , 9.91] |
| Zhang 2024[b] | 5 | 6 | 0 | 6 | 4.5% | 11.00 [0.74 , 163.49] |
| **Subtotal** | | **28** | | **28** | **40.3%** | **5.44 [2.30 , 12.91]** |
| Total events: | 23 | | 3 | | | |
| Test for overall effect: Z = 3.85 (P = 0.0001) | | | | | | |
| Heterogeneity: Chi² = 1.14, df = 3 (P = 0.77); I² = 0% | | | | | | |
| **Total** | | **57** | | **53** | **100.0%** | **3.72 [2.14 , 6.44]** |
| Total events: | 42 | | 9 | | | |
| Test for overall effect: Z = 4.68 (P < 0.00001) | | | | | | |
| Test for subgroup differences: Chi² = 3.42, df = 2 (P = 0.18), I² = 41.6% | | | | | | |
| Heterogeneity: Chi² = 4.24, df = 7 (P = 0.75); I² = 0% | | | | | | |

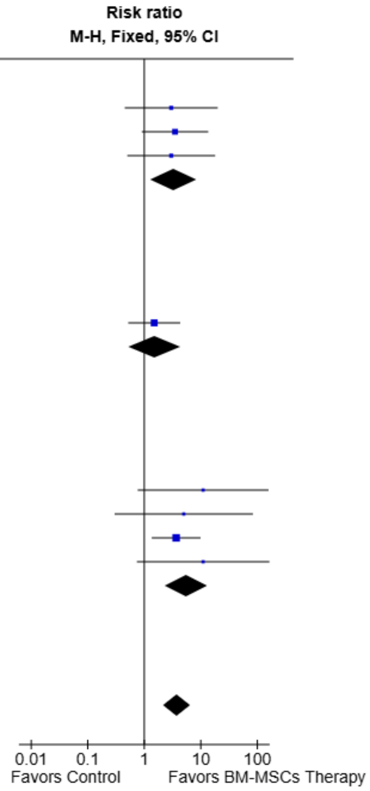


**Footnotes**

[a]BM-MSCs Transplantation

[b]BM-MSCs Secretome

[c]BM-MSCs Secretome and mIR-21 Exosomes

**Supplemental Fig. 77. Subgroup analysis of BM-MSC Therapy on pregnancy occurrence by route of administration.**

| Study or Subgroup | BM-MSCs Therapy Events | BM-MSCs Therapy Total | Control Events | Control Total | Weight | Risk ratio M-H, Fixed, 95% CI |
|---|---|---|---|---|---|---|
| **6.24.1 Mice** | | | | | | |
| Badawy 2017[a] | 5 | 5 | 0 | 5 | 4.5% | 11.00 [0.77 , 158.01] |
| Bahrehbar 2020[a] | 2 | 5 | 0 | 5 | 4.5% | 5.00 [0.30 , 83.69] |
| Park 2021a[a] | 7 | 12 | 2 | 12 | 17.9% | 3.50 [0.91 , 13.53] |
| Park 2023[a] | 11 | 12 | 3 | 12 | 26.9% | 3.67 [1.36 , 9.91] |
| Xie 2024[b] | 3 | 4 | 1 | 4 | 9.0% | 3.00 [0.50 , 17.95] |
| **Subtotal** | | **38** | | **38** | **62.7%** | **4.14 [2.07 , 8.28]** |
| Total events: | 28 | | 6 | | | |
| Test for overall effect: Z = 4.02 (P < 0.0001) | | | | | | |
| Heterogeneity: Chi² = 0.78, df = 4 (P = 0.94); I² = 0% | | | | | | |
| **6.24.2 Rats** | | | | | | |
| El-Derany 2021[a] | 3 | 5 | 1 | 5 | 9.0% | 3.00 [0.45 , 19.93] |
| Tang 2024[c] | 6 | 8 | 2 | 4 | 23.9% | 1.50 [0.52 , 4.32] |
| Zhang 2024[b] | 5 | 6 | 0 | 6 | 4.5% | 11.00 [0.74 , 163.49] |
| **Subtotal** | | **19** | | **15** | **37.3%** | **3.00 [1.21 , 7.42]** |
| Total events: | 14 | | 3 | | | |
| Test for overall effect: Z = 2.38 (P = 0.02) | | | | | | |
| Heterogeneity: Chi² = 2.54, df = 2 (P = 0.28); I² = 21% | | | | | | |
| **Total** | | **57** | | **53** | **100.0%** | **3.72 [2.14 , 6.44]** |
| Total events: | 42 | | 9 | | | |
| Test for overall effect: Z = 4.68 (P < 0.00001) | | | | | | |
| Test for subgroup differences: Chi² = 0.31, df = 1 (P = 0.58), I² = 0% | | | | | | |
| Heterogeneity: Chi² = 4.24, df = 7 (P = 0.75); I² = 0% | | | | | | |

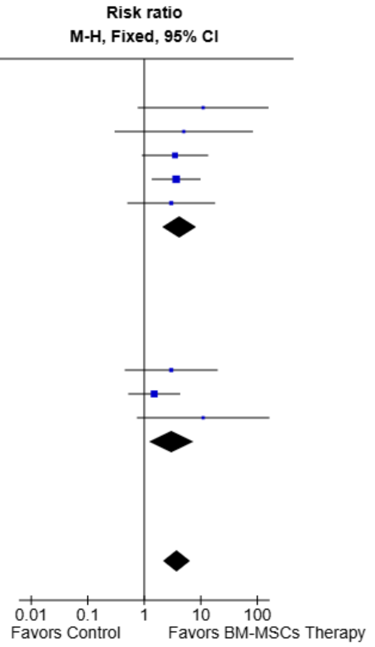


**Footnotes**

[a]BM-MSCs Transplantation

[b]BM-MSCs Secretome

[c]BM-MSCs Secretome and mIR-21 Exosomes

**Supplemental Fig. 78. Subgroup analysis of BM-MSC Therapy on pregnancy occurrence by animal model.**

| Study or Subgroup | BM-MSCs Therapy Events | BM-MSCs Therapy Total | Control Events | Control Total | Weight | Risk ratio M-H, Fixed, 95% CI |
|---|---|---|---|---|---|---|
| **6.25.1 Autoimmune** | | | | | | |
| Xie 2024[a] | 3 | 4 | 1 | 4 | 9.0% | 3.00 [0.50 , 17.95] |
| **Subtotal** | | **4** | | **4** | **9.0%** | **3.00 [0.50 , 17.95]** |
| Total events: | 3 | | 1 | | | |
| Test for overall effect: Z = 1.20 (P = 0.23) | | | | | | |
| Heterogeneity: Not applicable | | | | | | |
| **6.25.2 Chemotherapy (Cisplatin)** | | | | | | |
| Tang 2024[b] | 6 | 8 | 2 | 4 | 23.9% | 1.50 [0.52 , 4.32] |
| **Subtotal** | | **8** | | **4** | **23.9%** | **1.50 [0.52 , 4.32]** |
| Total events: | 6 | | 2 | | | |
| Test for overall effect: Z = 0.75 (P = 0.45) | | | | | | |
| Heterogeneity: Not applicable | | | | | | |
| **6.25.3 Chemotherapy (Cyclophosphamide)** | | | | | | |
| Badawy 2017[c] | 5 | 5 | 0 | 5 | 4.5% | 11.00 [0.77 , 158.01] |
| Bahrehbar 2020[c] | 2 | 5 | 0 | 5 | 4.5% | 5.00 [0.30 , 83.69] |
| Park 2021a[c] | 7 | 12 | 2 | 12 | 17.9% | 3.50 [0.91 , 13.53] |
| Park 2023[c] | 11 | 12 | 3 | 12 | 26.9% | 3.67 [1.36 , 9.91] |
| Zhang 2024[a] | 5 | 6 | 0 | 6 | 4.5% | 11.00 [0.74 , 163.49] |
| **Subtotal** | | **40** | | **40** | **58.2%** | **4.85 [2.35 , 10.01]** |
| Total events: | 30 | | 5 | | | |
| Test for overall effect: Z = 4.26 (P < 0.0001) | | | | | | |
| Heterogeneity: Chi² = 1.24, df = 4 (P = 0.87); I² = 0% | | | | | | |
| **6.25.4 Radiotherapy** | | | | | | |
| El-Derany 2021[c] | 3 | 5 | 1 | 5 | 9.0% | 3.00 [0.45 , 19.93] |
| **Subtotal** | | **5** | | **5** | **9.0%** | **3.00 [0.45 , 19.93]** |
| Total events: | 3 | | 1 | | | |
| Test for overall effect: Z = 1.14 (P = 0.26) | | | | | | |
| Heterogeneity: Not applicable | | | | | | |
| **Total** | | **57** | | **53** | **100.0%** | **3.72 [2.14 , 6.44]** |
| Total events: | 42 | | 9 | | | |
| Test for overall effect: Z = 4.68 (P < 0.00001) | | | | | | |
| Test for subgroup differences: Chi² = 3.23, df = 3 (P = 0.36), I² = 7.1% | | | | | | |
| Heterogeneity: Chi² = 4.24, df = 7 (P = 0.75); I² = 0% | | | | | | |

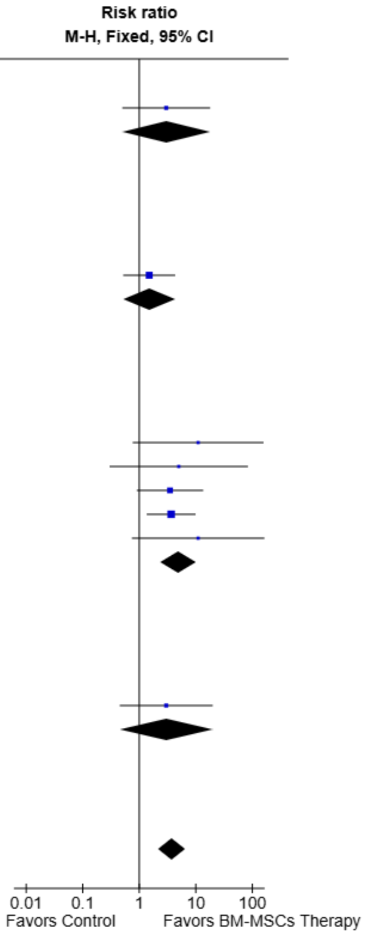


**Footnotes**

[a]BM-MSCs Secretome

[b]BM-MSCs Secretome and mIR-21 Exosomes

[c]BM-MSCs Transplantation

**Supplemental Fig. 79. Subgroup analysis of BM-MSC Therapy on pregnancy occurrence by POI induction model.**

| Study or Subgroup | BM-MSCs Therapy Mean | SD | Total | Control Mean | SD | Total | Weight | Std. mean difference IV, Fixed, 95% CI |
|---|---|---|---|---|---|---|---|---|
| **6.26.1 Intraovarian** | | | | | | | | |
| Park 2021a[a] | 5.67 | 3.08 | 12 | 0.92 | 2.15 | 12 | 29.7% | 1.73 [0.76 , 2.69] |
| **Subtotal** | | | **12** | | | **12** | **29.7%** | **1.73 [0.76 , 2.69]** |
| Test for overall effect: Z = 3.52 (P = 0.0004) | | | | | | | | |
| Heterogeneity: Not applicable | | | | | | | | |
| **6.26.2 Intraperitoneal** | | | | | | | | |
| Tang 2024[b] | 4.48 | 1.36 | 8 | 0.65 | 0.59 | 4 | 7.7% | 2.99 [1.10 , 4.88] |
| **Subtotal** | | | **8** | | | **4** | **7.7%** | **2.99 [1.10 , 4.88]** |
| Test for overall effect: Z = 3.10 (P = 0.002) | | | | | | | | |
| Heterogeneity: Not applicable | | | | | | | | |
| **6.26.3 Intravenous** | | | | | | | | |
| El-Derany 2021[a] | 5 | 2.95 | 5 | 0 | 2.53 | 5 | 11.5% | 1.64 [0.10 , 3.19] |
| Park 2023[a] | 4.5 | 2.95 | 12 | 1.25 | 2.53 | 12 | 35.9% | 1.14 [0.27 , 2.02] |
| Zhang 2024[c] | 5.06 | 4.43 | 6 | 0 | 0.53 | 6 | 15.2% | 1.48 [0.14 , 2.82] |
| **Subtotal** | | | **23** | | | **23** | **62.6%** | **1.32 [0.65 , 1.98]** |
| Test for overall effect: Z = 3.90 (P < 0.0001) | | | | | | | | |
| Heterogeneity: Chi² = 0.38, df = 2 (P = 0.83); I² = 0% | | | | | | | | |
| **Total** | | | **43** | | | **39** | **100.0%** | **1.57 [1.04 , 2.09]** |

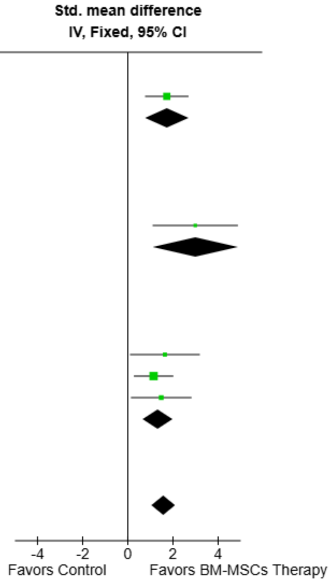


Test for overall effect: Z = 5.86 (P < 0.00001)
Test for subgroup differences: Chi² = 2.83, df = 2 (P = 0.24), I² = 29.4%
Heterogeneity: Chi² = 3.21, df = 4 (P = 0.52); I² = 0%

**Footnotes**
[a]BM-MSCs Transplantation
[b]BM-MSCs Secretome and mIR-21 Exosomes
[c]BM-MSCs Secretome

**Supplemental Fig. 80. Subgroup analysis of BM-MSC Therapy on number of offspring by route of administration.**

| Study or Subgroup | BM-MSCs Therapy Mean | SD | Total | Control Mean | SD | Total | Weight | Std. mean difference IV, Fixed, 95% CI |
|---|---|---|---|---|---|---|---|---|
| **6.27.1 Mice** | | | | | | | | |
| Park 2021a[a] | 5.67 | 3.08 | 12 | 0.92 | 2.15 | 12 | 29.7% | 1.73 [0.76 , 2.69] |
| Park 2023[a] | 4.5 | 2.95 | 12 | 1.25 | 2.53 | 12 | 35.9% | 1.14 [0.27 , 2.02] |
| **Subtotal** | | | **24** | | | **24** | **65.6%** | **1.41 [0.76 , 2.05]** |
| Test for overall effect: Z = 4.26 (P < 0.0001) | | | | | | | | |
| Heterogeneity: Chi² = 0.78, df = 1 (P = 0.38); I² = 0% | | | | | | | | |
| **6.27.2 Rats** | | | | | | | | |
| El-Derany 2021[a] | 5 | 2.95 | 5 | 0 | 2.53 | 5 | 11.5% | 1.64 [0.10 , 3.19] |
| Tang 2024[b] | 4.48 | 1.36 | 8 | 0.65 | 0.59 | 4 | 7.7% | 2.99 [1.10 , 4.88] |
| Zhang 2024[c] | 5.06 | 4.43 | 6 | 0 | 0.53 | 6 | 15.2% | 1.48 [0.14 , 2.82] |
| **Subtotal** | | | **19** | | | **15** | **34.4%** | **1.87 [0.98 , 2.77]** |
| Test for overall effect: Z = 4.11 (P < 0.0001) | | | | | | | | |
| Heterogeneity: Chi² = 1.75, df = 2 (P = 0.42); I² = 0% | | | | | | | | |
| **Total** | | | **43** | | | **39** | **100.0%** | **1.57 [1.04 , 2.09]** |

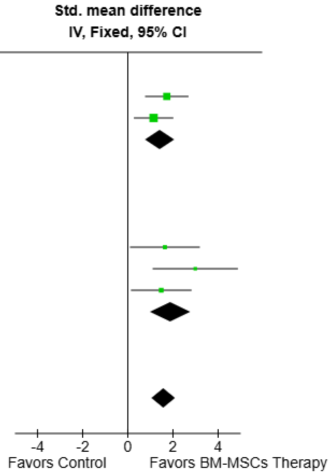


Test for overall effect: Z = 5.86 (P < 0.00001)
Test for subgroup differences: Chi² = 0.68, df = 1 (P = 0.41), I² = 0%
Heterogeneity: Chi² = 3.21, df = 4 (P = 0.52); I² = 0%

**Footnotes**
[a]BM-MSCs Transplantation
[b]BM-MSCs Exosomes and mIR-21 Exosomes
[c]BM-MSCs Secretome

**Supplemental Fig. 81. Subgroup analysis of BM-MSC Therapy on number of offspring by animal model.**

| Study or Subgroup | BM-MSCs Therapy Mean | SD | Total | Control Mean | SD | Total | Weight | Std. mean difference IV, Fixed, 95% CI |
|---|---|---|---|---|---|---|---|---|
| **6.28.1 Chemotherapy (Cisplatin)** | | | | | | | | |
| Tang 2024[a] | 4.48 | 1.36 | 8 | 0.65 | 0.59 | 4 | 7.7% | 2.99 [1.10 , 4.88] |
| **Subtotal** | | | **8** | | | **4** | **7.7%** | **2.99 [1.10 , 4.88]** |
| Test for overall effect: Z = 3.10 (P = 0.002) | | | | | | | | |
| Heterogeneity: Not applicable | | | | | | | | |
| **6.28.2 Chemotherapy (Cyclophosphamide)** | | | | | | | | |
| Park 2021a[b] | 5.67 | 3.08 | 12 | 0.92 | 2.15 | 12 | 29.7% | 1.73 [0.76 , 2.69] |
| Park 2023[b] | 4.5 | 2.95 | 12 | 1.25 | 2.53 | 12 | 35.9% | 1.14 [0.27 , 2.02] |
| Zhang 2024[c] | 5.06 | 4.43 | 6 | 0 | 0.53 | 6 | 15.2% | 1.48 [0.14 , 2.82] |
| **Subtotal** | | | **30** | | | **30** | **80.8%** | **1.42 [0.84 , 2.00]** |
| Test for overall effect: Z = 4.78 (P < 0.00001) | | | | | | | | |
| Heterogeneity: Chi² = 0.79, df = 2 (P = 0.67); I² = 0% | | | | | | | | |
| **6.28.3 Radiotherapy** | | | | | | | | |
| El-Derany 2021[b] | 5 | 2.95 | 5 | 0 | 2.53 | 5 | 11.5% | 1.64 [0.10 , 3.19] |
| **Subtotal** | | | **5** | | | **5** | **11.5%** | **1.64 [0.10 , 3.19]** |
| Test for overall effect: Z = 2.08 (P = 0.04) | | | | | | | | |
| Heterogeneity: Not applicable | | | | | | | | |
| **Total** | | | **43** | | | **39** | **100.0%** | **1.57 [1.04 , 2.09]** |

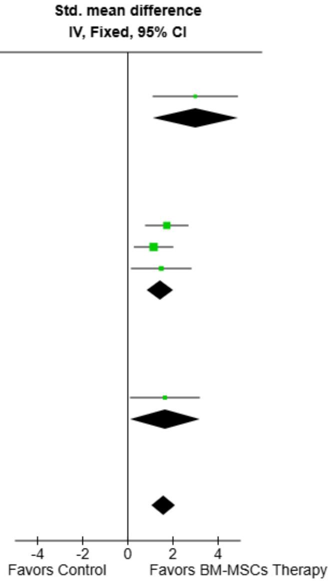


Test for overall effect: Z = 5.86 (P < 0.00001)
Test for subgroup differences: Chi² = 2.43, df = 2 (P = 0.30), I² = 17.6%
Heterogeneity: Chi² = 3.21, df = 4 (P = 0.52); I² = 0%

**Footnotes**
[a]BM-MSCs Secretome and miR-21 Exosomes
[b]BM-MSCs Transplantation
[c]BM-MSCs Secretome

**Supplemental Fig. 82. Subgroup analysis of BM-MSC Therapy on number of offspring by POI induction model.**

### Prisma 2020 Checklist

| Section and Topic | Item # | Checklist item | Location where item is reported |
|---|---|---|---|
| TITLE | | | |
| Title | 1 | Identify the report as a systematic review. | Title |
| Abstract | 2 | See the PRISMA 2020 for Abstracts checklist. | Abstract |
| INTRODUCTION | | | |
| Rationale | 3 | Describe the rationale for the review in the context of existing knowledge | 1. Introduction section |
| Objectives | 4 | Provide an explicit statement of the objective(s) or question(s) the review addresses. | End of 1. Introduction |
| METHODS | | | |
| Eligibility criteria | 5 | Specify the inclusion and exclusion criteria for the review and how studies were grouped for the syntheses. | 2.1 Eligibility Criteria |
| Information sources | 6 | Specify all databases, registers, websites, organisations, reference lists and other sources searched or consulted to identify studies. Specify the date when each source was last searched or consulted. | 2.2 Search strategy data extraction |
| Search strategy | 7 | Present the full search strategies for all databases, registers and websites, including any filters and limits used. | Table S1 (Supplementary material) |
| Selection process | 8 | Specify the methods used to decide whether a study met the inclusion criteria of the review, including how many reviewers screened each record and each report retrieved, whether they worked independently, and if applicable, details of automation tools used in the process. | 2.2 Search strategy data extraction |
| Data collection process | 9 | Specify the methods used to collect data from reports, including how many reviewers collected data from each report, whether they worked independently, any processes for obtaining or confirming data from study investigators, and if applicable, details of automation tools used in the process. | 2.2 Search strategy data extraction |
| Data items | 10a | List and define all outcomes for which data were sought. Specify whether all results that were compatible with each outcome domain in each study were sought (e.g. for all measures, time points, analyses), and if not, the methods used to decide which results to collect. | 2.3 Endpoints and subgroup analyses |
| | 10b | List and define all other variables for which data were sought (e.g. participant and intervention characteristics, funding sources). Describe any assumptions made about any missing or unclear information. | Table 1; 2.2 (data extraction). |
| Study risk of bias assessment | 11 | Specify the methods used to assess risk of bias in the included studies, including details of the tool(s) used, how many reviewers assessed each study and whether they worked independently, and if applicable, details of automation tools used in the process. | 2.4 Quality assessment (SYRCLE tool) |
| Effect measures | 12 | Specify for each outcome the effect measure(s) (e.g. risk ratio, mean difference) used in the synthesis or presentation of results. | 2.6 Statistical Analysis |
| Synthesis methods | 13a | Describe the processes used to decide which studies were eligible for each synthesis (e.g. tabulating the study intervention characteristics and comparing against the planned groups for each synthesis (item #5)). | 2.3 Endpoints and subgroup analyses |
| | 13b | Describe any methods required to prepare the data for presentation or synthesis, such as handling of missing summary statistics, or data conversions. | 2.6 Statistical Analysis; 2.2 (graph extraction) |
| | 13c | Describe any methods used to tabulate or visually display results of individual studies and syntheses. | 2.6 Statistical Analysis & summary tables/forest plots |

| | | | |
|---|---|---|---|
| | 13d | Describe any methods used to synthesize results and provide a rationale for the choice(s). If meta-analysis was performed, describe the model(s), method(s) to identify the presence and extent of statistical heterogeneity, and software package(s) used. | 2.5 Assessment of heterogeneity & 2.6 Statistical Analysis |
| | 13e | Describe any methods used to explore possible causes of heterogeneity among study results (e.g. subgroup analysis, meta-regression). | 2.5 Assessment of heterogeneity |
| | 13f | Describe any sensitivity analyses conducted to assess robustness of the synthesized results. | 2.5 Assessment of heterogeneity |
| Reporting bias assessment | 14 | Describe any methods used to assess risk of bias due to missing results in a synthesis (arising from reporting biases). | 2..4 Quality assessment (funnel plots) |
| Certainty assessment | 15 | Describe any methods used to assess certainty (or confidence) in the body of evidence for an outcome. | Not performed: GRADE not applicable to preclinical evidence; addressed via SYRCLE RoB, heterogeneity and publication-bias analyses |
| RESULTS | | | |
| Study selection | 16a | Describe the results of the search and selection process, from the number of records identified in the search to the number of studies included in the review, ideally using a flow diagram. | Results: 3.1 & Fig. 1 |
| | 16b | Cite studies that might appear to meet the inclusion criteria, but which were excluded, and explain why they were excluded. | Supplementary Material Excluded studies Table S2 |
| Study characteristics | 17 | Cite each included study and present its characteristics. | Results: 3.1 & Table 1 |
| Risk of bias in studies | 18 | Present assessments of risk of bias for each included study. | Results: 3.3 & Table 3 |
| Results of individual studies | 19 | For all outcomes, present, for each study: (a) summary statistics for each group (where appropriate) and (b) an effect estimate and its precision (e.g. confidence/credible interval), ideally using structured tables or plots. | Forest plots (Figures 2–7 & Supplementary forest plots) |
| Results of syntheses | 20a | For each synthesis, briefly summarise the characteristics and risk of bias among contributing studies. | Results: 3.2 |
| | 20b | Present results of all statistical syntheses conducted. If meta-analysis was done, present for each the summary estimate and its precision (e.g. confidence/credible interval) and measures of statistical heterogeneity. If comparing groups, describe the direction of the effect. | Results: 3.2 (Sections 3.2.1-3.2.2)Table 2 |
| | 20c | Present results of all investigations of possible causes of heterogeneity among study results. | Results: 3.2.3 & Supplemental figures |
| | 20d | Present results of all sensitivity analyses conducted to assess the robustness of the synthesized results. | Results: 3.2.3 |
| [1] Reporting biases | 21 | Present assessments of risk of bias due to missing results (arising from reporting biases) for each synthesis assessed. | Results: 3.2.3 Supplementary material (Funnel plots) |

| | | | |
|---|---|---|---|
| Certainty of evidence | 22 | Present assessments of certainty (or confidence) in the body of evidence for each outcome assessed. | Not performed: GRADE not applicable to preclinical evidence; addressed via SYRCLE RoB, heterogeneity and publication-bias analyses |
| DISCUSSION | | | |
| Discussion | 23a | Provide a general interpretation of the results in the context of other evidence. | Discussion: 4.1, 4.2 |
| | 23b | Discuss any limitations of the evidence included in the review. | Discussion: 4.4 |
| | 23c | Discuss any limitations of the review processes used. | Discussion: 4.4 |
| | 23d | Discuss implications of the results for practice, policy, and future research. | Discussion: 4.4 |
| OTHER INFORMATION | | | |
| Registration and protocol | 24a | Provide registration information for the review, including register name and registration number, or state that the review was not registered. | Methods (PROSPERO CRD42023449053) |
| | 24b | Indicate where the review protocol can be accessed, or state that a protocol was not prepared. | Methods |
| | 24c | Describe and explain any amendments to information provided at registration or in the protocol. | No amendments were made |
| Support | 25 | Describe sources of financial or non-financial support for the review, and the role of the funders or sponsors in the review. | Declarations: Funding |
| Competing interests | 26 | Declare any competing interests of review authors. | Declarations: Competing interests |
| Availability of data, code and other materials | 27 | Report which of the following are publicly available and where they can be found: template data collection forms; data extracted from included studies; data used for all analyses; analytic code; any other materials used in the review. | Declarations: Availability of data and material |

***From:*** **Page MJ, McKenzie JE, Bossuyt PM, Boutron I, Hoffmann TC, Mulrow CD, et al. The PRISMA 2020 statement: an updated guideline for reporting systematic reviews. BMJ 2021;372:n71. doi: 10.1136/bmj.n71. This work is licensed under CC BY 4.0. To view a copy of this license, visit https://creativecommons.org/licenses/by/4.0/**

**https://www.bmj.com/content/372/bmj.n160**
